# Genome-wide association study and transcriptomics reveal the genetic architecture of alkalinity tolerance in *Arabidopsis thaliana*

**DOI:** 10.64898/2026.03.01.708912

**Authors:** Neelam Jangir, Rishabh Kumar, Surbhi Vilas Tajane, Devanshu Verma, Rabisankar Mandi, Sucharita Dey, Ayan Sadhukhan

## Abstract

Alkalinity stress significantly restricts global plant productivity, yet the genetic basis for plant tolerance remains largely uncharacterized. In this study, a genome-wide association study was performed using 218 diverse natural *Arabidopsis thaliana* ecotypes to identify the top 73 SNPs associated with alkalinity tolerance, measured by relative root length in hydroponic growth media containing NaHCO_3 a_t pH 8.0. Prominent association peaks were localized near genes involved in lipid metabolism (*GGL20*), protein degradation (AT3G17570), and vesicle-mediated protein sorting (*VPS13B* and AT5G57210). Expression level and protein polymorphisms in these genes were associated with alkalinity tolerance. T-DNA mutants of *GGL20*, AT3G17570, and the chromatin-modifying gene *AFR1* showed alkaline hypersensitivity, reduced root length, iron content, and rosette size, and elevated hydrogen peroxide. Conversely, mutants of the DNA repair gene *ETG1* exhibited greater tolerance than wild type in hydroponics, solid media, and soil assays, confirming their role in alkalinity tolerance. Transcriptome and network analyses revealed that alkalinity responses significantly overlap with iron deficiency pathways, identifying hub genes involved in ribosome assembly and translation control. These findings provide a comprehensive map of the genetic and transcriptional landscape of alkalinity adaptation and offer promising candidate genes for engineering crops resilient to alkaline soil conditions.

## 1 Introduction

Soil pH, shaped by geology, weathering, environment, and vegetation, profoundly influences plant fitness, growth, species distribution, and agricultural productivity (Tsai and Schmidt, 2021). As a measure of H⁺ concentration in soil solution, pH dictates acidity or alkalinity and acts as the “master soil variable” controlling microbial activity, nutrient availability, and crop development (Zhang et al. 2019). High pH soils cover 30% of Earth’s surface, are prevalent in arid and semiarid regions, and are characterized by calcareous soils with pH 7.5–8.5 and bicarbonate (HCO_3⁻_) levels of 5–35 nmol L⁻¹ (Loeppert and Suarez, 1996). HCO_3⁻_ and carbonate (CO_3²_⁻) dominate as excess anions in such soils, buffering pH and limiting nutrient solubility. Most trace elements become less available as pH rises; a one-unit drop in pH increases metal solubility tenfold.

Alkalinity, high pH, and HCO_3⁻_ stresses are interlinked (Cao et al. 2022; Yang et al. 2024). Plants, being sessile, tightly regulate cytoplasmic and organelle pH, extending to the root apoplast, which governs cell wall properties and mineral availability (Martinière et al. 2018). High pH disrupts root membrane potential, ion uptake, and plant physiology (Busoms et al. 2023; Shi and Wang, 2005). Alkali stress from NaHCO₃ and Na₂CO₃ hydrolysis exacerbates this, inducing ionic toxicity, osmotic stress, and pH disruption, which destabilize cell pH, membrane integrity, photosynthesis, and root vitality (Zhang et al. 2017; Kaiwen et al. 2020). It triggers ion imbalance, antioxidant inhibition, osmotic imbalance, accumulation of reactive oxygen species (ROS)/malondialdehyde (MDA), leading to growth suppression (Fang et al. 2021).

External pH shifts trigger rapid transcriptional responses, indicating the presence of dedicated sensing mechanisms conserved across kingdoms (Lager et al. 2010; Tsai and Schmidt, 2021; Schmidt, 2022). The primary pH sensor complexes in roots are the root meristem growth factor (RGF)–RGF receptor (RGFR)–somatic embryogenesis receptor kinase (SERK), and plant elicitor peptide (Pep)–Pep receptor (PEPR)–BAK1/SERK3 (BAK) systems. RGF binds to RGFR and, under acidic conditions, recruits the co-receptor SERK, facilitating root meristem growth. Whereas, under alkaline conditions, the Pep ligand binds to PEPR and forms a complex with BAK to activate defense signaling (Liu et al. 2022). Plants tolerate high pH mainly by maintaining internal pH and ion homeostasis and improving nutrient acquisition under alkaline conditions (Li et al. 2023; Yang et al. 2024). Plants also adjust the expression and activity of ion transporters for Na⁺, K⁺, and H⁺, and employ signaling networks involving calcium, ROS, and protein kinases to coordinate responses to saline-alkali stress. At the cellular and organ level, they remodel cell walls and modify root system architecture to optimize access to more favorable nutrient pools (Li et al. 2023; Yang et al. 2024). A genome-wide association study (GWAS) in sorghum pinpointed a key negative regulator, *alkaline tolerance 1* (*AT1*) gene, encoding a G protein γ subunit. *AT1* modulates ROS homeostasis by inhibiting phosphorylation of plasma membrane intrinsic protein 2 (PIP2) aquaporins, promoting hydrogen peroxide (H_2O2)_ export under alkaline stress. *AT1* knockouts showed enhanced alkalinity tolerance and yield in sorghum, maize, rice, and foxtail millet, highlighting its potential for crop engineering (Zhang et al. 2023). A recent study in rice revealed Na⁺/K⁺ and ROS loci under alkalinity stress via GWAS and haplotype analysis (Ganapati et al. 2025). HCO_3⁻_ stress in alkalinity upregulates *slowly activating anion conductance homologue 3* (*SLAH3*) in *Glycine soja*. *GsSLAH3* overexpression in *Arabidopsis thaliana* boosts HCO_3⁻_ tolerance (Duan et al. 2018). Similarly, *ethylene response factor 6* (*GsERF6*) and *boron transporter 2* (*BOR2*) respond to HCO_3⁻_ (Yu et al. 2016; Duan et al. 2018). Loss-of-function mutants of *DNAJ homologue 3* (*J3*) show reduced plasma membrane (PM) H⁺-ATPase activity and alkalinity hypersensitivity (Yang et al. 2010). Under alkalinity stress, SOS3-like calcium-binding protein 3 (SCaBP3) dissociates from PM H⁺-ATPases upon Ca²⁺ sensing, activating *plasma membrane proton ATPase 2* (*AHA2*) via decreased *SCaBP3*/SOS2-like protein kinase 5 (*PKS5*) phosphorylation at Ser-931 and 14-3-3 transcription factor (TF) binding (Yang et al. 2019) In wheat, alkaline stress induces a Ca²⁺-binding *C-terminal centrin-like domain containing protein 1* (*CCD1*), which interacts with *small auxin up RNA 215* (*SAUR215*) to inhibit *D-clade type 2C protein phosphatases*. This prevents Thr947 dephosphorylation of PM H⁺-ATPase *TaHA2*, enhancing its activity (Cui et al. 2023), which in turn acidifies the rhizosphere and apoplast, increasing solubility and uptake of nutrients such as iron (Fe) and phosphorus (Khan et al. 2019; Li et al. 2023).

*A. thaliana*, a model in plant research, thrives optimally in acidic soils (pH 5.0–7.0) but exhibits natural genetic variation across a broader pH range of 5.0–8.5, reflecting ecotypic adaptations to diverse native environments. When grown on alkaline substrates, ecotypes from high-pH backgrounds display superior growth compared to those from acidic soils, demonstrating that plant fitness correlates strongly with native soil pH and CO ^2^⁻/HCO ⁻ content. Natural variation in large *A. thaliana* accession panels is exploited in GWAS to map phenotypes to candidate genes underlying complex traits, such as abiotic stress tolerance (Marik et al. 2026). Integrative GWAS and transcriptomics can effectively unravel high-pH tolerance genes in plants: GWAS identifies candidate loci, while transcriptomics validates their regulatory roles via differential expression under stress (Kobayashi et al. 2016; Butardo et al. 2017). A GWAS under low pH conditions used Central Asian *A. thaliana* ecotypes to identify a promoter polymorphism in *AtMATE* that lowers expression and Al tolerance in acid-soil natives, linking this variation to the exudation of organic acids (Nakano et al. 2020). GWAS in Iberian *A. thaliana* subpopulations (demes) under CO ^2^⁻ stress identified loci for alkalinity tolerance, Ca exclusion, and Fe acquisition. Similarly, another GWAS identified Fe homeostasis genes for chlorosis resistance in *A. thaliana* under high-pH conditions (Busoms et al. 2023; Casellas et al. 2023). Root carbonic anhydrase *βCA4.1* was found critical for HCO_3⁻_ tolerance in *A. thaliana* by mediating early membrane hyperpolarization and coordinating the upregulation of genes essential for Fe acquisition, anion transport, and CO_2/_water diffusion. Interestingly, *βCA4.1* was induced in a CO ^2^⁻-tolerant deme and downregulated in a sensitive one (Pérez-Martin et al. 2024). A transcriptome study unravelled thousands of pH-responsive genes (H⁺-ATPases, *NRT2*, *AMT1*, etc.) under pH ranging from 4.5 to 7.5 (Bailey et al. 2023). Genes involved in the jasmonate and salicylate (SA) pathways, along with glucosinolate biosynthesis, cell cycle, and carbohydrate metabolism-related genes, were rapidly induced in the shoots of CO ^2^⁻-tolerant *A. thaliana* demes, whereas Fe deficiency-responsive genes were induced in the sensitive ones (Pérez-Martin et al. 2021). A proteomics study showed extensive pH-dependent phosphosite remodeling on H⁺-ATPases, nutrient transporters (*NRT2.1*, *AMT1*, *PDR7/8*), and endocytic regulators (TPLATE complex), establishing phospho-networks which dynamically recalibrate ion uptake, H⁺ pumping, and growth-defense trade-offs (Jain et al. 2024). Another metabolome-based study of *A. thaliana* suspension cells revealed distinct HCO_3⁻_ signatures, particularly in carbon metabolism and flavonoid biosynthesis pathways (Misra et al. 2016).

Despite advances in this field, the genetic architecture underlying plants’ adaptive strategies for tolerance to high pH/alkalinity stress remains poorly understood. In this study, we employed GWAS to unravel hitherto unknown molecular players for HCO_3⁻_-induced high-pH stress tolerance under hydroponic growth conditions at pH 8, a condition that reflects moderate but physiologically relevant alkalinity stress. Using a diverse panel of 218 natural *A. thaliana* ecotypes, we performed GWA mapping to detect loci associated with phenotypic variation under alkaline conditions, followed by transcriptome profiling to characterize global gene expression changes induced by alkaline stress. Additionally, to identify core stress-responsive genes and pathways and to resolve the key regulatory mechanisms, we integrated GWAS and transcriptomic data via network analysis. Using reverse genetics with T-DNA insertional mutants from candidate genes identified via GWAS, we validated their functional roles in conferring tolerance to alkalinity stress. Collectively, this integrative approach provides an inclusive interpretation of how plants perceive and respond to alkalinity at the genetic and transcriptional levels. By unravelling the molecular mechanisms underlying adaptive responses of *A. thaliana*, our study advances our understanding of alkalinity stress biology and reveals candidate genes and regulatory mechanisms that may be used to improve crop yield on alkaline soils.

## 2 Materials and Methods

### 2.1 Mapping populations and phenotyping conditions

A diversity panel of 218 worldwide *A. thaliana* ecotypes was used for the current GWA study (Table S1). Seeds were sourced from the RIKEN Bioresource Research Centre, Japan, and the Nottingham Arabidopsis Stock Centre (NASC), United Kingdom. Geographic coordinates for the original collection sites of *A. thaliana* ecotypes were sourced from the 1001 Genomes database. Worldwide soil pH data were downloaded from OpenLandMap (https://s3.openlandmap.org/). The dataset included soil pH measurements in water at a depth of 0–10 cm (surface layer) with a spatial resolution of 250 m, covering the timeframe from 1950 through 2017. The pH scale ranged between 3 (acidic) and 10 (alkaline). Soil organic carbon stock in tons per hectare was predicted using the global compilation of soil ground observations (Hengl et at. 2017). Soil CaCO_3 d_ata were extracted from the WoSIS database (https://isric.org/explore/wosis). The coordinates of the *A. thaliana* ecotypes were mapped onto the global pH data using QGIS 3.4 Madeira long-term release software (https://qgis.org/). The seeds were freshly multiplied using the single-seed descent method. The culture room was maintained at 22 ± 3°C and 60 ± 5% humidity. SMD-type T8 LED lamps provided 40 μmol m^−2^ s^−1^ of photosynthetically active radiation in 12-hour illumination and 12-hour dark cycles. The seeds were imbibed in distilled water at 4°C for 3 days to achieve synchronized germination on N50 nylon meshes, which were floated on a hydroponic growth medium using photographic mounts and Styrofoam, as described previously (Sadhukhan et al. 2017; Marik et al. 2026). Plants were grown in a control growth medium of ¼-Hoagland’s solution (Hoagland & Arnon, 1938), pH 5.8, and an alkalinity stress-inducing high-pH growth medium consisting of ¼-Hoagland’s solution supplemented with 1 mM NaHCO_3,_ pH 8.0, adjusted with 7 M KOH. The alkalinity stress tolerance was determined as the ratio of root length in alkaline media to that in the control medium, expressed as a percentage, termed the relative root length (RRL).

### 2.2 GWAS

Genome-wide 105,856 single-nucleotide polymorphisms (SNPs) for 218 *A. thaliana* ecotypes were extracted from publicly available resources (Atwell et al. 2010; Cao et al. 2011; Horton et al. 2012) and used for association mapping using a mixed linear model (MLM), accounting for the population structure and kinship between ecotypes in TASSEL ver. 5.0 (Bradbury et al. 2007). Functional enrichment analysis of the GWAS-identified genes was performed using ShinyGO 0.85.1 (https://bioinformatics.sdstate.edu/go/).

### 2.3 Polymorphism analysis

The POLYMORPH web tool was used to mine denser polymorphisms of ‘high’ and ‘moderate’ effects for an available 151 ecotypes out of the 218 used in the GWAS from the 1001 Genomes database (http://tools.1001genomes.org/polymorph/). The denser polymorphism information was used in a local association study (LAS) of local genomic regions to assess association with the alkalinity tolerance variation.

#### 2.3.1 Gene expression level polymorphism analysis

Forty ecotypes were randomly selected from the 151 that had dense promoter/UTR polymorphism data in the 1001 database and grown for 10 days in hydroponics in ¼-Hoagland’s solution, pH 5.8, and transferred to ¼-Hoagland’s solution containing 1 mM NaHCO_3,_ pH 8.0, and grown for five days. Twenty seedlings were flash-frozen in liquid nitrogen for RNA isolation, followed by cDNA synthesis and reverse transcription quantitative polymerase chain reaction (RT-qPCR) analysis of genes showing significant promoter and 5’-UTR polymorphisms (MLM or GLM, *P* < 0.05) linked to alkalinity tolerance in the LAS population, as detailed in Section 2.5 with primers from Table S2.

#### 2.3.2 Promoter *cis-*element analysis

The promoter sequences of genes showing ELP were retrieved from TAIR and divided into eight-nucleotide stretches (octamers). These promoter octamer frequencies were quantified in genes induced (log_2F_C > 1) or repressed (log_2F_C < −1) from the in-house whole-seedling transcriptome (alkalinity (1): 10-day-old plants treated for 5 days with 1 mM NaHCO_3,_ pH 8.0), a public shoot transcriptome GSE164502 (alkalinity (2): 15-day-old plants treated for 3 h and 48 h with 10 mM NaHCO_3,_ pH 8.3; Pérez-Martin et al. 2021), root transcriptomes under low pH stress (GSE18982; Lager et al. 2010; GSE30095; Iyer-Pascuzzi et al. 2011), and a root transcriptome under Fe deficiency stress (GSE16964; Li et al. 2010). The relative appearance ratio (RAR) of each octamer in the promoters of stress-perturbed genes, compared to the *A. thaliana* genome-wide gene set, was calculated using Python scripts (Yamamoto et al. 2011; Marik et al. 2026). A one-sided Fisher’s exact test was used to evaluate the statistical significance of motif enrichment or overrepresentation, testing whether octamer frequencies in stress-responsive gene promoters exceeded those in the genomic background (RAR > 2 and *P* < 0.05).

### 2.4 Reverse genetic analysis

T-DNA insertion mutants targeting GWAS-identified candidate genes were identified through the Salk T-DNA Express database (http://signal.salk.edu/cgi-bin/tdnaexpress). Seeds for these lines were obtained from the NASC. The seeds were germinated and grown for three weeks in a hydroponic system, and then each seedling was transferred to two-inch pots with vermiculite, soilrite, and garden soil in a 1:2:1 ratio, along with Arasystem (https://www.arasystem.com/). After one month, leaf samples were taken from individual plants and snap-frozen in liquid nitrogen for DNA isolation using the CTAB method. Two PCR reactions were performed using DNA from both T-DNA insertion lines and the wild-type (WT; Col-0; N60000). While the first reaction used a T-DNA left border (LB) and a right primer (RP) to bind genomic DNA, the second used a left genomic primer (LP) and RP to check for homozygosity. The SALK-recommended LB primer, SALK LBb1.3 (http://signal.salk.edu/tdnaprimers.2.html), was used (Table S2). The single-seed descent method was used to multiply PCR-confirmed homozygous lines, and the resulting seeds were used for phenotyping. We measured gene expression levels in T-DNA insertion lines relative to wild-type controls to assess the degree of knockdown caused by insertions in 5’-UTRs, introns, or promoters. This analysis involved RT-qPCR on 15-day-old hydroponically grown seedlings subjected to 5 days of alkalinity stress, following the protocol detailed in Section 2.5. Mutants were grown under the same hydroponic conditions as used in the GWAS (section 2.1), and their RRL phenotypes were compared with the WT. To determine Fe concentration, aqueous root extracts were centrifuged at 10,000 rpm for 10 minutes, then digested with 98% HNO₃ in a Multiwave GO plus microwave digestion system (Anton Paar, Gurugram, India). Root Fe contents were then analyzed by inductively coupled plasma mass spectrometry (Agilent 7850 ICP-MS) and expressed as μg g⁻¹ of fresh tissue weight (Verma et al. 2025).

### 2.5 Transcriptome and quantitative PCR analysis

Col-0 seedlings were cultivated in ¼-Hoagland’s medium (pH 5.8) for 10 days, then exposed to ¼-Hoagland’s medium containing 1 mM NaHCO_3 (_pH 8.0) for five days to induce alkalinity stress, before snap-freezing in liquid nitrogen for transcriptome analysis. Total RNA extraction employed a CTAB-based method, with subsequent cDNA library preparation using the NEBNext^®^ Ultra^TM^ II RNA Library Prep Kit (New England Biolabs, MA, USA). Libraries underwent paired-end sequencing (2×150 bp) on the Illumina NovaSeq 6000 V1.5 platform (Illumina Inc., Gurgaon, India; 300 cycles). The raw data were deposited in NCBI SRA under accession PRJNA1270151. Raw reads were processed for quality control and adapter trimming using AdapterRemoval v2.3.2 (Schubert et al. 2016), then aligned to the *A. thaliana* reference genome (GCF_000001735.4) with HISAT2 v2.2.1 (Zhang et al. 2021), as detailed in Marik et al. (2026). Gene expression was quantified via FeatureCounts v2.0.3 (Liao et al. 2013) with default settings. Differential expression analysis employed edgeR v4.1.2 (Chen et al. 2025) with Fisher’s exact test and Benjamini-Hochberg *false discovery rate* (*FDR*) correction (*FDR* < 0.05; *N* = 3 biological replicates). These results were subsequently integrated with GWAS data. The in-house data captured the effects of alkalinity on young *A. thaliana* seedlings via a NaHCO_3-_induced increase in pH. We combined our transcriptome with public transcriptome datasets, including short- and long-term shoot responses to increased alkalinity, as well as root-specific signatures of low pH and Fe deficiency, for comparative stress-effect analysis. These included a shoot transcriptome from 15-day-old plants treated for 3 h or 48 h with 10 mM NaHCO_3,_ pH 8.3 (Pérez-Martin et al. 2021), a root transcriptome under low pH stress (Lager et al. 2010; Iyer-Pascuzzi et al. 2011), and another root transcriptome under Fe deficiency (Li et al. 2010). The overlapping genes in these datasets were compared using hierarchical clustering by TBtools (Chen et al. 2020). For verifying gene expression levels, the RNA was converted to cDNA with RevertAid reverse transcriptase (Thermo) and random hexamer primers, followed by RT-qPCR using TB Green^®^ Premix Ex Taq^TM^ II (Tli RNase H Plus; Takara) and the primers listed in Table S2 on a QuantStudio 5 Real-Time PCR System (Thermo). Primers were designed using Primer3 (https://primer3.ut.ee/).

### 2.6 Network analysis

Sixty-five genes delineated by the GWAS of alkalinity stress tolerance (MLM *P* < 10^−3^) were combined with genes induced (log_2F_C > 1) or repressed (log_2F_C < −1) by alkalinity in the in-house (356 genes) and public transcriptome (3737 genes) datasets. Coexpression and protein-protein interaction networks were constructed from the combined gene set using Search Tool for the Retrieval of Interacting Genes (STRING; Szklarczyk et al. 2025) with a high-confidence cutoff (0.7) and visualized in Cytoscape version 3.10.4 (Shannon et al. 2003). Hub genes were identified using the cytoHubba plugin version 0.1 in Cytoscape (Chin *et al*. 2014). Topological analysis was performed using the Maximal Clique Centrality (MCC) algorithm, which has been shown to efficiently identify essential genes by capturing both high- and low-degree nodes within biological networks. In parallel, densely connected modules were detected using the Molecular Complex Detection (MCODE) plugin (v2.0.3). Candidate hub genes were finalized based on a combination of high MCC scores and a minimum node degree ≥ 3. Functional enrichment analysis was subsequently performed using the STRING Cytoscape plugin. Enriched biological process terms were retrieved and visualized using STRING donut charts, in which colored segments surrounding each node represent significantly enriched functional categories. To minimize redundancy among enrichment terms, an overlap cutoff of 0.5 was applied, ensuring the retention of biologically distinct processes.

## 3. Results

### 3.1 Natural variation in alkalinity stress tolerance in *Arabidopsis thaliana* under hydroponic conditions

An extensive variation in the RRL phenotype was observed by growing 218 *A. thaliana* ecotypes (Table S1) originating from diverse soil pH (4.9−7.7) (Fig. 1A) and soil CO ^2^⁻ (1.5−220 g Kg^−1^) regions (Fig. S1) for 14 days on an alkaline (pH 8.0) hydroponic medium containing NaHCO_3 (_Fig. 1B). The RRL, as a measure of alkalinity stress tolerance, varied from 12.82% to 76.45%, having a median RRL of 39.97% (*H* ^2^ = 0.97). Alkalinity stress tolerance was not correlated with the soil pH at the ecotypes’ geographic origins. However, a weak positive correlation (*R^2^*= 0.02) was observed between the alkalinity tolerance and the soil CO ^2^⁻ content at the origins (Fig. 1C). Most ecotypes highly sensitive to alkalinity (26 out of 40), showing a drastic reduction in their root length (RRL < 30%) originated from low CO ^2^⁻ (0−50 g Kg^−1^) areas, e.g., Is-0 (RRL 14.9%), Pro-0 (15.1%), Buckhorn Pass (15.2%), Ru3.1-31 (18.2%), Nz-1 (18.3%), CIBC-5 (19.7%), and Hh-0 (20.5%). On the other hand, most ecotypes (10 out of 17) highly tolerant to alkalinity (RRL > 60%) originated from high (> 100 g Kg^−1^) and moderate (50−100 g Kg^−1^) CO ^2^⁻ areas, like Sij-4 (60.9%), Gie-0 (61.3%), RRS-10 (62%), Fei-0 (RRL 63.4%), Bor-1 (63.5%), Nok-1 (64%), Castelfed-4 (66.8%), La-1 (69.8%), Gu-0 (76.1%), and Li-6 (76.5%).

**Fig. 1.**
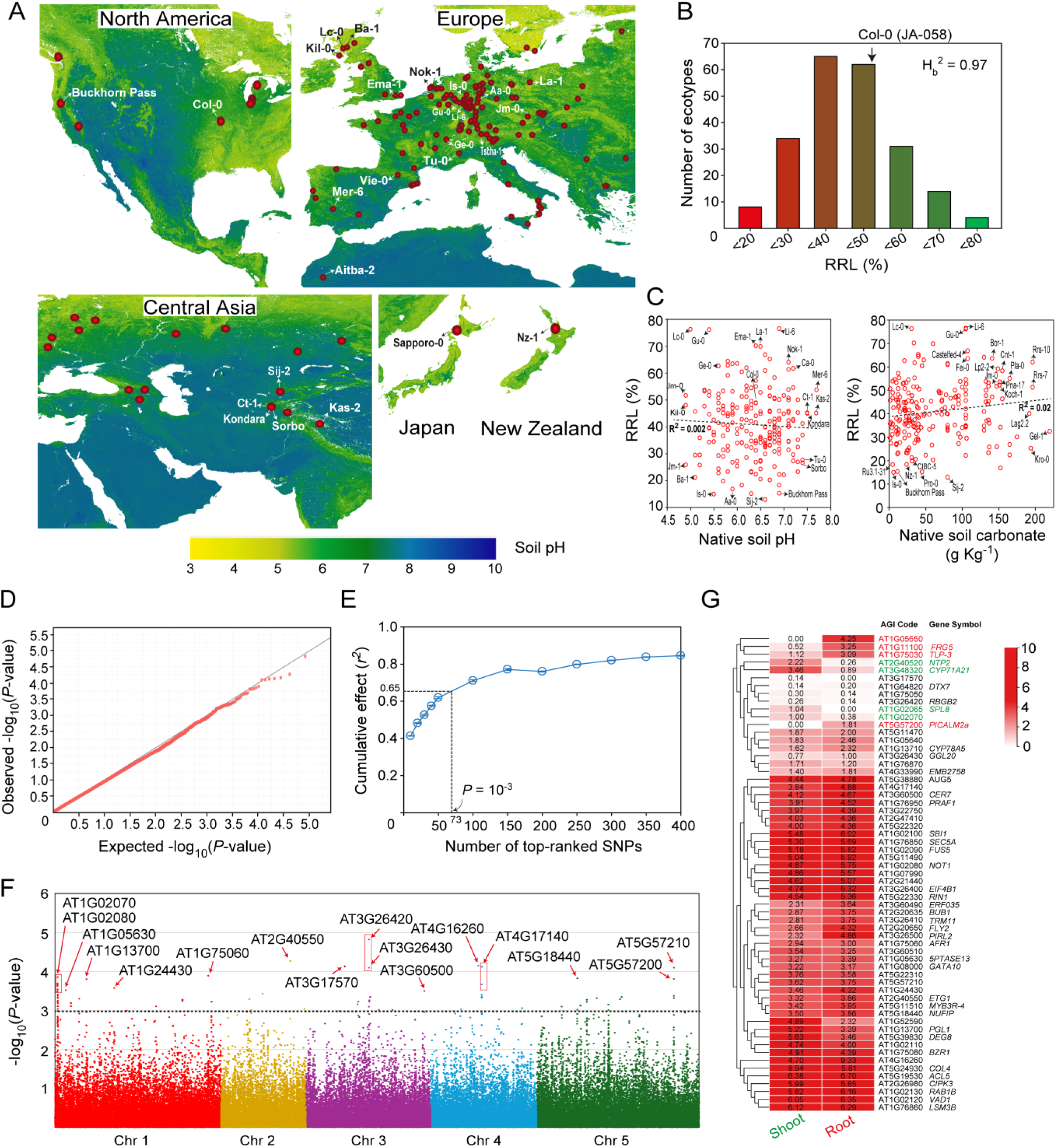
Genome-wide association study of alkalinity stress tolerance in *Arabidopsis thaliana*. Two hundred eighteen *A. thaliana* ecotypes were grown hydroponically from seed in modified ¼-Hoagland’s solution, pH 5.8 (control), or the same media supplemented with 1 mM sodium bicarbonate (NaHCO_3)_, pH 8.0, for two weeks, placed on solidified agar, and photographed. The relative root length (RRL; %) in alkaline/control media was used as a stress tolerance phenotype. The average data of 20 seedlings for each *A. thaliana* ecotype were used. (**A**) The 218 *A. thaliana* ecotypes utilized in the GWAS have their native geographic distribution displayed (https://www.google.com/maps). Parts of North America, Japan, New Zealand, and Eurasia are depicted on the maps (not to scale). The color scale shows the native soil pH at 10 cm depth (https://openlandmap.org/). The locations of the ecotypes are indicated by red dots. (**B**) The frequency distribution of RRL traits in 218 *A. thaliana* ecotypes is displayed as a histogram. The reference genotype, Columbia (Col-0), is indicated by an arrow. The ratio of genetic variance to the total of genetic and residual variance, or broad-sense heritability (H_b2_), is displayed. (**C**) A linear regression is indicated by a dotted line in the correlation plot of RRL vs soil pH of the ecotypes’ precise native geographical areas. The correlation plot also indicates the sensitive and tolerant ecotypes. The findings of a genome-wide association study (GWAS) examining single-nucleotide polymorphisms (SNPs) and alkalinity tolerance traits in 218 global *A. thaliana* ecotypes, along with population structure and kinship (mixed linear model; MLM), are shown in the next sections. (**D**) The observed versus expected *P*-value distribution of SNPs in the GWAS is displayed in a quantile-quantile (Q-Q) plot. The null distribution of *P*-values is shown by the straight line. (**E**) Ridge regression analysis results are displayed. The number of top-ranking SNPs identified by the GWAS is plotted against the mean correlation coefficient (*r²*) between the expected and observed RRLs, along with the standard deviation for 100 iterations. (**F**) A Manhattan plot of the significance of association (‒log_10*P*_-value) versus physical locations on the five *A. thaliana* chromosomes displays the GWAS results. The protein-coding genes closest to the top-ranking SNPs are indicated by arrows. (**G**) The heatmap displays a hierarchical grouping of genes associated with the most significant SNPs (MLM *P* < 10^‒3^) identified by the GWAS based on their expression levels in the root and shoot tissues of seedlings (Mergner et al. 2020). Gene expression levels, represented as transcripts per million (TPM), are depicted after correction using a logarithmic base-2 transformation. Higher expression levels are denoted by darker shades of red.

### 3.2 A GWAS identifies alkalinity tolerance genes using natural genetic variation in *Arabidopsis thaliana*

GWA mapping of natural variation in seedling tolerance to alkalinity stress revealed multiple significant loci linked to this phenotype. This was evident from deviations of observed *P*-values from expected values at the tail of the Q-Q plot (Fig. 1D). We used the stringent MLM for GWAS analyses, which corrects for kinship and population structure. This approach detected 73 SNPs at *P* < 10⁻^3^, cumulatively explaining 65% of the phenotypic variation (Fig. 1E), and seven SNPs at *P* < 10⁻^4^. Sixty-five protein-coding genes were located within haploblocks of the GWAS-identified SNPs (*P* < 10⁻^3^), whereas nine were closest to SNPs at *P* < 10⁻^4^ (Table 1). The highest peak in the Manhattan plot was in chromosome 3 at the SNPs *Chr 3:9675996* and *Chr3:9675361*, where an *RNA-binding glycine-rich protein 2* (*RBGB2*; AT3G26420), contributing to freezing tolerance, and an extracellular GDS(L) family lipase (*GGL20*; AT3G26430), playing roles in plant growth and stress responses through lipid metabolism and signaling, were located (Fig. 1F). Another significant SNP *Chr 3: 6009787* lies on an exon of AT3G17570, encoding an F-box/kelch-repeat protein functioning in ubiquitin-mediated protein degradation via the SCF E3 ligase complex, targeting regulatory proteins for turnover in processes like stress response, development, and hormone signaling. *Chr2:16936596* is located on an exon of *E2F target gene 1* (*ETG1*; AT2G40550), required for sister chromatid cohesion and post-replicative homologous recombination DNA repair in the cell cycle. A prominent GWAS peak in chromosome 4 (*Chr4:9200675* and *Chr4:9629622*) is associated with a protein involved in intracellular trafficking and vesicle-mediated protein sorting (*VPS13B*; AT4G17140) and AT4G16260, encoding a putative beta-1,3-endoglucanase. At chromosome 5, *Chr5:23181290* is associated with an ENTH/ANTH/VHS superfamily protein *phosphatidylinositol binding clathrin assembly protein 2A* (*PICALM2A*; AT5G57200), and Ypt/Rab-GAP domain of gyp1p superfamily protein (AT5G57210), both functioning in vesicle trafficking and endomembrane dynamics linked to nutrient uptake and hormonal signaling for root growth. Another peak in chromosome 1 contained a zinc ion-binding protein (AT1G02070) and *negative on TATA-less 1* (*NOT1*; AT1G02080), a core scaffold subunit of the carbon catabolite repression 4 (CCR4)-NOT deadenylase complex involved in mRNA deadenylation and turnover. AT1G75060, associated with *Chr1:28184486*, encodes SAP30 function-related 1 (AFR1), a Sin3-associated polypeptide (SAP) protein. It functions in photoperiodic flowering regulation and epigenetic repression by interacting with histone deacetylase complexes, thereby modulating gene expression. Two more significant SNPs on chromosome 1, *Chr1:4697671* and *Chr1:1686725*, were associated with *6-phosphogluconolactonase 1* (*PGL1*; AT1G13700) and *inositol-polyphosphate 5-phosphatase 13* (*5PTase13*; AT1G05630), respectively. PGL1 catalyzes the hydrolysis of 6-phosphogluconolactone to 6-phosphogluconate in the pentose phosphate pathway, thereby supporting NADPH production for defense against oxidative stress. It appears in phosphate-resupply proteomes, suggesting roles in acclimation to nutrient stress (Smith et al. 2025). 5PTase13 hydrolyzes inositol 1,4,5-trisphosphate to inositol 1,4-bisphosphate, modulating calcium signaling and auxin homeostasis. We assessed the organ-specificity of the GWAS-identified genes by analyzing their expression patterns across publicly available experiments. Our results revealed that most of these genes were expressed in both shoots and roots, suggesting coordinated regulation between these organs to sustain root growth under alkaline stress. However, *PICALM2A*, *TLP-3*, *FRG5*, and AT1G05650 had higher expression in roots, whereas *NTP2*, *CYP71A21*, *SPL8*, and *AT1G02070* showed greater shoot-specific expression (Fig. 1G). Among the GWAS-identified genes for alkalinity tolerance (Table 1), several showed changes in gene expression upon alkalinity exposure. AT1G05650, AT3G22750, and AT4G16260 were upregulated (log_2f_old-change > 1), while *ACL5*, *AUG5*, *CIPK3*, *CYP78A5*, *FLY2*, *FRG5*, *5PTASE13*, *MYB3R-4*, AT1G02110, AT1G24430, and AT5G22310 were downregulated (log_2F_C < −1) in response to alkalinity, as indicated by our in-house and various public transcriptome datasets (Table S3). The GWAS-identified genes (*P* < 10^−3^) belonged to various enriched (*FDR* < 0.05) gene ontology (GO) biological processes, including regulation of growth and development, carbohydrate, polyamine and valine metabolism, cell division, post-replication DNA repair, histone methylation, transcription regulation, mRNA decay, splicing-related snRNA 3’-end processing, snoRNP assembly linked to ribosome biogenesis, protein modifications and Golgi-to-plasma membrane transport, ethylene, brassinosteroid and SA signaling, as well as photosystem II repair, emphasizing the pathways essential for sustaining root elongation amid alkalinity stress (Table S4).

**Table 1.**
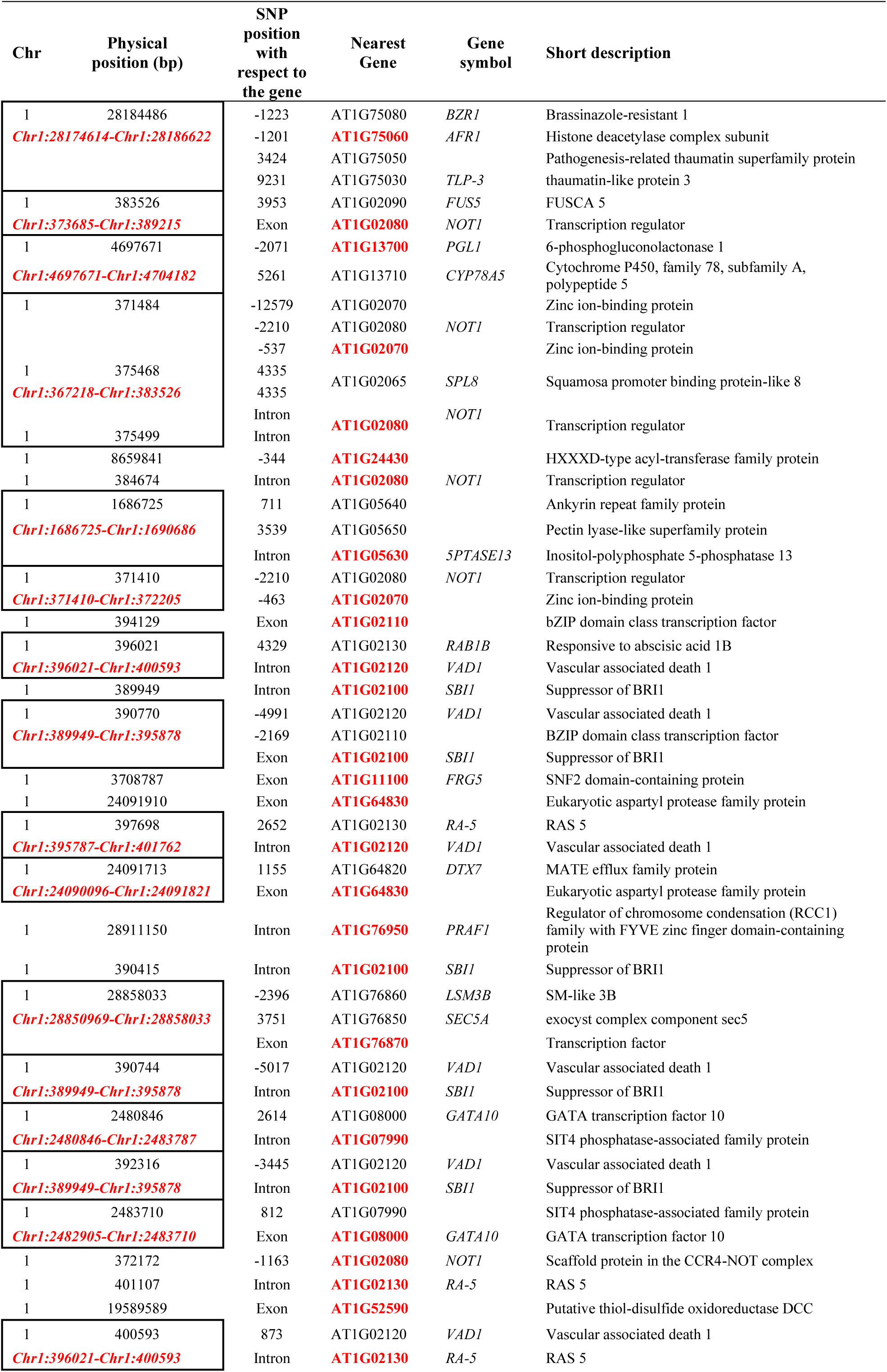
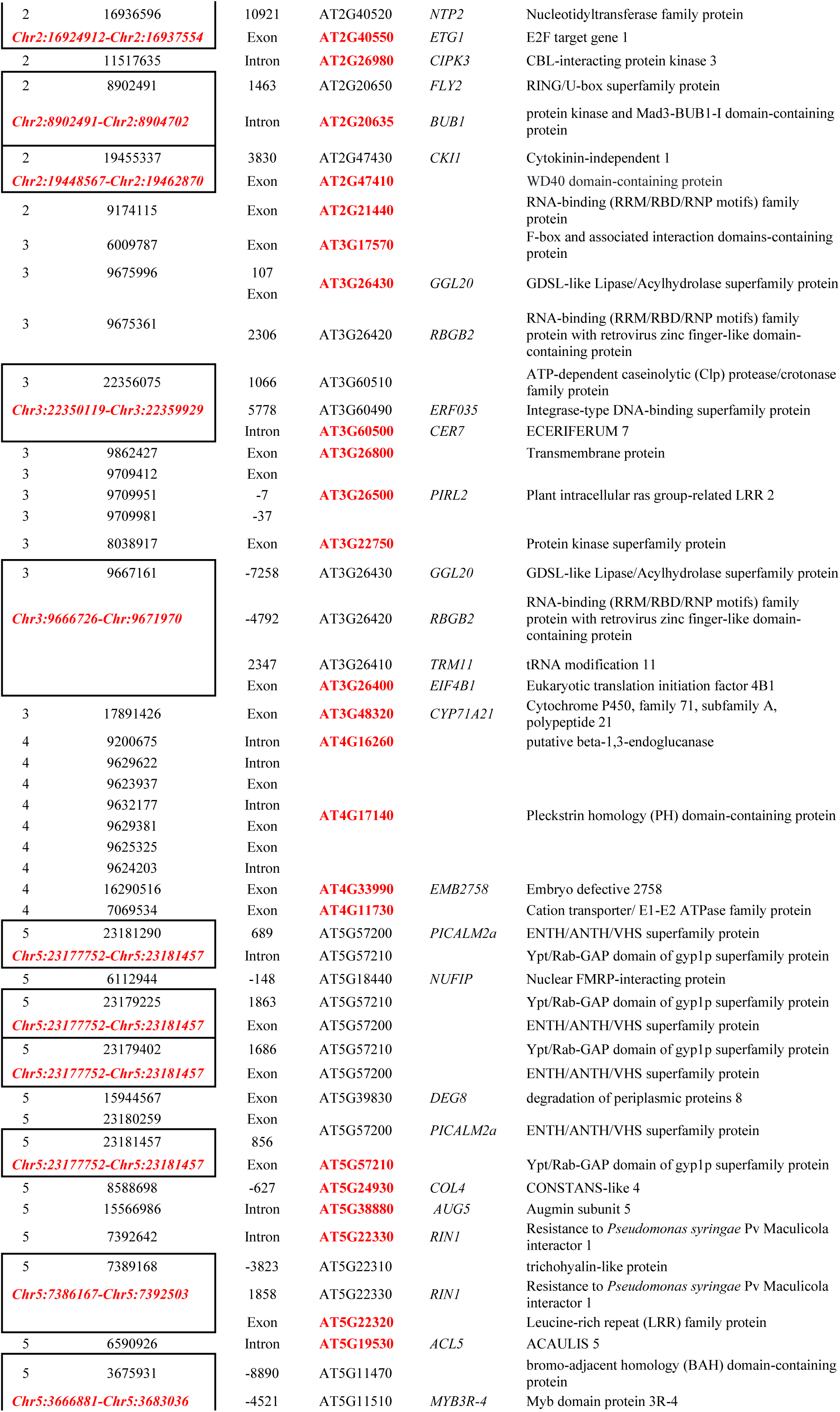
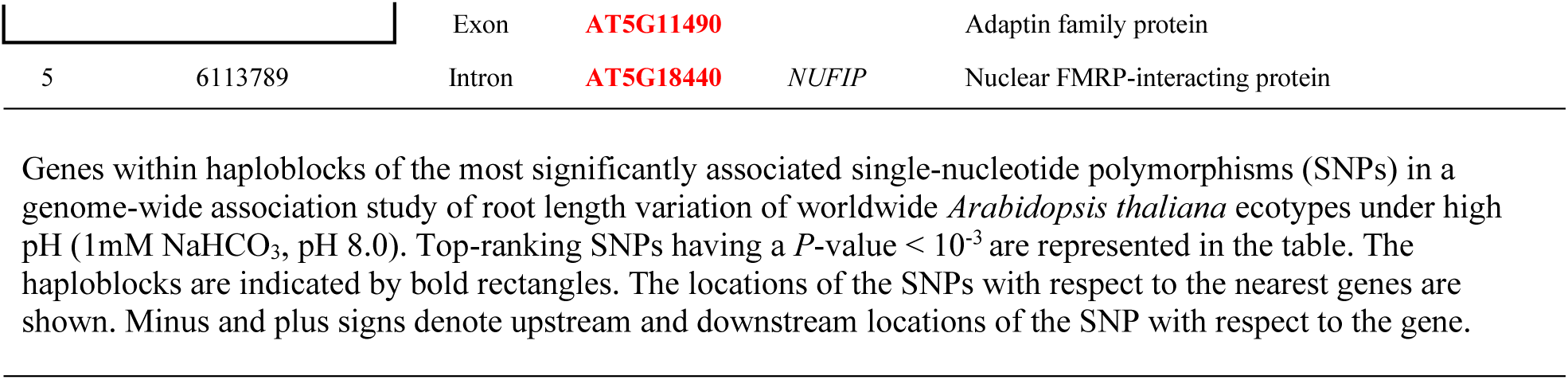
Genes associated with the most significant SNPs in a genome-wide association study of alkalinity stress tolerance in *Arabidopsis thaliana*.

### 3.3 GWAS-identified genes exhibit expression and protein variants linked to alkalinity tolerance

For the 27 top-ranked genes linked to the most-significant GWAS SNPs (*P* < 10^−3.5^), higher-density polymorphism data were obtained from 151 out of 218 ecotypes in the 1001 Genomes database. Among these, 18 genes exhibited promoter and 5’-UTR variants that were significantly associated with alkalinity stress tolerance, as revealed by LAS in targeted genomic regions with denser SNP coverage (Table S5). Ten genes exhibited ‘high-impact’ protein polymorphisms, including splice variants, loss or gain of stop codons, and frameshift mutations, which were mostly associated with a minority of the ecotypes. Out of these, a stop codon gain (Tyr344*) in AT3G17570 in six ecotypes, Kin-0 (RRL 36.86%), Ga-0 (39.39%), and Fei-0 (63.4%) from high CO ^2^⁻ areas, and HR-5 (47.58%), Sq-8 (47.66%), TuWa1-2 (53.77%), and Ema-1 (70.09%) from low CO ^2^⁻ areas, was significantly associated with alkalinity tolerance (Table S5). Eight genes had ‘moderate-impact’ amino acid substitutions significantly associated (MLM or GLM *P* < 0.05) with the RRL phenotype in the LAS. These included Pro51Thr and Thr1616Ser in AT1G02080, Leu976Ser in 5PTASE13, Gly2Glu, Pro23Leu, Ala36Ser, Val88Met, and Asp282Glu in AT1G24430, Leu230Pro and Asn239Asp in AT3G17570, Asp195Asn, Val203Ala, Pro236Leu, His243Tyr and Lys261Arg in GGL20, His2378Arg and Tyr3231His in VPS13B, Ala494Pro, Val505Met, Ser538Ile, Ser538Arg, and Tyr584His in AT5G57200, Asp375Tyr, Glu491Val, Leu516Met, Lys531Arg, Lys547Asn, Lys547Asn, Ser677del (amino acid deletion), and Glu721Gly in AT5G57210.

Expression variation in the top 10 genes, ranked by GWAS *P*-value and featuring promoter (1000 bp upstream of TSS) or 5’-UTR polymorphisms significantly linked to alkalinity tolerance via LAS (Table S5), was confirmed in 40 randomly selected ecotypes (Fig. S2) using primers listed in Table S2. Among them, three genes, *GGL20*, *CER7* (AT3G60500), and AT5G57210, exhibited significant differences in expression levels between SNP alleles on their promoters and UTRs (Fig. 2). *Chr3:9672189* on the 5’-UTR and the GWAS-detected SNP *Chr3:9675996* on the 3’-UTR of *GGL20* were associated with higher expression levels in ecotypes containing the tolerant SNP alleles. Similarly, in *CER7*, higher expression was linked to tolerant alleles at *Chr3:22354096*, *Chr3:22354173*, *Chr3:22354195*, *Chr3:22354196*, and *Chr3:22354206*. But in AT5G57210, a higher expression was observed in sensitive SNP alleles at the GWAS-identified SNP, *Chr5:23181290* on the CDS, and several SNPs in the promoter, including *Chr5:23184164*, *Chr5:23184517*, *Chr5:23184573*, *Chr5:23184635*, and *Chr5:23184863*. Next, we examined the *cis*-elements at these SNP positions by analyzing sequence octamers overrepresented in high- or low-pH or Fe-deficiency stress conditions (Fig. 2, lower panels). The analysis indicated that *Chr3:9672189* on the 5’-UTR of *GGL20* was 7 bp upstream of an octamer “GCAACTCT” that is enriched (RAR > 2; *P* < 0.05) in the promoter regions of genes downregulated by Fe deficiency yet upregulated under low pH stress. In the 5’-UTR of *CER7*, *Chr 3:22354096* falls on the octamer “CTCATCTC”, which is enriched in genes suppressed by Fe deficiency. SNPs *Chr3:22354195* and *Chr3:22354196* occur within the octamer “TGTAAGGG”, which is associated with genes suppressed by both alkalinity and Fe deficiency stresses. On the AT5G57210 promoter, SNP *Chr5:23184164* aligns with “TTACACAT” (enriched in alkalinity-induced genes but suppressed under Fe deficiency), *23184517* with “GAAACCTA” (low pH-induced but Fe deficiency-suppressed), *Chr5:23184573* with “CAAATGTT” (Fe deficiency-suppressed, overlapping a MYC *cis*-element), and *Chr5:*23184863 with “AAGAATTT” (also Fe deficiency-suppressed). Several other SNPs with non-significant associations with alkalinity tolerance in the LAS were located within overrepresented octamers containing known cis-elements (Fig. 2). Our analysis revealed that SNPs within cis-regulatory elements responsive to pH and Fe deficiency stresses drive natural variation in the expression of GWAS-identified genes under alkaline conditions.

**Fig. 2.**
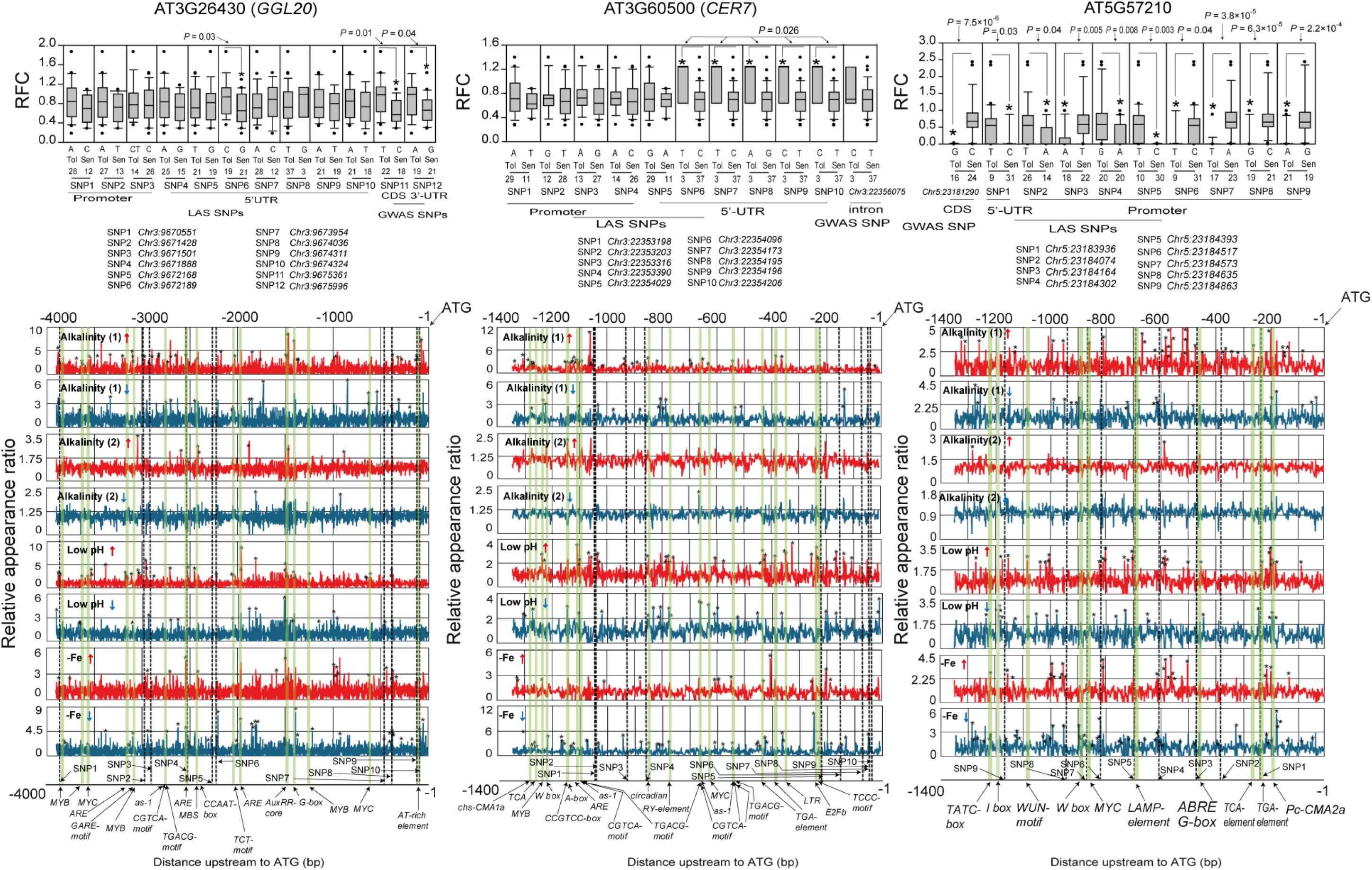
Expression level polymorphisms in genes identified by alkalinity stress GWAS in *Arabidopsis thaliana*. Genes associated with the top-ranking single-nucleotide polymorphisms (SNPs) identified in the genome-wide association study (GWAS; Table 1, S3), which exhibit expression level differences between ecotypes, are presented. Box plots depict genes with significantly associated promoter and 5’-UTR polymorphisms identified in a local association study (LAS) using 40 randomly selected ecotypes from the 218 ecotypes analyzed in the GWAS (Table S5). Each data point represents the average of three biological replicates, each comprising 10 plants. X-axis: ecotypes grouped by SNP alleles (Chr: physical position in bp; N = allele count; sequential numbering of SNP positions). Asterisks: *P* < 0.05 (Student’s *t*-test); left numbers: mean fold change. Beneath the box plots, line graphs illustrate the enrichment of promoter motifs (analyzed as 8-nucleotide octamers) in genes upregulated or downregulated by alkalinity, low pH, and iron deficiency treatments across different experiments. These include our in-house whole-seedling transcriptome (10-day-old plants treated for 5 days with 1 mM NaHCO_3,_ pH 8.0), denoted by *alkalinity (1)*, and a publicly available shoot transcriptome GSE164502 (15-day old plants treated for 3 h/48h with 10 mM NaHCO_3,_ pH 8.3 denoted by *alkalinity (2)* (Pérez-Martin et al. 2021), root transcriptome under low pH stress (GSE18982; Lager et al. 2010; GSE30095; Iyer-Pascuzzi et al. 2011), and root transcriptomes under iron deficiency stress (GSE16964; Li et al. 2010), with mappings of the promoter SNPs overlaid. The relative appearance ratio quantifies the frequency with which each octamer occurs in promoters of stress-responsive genes compared to the *A. thaliana* genome-wide promoter set. Asterisks atop peaks denote significant enrichment (Fisher’s exact test; *P* < 0.05). Green lines highlight the overrepresented octamers that overlap with known cis-regulatory elements from the PlantCARE database, as indicated by the arrows below.

Due to the absence of experimentally resolved structures in PDB, we predicted protein structures *ab initio* using the AlphaFold 3 server (Jumper et al. 2021). Haplotypes chosen for further structural studies were those that involved amino acid changes that markedly alter physicochemical properties, such as polarity or charge, or that induce significant disruptions via frameshifts or premature truncations (Table S6). The F-box domain–containing protein AT3G17570 with Kelch repeats was found to carry frameshift mutations that introduce a premature stop codon leading to truncations in the F-box domain in H1 (Kz-9; 23.2%) from a low CO_32_⁻ region, H11 (Ct-1; 45.1%, Ms-0; 51.2% and En-1; 53.2%) from low and moderate CO_32_⁻ regions, and H13 (Gy-0; 53.1%) low CO_32_⁻ region. Additionally, H10 (Kin-0; 36.9%, and Ga-0; 39.4% from high-CO_32_⁻ regions and HR-5; 47.58%, Sq-8; 47.7%, and Ema-1; 70.1% from low-CO_32_⁻ regions) and H14 (TuWa1-2; 53.8%) from low-CO_32_⁻ regions also contained truncations affecting the last Kelch motif (Fig. 3A). F-box Kelch (FBK) repeat proteins act as substrate-recognition components of the SCF (Skp1–Cullin1–F-box) E3 ubiquitin ligase complex within the ubiquitin–proteasome system (UPS). These proteins are widely conserved across plants and have been implicated in regulating growth and development, secondary metabolism, and stress responses (Wu et al. 2025). Thus, disruption of the F-box/Kelch repeat region may potentially alter substrate recognition and, in turn, affect SCF-mediated ubiquitination and downstream regulatory processes essential for alkalinity tolerance. GGL20, a member of the GDSL esterase/lipase family, is involved in diverse aspects of plant growth, development, and stress responses (Shen et al. 2022). A frameshift mutation resulting in a premature stop codon in moderately sensitive haplotypes H21 (Gel-1; 32.4%, and Kin-0; 36.9%) from high CO_32_⁻ areas and H48 (Stepn-1; 42.8%) from a low CO_32_⁻ area truncates the protein and disrupts the GDSL catalytic domain, thereby compromising enzymatic activity. Members of the GDSL family have been associated with root and seedling development, vegetative growth, and epidermal patterning, likely through their roles in lipid modification and cell wall remodeling. Disruption of these processes may collectively contribute to alkalinity sensitivity. In VPS13B, Wt-5 (RRL 32.99%) from a low CO_32_⁻ region gained a premature stop codon at *Chr4:9618843* (Met3231*) due to a frameshift mutation at *Chr4:9618868* (Met3222), leading to a truncated protein. In AT3G26430, haplotype H21 (Gel-1; 32.36% and Kin-0; 36.86%), originating from high-CO_32_⁻ environments, harbored a frameshift mutation resulting in a premature stop codon at Phe229*. In contrast, H48 (Stepn-1; 42.8%), originating from a low CO_32_⁻ environment, carried a frameshift at *Chr3:9674928* (Ala146), leading to a premature stop codon at (Asp179*). Molecular dynamics simulations of key protein haplotypes (Table S6) in the top GWAS-identified genes revealed how RRL-linked amino acid variants from LAS alter overall structural dynamics and local interactions (Fig. S3), as described in the Supporting results and discussion.

**Fig. 3.**
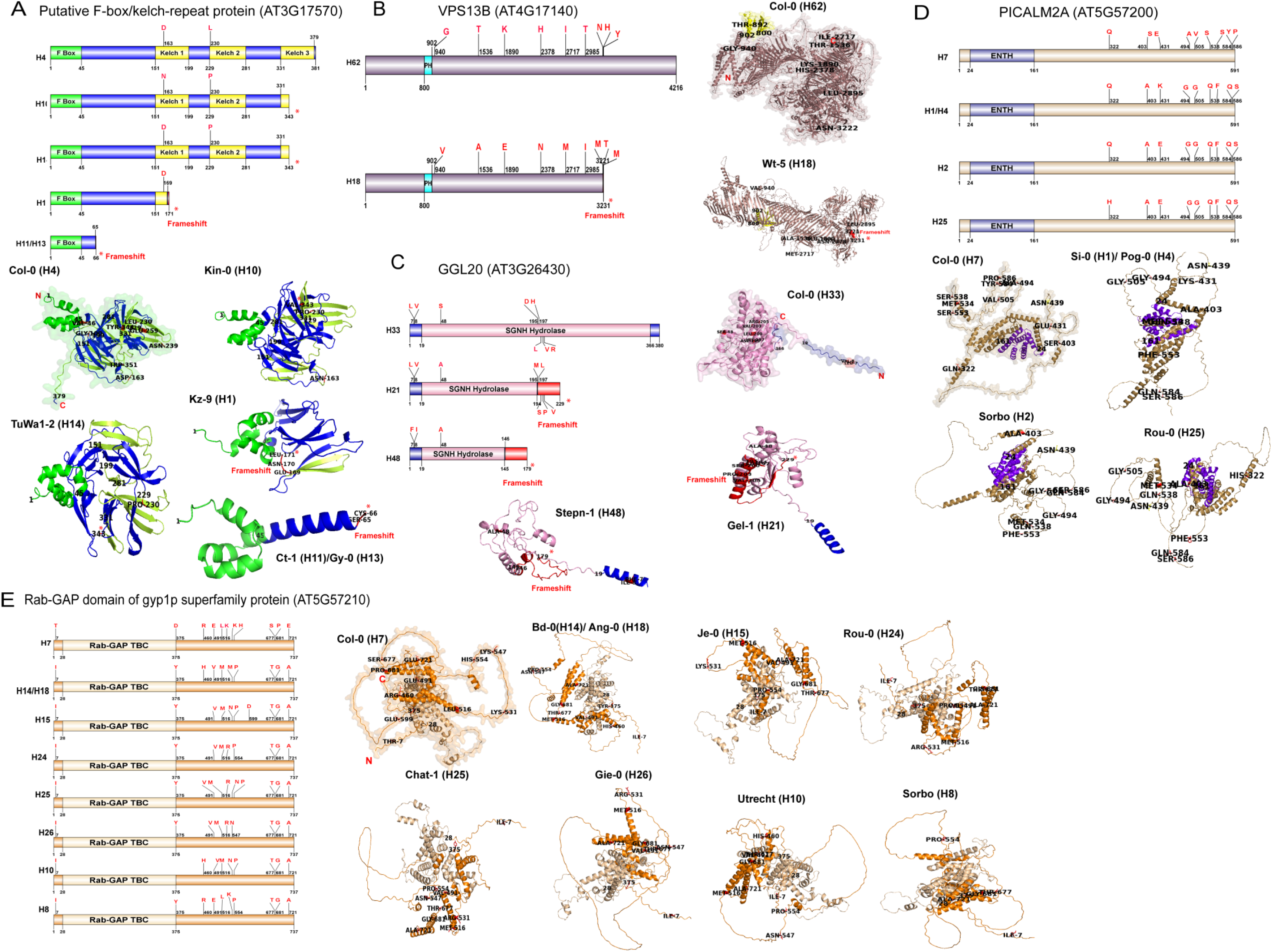
Protein polymorphisms in GWAS-delineated genes. High- and moderate-impact protein polymorphisms in top GWAS candidate genes were detected using the POLYMORPH tool from the 1001 Genomes Project (http://tools.1001genomes.org/polymorph/). Amino acid haplotypes (H1, H2, etc.; see Table S6) were derived from these variants. This figure highlights representative haplotypes with major structural divergences, alongside AlphaFold-predicted 3D models (https://alphafold.ebi.ac.uk/). InterPro domains (https://www.ebi.ac.uk/interpro/) are marked. **(A)** Putative F-box/Kelch-repeat protein (AT3G17570) containing F-box domain and Kelch repeats **(B)** Pleckstrin homology (PH) domain-containing protein (VPS13B; AT4G17140) **(C)** GDSL esterase/lipase (AT3G26430) like superfamily protein-containing the SGNH Hydrolase superfamily domain **(D)** ENTH/ANTH/VHS superfamily protein (PICALM2A; AT5G57200), displaying the ENTH domain **(E)** Ypt/Rab-GAP domain of gyp1p superfamily protein (AT5G57210) showing Rab-GAP TBC domain. Amino acid substitution positions are indicated within the amino acid sequence maps. Asterisks (*) denote premature stop codon gains, while red patches highlight amino acid sequence alterations arising from frameshift mutations.

### 3.4 T-DNA insertion mutants of GWAS-delineated genes show differential alkalinity tolerance from wild-type plants

Homozygous T-DNA insertional mutants for genes closest to the highest-ranking GWAS SNPs underwent phenotyping in hydroponic setups mirroring the conditions of the original GWAS. We used a range of NaHCO_3 d_oses, 1 mM, 1.25 mM, and 2.5 mM, with a pH set to 8.0 for each, to test the alkalinity tolerance of the T-DNA insertion lines (Fig. 4A). Altogether we used 11 confirmed lines viz., *rbgb2* (SALK_109798C), *etg1* (SALK_071046C), *at3g17570* (SALK_100641C), *vps13b* (SALK_096315C), *ggl20* (SALK_100641C), *at5g57210* (SALK_021950C), *afr1* (SALK_110828C), *not1* (SALK_114704C), *picalm2a* (SALK_133151C), *pgl1* (SALK_033902C), and *5ptase13* (SALK_030498C) (Fig. S4). Out of these, *vps13b*, *ggl20*, and *at5g57210* had the T-DNA inserted in exons, whereas *at3g17570* had the T-DNA inserted in the promoter, *afr1*, *etg1*, and *rbgb2* in introns, and *not1* and *picalm2a* in 5’-UTRs. The T-DNA insertions in the promoter, 5’ UTR, and introns suppressed gene expression, leading to gene knockdowns. The knockdown line *rbgb2* showed an average 39% reduction in gene expression, while *etg1*, *afr1*, *not1*, *picalm2a*, and *pgl1* showed reductions of 42%, 45%, 30%, 55%, and 11%, respectively (Fig. S5). At 1 mM NaHCO_3,_ *ggl20*, *at3g17570*, and *afr1* showed sensitivity to alkalinity, with average RRL reductions of 44%, 33%, and 19% from the WT. On the other hand, *etg1* showed tolerance, with an RRL enhancement of 35%. At 2.5 mM, more drastic RRL reductions of 54%, 34%, and 38% were observed for *ggl20*, *at3g17570*, and *afr1*, whereas 21%, 55%, 56%, and 21% increases were observed for *at4g17140*, *at5g57210*, *etg1*, and *rbgb2*, respectively (Fig. 4B). We estimated the Fe contents in the root tissues of six T-DNA lines, which consistently showed differences in RRL than the WT under all concentrations of NaHCO_3 i_n the hydroponic experiment, viz. *ggl20*, *at3g17570*, *afr1*, *vps13b*, *at5g57210*, and *etg1*. The Fe contents of the alkalinity-sensitive mutants *ggl20*, *at3g17570*, and *afr1* showed significant average decreases of 39%, 68.4%, and 32.5% relative to WT, while other elements, viz., Ca, Mg, Mn, Co, Ni, Zn, and Cu levels were not significantly different from the WT (Fig. 4C).

**Fig. 4.**
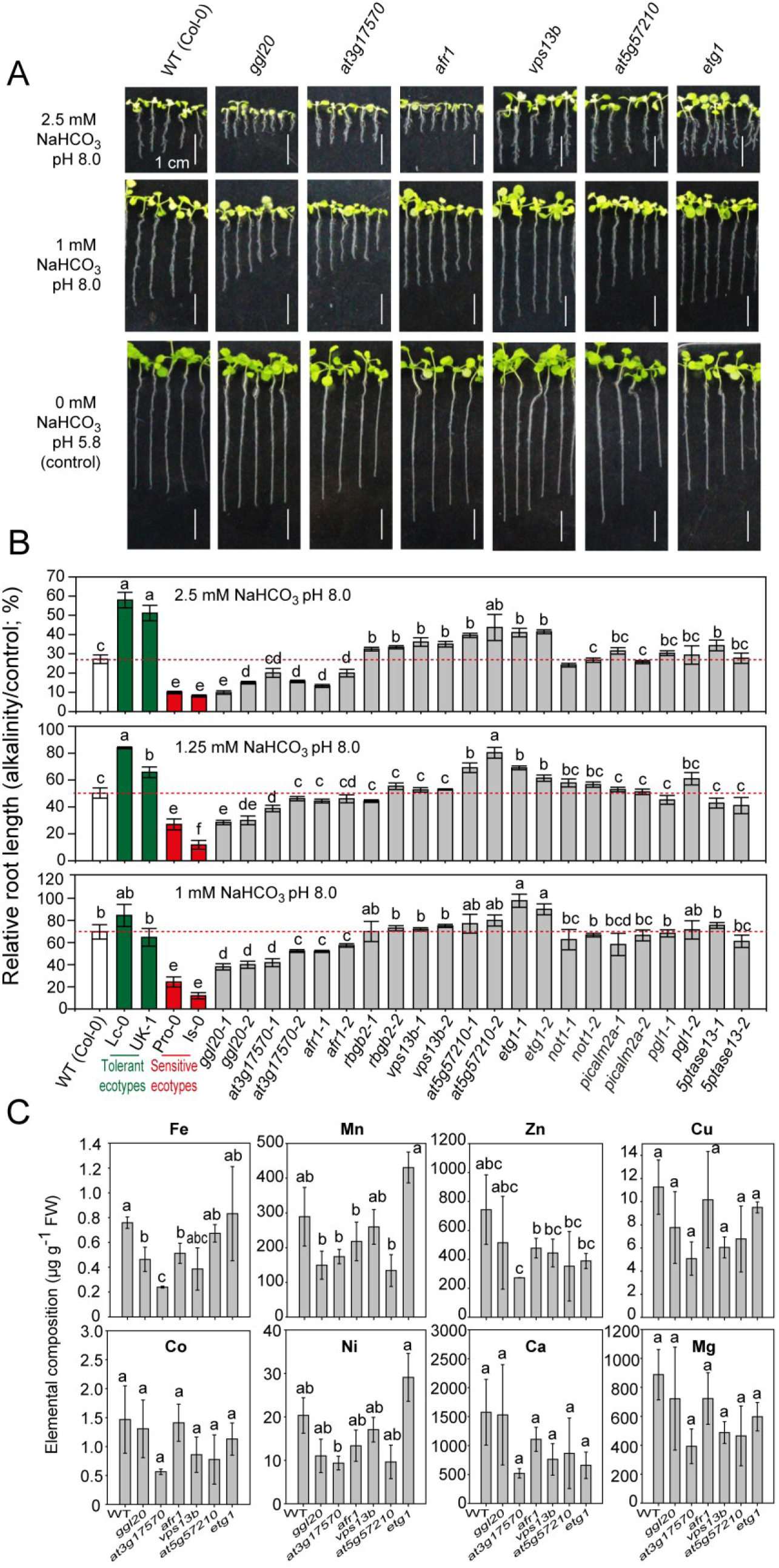
Alkalinity stress tolerance of T-DNA insertion mutants of GWAS-identified genes. **(A)** T-DNA insertion mutants and wild type (WT; Col-0) after growth under 14-day alkalinity stress in hydroponics (¼-Hoagland’s medium supplemented with 1, 1.25, and 2.5 mM NaHCO_3,_ pH 8.0) vs. control (¼-Hoagland’s medium, pH 5.8). Scale: 1 cm. **(B)** The graphs show the relative root length (RRL) in pH 8.0 versus pH 5.8 media (mean ± standard error; *N* = 20 seedlings). WT, two independent homozygous T-DNA lines per gene (progenies of a single T-DNA insertion event), and GWAS controls (tolerant ecotypes: Lc-0, UK-1; sensitive ecotypes: Pro-0, Is-0) are shown. The red dashed lines indicate the WT means. Different letters above the bars indicate significant differences (Tukey’s HSD test; *P* < 0.05). **(C)** Root elemental contents of hydroponically grown WT and mutants under 2.5 mM NaHCO_3,_ pH 8.0, after 14 days, measured by inductively coupled plasma mass spectrometry (ICP-MS), expressed per gram of root fresh weight (FW). The graphs show mean ± standard error; *N* = 3 biological replicates, each consisting of 100 seedlings.

### 3.5 Transcriptome profiling and network analysis uncover key processes regulating alkalinity stress tolerance in *Arabidopsis thaliana*

Ten-day-old Col-0 seedlings exposed to alkaline (1mM NaHCO_3,_ pH 8.0) hydroponic media for five days versus control (pH 5.8) resulted in the upregulation of 116 genes (log_2F_C > 1; *FDR* < 0.05) and the downregulation (log_2F_C < −1; *FDR* < 0.05) of 226 genes (Fig. 5A, Table S7). The expression levels of selected genes were validated by RT-qPCR (Fig. 5B). A functional enrichment analysis (*FDR* < 0.05) of the identified genes revealed key biological processes involved in adaptation to alkalinity. While the upregulated genes were primarily involved in cellular responses to Fe deficiency and nutrient levels, the downregulated genes were responsive to stress, ROS, including H_2O2,_ and were also involved in suberin biosynthesis, secondary metabolism and Fe ion homeostasis (Fig. 5C). We compared our in-house seedling transcriptome data with transcriptome results from rosettes under alkalinity stress (Pérez-Martín et al. 2021), low pH stress in roots (Lager et al. 2010; Iyer-Pascuzzi et al. 2011), and Fe deficiency in roots (Li et al. 2010), revealing substantial overlap in differentially expressed genes (Fig. 5D). To dissect unique and shared genes perturbed by alkalinity, low pH, and Fe deficiency, we conducted a thorough comparative analysis across these conditions. Hierarchical clustering analysis identified five distinct gene groups, some exhibiting opposing expression patterns between high- and low-pH conditions (Fig. 5E). Group 1 comprised genes repressed under alkalinity and Fe deficiency but induced under low pH. Enriched biological processes for this group included Fe sequestration, biosynthesis of cutin, suberin, and wax, lipid metabolism, gibberellin metabolism, and secondary metabolism. This cluster also contained genes directly regulated by the low-pH master TF, sensitive to proton rhizotoxicity 1 (STOP1) (Sawaki et al. 2009), as well as a subset regulated by its paralog, *STOP2*, including *CIPK23*, *SIF1*, *pHRK1*, *DGR1*, AT4G30670, and AT2G28270 (Kobayashi et al. 2014). Group 2 included genes involved in circadian rhythms that were suppressed by all conditions. The third group included genes induced by alkalinity and Fe deficiency but suppressed by low pH. This included genes of phytohormone signaling, transport of oxygen, Fe, copper, and zinc ions, response to Fe starvation and cellular nutrient levels, and biosynthesis of secondary metabolites, including coumarin, flavonoids, and polypropanoids. Group 4 included genes suppressed in our transcriptome and induced by both Fe deficiency and another alkalinity study (Pérez-Martín et al. 2021) but suppressed by low pH. These genes are involved in plant defense responses, hypoxia and heat tolerance, ROS detoxification, and the biosynthesis of indole glucosinolates, olefins, oxylipins, and ethylene. Finally, Group 5 comprised genes induced across all conditions, encompassing general stress responses, glycolysis/gluconeogenesis, hypoxia, Fe, phosphate, and H⁺ transport, as well as the metabolism of aldehydes, coumarin, and vitamin B6. Among genes induced by alkalinity (clusters 3 and 5), several genes could be found that may have a direct role in cellular pH homeostasis by altering H^+^ flux across membranes. These included H^+^ transporters, such as H^+^-ATPase *AHA7*, and H^+^ cotransporters, like *NRT2.6* and *NRAMP4*. *pHRK1* (AT3G46270) functions as a pH-receptor kinase in Arabidopsis, playing a crucial role in sensing and responding to elevated external pH levels (Bailey et al. 2023). Upregulation of the auxin signaling gene *SAUR37* could contribute to alkalinity tolerance by regulating H^+^ pumping via enhanced H^+^-ATPase activity (Spartz et al. 2014). *PDC1*, activated by hypoxia or flooding, lowers cytosolic pH by increasing lactic acid accumulation, triggering a pH-stat mechanism that shifts glycolysis toward *PDC1*-dependent ethanol production. This process consumes H^+^ indirectly by sustaining the fermentation flux and preventing excessive H⁺ buildup (Kürsteiner et al. 2003). *ADH1* supports fermentation under hypoxia, detoxifies reactive aldehydes, and enhances survival in waterlogged or saline conditions via NAD-dependent catalysis. The activation of *ADH1* suggests potential roles in metabolic adaptation to high-pH-induced hypoxia (Luna et al. 2017). The genes for Fe uptake in cluster 3, *IRT1*, *IRT2*, *FRO3*, and *IREG2*, were induced under alkalinity, as it induces Fe deficiency (Hsieh & Waters, 2016). *IRT1* associates with Fe uptake under alkaline-saline conditions in GWAS studies, thereby aiding metal homeostasis and indirectly influencing pH tolerance (Almira Casellas et al. 2022). Under alkaline conditions, the cell wall stiffens due to Fe scarcity (Saleem et al. 2023). Hence, the induction of *XTH13* and *EXP17* (cluster 3) may help remodel cell walls to maintain cell elongation under alkaline conditions.

**Fig. 5.**
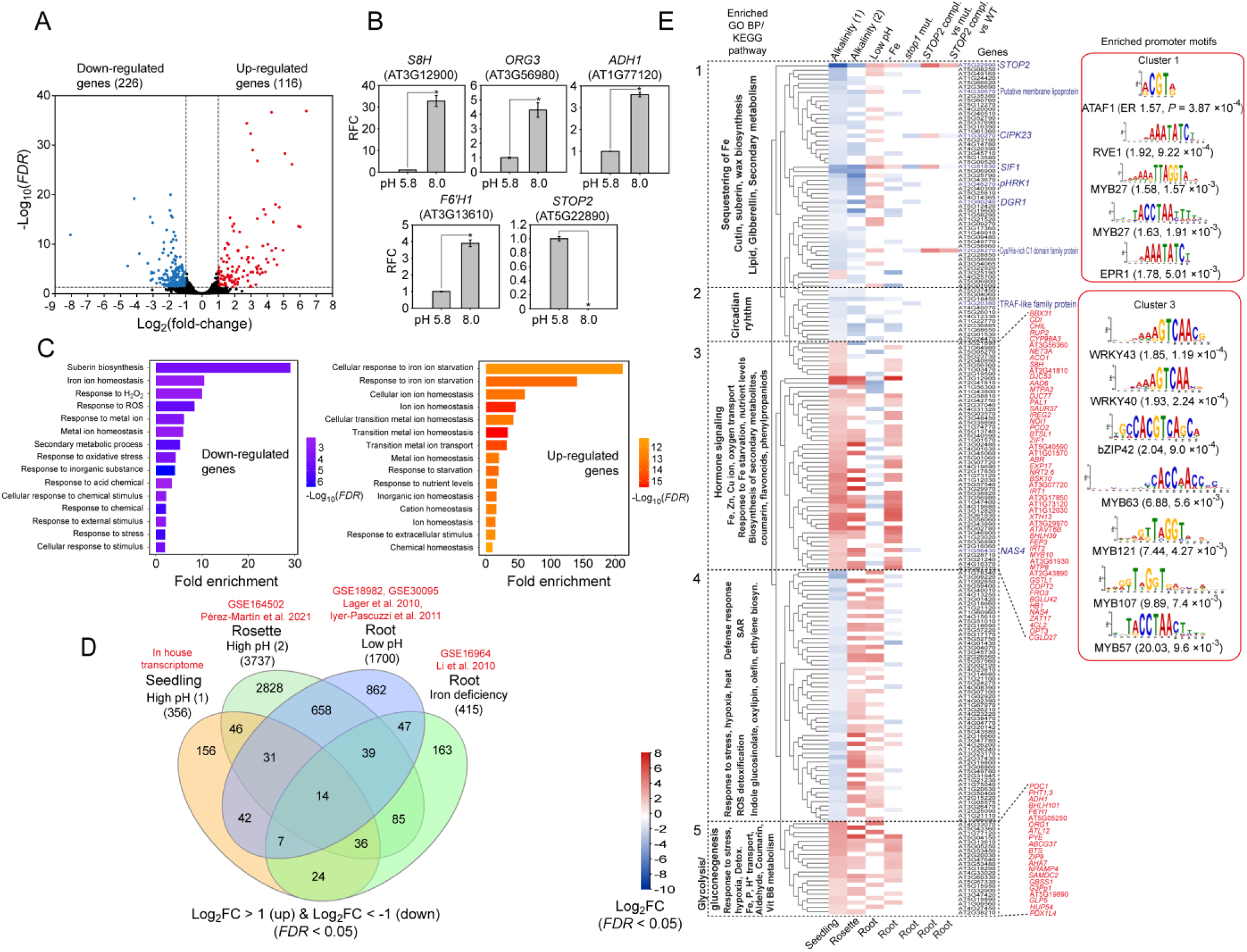
Transcriptome analysis under alkalinity stress in *Arabidopsis thaliana*. *A. thaliana* seedlings were grown hydroponically for 10 days in ¼-Hoagland’s solution, pH 5.8, and then transferred to ¼-Hoagland’s solution with 1 mM NaHCO_3,_ pH 8.0, and grown further for five days to provide stress. Control plants were continuously grown in ¼-Hoagland’s solution for 15 days. The transcriptomes of stressed and control plants were sequenced in three biological replicates, each consisting of 10 whole seedlings, and gene expression was analyzed for differential expression. **(A)** A volcano plot of ‒log_10(_*false discovery rate*) versus log_2(_fold change) shows the distribution of up- (red) and down-regulated (blue) genes. The dotted lines indicate the cutoffs for *FDR* (< 0.05) and fold change (> 1 for up- and < ‒1 for down-regulated genes). **(B)** Reverse transcription quantitative polymerase chain reaction (RT-qPCR) validation of the transcriptome analysis is shown for five highly perturbed genes, *S8H* (AT3G12900), *ORG3* (AT3G56980), *ADH1* (AT1G77120), *F6’H1* (AT3G13610), and *STOP2* (AT5G22890), using *UBQ1* (AT3G52590) as the internal control. The asterisks indicate significant differences between control (pH 5.8) and alkalinity stress (pH 8.0). **(C)** The enriched gene ontology (GO) biological processes for up- and down-regulated genes are shown, along with fold enrichment and a color scale indicating ‒log_10(_*FDR*) value. **(D)** A Venn diagram shows the common genes between different differential gene expression studies in *A. thaliana* (see Methods and Fig. 2 caption for dataset details). The expression fold change cutoffs used are indicated. **(E)** The heatmap displays a hierarchical clustering of genes perturbed in high-pH, low-pH, and iron-deficiency conditions. Additionally, genes up- or down-regulated in the *sensitive to proton rhizotoxicity 1* (*stop1*) mutant (Sawaki et al. 2009), as well as those in the STOP2-complemented line in the *stop1* mutant background, compared to the *stop1* mutant and wild-type (WT) (Kobayashi et al. 2014), are indicated with gene symbols/short descriptions in blue. The enriched GO biological processes and Kyoto Encyclopaedia of Genes and Genomes (KEGG) pathways for each gene cluster are shown. The enriched promoter motifs in clusters 1 and 3, calculated using Simple Enrichment Analysis (SEA) from the MEME suite (https://meme-suite.org/), are shown (*E*-values < 10). The binding transcription factor, motif enrichment ratio (ER; cluster gene set/genome-wide genes), and Fisher’s test *P*-value are shown below.

We analyzed gene coexpression and protein-protein interaction networks to elucidate interrelationships among genes identified through GWAS and transcriptome profiling (Fig. 6). Applying a stringent confidence threshold of 0.7 revealed robust interconnectivity: one large network containing seven GWAS genes and numerous transcriptome genes, alongside 17 smaller networks involving three additional GWAS genes (Fig. 6A). The large network (1) comprised four major clusters. Clusters (i) and (ii) predominantly feature downregulated genes associated with cell cycle and replication regulation; cluster (iii) is linked to ribosome assembly and translation control; and cluster (iv) encompasses a blend of upregulated and downregulated genes tied to amino acid and fatty acid metabolism, along with the mevalonate pathway (Fig. 6B). Hub genes exhibiting the highest interconnectivity within this network included the ribosomal protein large subunit (RPL) and small subunit (RPS) genes, as well as a small nuclear ribonucleoprotein *SNRNP48,* involved in splicing (Fig. 6C). The smaller networks encompassed distinct functional gene groups associated with stress responses, signal transduction, cytoskeleton organization, cell wall biogenesis, misfolded protein response, protein localization, autophagy, sphingolipid metabolism, diterpenoid biosynthesis, endosomal transport, ammonium ion transport, cell differentiation, and deadenylation-dependent mRNA decay.

**Fig. 6.**
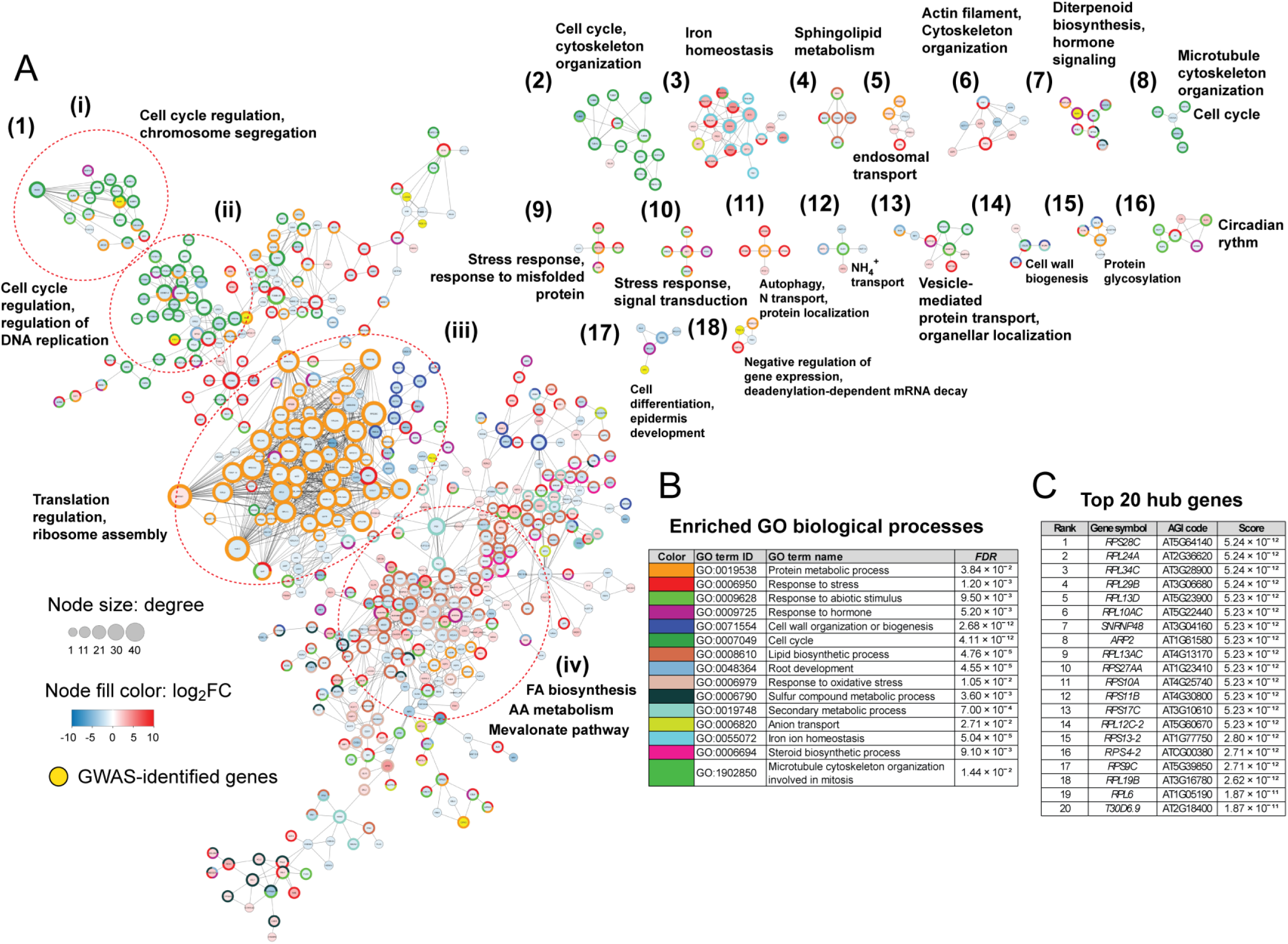
Network of genes regulating alkalinity stress in *Arabidopsis thaliana*. Gene coexpression and protein-protein interaction networks of genes identified by both GWAS and transcriptomic analysis of alkalinity stress in *A. thaliana*. Sixty-five genes from haploblocks associated with SNPs showing *P*-values below 10^‒3^ in the GWAS (detailed in Table 1, S3), together with 342 genes differentially expressed (‒1 < log_2f_old-change > 1; *false discovery rate* < 0.05) by alkalinity in our transcriptome in 15-day-old whole seedling tissues, and 3115 genes differentially expressed in another study of alkalinity stress in rosette tissues, GSE164502 (Pérez-Martin et al. 2021) underwent analysis for network analysis via the STRING platform, applying a stringent confidence threshold of 0.7. Nodes appear as circles, with connecting lines as edges; node dimensions reflect the extent of predicted gene interconnectivity computed by the tool. GWAS-identified genes are featured by yellow-colored nodes, while differentially expressed genes appear in blue and red hues scaled to log_2(_FC). Surrounding node rings highlight significant gene ontology (GO) biological process terms, per the color legend in the table, where enrichments carry Fisher’s exact test *P*-values adjusted by Benjamini-Hochberg *FDR*. The network hub genes are listed by gene symbol, Arabidopsis Genome Initiative (AGI) code, and score. Networks appear numbered with Indo-Arabic numerals, while gene clusters inside the main network use Roman numerals and are outlined by red dotted ellipses. The enriched GO biological processes for each cluster/ small network are shown.

## 4 Discussion

*A. thaliana* ecotypes show no correlation between NaHCO_3 t_olerance and native soil pH because most ecotypes originate from a narrow pH range (5−7), but tolerance weakly correlates with native soil CO ^2^⁻ (Fig. 1C). Adaptation is thus driven by CO_3_ ^2−^/HCO ^−^ ions, not pH. In small-scale local adaptation studies, *A. thaliana* demes show a classic “home-site advantage” on CO ^2^⁻-rich soils: demes from high-CO ^2^⁻ native sites perform better there than foreign demes from low-CO ^2^⁻ sites, and this fitness difference correlates strongly with native soil CO ^2^⁻, not with native soil pH or organic carbon content (Terés et al. 2019). This pattern meets the criterion of local adaptation (local demes outperforming foreign ones) and indicates that soil CO ^2^⁻, rather than pH, is the primary environmental driver of divergent selection among *A. thaliana* ecotypes. In the current GWAS for alkalinity tolerance, the identified candidate genes are those involved in growth, cell division, DNA repair, transcriptional regulation, mRNA decay, protein processing, photosystem repair, and phytohormone signaling (Table S4), not general pH-sensing or pH-regulating genes, because CO ^2^⁻ stress imposes a multifactorial ionic/nutrient imbalance (Na⁺, HCO_3⁻_, high Ca²⁺, Fe deficiency) that disrupts core cellular processes, so natural variation reflects adaptation in fundamental growth and stress-response pathways rather than in short-term pH regulation.

Reverse genetic analysis under various culturing formats revealed significant phenotypic differences between the mutants of *AFR1*, AT3G17570, *GGL20*, and *ETG1* and the WT (Fig. 4, S6, S7). *AFR1*, an evening-expressed core component of the Arabidopsis Switch-Independent 3 (Sin3)-Histone Deacetylase (HDAC) complex, binds the promoters of clock genes *Circadian Clock Associated 1* and *Pseudo-Response Regulator 9* to catalyze H3 deacetylation, thereby repressing their expression during dusk to facilitate proper circadian decline (Lee et al. 2019). Knockout hypersensitivity to HCO_3⁻_ indicates that *AFR1* positively regulates alkalinity tolerance, possibly by temporally gating stress-responsive chromatin remodeling. The AT3G17570 gene product is an F-box protein featuring interaction domains: an N-terminal F-box for SKP1 binding and C-terminal FBA domains that facilitate SCF E3 ubiquitin ligase complex formation, enabling targeted protein ubiquitination and degradation. This structure supports roles in selective proteolysis, potentially regulating stress signaling pathways through protein ubiquitination and turnover (Malik et al. 2020). The presence of amino acid polymorphisms significantly associated with alkalinity tolerance, drastic protein truncations linked mostly to low CO_32_⁻ regions (Table S5, S6), as well as hypersensitivity of the knockdown line to HCO_3⁻_ (Fig. 4, S6) indicates reduced F-box activity failing to appropriately degrade proteins (e.g., aquaporins or ROS scavengers), disrupting H₂O₂ homeostasis and exacerbating oxidative damage under high pH/HCO_3⁻_ conditions (Fig. 4G, H). *GGL20* encodes a GDSL-motif lipase with specificity for long-chain substrates, such as p-nitrophenyl palmitate, but not choline esters, indicating extracellular lipid hydrolysis (Muralidharan et al. 2013). Its signal peptide suggests apoplastic localization for roles in cuticular wax modification, pathogen defence, lipid signaling, and growth regulation (Gao et al. 2017; Shen et al. 2022). Detection of significant promoter and UTR polymorphisms leading to expression level variation (Fig. 2) along with amino acid polymorphisms leading to predicted protein structure variations (Fig. S3) place *GGL20* as a prominent candidate for alkalinity tolerance, validated by the sensitivity of its knockout to HCO_3⁻_ (Fig. 4). *ETG1* encodes a nuclear-localized protein targeted by E2Fa-DPa TFs, essential for cell cycle progression, sister chromatid cohesion, and DNA repair via regulation of centromeric histone deposition and cohesion factors. Given its knockout confers HCO_3⁻_ tolerance (Fig. 4), *ETG1* likely acts as a negative regulator of alkalinity tolerance. HCO_3⁻_ triggers ROS accumulation and DNA strand breaks; *ETG1* knockouts possibly stabilize cohesion/DNA repair machinery (e.g., via reduced E2F targets), enabling survival through G2/M arrest and repair rather than WT’s error-prone progression. T-DNA mutants sensitive to HCO_3⁻_ showed no significant difference in root length from the WT under low pH (4.7) or Fe deficiency (at pH 5.8), suggesting a specific defect in HCO_3⁻_-induced alkalinity stress responses, rather than general pH homeostasis or Fe deficiency, which are ruled out by the insensitivity of the mutants to such conditions (Fig. S8). On the other hand, these mutants accumulate 32-68% less Fe than WT under alkaline conditions (Fig. 4C), indicating that the affected genes promote HCO_3⁻_-specific Fe homeostasis, such as through enhanced mobilization, intracellular compartmentation, or oxidative stress protection under high pH, independent of standard Fe deficiency responses under neutral/ low pH conditions.

The transcriptome analysis of wild-type Col-0 seedlings exposed to HCO_3⁻_ stress revealed key adaptive processes under alkalinity, particularly through comparisons with other gene expression studies (Fig. 5E). A set of genes induced by high pH are repressed at low pH (cluster 3) and another set is suppressed by high pH and induced at low pH (cluster 1) due to antagonistic *cis*-regulatory mechanisms and context-specific signaling pathways that maintain pH homeostasis (Fig. 5). Low pH rapidly activates genes enriched for a CGCG-box motif (overlapping with an ABA-responsive element (ABRE)-related element) bound by CAMTA TFs via Ca²⁺/calmodulin signaling, promoting acid tolerance responses, such as H^+^ extrusion via H⁺-ATPases (Lager et al. 2010). Our enrichment analysis identified enriched motifs, viz., (A/G)CGT(A/G) (ATAF1-binding site) and AATATC (RVE1, EPR1-binding), in the promoters of low pH-induced cluster 1 genes. High pH-responsive genes of cluster 3, in contrast, have enriched GTCAA motifs, which are binding sites for WRKY TFs (Fig. 5). Under low pH conditions, these genes are repressed to prevent maladaptive overlaps. This bidirectional regulation, possibly influenced by pH-dependent promoter accessibility and/or specific TFs like ATAF1 and STOP1/2 for low pH versus alkalinity-specific WRKYs, minimizes metabolic conflicts: low pH prioritizes acid coping through mechanisms such as ion homeostasis and metabolic adjustments, while high pH drives rhizosphere acidification for tolerance. H^+^ efflux under low pH and rhizosphere acidification are distinct, opposing processes mediated by H⁺-ATPases in roots, despite mechanistic similarities. H^+^ efflux occurs during acidic (low pH) stress, where roots pump H⁺ outward via AHA2/AHA7, protecting cytosolic pH, and facilitating nutrient uptake, such as phosphorus. In contrast, rhizosphere acidification occurs under alkaline (high pH) conditions to deliberately lower rhizosphere pH by enhancing H⁺ extrusion or organic acid secretion, thereby solubilizing nutrients such as Fe³⁺ and countering alkalinity. The common MYB binding motifs (ACCTAA and TTAGGT) between the promoters of clusters 1 and 3 indicate shared MYB TF-mediated regulatory mechanisms that provide a constitutive layer of pH stress response, such as ion homeostasis or membrane maintenance, modulated by context-specific TFs like CAMTA or WRKY to drive opposing expression patterns while maintaining homeostasis.

The network of genes identified by the GWAS, as well as those perturbed under alkalinity stress, revealed several functional clusters (Fig. 6). These included genes suppressed under alkalinity stress, including those involved in microtubule cytoskeleton organization, cell cycle, protein metabolism, and ribosome biogenesis. On the other hand, genes involved in Fe homeostasis, autophagy, protein localization, vesicle-mediated protein localization, and response to misfolded proteins were induced by alkalinity. We also identified several upregulated genes responsive to oxidative stress and hormone signaling, which clustered with downregulated genes involved in lipid metabolism. The gene network patterns under alkalinity stress suggest that plants activate protective mechanisms, such as autophagy and Fe homeostasis, to recycle nutrients and combat oxidative damage, while halting energy-intensive processes, such as cell cycle progression, ribosome biogenesis, and microtubule organization, to conserve resources (Li et al. 2022). Suppression of microtubule cytoskeleton and cell cycle genes indicates a stress-induced arrest in cell division and expansion, allowing reallocation of resources toward survival rather than growth, a common adaptation seen in salt and alkaline conditions (Kumar et al. 2022). Previous studies indicate a strong connection between alkalinity tolerance and stabilization of the microfilament cytoskeleton (Liu and Guo, 2011). Upregulation of autophagy, protein localization, and vesicle-mediated transport indicates enhanced degradation and recycling of misfolded proteins and damaged organelles, thereby maintaining cellular homeostasis under high-pH-induced proteotoxic stress (Zheng et al. 2018). Induced Fe homeostasis genes likely counter alkalinity-induced Fe deficiency, supporting antioxidant enzymes to mitigate ROS, with hormone signaling coordinating this response (Tripathi et al. 2018). Clustering of upregulated oxidative stress/hormone genes with downregulated lipid metabolism genes suggests ROS-mediated lipid peroxidation, prompting reduced lipid synthesis to prevent membrane damage and stabilize cellular integrity (Savchenko and Tikhonov, 2021).

To summarize, this study established a comprehensive genetic map of alkalinity tolerance in *A. thaliana* by integrating GWAS and transcriptomics. Key pathways involve lipid metabolism, protein degradation, and DNA repair. Natural variation is driven by specific polymorphisms: promoter and 5’-UTR SNPs in *GGL20*, *CER7*, and AT5G57210 modulate stress-responsive expression, while high-impact protein truncations, as in AT3G17570 and frameshifts in VPS13B and GGL20, alter functional domains essential for tolerance. Moderate-impact amino acid substitutions in genes like AT5G57210 and *5PTASE13* further differentiate tolerant ecotypes. Functional validation identified *ETG1* as a negative regulator, whereas *GGL20*, AT3G17570, and *AFR1* are essential for survival under alkaline conditions. These findings demonstrate that alkalinity adaptation extensively overlaps with Fe-deficiency pathways. Future research should investigate the precise regulatory mechanisms of chromatin-modifying genes such as *AFR1* in temporal stress gating, clarify the roles of hub genes involved in ribosome assembly and translation control, and explore the interplay between pH-sensing complexes and nutrient-specific signaling. Translational engineering of these candidate orthologs in major crops will be crucial for developing resilient varieties for alkaline soil regions.

## Author contributions

Neelam Jangir: Investigation, Writing – original draft. Rishabh Kumar: Investigation, data visualization, Writing – original draft. Surbhi Vilas Tajane: Investigation, data visualization, Writing – original draft. Devanshu Verma: Investigation. Rabisankar Mandi: Investigation. Sucharita Dey: Supervision, Resources. Ayan Sadhukhan: Funding acquisition, Conceptualization, Supervision, Visualization, Writing – original draft, Review & editing.

## Acknowledgements

The authors thank the Department of Bioscience and Bioengineering, IIT Jodhpur, for providing infrastructure support, and the Science and Engineering Research Board and the Department of Biotechnology, Govt. of India, for funding. RM and NJ received doctoral fellowships from the University Grants Commission, India, DV from the Department of Biotechnology, and RK from the Ministry of Education, Government of India.

## Data availability

The raw transcriptomic data are available in the NCBI Sequence Read Archive under accession PRJNA1270151.

