## Supplementary data for "Genome-wide association study and transcriptomics reveal the genetic architecture of alkalinity tolerance in *Arabidopsis thaliana*"

**Supporting Information:**

### 1. Supporting Methods

#### 1.1 *In silico* protein structure analysis

All-atom molecular dynamics (MD) simulations of the apo proteins were performed in GROMACS 2024 using the CHARMM27 force field (Ponder & Case, 2003). To enhance conformational sampling, each of the four protein systems was simulated for 200 ns in explicit solvent, and each simulation was independently repeated twice. Standard protonation states corresponding to neutral pH were assigned to all titratable residues. Each protein was positioned at the center of a cubic box, which was subsequently filled with TIP3P water molecules. The overall charge of the system was neutralized by adding appropriate numbers of $Na^+$ and $Cl^-$ ions, and energy minimization using the steepest-descent algorithm was performed to eliminate steric clashes and other unfavorable interactions. The minimized systems were heated from 0 to 300 K over 100 ps under NVT conditions, followed by 100 ps of equilibration in the NPT ensemble to stabilize pressure and density. Production MD simulations of 200 ns were then conducted with a 2 fs integration time step, and for each protein, the trajectory with the most favorable stability indicators (e.g., backbone RMSD and radius of gyration) was retained for downstream analysis. Long-range electrostatics were computed with the Particle Mesh Ewald (PME) scheme, and periodic boundary conditions were imposed in all three dimensions. Trajectories were analyzed using built-in GROMACS utilities and in-house Python scripts. Hydrogen bonds were identified with HBPLUS (McDonald & Thornton, 1994). Ionic interactions were defined as contacts between side-chain atoms of positively charged residues (Lys, Arg, His) and negatively charged residues (Asp, Glu) within 4.0 Å (Gowri Shankar et al. 2007), and hydrophobic interactions as side-chain contacts between hydrophobic residues within 4.0 Å (Marik et al. 2026).

#### 1.2 Phenotyping T-DNA mutants in Murashige Skoog (MS) media

Plants were also phenotyped by growing in ½-MS media with 1.5% sucrose, pH 5.8, alone or supplemented with 3.5 mM $NaHCO_3$, pH 8.0, adjusted with KOH before autoclaving. Media were solidified with 0.7% agar. Seeds were surface-sterilized with 1% sodium hypochlorite and 70% ethanol, then rinsed 5 times with deionized water before aseptic sowing on solid media. The plants were photographed after 14 days of growth. Ten-day-old seedlings grown on control agar-solidified ½-MS medium (pH 5.8) were aseptically transferred to ½-MS agar plates supplemented with 3.5 mM $NaHCO_3$ (pH 8.0) for 72 h. Thereafter, plants were transferred to 0.006% (w/v) bromocresol purple-containing water-agar plates, pH 6.0. Plates were incubated vertically for 48 h, after which they were photographed, and the color change in the media from purple to yellow around the roots was recorded as a readout of rhizosphere acidification.

#### 1.3 Phenotyping T-DNA mutants in soil

For soil culture, plants were first grown hydroponically for three weeks and then transferred to a 1:2:1 mixture of desert soil, Soil Rite, and vermiculite. Control plants were watered with only deionized water (final soil pH 8.4), while stressed plants were watered with 7 mM or 10 mM $NaHCO_3$ every 7 days and with deionized water every 2 days. The plants were photographed after one month of growth. The shoot biomass and relative shoot diameter (alkaline soil/control) were recorded. The final soil pH was measured by suspending the soil in 5 times (weight/volume) deionized water, vortexing for 1 hour, performing a brief centrifugation, and measuring the supernatant pH. The electrical conductivity (EC) was measured using an Eutech™ PC 700 Multi-parameter Meter (Thermo Fisher Scientific, MA, USA). Chlorophyll contents in rosette leaves were measured using the method of Arnon (1949). Leaf $H_2O_2$ and MDA levels were estimated as described previously (Elstner and Heupel, 1976; Heath and Packer, 1968).

### 2. Supporting Results and Discussion

#### 2.1 Amino acid substitutions linked to alkalinity tolerance drive divergent dynamics in polymorphic proteins

Molecular dynamics simulations across haplotypes carrying LAS-associated contrasting (sensitive and tolerant) SNP alleles indicated that amino acid substitutions that alter side-chain polarity or charge led to transient shifts in local protein structures, accompanied by changes in hydrogen bonds, hydrophobic contacts, and ionic interactions among residues. In GGL20, the possible ligand (palmitate)-binding site (vide UniProt) was located at residues 97 and 99, with additional hydrophobic interactions involving amino acid residues 28, 82, and 84, spatially distant from the LAS-associated polymorphism positions at 195, 203, 236, 243, and 261. GGL20 protein variants Col-0 and Uk-1 (Fig. S3A) exhibit the lowest backbone RMSD values, indicating greater overall structural stability, whereas Ba-1 and Gu-0 show moderately higher RMSD values, and Rovero-1 shows the largest deviations, suggesting increased global flexibility (Fig. S3B). The RMSF profiles largely overlap across most residues, confirming that polymorphic variation does not induce widespread changes in flexibility (Fig. S3C). Local interaction analysis revealed that polymorphic residues, such as Asp195Asn, Val203Ala, and Arg261Lys, form highly persistent hydrogen bonds across all variants, thereby contributing to overall structural stability. Notably, Asp195 in Col-0, Ba-1, and Rovero-1 forms more hydrogen bonds compared to Asn195 in Uk-1 and Gu-0, indicating enhanced local stabilization by the charged residue. Polymorphic sites, particularly at positions 236 and 243, exhibit variability in hydrophobic and ionic interactions, reflecting locally altered interactions; Pro236 shows more hydrophobic interactions than Leu236 (Fig. S3D). Interestingly, some haplotypes are associated with low soil carbonate levels at the ecotypes' geographical origins (Fig. S3E). The AT3G17570 protein in both Col-0 and Buckhorn Pass (Fig. S3F) equilibrates and remains stable throughout the 200 ns simulation, with minimal backbone RMSD fluctuations, indicating that the amino acid substitutions at positions 230 and 239 do not disrupt the global fold and protein dynamics. The RMSF profiles largely overlap across most residues, suggesting that these polymorphisms do not induce significant changes in overall flexibility. With overlapping RMSD/RMSF profiles indicating that substitutions at these specific positions do not measurably alter overall structure or dynamics (Fig. S3G, H). The residue Pro230 shows a higher number of hydrophobic interactions in Buckhorn Pass, compared with Leu230 in Col-0. These localized structural enhancements, characterized by greater overall stability (low RMSD), persistent hydrogen bonding (e.g., Asp195 in GGL20), and optimized hydrophobic/ionic interactions (e.g., Pro236/Leu236 in GGL20, Pro230/Leu230 in AT3G17570), in tolerant haplotypes likely preserve critical functions like ligand binding or signaling under alkalinity stress, conferring ecotypic tolerance unlike the flexibility/disruptions in sensitive variants.

#### 2.2 T-DNA insertion mutants of GWAS-delineated genes show differential alkalinity tolerance from wild-type plants in solid media and soil

The six T-DNA lines, which consistently showed differences in RRL from the WT under all concentrations of $NaHCO_3$ in the hydroponic experiment, viz. *ggl20*, *at3g17570*, *afr1*, *vps13b*, *at5g57210*, and *etg1*, were additionally phenotyped in solidified MS media containing sucrose (Fig. S6) and in soil (Fig. S7). In solid media, *ggl20*, *at3g17570*, and *afr1* showed sensitivity, with 74%, 78%, and 63% decreases in RRL relative to WT, respectively, while etg1 showed tolerance, with a 70% increase in RRL.

In the soil culture assay, alkalinity stress was induced by irrigation with 7 mM and 10 mM $NaHCO_3$, raising soil pH to 8.8 and 9.0, respectively. The EC of the soil irrigated with 0 mM, 7 mM, and 10 mM $NaHCO_3$ reached 335, 370, and 379 µS $cm^{-1}$ at the end of the experiment, reflecting negligible salinity contributions from the added $NaHCO_3$, indicative of non-saline conditions primarily driven by alkalinity-induced pH elevation rather than osmotic stress from dissolved ions. Sensitive genotypes exhibited smaller, curled, and pigmented leaves after one month of growth in soil culture, whereas tolerant genotypes displayed larger, expanded, and green leaves (Fig. S7A). At 10 mM $NaHCO_3$, the relative rosette diameters of *ggl20*, *at3g17570*, and *afr1* were significantly smaller than those of the wild type by 22%, 28%, and 26%, respectively, whereas *vps13b* and *AT5G57210* were larger by 18% and 27%, respectively (Fig. S7B). In terms of shoot biomass, *ggl20*, *AT3G17570*, and *afr1* showed sensitivity (reductions of 13%, 32%, and 37%, respectively), while *vps13b*, *AT5G57210*, and *etg1* exhibited tolerance (increases of 65%, 12%, and 54%, respectively) (Fig. S7C). Additionally, *AT3G17570* and *afr1* had lower chlorophyll content (by 23% and 53%, respectively), whereas *vps13b* showed higher content (by 30%) (Fig. S7D). The $H_2O_2$ contents of the sensitive mutants *at3g17570* and *afr1* showed a significant increase of 29% and 48%, respectively, whereas the tolerant mutants *vps13b*, *at5g57210*, and *etg1* showed corresponding increases of 37%, 32%, and 15%, respectively, compared to the WT (Fig. S7E). The extent of lipid peroxidation, estimated by MDA content, increased 1.9- and 2.7-fold in sensitive mutants *at3g17570* and *afr1*, respectively (Fig. S7F).

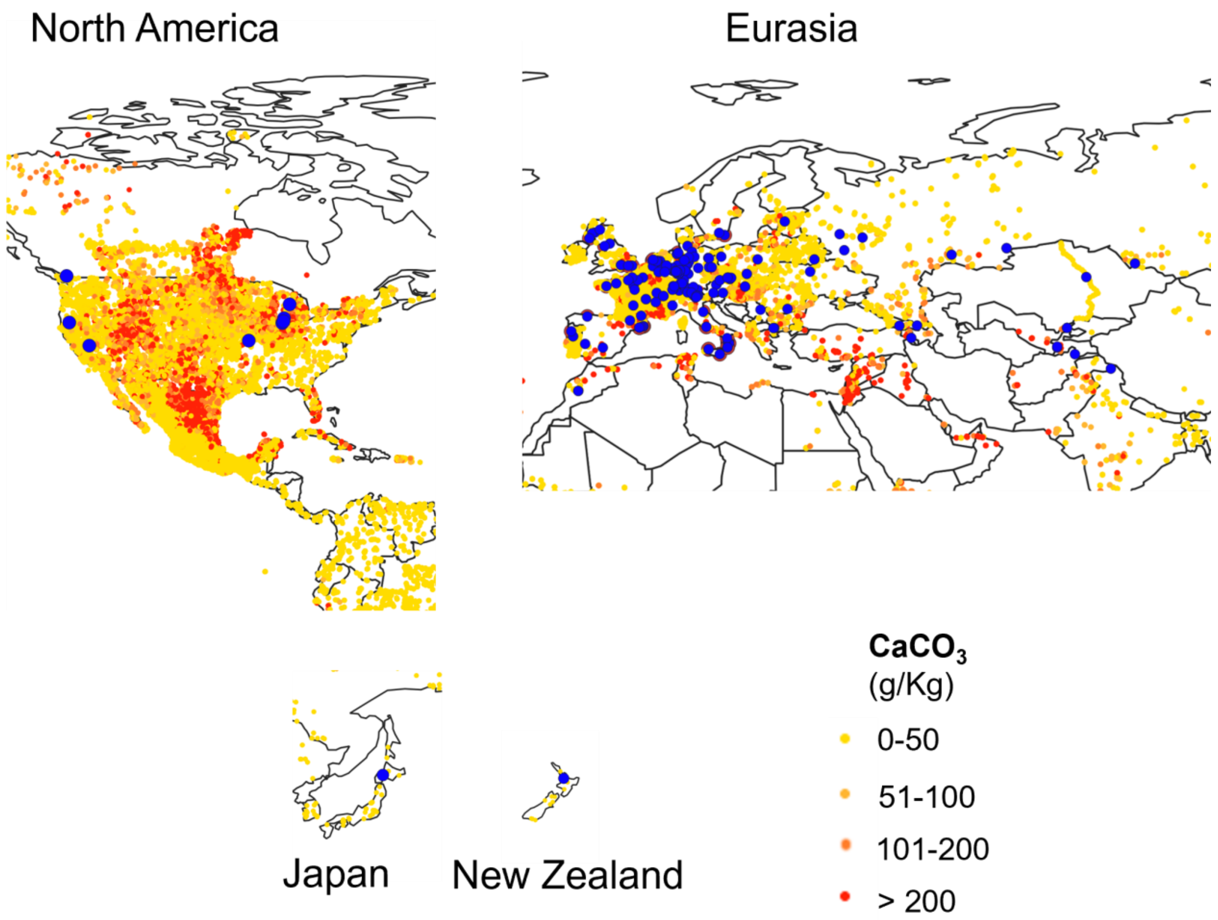


**Fig. S1. Global distribution of *Arabidopsis thaliana* ecotypes overlaid on soil calcium carbonate ($CaCO_3$) content**. Ecotype locations are indicated by dark blue dots. Soil $CaCO_3$ content is shown in g·kg$^{-1}$; white areas correspond to regions with unknown or not reported $CaCO_3$ levels. Soil CaCO3 measurements were obtained from the WoSIS database (https://isric.org/explore/wosis). Global pH data layers were overlaid with the geographic coordinates of *A. thaliana* ecotypes via QGIS 3.4 Madeira (long-term release) software (https://qgis.org/).

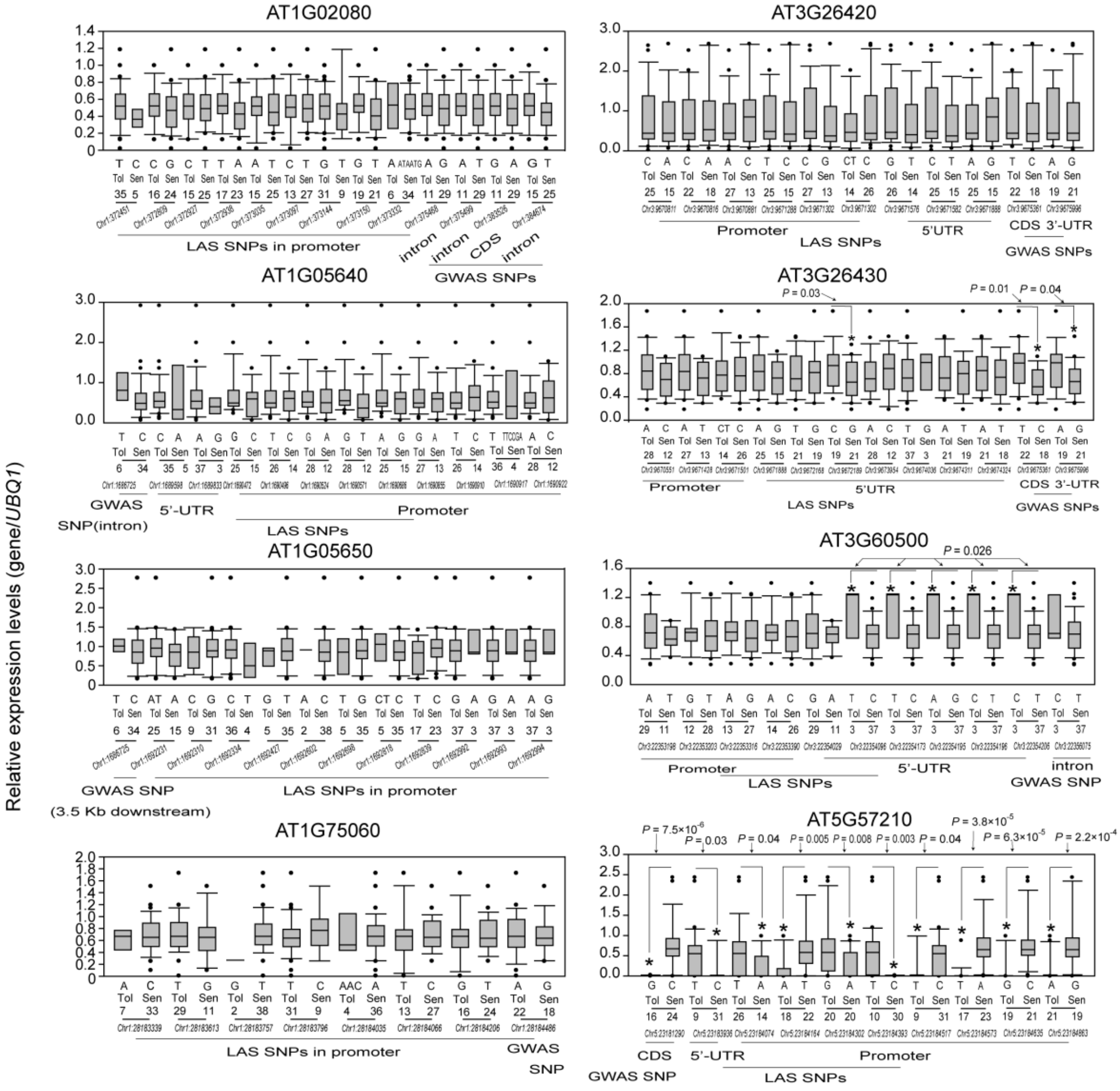


**Fig. S2. Expression level polymorphisms in genes identified by a GWAS of alkalinity stress tolerance in *Arabidopsis thaliana.*** Box plots depict genes with significantly associated promoter and 5'-UTR polymorphisms identified in a local association study (LAS) using 40 randomly selected ecotypes from the 218 ecotypes analyzed in the GWAS (Table S5). Each data point represents the average of three biological replicates, each comprising 10 plants. X-axis: ecotypes grouped by SNP alleles (Chr: physical position in bp; N = allele count; sequential numbering of SNP positions). Asterisks: $P < 0.05$ (Student's *t*-test); left numbers: mean fold change.

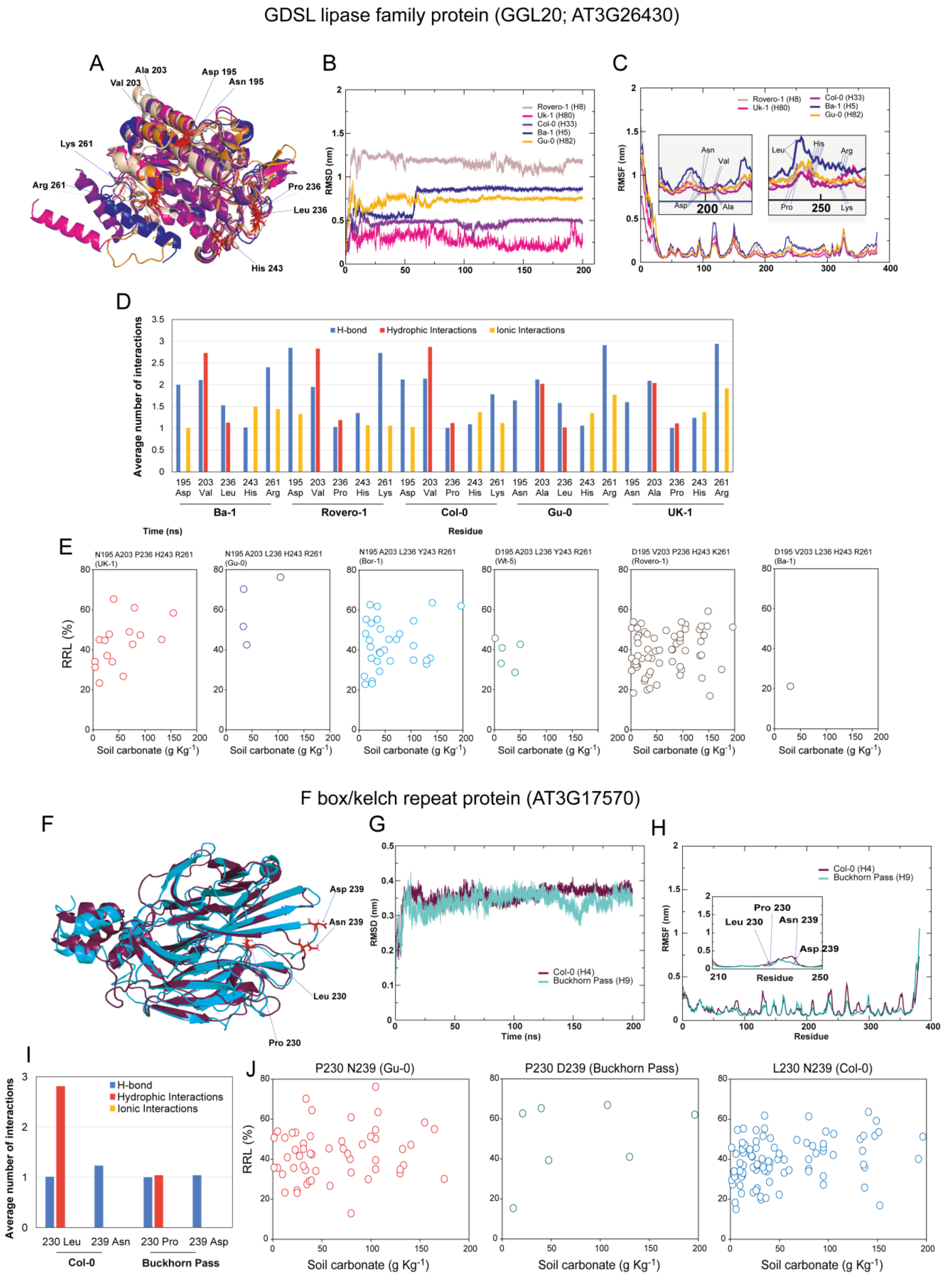


**Fig. S3 Effect of amino acid substitutions on predicted protein dynamics, stability, and conformational landscape.** Moderate-impact amino acid substitutions in two top-ranking GWAS candidate genes, a GDSL-lipase family protein (GGL20; AT3G26430) and an F-box/Kelch-repeat protein (AT3G17570), were identified using the POLYMORPH 1000 Genomes variants web tool (http://tools.1001genomes.org/polymorph/), and amino acid sequence haplotypes were constructed (Table S5). Representative haplotypes with contrasting amino acid substitutions were modelled, and their AlphaFold-predicted three-dimensional structures (https://alphafold.ebi.ac.uk/) are shown as structural superpositions after molecular dynamics (MD) simulations (A, F). Root-mean-square deviation (RMSD) plots over simulation time depict backbone deviations from the initial conformation, indicating overall structural stability and conformational changes during MD simulations (B, G). Backbone root-mean-square fluctuation (RMSF) profiles as a function of residue number illustrate the flexibility and dynamic behavior of individual residues during MD simulations (C, H); the color key for the curves is identical between panels (B) and (C), and between (G) and (H). Predicted intramolecular interactions (hydrogen bonds, hydrophobic interactions, and ionic interactions) involving polymorphic amino acids and their neighboring residues are shown as bar graphs (D, I); the average number of interactions was calculated as the total interaction count divided by the number of frames in which the interaction was observed. Climate association of protein haplotypes is presented as correlation plots of alkalinity-tolerance phenotype (relative root length, RRL%) versus soil carbonate content (g $kg^{-1}$) (E, J).

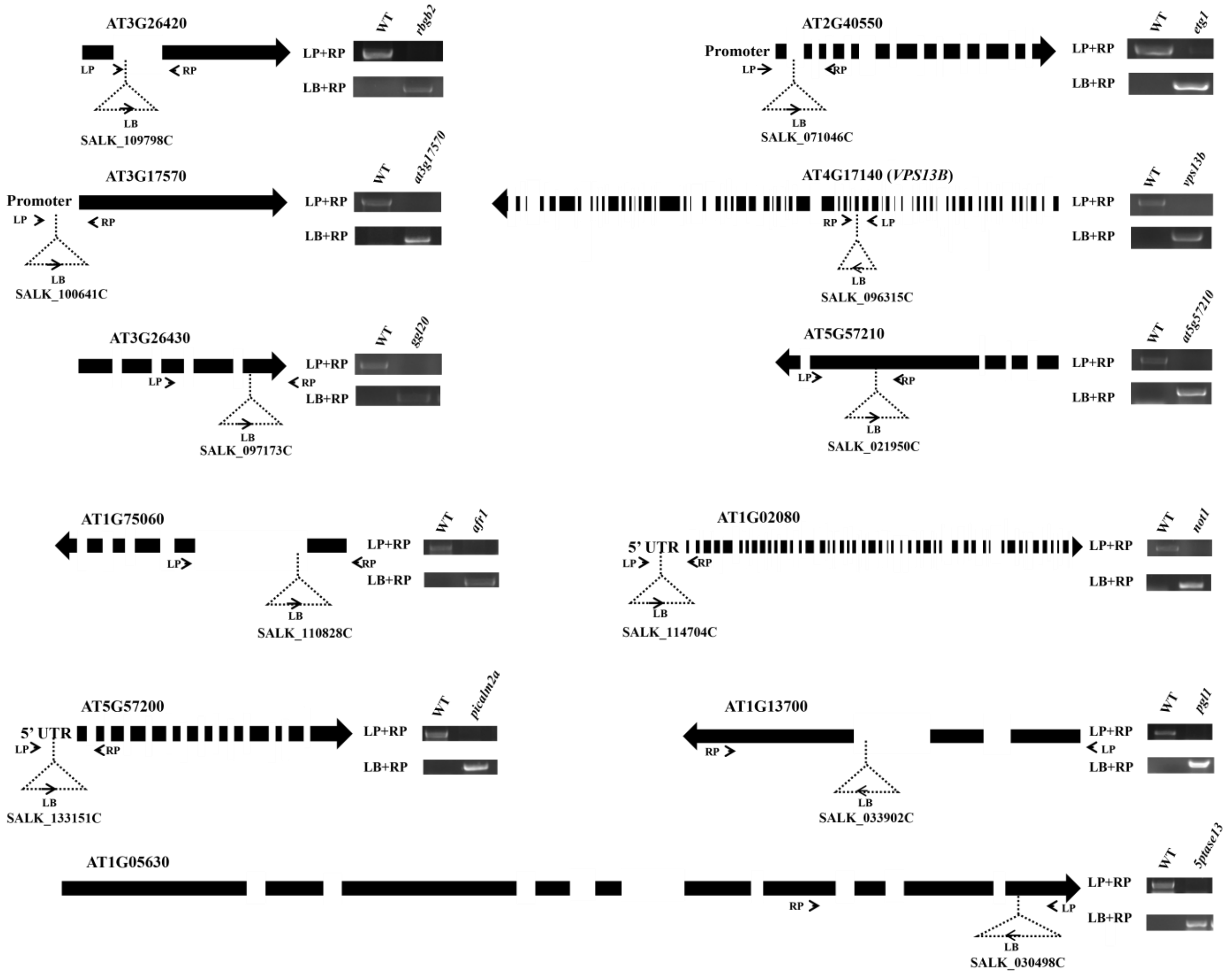


**Fig. S4 Homozygosity validation of the TDNA insertion mutants used in this study**. TDNA insertion mutants used in the study, viz., *rbgb2* (SALK_109798C), *etg1* (SALK_071046C), *at3g17570* (SALK_100641C), *vps13b* (SALK_096315C), *ggl20* (SALK_100641C), *at5g57210* (SALK_021950C), *afr1* (SALK_110828C), *not1* (SALK_114704C), *picalm2a* (SALK_133151C), *pgl1* (SALK_033902C), and *5ptase13* (SALK_030498C) were tested for homozygosity using PCR-based method. The list of primers is available in Table S2. Dotted triangles represents location of the inserted T-DNA, where left border of T-DNA (LB), reverse genomic primer (RP) and forward genomic primer (LP) are labelled together with gel images of amplification. Thick black arrows denotes gene coding sequences from the 5' to 3' direction, where intermediate slits indicate introns.

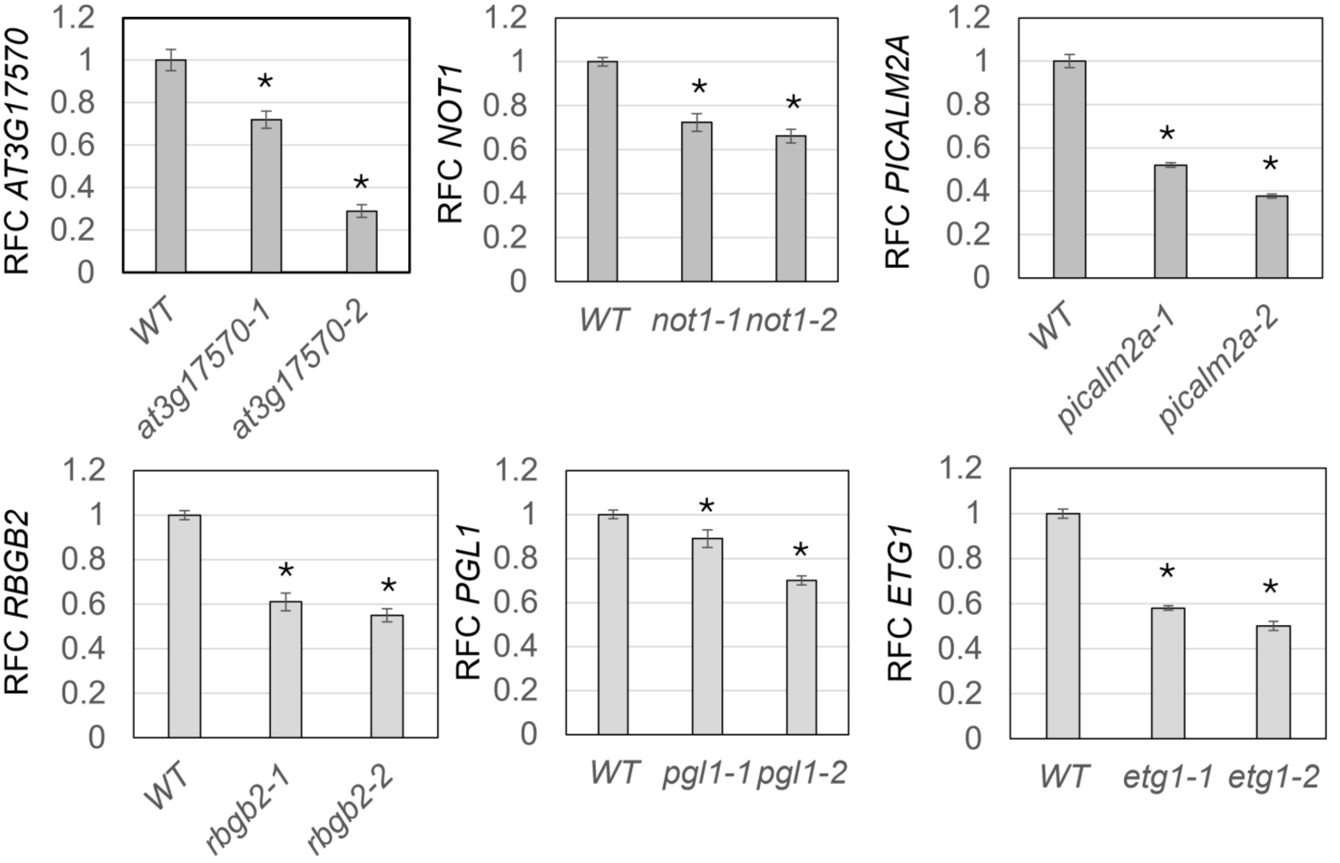


**Fig. S5 Gene expression levels of knockdown lines**. T-DNA insertion events in the promoter, 5'-UTR or introns were tested for expression level reduction. The confirmed homozygous mutants: *at3g17570* (SALK_100641C; promoter), *not1* (SALK_114704C, 5'-UTR), *picalm2a* (SALK_133151C; 5'-UTR), *rbgb2* (SALK_109798C; intron), *pgl1* (SALK_033902C; intron), and *etg1* (SALK_071046C; intron) were grown in ¼-Hoagland's media supplemented with 1 mM $NaHCO_3$, pH 8.0 for 14 days. The seedlings were harvested, followed by RNA isolation and quantitative PCR (qPCR) analysis with primers listed in Table S2. Fold change in expression levels of two independent knockdown lines with respect to the wild type (WT) is represented. *AtUBQ1* was used as an internal control for normalization of expression levels, and fold changes were calculated using the $2^{-\Delta\Delta Ct}$ method. Asterisks indicate significant differences between WT and knockdown lines (Student's *t*-test, *N* = 3 biological replicates).

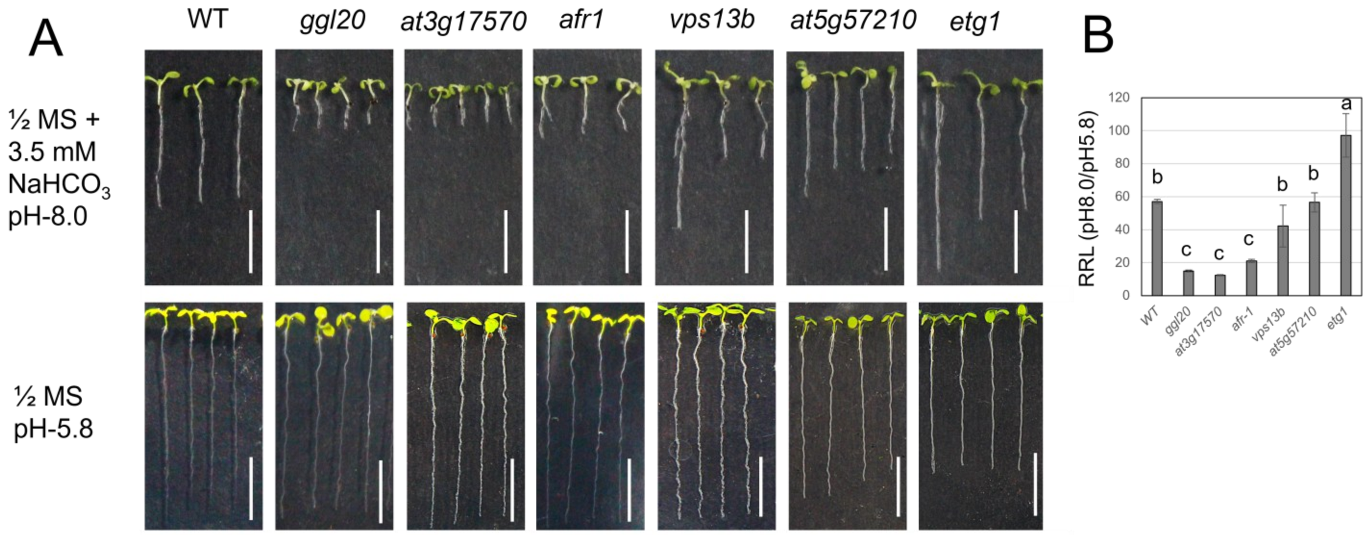


**Fig. S6 Alkalinity stress tolerance of T-DNA insertion lines in solid media**. (**A**) Wild-type (Col-0) and T-DNA mutants: *ggl20*, *at3g17570*, *afr1, vps13b*, *at5g57210*, and *etg1* were grown in ½ MS media with 1.5% sucrose alone (pH 5.8) or that supplemented with 3.5 mM $NaHCO_3$ (pH 8.0) for 14 days and photographed. Scale bar = 1 cm. (**B**) Relative root length (RRL) was calculated by dividing the average root length of bicarbonate-treated plants by the control plants and expressed as a percentage. The average RRL of 20 plants is shown with standard error. Significant differences ($P < 0.05$; Tukey's test) are shown by different lowercase letters above the bars.

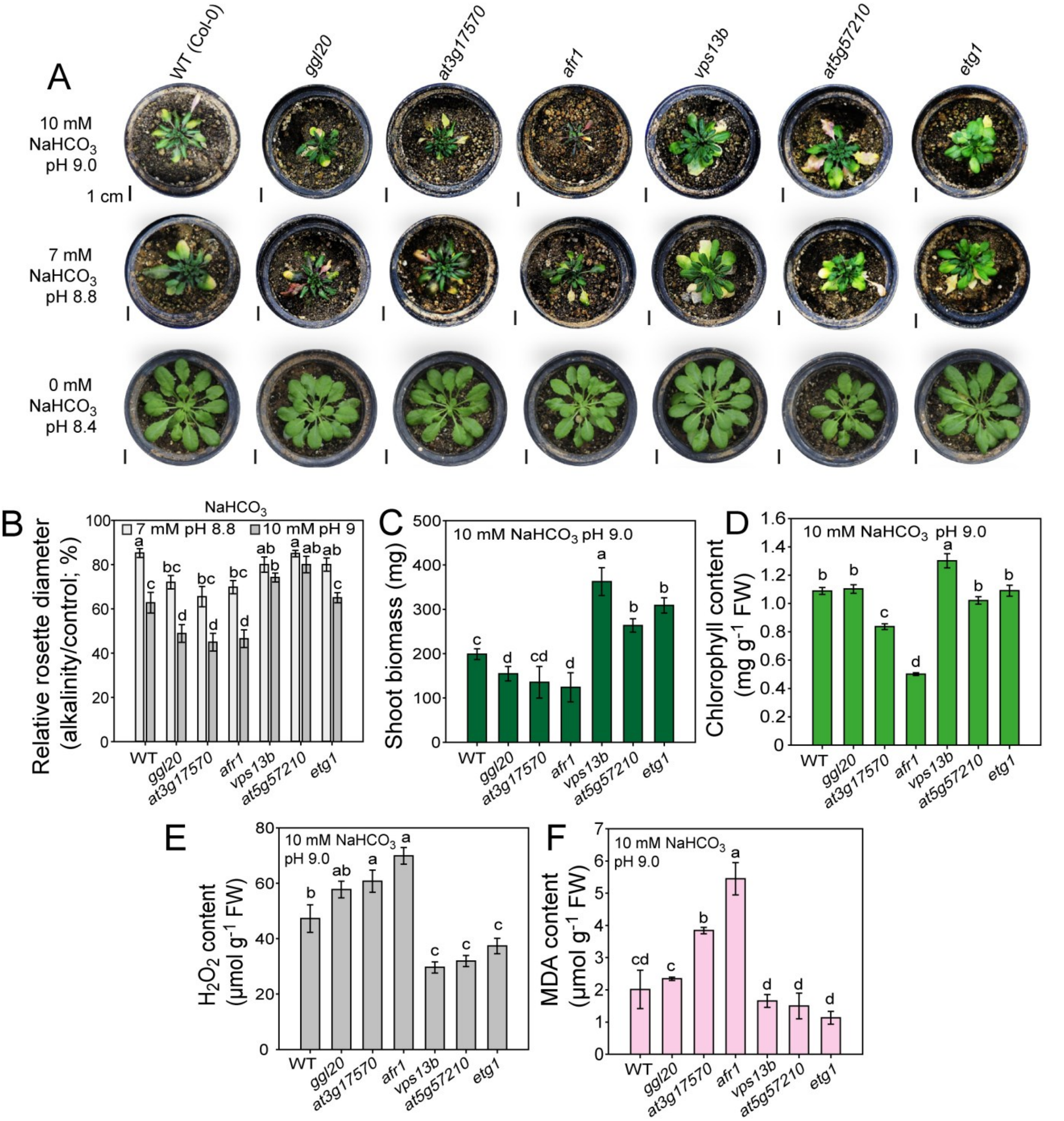


**Fig. S7 Alkalinity stress tolerance of T-DNA insertion lines in soil**. **(A)** Alkalinity tolerance test in soil. T-DNA insertion mutants and WT grown in a 1:2:1 mixture of desert soil, soil rite, and vermiculite, watered with deionized water (final pH 8.4), or the same soil mixtures watered with 7 mM $NaHCO_3$ (pH 8.8) and 10 mM $NaHCO_3$ (pH 9.0) are shown. Scale: 1 cm. The graphs show relative rosette diameter (alkaline soil vs. pH 6.5) **(B)**, shoot biomass **(C)**, together with hydrogen peroxide **(D)**, chlorophyll **(E)**, and malonaldehyde **(F)**, measured spectrophotometrically. The graphs show mean ± standard error; $N = 10$. Different letters above the bars indicate significant differences (Tukey's HSD test; $P < 0.05$).

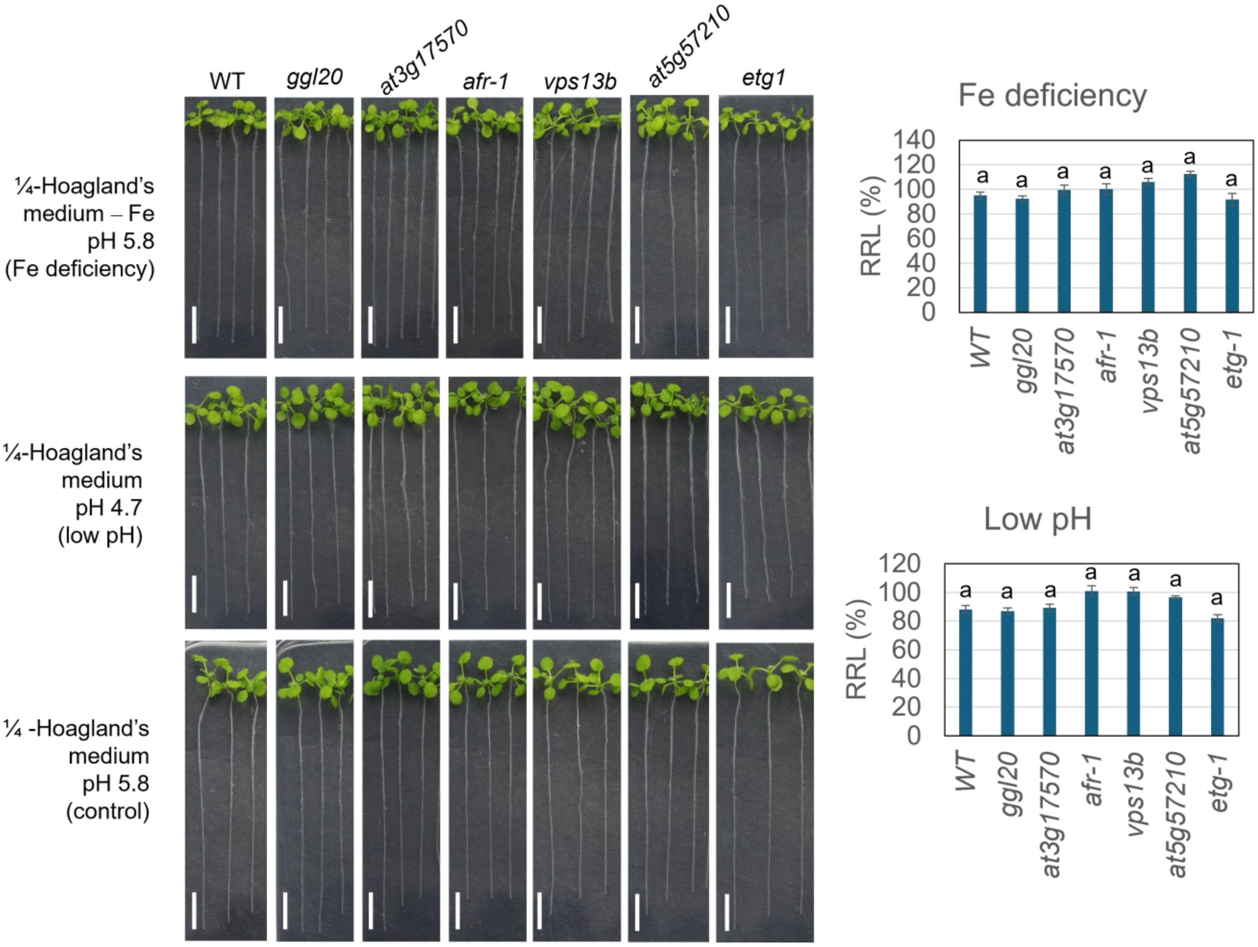


**Fig. S8 Low pH and iron deficiency stress tolerance of T-DNA insertion lines (A)** Wild-type (Col-0) and T-DNA mutants: *ggl20*, *at3g17570*, *afr1, vps13b*, *at5g57210*, and *etg1* were grown in ¼-Hoagland's solution, pH 5.8 (control) or ¼-Hoagland's solution, pH 4.7 (adjusted with HCl) or a modified ¼-Hoagland's solution without iron (pH 5.8) in hydroponics for 14 days, transferred to agar plates and photographed. Scale bar = 1 cm. **(B)** Relative root length (RRL) was calculated by dividing the average root length of low pH or Fe-deficiency-treated plants by the control plants and expressed as a percentage. The average RRL of 20 plants is shown with standard error. Insignificant differences ($P < 0.05$; Tukey's test) are shown by the same lowercase letters above the bars

**Fig. S9 Rhizosphere acidification of T-DNA insertion lines (A)** Wild-type (Col-0) and T-DNA mutants: *ggl20*, *at3g17570*, *afr1, vps13b*, *at5g57210*, and *etg1* were grown in ½-MS media with 1.5% sucrose (pH 5.8) for 10 days, transferred to ½-MS media supplemented with 3.5 mM $NaHCO_3$ (pH 8.0) for 72 h, and again transferred to water agar plates containing 0.006% bromocresol purple (pH 6.0) and photographed after 48 h. Scale bar = 1 cm. The color scale indicates bromocresol purple color changes corresponding to media pH (yellow at pH < 5.2; purple at pH > 6.8).

#### Table S1. List of ecotypes used in the study

| Ecotype | High pH tolerance (RRL; %) | Latitude | Longitude | Country | Region | Soil pH (0-10 cm depth) | Soil organic carbon (0-30 cm depth) tons/hectare | Soil $CaCO_3$ (0-30 cm depth) (g/Kg) |
|---|---|---|---|---|---|---|---|---|
| Sij-2 | 12.8 | 41.45 | 70.05 | Uzbekistan | Central Asia | 6.6 | 111 | 80 |
| Is-0 | 14.9 | 50.5 | 7.5 | Germany | Western Europe | 5.5 | 120 | 6 |
| Pro-0 | 15.1 | 43.25 | -6 | Portugal | Southern Europe | 6.3 | 103 | 45 |
| Buckhorn Pass | 15.2 | 41.3599 | -122.755 | United States of America | Northern America | 6.9 | 76 | 11.5 |
| Aa-0 | 16.8 | 50.9167 | 9.57073 | Germany | Western Europe | 5.9 | 103 | 152.5 |
| Ru3.1-31 | 18.2 | 48.56 | 9.16 | Germany | Western Europe | 6.1 | 83 | 5 |
| Nz-1 | 18.3 | -37.7871 | 175.283 | New Zealand | Australia and New Zealand | 6.9 | 116 | 24 |
| CIBC-5 | 19.7 | 51.4083 | -0.6383 | United Kingdom | Northern Europe | 5.7 | 96 | 31.5 |
| Go-0 | 20.1 | 51.5338 | 9.9355 | Germany | Western Europe | 6.7 | 118 | 60 |
| Hh-0 | 20.5 | 54.4175 | 9.88682 | Germany | Western Europe | 6.5 | 113 | 36.5 |
| Ba-1 | 21.1 | 56.5459 | -4.79821 | United Kingdom | Northern Europe | 5.1 | 237 | 31 |
| Fr-4 | 22.1 | 50.1102 | 8.6822 | Germany | Western Europe | 7.0 | | 105 |
| Tha-1 | 22.2 | 52.08 | 4.3 | Netherlands | Western Europe | 6.9 | 102 | 50 |
| Pla-1 | 22.3 | 41.5 | 2.25 | Spain | Southern Europe | 6.4 | 79 | 165 |
| Baa-1 | 22.7 | 51.3333 | 6.1 | Belgium | Western Europe | 5.6 | 62 | 11.5 |
| Si-0 | 23.0 | 50.8738 | 8.02341 | Germany | Western Europe | 6.3 | 122 | 25 |
| Kz-9 | 23.2 | 49.5 | 73.1 | Kazakhstan | Central Asia | 7.3 | 95 | 12.5 |
| Zdr-6 | 23.2 | 49.3853 | 16.2544 | Czech Republic | Eastern Europe | 6.3 | 68 | 136 |
| Bsch-0 | 23.3 | 50.0167 | 8.6667 | Germany | Western Europe | 5.8 | 108 | 23.5 |
| Rovero-1 | 24.2 | 46.2543 | 11.167 | Switzerland | Western Europe | 6.5 | 86 | 26.5 |
| Do-0 | 24.3 | 50.7224 | 8.2372 | Germany | Western Europe | 5.6 | 100 | 25 |
| Kro-0 | 25.1 | 50.0742 | 8.96617 | Germany | Western Europe | 6.8 | 75 | 194 |
| Pt-0 | 25.3 | 53.476 | 10.6065 | Germany | Western Europe | 6.0 | 81 | 27.5 |
| Uod-1 | 25.7 | 48.3 | 14.45 | Czech Republic | Eastern Europe | 5.2 | 81 | 137 |
| Jm-1 | 25.8 | 49 | 15 | Czech Republic | Eastern Europe | 4.9 | 149 | 147.5 |
| Sorbo | 26.5 | 38.35 | 68.48 | Uzbekistan | Central Asia | 7.4 | 41 | 58.5 |
| Rennes-1 | 26.7 | 48.5 | -1.41 | France | Western Europe | 5.9 | 73 | 10 |
| Pla-3 | 26.8 | 41.5 | 2.25 | Spain | Southern Europe | 6.4 | 79 | 165 |
| Lz-0 | 26.9 | 46 | 3.3 | France | Western Europe | 7.3 | 71 | 10 |
| Chi-1 | 27.1 | 53.7502 | 34.7361 | Russian Federation | Eastern Europe | 6.5 | 113 | 10 |
| Lago-1 | 27.1 | 39.18 | 16.26 | Italy | Southern Europe | 6.0 | 91 | 95 |
| Star-8 | 27.2 | 48.43 | 8.82 | Germany | Western Europe | 7.2 | 83 | 2.5 |
| Zu-1 | 27.4 | 47.3667 | 8.55 | Switzerland | Western Europe | 6.5 | 128 | 35 |
| Tu-0 | 27.6 | 45 | 7.5 | France | Western Europe | 7.4 | 75 | 15 |
| Pn-0 | 28.5 | 48.0653 | -2.96591 | France | Western Europe | 6.2 | 77 | 40 |
| Old-1 | 28.9 | 53.1667 | 8.2 | Germany | Western Europe | 5.7 | 157 | 37.5 |
| Alst-1 | 29.2 | 54.8 | -2.4333 | United Kingdom | Northern Europe | 5.6 | 163 | 39.5 |
| Ty-0 | 29.4 | 56.4278 | -5.23439 | United Kingdom | Northern Europe | 5.1 | 215 | 106 |
| Slavi-1 | 29.5 | 41.43 | 23.65 | Greece | Southern Europe | 5.2 | 109 | 1.5 |
| Nd-1 | 29.6 | 50 | 10 | Germany | Western Europe | 7.2 | 74 | 21.5 |
| Se-0 | 29.8 | 38.3333 | -3.53333 | Spain | Southern Europe | 6.6 | 48 | 97.5 |
| Je-0 | 30.0 | 50.927 | 11.587 | Germany | Western Europe | 6.9 | 69 | 175 |
| Tu-Scha-9 | 30.8 | 48.53 | 9.05 | Germany | Western Europe | 6.8 | 85 | 27.5 |

| | | | | | | | | |
|---|---|---|---|---|---|---|---|---|
| In-0 | 31.1 | 47.5 | 11.5 | Germany | Western Europe | 5.8 | 163 | 4.5 |
| Sei-0 | 31.1 | 46.5438 | 11.5614 | Italy | Southern Europe | 6.4 | 139 | 11.5 |
| Old-2 | 31.3 | 53.1667 | 8.2 | Germany | Western Europe | 5.7 | 157 | 37.5 |
| Timpo-1 | 31.3 | 39.27 | 16.27 | Italy | Southern Europe | 7.1 | 75 | 95 |
| Ciste-2 | 31.7 | 41.62 | 12.87 | Italy | Southern Europe | 6.9 | 76 | 12.5 |
| Po-1 | 32.2 | 50.7167 | 7.1 | Germany | Western Europe | 6.3 | 89 | 21 |
| Gel-1 | 32.4 | 51.0167 | 5.86667 | Netherlands | Western Europe | 6.6 | 69 | 220 |
| Ra-0 | 32.4 | 46 | 3.3 | France | Western Europe | 7.3 | 71 | 10 |
| Rsch-0 | 32.7 | 45.5333 | 4.85 | France | Western Europe | 6.2 | 75 | 40 |
| Nw-4 | 32.8 | 50.5 | 8.5 | Germany | Western Europe | 6.6 | 61 | 80 |
| Rsch-4 | 32.8 | 56.3 | 34 | Russian Federation | Eastern Europe | 5.8 | 122 | 130 |
| Wt-5 | 33.0 | 52.3 | 9.3 | Germany | Western Europe | 6.7 | 83 | 13 |
| Wt-1 | 33.1 | 52.3 | 9.3 | Germany | Western Europe | 6.7 | 83 | 13 |
| Me-0 | 33.2 | 51.9183 | 10.1138 | Germany | Western Europe | 6.7 | 69 | 55 |
| Sij-1 | 33.2 | 41.45 | 70.05 | Uzbekistan | Central Asia | 6.6 | 111 | 80 |
| Sp-0 | 33.8 | 52.5339 | 13.181 | Germany | Western Europe | 6.9 | 84 | 37.5 |
| Wa-1 | 33.9 | 52.3 | 21 | Poland | Eastern Europe | 6.4 | 77 | 4 |
| Np-0 | 33.9 | 52.6969 | 10.981 | Germany | Western Europe | 6.0 | 87 | 80 |
| Est-1 | 33.9 | 58.3 | 25.3 | Estonia | Northern Europe | 6.1 | 114 | 100 |
| Sg-1 | 34.0 | 47.6667 | 9.5 | Switzerland | Western Europe | 6.6 | 83 | 10 |
| Sha | 34.0 | 37.29 | 71.3 | Tajikistan | Central Asia | 6.5 | 217 | 44 |
| Ma-2 | 34.1 | 50.8167 | 8.7667 | Germany | Western Europe | 6.1 | 102 | 152.5 |
| Ga-2 | 34.2 | 50.3 | 8 | Germany | Western Europe | 5.4 | 99 | 105 |
| Kl-5 | 34.2 | 50.95 | 6.9666 | Germany | Western Europe | 6.8 | 101 | 35 |
| El-0 | 34.4 | 51.5105 | 9.68253 | Germany | Western Europe | 5.5 | 98 | 60 |
| Bolin-1 | 34.4 | 44.46 | 25.74 | Romania | Eastern Europe | 7.1 | 67 | 33.5 |
| Pr-0 | 34.6 | 50.1448 | 8.60706 | Germany | Western Europe | 7.1 | 69 | 105 |
| Vezzano-2 | 34.6 | 46.6297 | 10.817 | Switzerland | Western Europe | 6.3 | 82 | 26.5 |
| Pog-0 | 34.7 | 49.2655 | -123.206 | Canada | Northern America | 6.4 | 175 | 130 |
| Or-0 | 34.8 | 50.3827 | 8.01161 | Germany | Western Europe | 6.9 | 89 | 105 |
| Rd-0 | 34.9 | 50.5 | 8.5 | Germany | Western Europe | 6.6 | 61 | 80 |
| Co-3 | 35.1 | 40.12 | -8.25 | Portugal | Southern Europe | 7.3 | 67 | 90 |
| Bch-1 | 35.5 | 49.5166 | 9.3166 | Germany | Western Europe | 6.7 | 65 | 21.5 |
| Ak-1 | 35.5 | 48.0683 | 7.62551 | France | Western Europe | 6.5 | 112 | 40 |
| Borsk-2 | 35.5 | 53.04 | 51.75 | Russian Federation | Eastern Europe | 6.5 | 146 | 5 |
| Ts-1 | 35.6 | 41.7194 | 2.93056 | France | Western Europe | 6.5 | 82 | 1.5 |
| Da-0 | 35.8 | 49.8724 | 8.65081 | Germany | Western Europe | 6.7 | 87 | 23.5 |
| Uod-7 | 35.8 | 48.3 | 14.45 | Czech Republic | Eastern Europe | 5.2 | 81 | 137 |
| Petro-1 | 35.9 | 44.34 | 21.46 | Romania | Eastern Europe | 6.8 | 59 | 33.5 |
| Ey1.5-2 | 36.2 | 48.43 | 8.77 | Germany | Western Europe | 6.5 | 97 | 2.5 |
| NFA-8 | 36.5 | 51.4083 | -0.6383 | United Kingdom | Northern Europe | 5.7 | 96 | 31.5 |
| Toufl-1 | 36.7 | 31.47 | -7.42 | Morocco | Northern Africa | 6.7 | 86 | 22.5 |
| Br-0 | 36.8 | 49.2 | 16.6166 | Czech Republic | Eastern Europe | 6.9 | 63 | 13.5 |
| Lo-2 | 36.9 | 47.6166 | 7.6666 | Germany | Western Europe | 7.4 | 110 | 28.5 |
| Tu-SB30-3 | 36.9 | 48.53 | 9.06 | Germany | Western Europe | 6.9 | 87 | 27.5 |
| Kin-0 | 36.9 | 44.46 | -85.37 | United States of America | Northern America | 5.5 | 139 | 133.5 |
| Zdr-1 | 37.2 | 49.3853 | 16.2544 | Czech Republic | Eastern Europe | 6.3 | 68 | 136 |
| Kl-0 | 37.3 | 50.95 | 6.9666 | Germany | Western Europe | 6.8 | 101 | 35 |
| Di-1 | 37.3 | 47 | 5 | France | Western Europe | 6.9 | 67 | 15 |
| HKT2-4 | 37.3 | 48.14 | 9.4 | Germany | Western Europe | 6.0 | 95 | 27.5 |

| | | | | | | | | |
|---|---|---|---|---|---|---|---|---|
| Ep-0 | 37.9 | 50.1721 | 8.38912 | Germany | Western Europe | 6.7 | 110 | 105 |
| Wl-0 | 37.9 | 47.9299 | 10.8134 | Switzerland | Western Europe | 5.5 | 111 | 10 |
| Pf-0 | 38.2 | 48.5479 | 9.11033 | Germany | Western Europe | 6.8 | 86 | 27.5 |
| Ven-1 | 38.3 | 52.0333 | 5.55 | Netherlands | Western Europe | 6.2 | 92 | 40.5 |
| Bsch-2 | 38.7 | 50.0167 | 8.6667 | Germany | Western Europe | 5.8 | 108 | 23.5 |
| An-1 | 38.8 | 51.2167 | 4.4 | Belgium | Western Europe | 6.9 | 70 | 39 |
| Hs-0 | 39.1 | 52.24 | 9.44 | Germany | Western Europe | 6.0 | 105 | 47.5 |
| Ga-0 | 39.4 | 50.3 | 8 | Germany | Western Europe | 5.4 | 99 | 105 |
| Boot-1 | 39.6 | 54.4 | -3.2667 | United Kingdom | Northern Europe | 5.4 | 230 | 55 |
| Pa-1 | 39.6 | 38.07 | 13.22 | Italy | Southern Europe | 6.8 | 108 | 72 |
| Nw-0 | 39.8 | 50.5 | 8.5 | Germany | Western Europe | 6.6 | 61 | 80 |
| Ws-2 | 39.9 | 52.3 | 30 | Belarus | Eastern Europe | 6.1 | 120 | 48.5 |
| Yeg-1 | 40.0 | 39.8692 | 45.3622 | Armenia | Western Asia | 6.6 | 99 | 79 |
| Lag2-2 | 40.0 | 41.8296 | 46.2831 | Georgia | Western Asia | 7.1 | 87 | 192 |
| Utrecht | 40.0 | 52.0918 | 5.1145 | Netherlands | Western Europe | 6.8 | 101 | 32.5 |
| Omo-2-3 | 40.5 | 56.1509 | 15.7735 | Sweden | Northern Europe | 5.8 | 129 | 65 |
| Ag-0 | 40.8 | 45 | 1.3 | France | Western Europe | 5.7 | 85 | 15 |
| Nok-3 | 40.9 | 52.24 | 4.45 | Netherlands | Western Europe | 7.1 | 92 | 130 |
| Aitba 2 | 41.1 | 31.48 | -7.45 | Morocco | Northern Africa | 7.4 | 49 | 22.5 |
| Ge-1 | 41.4 | 46.5 | 6.08 | France | Western Europe | 5.6 | 170 | 21 |
| Hn-0 | 41.4 | 51.3472 | 8.28844 | Germany | Western Europe | 5.9 | 133 | 22 |
| TueV-13 | 41.5 | 48.52 | 9.05 | Germany | Western Europe | 6.4 | 88 | 27.5 |
| Pa-3 | 41.6 | 38.07 | 13.22 | Italy | Southern Europe | 6.8 | 108 | 72 |
| Kelsterbach-4 | 42.0 | 50.0667 | 8.5333 | Germany | Western Europe | 6.7 | 73 | 105 |
| Nie1-2 | 42.1 | 48.52 | 8.8 | Germany | Western Europe | 6.3 | 100 | 21 |
| Nd-0 | 42.2 | 50 | 10 | Germany | Western Europe | 7.2 | 74 | 21.5 |
| Chat-1 | 42.3 | 48.0717 | 1.33867 | France | Western Europe | 7.1 | 56 | 40 |
| Mh-0 | 42.5 | 50.95 | 7.5 | Germany | Western Europe | 6.2 | 122 | 6 |
| Ove-0 | 42.5 | 53.3422 | 8.42255 | Germany | Western Europe | 5.5 | 144 | 50 |
| Gr-1 | 42.5 | 47 | 15.5 | Austria | Western Europe | 6.9 | 78 | 76.5 |
| Stepn-1 | 42.7 | 54.06 | 60.48 | Russian Federation | Eastern Europe | 7.2 | 126 | 38.5 |
| Db-0 | 42.7 | 50.3055 | 8.324 | Germany | Western Europe | 6.1 | 104 | 12.5 |
| Bs-1 | 42.8 | 47.5 | 7.5 | Switzerland | Western Europe | 6.8 | 79 | 87.5 |
| Jl-3 | 43.1 | 49.2 | 16.6166 | Czech Republic | Eastern Europe | 6.9 | 63 | 13.5 |
| Apost-1 | 44.0 | 39.01 | 16.47 | Italy | Southern Europe | 5.8 | 86 | 95 |
| No-0 | 44.0 | 51.0581 | 13.2995 | Czech Republic | Eastern Europe | 6.4 | 98 | 135 |
| Tscha-1 | 44.5 | 47.0748 | 9.9042 | Switzerland | Western Europe | 5.9 | 120 | 23 |
| Ka-0 | 44.6 | 47 | 14 | Italy | Southern Europe | 5.4 | 140 | 37.5 |
| Mir-0 | 44.8 | 44 | 12.37 | Italy | Southern Europe | 7.0 | 62 | 20 |
| Kr-0 | 44.8 | 51.3317 | 6.55934 | Germany | Western Europe | 6.5 | 94 | 22.5 |
| Bd-0 | 44.8 | 52.4584 | 13.287 | Germany | Western Europe | 5.8 | 123 | 12.5 |
| Kondara | 44.9 | 38.48 | 68.49 | Tajikistan | Central Asia | 7.5 | 42 | 132.5 |
| NFA-10 | 45.0 | 51.4083 | -0.6383 | United Kingdom | Northern Europe | 5.7 | 96 | 31.5 |
| Ct-1 | 45.1 | 37.3 | 15 | Italy | Southern Europe | 7.5 | 53 | 72 |
| Moran-1 | 45.2 | 39.83 | 16.17 | Italy | Southern Europe | 7.3 | 78 | 31 |
| Ha-0 | 45.2 | 52.3721 | 9.73569 | Germany | Western Europe | 7.1 | 82 | 57.5 |
| Dra-2 | 45.2 | 49.4167 | 16.2667 | Czech Republic | Eastern Europe | 5.9 | 82 | 136 |
| Voeran-1 | 45.5 | 46.36 | 11.23 | Switzerland | Western Europe | 6.2 | 141 | 26.5 |
| Ob-1 | 45.5 | 50.2 | 8.5833 | Germany | Western Europe | 6.7 | 71 | 105 |
| Bla-1 | 45.6 | 41.6833 | 2.8 | France | Western Europe | 6.5 | 80 | 1.5 |

| | | | | | | | | |
|---|---|---|---|---|---|---|---|---|
| Ting-1 | 45.6 | 56.5 | 14.9 | Sweden | Northern Europe | 5.1 | 175 | 45 |
| Mammo-1 | 45.8 | 38.36 | 16.23 | Italy | Southern Europe | 6.7 | 74 | 95 |
| Leb-3 | 45.8 | 51.65 | 80.82 | Russian Federation | Eastern Europe | 7.1 | 116 | 72.5 |
| Cit-0 | 45.8 | 43.3779 | 2.54038 | France | Western Europe | 6.7 | 90 | 10 |
| Ciste-1 | 45.8 | 41.62 | 12.87 | Italy | Southern Europe | 6.9 | 76 | 12.5 |
| Bu-0 | 46.2 | 50.5 | 9.5 | Germany | Western Europe | 5.6 | 103 | 155 |
| Yo-0 | 46.5 | 37.45 | -119.35 | United States of America | Northern America | 5.7 | 91 | 84 |
| Lip-0 | 46.6 | 50 | 19.3 | Poland | Eastern Europe | 6.2 | 75 | 5 |
| Hi-0 | 46.8 | 52 | 5 | Netherlands | Western Europe | 6.3 | 159 | 90 |
| Wc-1 | 47.0 | 52.6 | 10.0667 | Germany | Western Europe | 5.9 | 98 | 15 |
| Bor-4 | 47.1 | 49.4013 | 16.2326 | Czech Republic | Eastern Europe | 6.0 | 73 | 141 |
| Kas-2 | 47.2 | 35 | 77 | Jammu and Kashmir | Eastern Asia | 7.7 | | 91.5 |
| Mammo-2 | 47.5 | 38.38 | 16.22 | Italy | Southern Europe | 6.2 | 117 | 95 |
| HR-5 | 47.6 | 51.4083 | -0.6383 | United Kingdom | Northern Europe | 5.7 | 96 | 31.5 |
| Blh-1 | 47.6 | 48 | 19 | Hungary | Eastern Europe | 5.3 | 103 | 132.5 |
| Sq-8 | 47.7 | 51.4083 | -0.6383 | United Kingdom | Northern Europe | 5.7 | 96 | 31.5 |
| Kil-0 | 48.0 | 55.6395 | -5.66364 | United Kingdom | Northern Europe | 4.9 | 192 | 78 |
| Na-1 | 48.1 | 47.5 | 1.5 | France | Western Europe | 5.9 | 58 | 12.5 |
| Fi-1 | 48.1 | 50.5 | 8.0167 | Germany | Western Europe | 6.3 | 82 | 105 |
| Altenb-2 | 48.7 | 46.3716 | 11.2376 | Switzerland | Western Europe | 6.4 | 109 | 26.5 |
| Ang-0 | 48.8 | 50.3 | 5.3 | Belgium | Western Europe | 5.6 | 106 | 70.5 |
| KNO-18 | 48.9 | 41.2816 | -86.621 | United States of America | Northern America | 6.4 | 77 | 68.5 |
| Wal-HasB-4 | 48.9 | 48.6 | 9.19 | Germany | Western Europe | 6.4 | 80 | 5 |
| Col-0 | 49.5 | 38.3 | -92.3 | United States of America | Northern America | 6.3 | 66 | 68 |
| Dr-0 | 49.9 | 51.051 | 13.7336 | Czech Republic | Eastern Europe | 7.1 | 59 | 132.5 |
| Co-2 | 50.0 | 40.12 | -8.25 | Portugal | Southern Europe | 7.3 | 67 | 90 |
| Li-2:1 | 50.1 | 50.3833 | 8.0666 | Germany | Western Europe | 6.9 | 96 | 105 |
| Com-1 | 50.4 | 49.416 | 2.823 | France | Western Europe | 6.8 | 58 | 41 |
| Ll-0 | 50.6 | 41.59 | 2.49 | France | Western Europe | 5.6 | 81 | 1.5 |
| Lecho-1 | 50.6 | 41.43 | 23.5 | Bulgaria | Eastern Europe | 6.8 | 66 | 12.5 |
| Mrk-0 | 50.7 | 49 | 9.3 | Germany | Western Europe | 7.1 | 67 | 21.5 |
| RRS-7 | 51.1 | 41.5609 | -86.4251 | United States of America | Northern America | 6.2 | 106 | 196.5 |
| Ms-0 | 51.2 | 55.7522 | 37.6322 | Russian Federation | Eastern Europe | 6.1 | 138 | 100 |
| Stepn-2 | 51.4 | 54.09 | 60.46 | Russian Federation | Eastern Europe | 6.7 | 138 | 38.5 |
| Mnz-0 | 51.4 | 50.001 | 8.26664 | Germany | Western Europe | 6.7 | 84 | 33.5 |
| Jm-0 | 51.6 | 49 | 15 | Czech Republic | Eastern Europe | 4.9 | 149 | 147.5 |
| Pu2-23 | 51.6 | 49.42 | 16.36 | Czech Republic | Eastern Europe | 5.4 | 95 | 136 |
| Koch-1 | 52.1 | 50.3553 | 29.3244 | Ukraine | Eastern Europe | 5.6 | 94 | 151 |
| Da(1)-12 | 52.49 | N/A | N/A | Ghana | Western Africa | 7.0 | | 105.5 |
| Sapporo-0 | 52.7 | 43.0553 | 141.346 | Japan | Eastern Asia | 6.5 | 157 | 19 |
| Gy-0 | 53.0 | 49 | 2 | France | Western Europe | 5.4 | 85 | 20 |
| Dra-1 | 53.1 | 49.4167 | 16.2667 | Czech Republic | Eastern Europe | 5.9 | 82 | 136 |
| En-1 | 53.2 | 50 | 8.5 | Germany | Western Europe | 5.3 | 121 | 23.5 |
| Pi-2 | 53.2 | 47.04 | 10.51 | Switzerland | Western Europe | 5.7 | 125 | 10 |
| Pna-17 | 53.4 | 42.0945 | -86.3253 | United States of America | Northern America | 6.2 | 87 | 151.5 |
| Blh-2 | 53.6 | 48 | 19 | Hungary | Eastern Europe | 5.3 | 103 | 132.5 |
| Mer-6 | 53.7 | 38.92 | -6.34 | Spain | Southern Europe | 7.7 | 39 | 97.5 |
| TuWa1-2 | 53.8 | 48.53 | 9.04 | Germany | Western Europe | 7.1 | 84 | 5 |
| Bozen-1 | 54.4 | 46.513 | 11.331 | Switzerland | Western Europe | 6.3 | 90 | 26.5 |
| Ob-0 | 54.5 | 50.2 | 8.5833 | Germany | Western Europe | 6.7 | 71 | 105 |

| | | | | | | | | |
|---|---|---|---|---|---|---|---|---|
| Rou-0 | 54.5 | 49.4424 | 1.09849 | France | Western Europe | 7.2 | 77 | 7.5 |
| Pla-0 | 54.9 | 41.5 | 2.25 | Spain | Southern Europe | 6.4 | 79 | 165 |
| Mz-0 | 55.2 | 50.3 | 8.3 | Germany | Western Europe | 6.2 | 107 | 35 |
| Ei-2 | 55.2 | 50.3 | 6.3 | Germany | Western Europe | 6.2 | 139 | 13.5 |
| Bu-4 | 57.1 | 49 | 0.5 | France | Western Europe | 6.4 | 62 | 12.5 |
| Cnt-1 | 58.2 | 51.3 | 1.1 | United Kingdom | Northern Europe | 6.8 | 69 | 155 |
| Lp2-2 | 59.0 | 49.38 | 16.81 | Czech Republic | Eastern Europe | 5.8 | 65 | 149 |
| Bak-7 | 60.4 | 41.7942 | 43.4767 | Georgia | Western Asia | 6.1 | 98 | 45 |
| Sij-4 | 60.8 | 41.45 | 70.05 | Uzbekistan | Central Asia | 6.6 | 111 | 80 |
| Gie-0 | 61.3 | 50.584 | 8.67825 | Germany | Western Europe | 7.1 | 87 | 80 |
| Ca-0 | 61.6 | 50.2981 | 8.26607 | Germany | Western Europe | 7.2 | 82 | 35 |
| RRS-10 | 61.9 | 41.5609 | -86.4251 | United States of America | Northern America | 6.2 | 106 | 196.5 |
| Ge-0 | 62.6 | 46.5 | 6.08 | France | Western Europe | 5.6 | 170 | 21 |
| Hl-3 | 62.8 | 52.1444 | 9.37827 | Germany | Western Europe | 5.8 | 95 | 47.5 |
| Fei-0 | 63.4 | 40.92 | -8.54 | Portugal | Southern Europe | 5.9 | 92 | 107.5 |
| Bor-1 | 63.5 | 49.4013 | 16.2326 | Czech Republic | Eastern Europe | 6.0 | 73 | 141 |
| Nok-1 | 64.0 | 52.24 | 4.45 | Netherlands | Western Europe | 7.1 | 92 | 130 |
| Vie-0 | 64.4 | 42.63 | 0.76 | France | Western Europe | 5.8 | 156 | 40 |
| UK-1 | 65.1 | 48.0333 | 7.7667 | France | Western Europe | 6.7 | 89 | 40 |
| Castelfed-4 | 66.8 | 46.3378 | 11.2928 | Switzerland | Western Europe | 6.6 | 105 | 107.5 |
| La-1 | 69.8 | 52.7333 | 15.2333 | Poland | Eastern Europe | 6.5 | 85 | 60 |
| Ema-1 | 70.1 | 51.3 | 0.5 | United Kingdom | Northern Europe | 6.4 | 77 | 34 |
| Gu-0 | 76.1 | 50.3 | 8 | Germany | Western Europe | 5.4 | 99 | 105 |
| Lc-0 | 76.2 | 57 | -4 | United Kingdom | Northern Europe | 5.0 | 193 | 31 |
| Li-6 | 76.5 | 50.3833 | 8.0666 | Germany | Western Europe | 6.9 | 96 | 105 |

**Table S2. Oligonucleotide sequences used in the study**

| AGI code | Primer name | Sequence (5'--> 3') |
|---|---|---|
| **Primers for ELP analysis** | | |
| AT3G52590 | *UBQ1* _RT_Fw | TCGTAAGTACAATCAGGATAAGATG |
| | *UBQ1* _RT_Rv | CACTGAAACAAGAAAAACAAACCCT |
| AT1G02070 | AT1G02070_RT_Fw | ACCGTCACGATCAAGTAGCCA |
| | AT1G02070_RT_Rv | GCTTGTACCTTCGTGTGGCC |
| AT1G75060 | AT1G75060_RT_Fw | GGATGAGCTTCAGGTTATTGTGGG |
| | AT1G75060_RT_Rv | TGTTTCTGGCTTCTTTGGTCTTCC |
| AT3G26420 | AT3G26420_ RT_Fw | CGCTCCCGTGGTTTTGGATT |
| | AT3G26420_RT_Rv | TATCTCTGCCTGCACCACCC |
| AT3G26430 | AT3G26430_RT_Fw | AGCCATGGTGCCAATTTCGC |
| | AT3G26430_RT_Rv | TTGGACTCACACCGCTCTGA |
| AT3G60500 | AT3G60500_RT_Fw | TGAACGGCTCTGGAAATGCG |
| | AT3G60500_RT_Rv | ATCCTTCTTCCCTAATTCTCCGGT |
| AT5G57210 | AT5G57210_RT_Fw | TCCCAAACCCAGACACCGAT |
| | AT5G57210_RT_Rv | TCCGGGATAGAAAGAGGCAGA |
| AT1G02080 | AT1G02080_RT_Fw | GCCTTCATCAGATGTGCGCC |
| | AT1G02080_RT_Rv | GACCCAACCTCCCGAAACCA |
| AT1G05640 | AT1G05640_RT_Fw | AGCTCATGTGGTTGGCTTGT |
| | AT1G05640_RT_Rv | GCCAGCCAAATGTCCTCCTT |
| AT1G05650 | AT1G05650_RT_Fw | AACCCAAGTGACGTACAAGAACA |
| | AT1G05650_RT_Rv | TGTGATTCCCGTGCATGGATT |
| AT4G16260 | AT4G16260_RT_Fw | ATCTTGCGTCGCGTGTGAGA |
| | AT4G16260_RT_Rv | ATCACTCAACCGCCGTACCG |
| **Primers for homozygosity assessment of T-DNA insertion lines** | | |
| AT3G26420 | N867804_KO_LP | TTGCATGTCTCGTCATCTTTG |
| | N867804_KO_RP | CTCTCTTGCAAAATGTCCAGG |
| AT2G40550 | N679804_KO_LP | AAGTCGGCATGGTTTATGTTG |
| | N679804_KO_RP | GTGGAAGAACCCTCAGGAAAC |
| AT3G17570 | N668749_KO_LP | CATGGAAACGTGAACGAAAAC |
| | N668749_KO_RP | AAGTGTACCACCGTTTGCAAG |
| AT4G17140 | N678858_KO_LP | TATGTTTGCCTTTGACGAACC |
| | N678858_KO_RP | TACGCAATCCATTCTCATTCC |
| AT3G26430 | N680515_KO_LP | ACATTCGACATCGGTCAGAAC |
| | N680515_KO_RP | TCATCAAATACGTTTCACTTCATTC |
| AT5G57210 | N661789_KO_LP | AATATGTTCAAGCATGGACCG |
| | N661789_KO_RP | CTTCTCTTCACAGCCATCTCG |
| AT1G75060 | N668850_KO_LP | CAAGAACACTAAGCGCATTCC |
| | N668850_KO_RP | CAATCCTTCACAAAATGCACC |
| AT1G02080 | N868048_KO_LP | TGGAGGGAGACTTAGACCATG |
| | N868048_KO_RP | GGGATTTACGAGAACGCTCTC |
| AT5G57200 | N676976_KO_LP | CTGCATCCAAACAATTGGAAG |
| | N676976_KO_RP | ACGAACATGACGTTCTTTTGG |
| AT1G13700 | N658610_KO_LP | TCAATGTGCAAAATGTGAACC |

| | | |
|---|---|---|
| | N658610_KO_RP | AGAGAGCTGTCTGGTAAGGGC |
| AT1G05630 | N672499_KO_LP | GAGAAGCAGTGACGGACTTTG |
| | N672499_KO_RP | GGTTGGAAAAGTCTTCCAAGG |
| T-DNA left border primer | SALK-LB-1.3 | ATTTTGCCGATTTCGGAAC |

**Primers for qPCR validation of transcriptome**

| | | |
|---|---|---|
| AT3G12900 | AT3G12900 LP (RT) | GGCACCAAATCCCTCCCAGA |
| | AT3G12900 RP (RT) | TTTTGCCGTCGTGTGGTTGG |
| AT3G56980 | AT3G56980 LP (RT) | GTGGACATGTCATCTTCAAGGTCT |
| | AT3G56980 RP (RT) | ACATCCTCTGACTTAACTCTTCGC |
| AT1G77120 | AT1G77120 LP (RT) | CCGGAGAATGTGGGGAGTGT |
| | AT1G77120 RP (RT) | CCTCGCTCGGTGTTGATCCT |
| AT3G13610 | AT3G13610 LP (RT) | CCTGAGGTGATTGCAAACGGA |
| | AT3G13610 RP (RT) | TCCATCGTGTGCCTTCCTGA |
| AT5G22890 | AT5G22890 LP (RT) | TGGGTTTGCTCTTGTGGGAC |
| | AT5G22890 RP (RT) | TTTAGGGTGATTGTCGGCGG |

**Primers for qPCR validation of knockdown T-DNA insertion lines**

| | | |
|---|---|---|
| AT3G17570 | AT3G17570_RT-Fw | CGAGGCCAAAGACTTGTCGT |
| | AT3G17570_RT-Rv | GTCTGTATCACAAACCACCACCA |
| AT5G57200 | AT5G57200_RT-Fw | GTTACGGATTTGGAGCCACGG |
| | AT5G57200_RT-Rv | AGGATCTTGTTGCACCCCGA |
| AT2G40550 | AT2G40550_RT-Fw | TGGCTCGTATGATGTCTGTGAGT |
| | AT2G40550_RT-Rv | ACGGTGGTTTTACTTGAGCCTC |
| AT1G13700 | AT1G13700_RT-Fw | TGCCCGTCATCAACTCAGCT |
| | AT1G13700_RT-Rv | TGCAGGCAGAGACAGAGAGC |
| AT3G26420 | AT3G26420_ RT_Fw | CGCTCCCGTGGTTTTGGATT |
| | AT3G26420_RT_Rv | TATCTCTGCCTGCACCACCC |
| AT1G75060 | AT1G75060_RT_Fw | GGATGAGCTTCAGGTTATTGTGGG |
| | AT1G75060_RT_Rv | TGTTTCTGGCTTCTTTGGTCTTCC |
| AT1G02080 | AT1G02080_RT_Fw | GCCTTCATCAGATGTGCGCC |
| | AT1G02080_RT_Rv | GACCCAACCTCCCGAAACCA |

### Table S3 Genes associated with the most significant SNPs in a genome-wide association study of high pH stress tolerance in *Arabidopsis thaliana*

| Chr | Physical position (bp) | GWAS *P*-value | RRL tolerant allele | Frequency of tolerant allele | RRL sensitive allele | Frequency of sensitive allele | *t*-test *P*-value | SNP position with respect to the gene | Nearest Gene | Gene symbol | Expression levels: 1 mM $NaHCO_3$ pH 8 In-house transcriptome log$_2$FC | *P*-value | *FDR* | 10 mM $NaHCO_3$ pH 8.3 GSE164502 log$_2$FC | *FDR* | Low pH 4.5 GSE18982 log$_2$FC | *FDR* | Low Fe GSE16964 log$_2$FC | *FDR* |
|---|---|---|---|---|---|---|---|---|---|---|---|---|---|---|---|---|---|---|---|
| 1 | 28184486 | $1.28 \times 10^{-4}$ | 119/A | 44.2 ± 1.2 | 99/G | 37.2 ± 1.2 | $3.15 \times 10^{-5}$ | -1223 | AT1G75080 | *BZR1* | | | | | | | | | |
| ***Chr1:28174614-Chr1:28186622*** | | | | | | | | -1201 | **AT1G75060** | *AFR1* | | | | | | | | | |
| | | | | | | | | 3424 | AT1G75050 | | | | | | | | | | |
| | | | | | | | | 9231 | AT1G75030 | *TLP-3* | | | | | | | | | |
| 1 | 383526 | $1.39 \times 10^{-4}$ | 75/G | 45.3 ± 1.5 | 143/A | 38.7 ± 1 | $1.68 \times 10^{-4}$ | 3953 | AT1G02090 | *FUS5* | | | | | | | | | |
| ***Chr1:373685-Chr1:389215*** | | | | | | | | Exon | **AT1G02080** | *NOT1* | | | | | | | | | |
| 1 | 4697671 | $1.56 \times 10^{-4}$ | 41/A | 47 ± 2.3 | 177/G | 39.6 ± 0.9 | $5.71 \times 10^{-4}$ | -2071 | **AT1G13700** | *PGL1* | -0.67 | $4.90 \times 10^{-2}$ | | | | | | | |
| ***Chr1:4697671-Chr1:4704182*** | | | | | | | | 5261 | AT1G13710 | *CYP78A5* | -0.57 | $2.70 \times 10^{-2}$ | | -3.3 | $1.62 \times 10^{-3}$ | | | | |
| 1 | 371484 | $2.18 \times 10^{-4}$ | 101/T | 44.2 ± 1.2 | 117/A | 38.3 ± 1.2 | $4.04 \times 10^{-4}$ | -12579 | AT1G02070 | | | | | | | | | | |
| | | | | | | | | -2210 | AT1G02080 | *NOT1* | | | | | | | | | |
| | | | | | | | | -537 | **AT1G02070** | | | | | | | | | | |
| 1 | 375468 | $2.02 \times 10^{-4}$ | 77/A | 45.2 ± 1.5 | 141/G | 38.8 ± 1 | $2.36 \times 10^{-4}$ | 4335 | AT1G02065 | *SPL8* | | | | | | | | | |
| ***Chr1:367218-Chr1:383526*** | | | | | | | | 4335 | | | | | | | | | | | |
| | | | | | | | | Intron | **AT1G02080** | *NOT1* | | | | | | | | | |
| 1 | 375499 | $2.31 \times 10^{-4}$ | 77/A | 45.1 ± 1.5 | 141/T | 38.8 ± 1 | $2.78 \times 10^{-4}$ | Intron | | | | | | | | | | | |
| 1 | 8659841 | $2.65 \times 10^{-4}$ | 169/C | 42.6 ± 1 | 49/A | 35.6 ± 1.4 | $4.40 \times 10^{-4}$ | -344 | **AT1G24430** | | -1.28 | $6.00 \times 10^{-3}$ | | | | | | | |
| 1 | 384674 | $2.89 \times 10^{-4}$ | 92/G | 44.4 ± 1.3 | 126/T | 38.6 ± 1.1 | $5.62 \times 10^{-4}$ | Intron | **AT1G02080** | *NOT1* | | | | | | | | | |
| 1 | 1686725 | $3.00 \times 10^{-4}$ | 27/T | 49.3 ± 2.5 | 191/C | 39.8 ± 0.9 | $1.80 \times 10^{-4}$ | 711 | AT1G05640 | | | | | | | | | | |
| ***Chr1:1686725-Chr1:1690686*** | | | | | | | | 3539 | AT1G05650 | | 2.82 | $3.4 \times 10^{-5}$ | $3.00 \times 10^{-3}$ | | | | | | |
| | | | | | | | | Intron | **AT1G05630** | *5PTASE13* | | | | -1.4 | $2.84 \times 10^{-2}$ | -0.7 | $7.92 \times 10^{-4}$ | | |
| 1 | 371410 | $3.23 \times 10^{-4}$ | 74/C | 45.2 ± 1.6 | 144/G | 38.9 ± 1 | $3.39 \times 10^{-4}$ | -2210 | AT1G02080 | *NOT1* | | | | | | | | | |
| ***Chr1:371410-Chr1:372205*** | | | | | | | | -463 | **AT1G02070** | | | | | | | | | | |
| 1 | 394129 | $3.90 \times 10^{-4}$ | 93/A | 44.8 ± 1.4 | 125/G | 38.2 ± 1.1 | $8.43 \times 10^{-5}$ | Exon | **AT1G02110** | | | | | -1.0 | $1.31 \times 10^{-3}$ | -0.5 | $2.06 \times 10^{-3}$ | | |
| 1 | 396021 | $4.09 \times 10^{-4}$ | 113/T | 44 ± 1.2 | 105/A | 37.8 ± 1.2 | $1.64 \times 10^{-4}$ | 4329 | AT1G02130 | *RAB1B* | | | | | | | | | |
| ***Chr1:396021-Chr1:400593*** | | | | | | | | Intron | **AT1G02120** | *VAD1* | | | | | | 0.2 | $5.56 \times 10^{-2}$ | | |
| 1 | 389949 | $4.56 \times 10^{-4}$ | 103/C | 44.4 ± 1.3 | 115/T | 38 ± 1.1 | $1.19 \times 10^{-4}$ | Intron | **AT1G02100** | *SBI1* | | | | | | | | | |
| 1 | 390770 | $4.79 \times 10^{-4}$ | | | | | | -4991 | AT1G02120 | *VAD1* | | | | | | | | | |
| ***Chr1:389949-Chr1:395878*** | | | | | | | | -2169 | AT1G02110 | | | | | -1.0 | $1.31 \times 10^{-3}$ | | | | |

| | | | | | | | | | | | | | | | | | | |
|---|---|---|---|---|---|---|---|---|---|---|---|---|---|---|---|---|---|---|
| | | | 110/C | 44.1 ± 1.2 | 108/A | 37.9 ± 1.2 | $1.74 \times 10^{-4}$ | Exon | **AT1G02100** | *SBI1* | | | | | | | | |
| 1 | 3708787 | $4.99 \times 10^{-4}$ | 142/C | 43 ± 1.1 | 76/G | 37.3 ± 1.5 | $1.20 \times 10^{-3}$ | Exon | **AT1G11100** | *FRG5* | -1.33 | $1.00 \times 10^{-2}$ | | | 0.2 | $4.95 \times 10^{-2}$ | 0.8 | $3.05 \times 10^{-4}$ |
| 1 | 24091910 | $5.28 \times 10^{-4}$ | 18/G | 49.5 ± 3.4 | 200/C | 40.2 ± 0.9 | $2.28 \times 10^{-3}$ | Exon | **AT1G64830** | | | | | | -0.3 | $9.36 \times 10^{-2}$ | | |
| 1 | 397698 | $5.56 \times 10^{-4}$ | 96/C | 44.6 ± 1.4 | 122/T | 38.2 ± 1.1 | $1.22 \times 10^{-4}$ | 2652 | AT1G02130 | *RA-5* | | | | | | | | |
| ***Chr1:395787-Chr1:401762*** | | | | | | | | Intron | **AT1G02120** | *VAD1* | | | | | | | | |
| 1 | 24091713 | $5.63 \times 10^{-4}$ | 17/T | 49.7 ± 3.6 | 201/G | 40.3 ± 0.9 | $2.43 \times 10^{-3}$ | 1155 | AT1G64820 | *DTX7* | | | | | | | | |
| ***Chr1:24090096-Chr1:24091821*** | | | | | | | | Exon | **AT1G64830** | | | | | | | | | |
| 1 | 28911150 | $5.91 \times 10^{-4}$ | 183/T | 42.2 ± 1 | 35/C | 34.8 ± 1.9 | $1.16 \times 10^{-3}$ | Intron | **AT1G76950** | *PRAF1* | | | | | | | 0.9 | $5.48 \times 10^{-4}$ |
| 1 | 390415 | $5.91 \times 10^{-4}$ | 94/C | 44.6 ± 1.4 | 124/T | 38.3 ± 1.1 | $1.55 \times 10^{-4}$ | Intron | **AT1G02100** | *SBI1* | | | | | | | | |
| 1 | 28858033 | $5.97 \times 10^{-4}$ | 190/A | 42.1 ± 1 | 28/T | 33.4 ± 1.7 | $4.80 \times 10^{-4}$ | -2396 | AT1G76860 | *LSM3B* | | | | | | | | |
| ***Chr1:28850969-Chr1:28858033*** | | | | | | | | 3751 | AT1G76850 | *SEC5A* | | | | | -0.3 | $2.97 \times 10^{-2}$ | | |
| | | | | | | | | Exon | **AT1G76870** | | | | | | | | | |
| 1 | 390744 | $6.43 \times 10^{-4}$ | 111/G | 44 ± 1.2 | 107/T | 37.9 ± 1.2 | $2.26 \times 10^{-4}$ | -5017 | AT1G02120 | *VAD1* | | | | | | | | |
| ***Chr1:389949-Chr1:395878*** | | | | | | | | Intron | **AT1G02100** | *SBI1* | | | | | | | | |
| 1 | 2480846 | $6.45 \times 10^{-4}$ | 28/A | 47.7 ± 2.7 | 190/T | 40 ± 0.9 | $1.99 \times 10^{-3}$ | 2614 | AT1G08000 | *GATA10* | | | | | | | | |
| ***Chr1:2480846-Chr1:2483787*** | | | | | | | | Intron | **AT1G07990** | | | | | | | | | |
| 1 | 392316 | $6.53 \times 10^{-4}$ | 114/T | 43.9 ± 1.2 | 104/G | 37.9 ± 1.2 | $2.89 \times 10^{-4}$ | -3445 | AT1G02120 | *VAD1* | | | | | | | | |
| ***Chr1:389949-Chr1:395878*** | | | | | | | | Intron | **AT1G02100** | *SBI1* | | | | | | | | |
| 1 | 2483710 | $7.67 \times 10^{-4}$ | 25/G | 47.8 ± 3.1 | 193/A | 40.1 ± 0.9 | $3.74 \times 10^{-3}$ | 812 | AT1G07990 | | | | | | | | | |
| ***Chr1:2482905-Chr1:2483710*** | | | | | | | | Exon | **AT1G08000** | *GATA10* | | | | | | | | |
| 1 | 372172 | $7.73 \times 10^{-4}$ | 90/A | 44.4 ± 1.3 | 128/T | 38.7 ± 1.1 | $7.40 \times 10^{-4}$ | -1163 | **AT1G02080** | *NOT1* | | | | | | | | |
| 1 | 401107 | $7.75 \times 10^{-4}$ | 96/A | 44.5 ± 1.4 | 122/G | 38.3 ± 1.1 | $1.84 \times 10^{-4}$ | Intron | **AT1G02130** | *RA-5* | | | | | | | | |
| 1 | 19589589 | $7.84 \times 10^{-4}$ | 70/T | 45.6 ± 1.6 | 148/A | 38.9 ± 1 | $1.67 \times 10^{-4}$ | Exon | **AT1G52590** | | | | | | | | 0.3 | $4.95 \times 10^{-2}$ |
| 1 | 400593 | $9.08 \times 10^{-4}$ | 116/C | 43.8 ± 1.2 | 102/A | 37.8 ± 1.2 | $3.40 \times 10^{-4}$ | 873 | AT1G02120 | *VAD1* | | | | | | | | |
| ***Chr1:396021-Chr1:400593*** | | | | | | | | Intron | **AT1G02130** | *RA-5* | | | | | | | | |
| 2 | 16936596 | $5.40 \times 10^{-5}$ | 149/A | 42.9 ± 1.1 | 69/G | 37 ± 1.3 | $1.06 \times 10^{-3}$ | 10921 | AT2G40520 | *NTP2* | | | | | | | | |
| ***Chr2:16924912-Chr2:16937554*** | | $5.40 \times 10^{-5}$ | | | | | | Exon | **AT2G40550** | *ETG1* | | | | | | | 0.4 | $9.55 \times 10^{-3}$ |
| 2 | 11517635 | $3.69 \times 10^{-4}$ | 56/C | 46 ± 1.7 | 162/A | 39.3 ± 1 | $4.52 \times 10^{-4}$ | Intron | **AT2G26980** | *CIPK3* | | | -1.5 | $7.50 \times 10^{-3}$ | -0.3 | $4.80 \times 10^{-2}$ | -0.6 | $8.40 \times 10^{-3}$ |
| 2 | 8902491 | $5.11 \times 10^{-4}$ | 187/A | 42.3 ± 1 | 31/T | 33 ± 1.6 | $9.12 \times 10^{-5}$ | 1463 | AT2G20650 | *FLY2* | | | -1.3 | $1.14 \times 10^{-2}$ | | | | |
| ***Chr2:8902491-Chr2:8904702*** | | | | | | | | Intron | **AT2G20635** | *BUB1* | | | | | | | | |
| 2 | 19455337 | $9.13 \times 10^{-4}$ | 202/T | 41.7 ± 0.9 | 16/C | 32.7 ± 3.1 | $5.20 \times 10^{-3}$ | 3830 | AT2G47430 | *CKI1* | | | | | | | | |
| ***Chr2:19448567-Chr2:19462870*** | | | | | | | | Exon | **AT2G47410** | | | | | | | | | |

| | | | | | | | | | | | | | | | | | | |
|---|---|---|---|---|---|---|---|---|---|---|---|---|---|---|---|---|---|---|
| 2 | 9174115 | $9.58 \times 10^{-4}$ | 49/A | 44.8 ± 1.7 | 169/T | 39.9 ± 1 | $1.62 \times 10^{-2}$ | Exon | **AT2G21440** | | | | | | -0.3 | $1.64 \times 10^{-2}$ | | |
| 3 | 6009787 | $7.38 \times 10^{-5}$ | 23/A | 50.8 ± 3.2 | 195/T | 39.9 ± 0.9 | $5.27 \times 10^{-5}$ | Exon | **AT3G17570** | | | | | | | | | |
| 3 | 9675996 | $1.51 \times 10^{-5}$ | 69/A | 45.3 ± 1.7 | 149/G | 39 ± 1 | $4.11 \times 10^{-4}$ | 107 | **AT3G26430** | *GGL20* | | | | | | | | |
| 3 | 9675361 | $7.97 \times 10^{-5}$ | 79/T | 44.8 ± 1.5 | 139/C | 38.9 ± 1 | $6.90 \times 10^{-4}$ | Exon | | | | | | | | | | |
| | | | 79/T | 44.8 ± 1.5 | 139/C | 38.9 ± 1 | $6.90 \times 10^{-4}$ | 2306 | AT3G26420 | *RBGB2* | | | | | | | | |
| 3 | 22356075 | $3.15 \times 10^{-4}$ | 14/C | 50.9 ± 4.2 | 204/T | 40.3 ± 0.9 | $1.87 \times 10^{-3}$ | 1066 | AT3G60510 | | | | | | 0.2 | $4.12 \times 10^{-2}$ | | |
| ***Chr3:22350119-Chr3:22359929*** | | | | | | | | 5778 | AT3G60490 | *ERF035* | | | | | | | 0.3 | $1.83 \times 10^{-1}$ |
| | | | | | | | | Intron | **AT3G60500** | *CER7* | | | | | | | | |
| 3 | 9862427 | $4.51 \times 10^{-4}$ | 30/C | 46.6 ± 2.5 | 188/T | 40.1 ± 0.9 | $7.52 \times 10^{-3}$ | Exon | **AT3G26800** | | | | | | | | -0.3 | $6.76 \times 10^{-2}$ |
| 3 | 9709412 | $5.50 \times 10^{-4}$ | 30/T | 46.9 ± 2.7 | 188/G | 40.1 ± 0.9 | $5.14 \times 10^{-3}$ | Exon | | | | | | | | | | |
| 3 | 9709951 | $5.50 \times 10^{-4}$ | 30/C | 46.9 ± 2.7 | 188/T | 40.1 ± 0.9 | $5.14 \times 10^{-3}$ | -7 | **AT3G26500** | *PIRL2* | | | | | | | | |
| 3 | 9709981 | $5.94 \times 10^{-4}$ | 33/C | 46.8 ± 2.7 | 185/T | 40 ± 0.9 | $3.31 \times 10^{-3}$ | -37 | | | | | | | | | | |
| 3 | 8038917 | $5.95 \times 10^{-4}$ | 193/G | 41.9 ± 0.9 | 25/A | 34.3 ± 2.4 | $3.84 \times 10^{-3}$ | Exon | **AT3G22750** | | | | 1.1 | $2.53 \times 10^{-3}$ | -0.8 | $6.4 \times 10^{-4}$ | | |
| 3 | 9667161 | $7.94 \times 10^{-4}$ | 116/C | 43.4 ± 1.3 | 102/T | 38.3 ± 1.2 | $2.32 \times 10^{-3}$ | -7258 | AT3G26430 | *GGL20* | | | | | | | 0.3 | $3.65 \times 10^{-2}$ |
| ***Chr3:9666726-Chr:9671970*** | | | | | | | | -4792 | AT3G26420 | *RBGB2* | | | | | | | | |
| | | | | | | | | 2347 | AT3G26410 | *TRM11* | | | | | | | | |
| | | | | | | | | Exon | **AT3G26400** | *EIF4B1* | | | | | | | | |
| 3 | 17891426 | $9.58 \times 10^{-4}$ | 70/A | 45.4 ± 1.6 | 148/T | 38.9 ± 1 | $2.82 \times 10^{-4}$ | Exon | **AT3G48320** | *CYP71A21* | | | | | | | | |
| 4 | 9200675 | $7.01 \times 10^{-5}$ | 50/G | 46.9 ± 2.1 | 168/A | 39.3 ± 0.9 | $1.11 \times 10^{-4}$ | Intron | **AT4G16260** | | | | 2.2 | $1.63 \times 10^{-3}$ | 0.7 | $7.80 \times 10^{-4}$ | | |
| 4 | 9629622 | $7.55 \times 10^{-5}$ | 30/T | 49.7 ± 2.6 | 188/C | 39.6 ± 0.9 | $2.98 \times 10^{-5}$ | Intron | | | -0.63 | 4.7E-02 | | | -0.3 | $1.24 \times 10^{-2}$ | | |
| 4 | 9623937 | $2.12 \times 10^{-4}$ | 35/C | 47.9 ± 2.3 | 183/T | 39.7 ± 0.9 | $3.08 \times 10^{-4}$ | Exon | | | | | | | | | | |
| 4 | 9632177 | $2.12 \times 10^{-4}$ | 35/A | 47.9 ± 2.3 | 183/G | 39.7 ± 0.9 | $3.08 \times 10^{-4}$ | Intron | **AT4G17140** | | | | | | | | | |
| 4 | 9629381 | $3.96 \times 10^{-4}$ | 31/G | 48.2 ± 2.5 | 187/A | 39.8 ± 0.9 | $4.59 \times 10^{-4}$ | Exon | | | | | | | | | | |
| 4 | 9625325 | $4.45 \times 10^{-4}$ | 31/T | 48.2 ± 2.5 | 187/C | 39.8 ± 0.9 | $4.49 \times 10^{-4}$ | Exon | | | | | | | | | | |
| 4 | 9624203 | $4.70 \times 10^{-4}$ | 31/A | 48.1 ± 2.6 | 187/G | 39.8 ± 0.9 | $5.58 \times 10^{-4}$ | Intron | | | | | | | | | | |
| 4 | 16290516 | $8.71 \times 10^{-4}$ | 152/T | 42.5 ± 1.1 | 66/A | 37.6 ± 1.5 | $7.63 \times 10^{-3}$ | Exon | **AT4G33990** | *EMB2758* | | | | | | | | |
| 4 | 7069534 | $9.28 \times 10^{-4}$ | 141/T | 42.8 ± 1.1 | 77/C | 37.7 ± 1.4 | $3.57 \times 10^{-3}$ | Exon | **AT4G11730** | | | | | | | | | |
| 5 | 23181290 | $7.98 \times 10^{-5}$ | 56/G | 46.3 ± 1.7 | 162/C | 39.2 ± 1 | $1.88 \times 10^{-4}$ | 689 | AT5G57200 | *PICALM2a* | | | | | -0.2 | $7.62 \times 10^{-2}$ | | |
| ***Chr5:23177752-Chr5:23181457*** | | | | | | | | Intron | AT5G57210 | | | | | | | | | |
| 5 | 6112944 | $1.50 \times 10^{-4}$ | 43/G | 47.2 ± 2.1 | 175/C | 39.5 ± 0.9 | $2.08 \times 10^{-4}$ | -148 | AT5G18440 | *NUFIP* | | | | | | | | |
| 5 | 23179225 | $1.55 \times 10^{-4}$ | 54/G | 45.9 ± 1.8 | 164/A | 39.4 ± 1 | $7.95 \times 10^{-4}$ | 1863 | AT5G57210 | | | | | | | | | |
| ***Chr5:23177752-Chr5:23181457*** | | | | | | | | Exon | AT5G57200 | | | | | | | | | |
| 5 | 23179402 | $4.20 \times 10^{-4}$ | 63/C | 45.1 ± 1.6 | 155/G | 39.4 ± 1 | | 1686 | AT5G57210 | | | | | | | | | |

| | | | | | | | | | | | | | | |
|---|---|---|---|---|---|---|---|---|---|---|---|---|---|---|
| *Chr5:23177752-Chr5:23181457* | | | | | | | $1.97 \times 10^{-3}$ | Exon | AT5G57200 | | | | | |
| 5 | 15944567 | $4.48 \times 10^{-4}$ | 176/T | 42.4 ± 1 | 42/C | 35.1 ± 1.7 | $5.67 \times 10^{-4}$ | Exon | AT5G39830 | *DEG8* | | | | |
| 5 | 23180259 | $4.79 \times 10^{-4}$ | 60/A | 45.2 ± 1.7 | 158/G | 39.4 ± 1 | $2.06 \times 10^{-3}$ | Exon | AT5G57200 | *PICALM2a* | | | | |
| 5 | 23181457 | $4.79 \times 10^{-4}$ | 60/A | 45.2 ± 1.7 | 158/G | 39.4 ± 1 | $2.06 \times 10^{-3}$ | 856 | | | | | | |
| *Chr5:23177752-Chr5:23181457* | | | | | | | | Exon | **AT5G57210** | | | | | |
| 5 | 8588698 | $4.80 \times 10^{-4}$ | 19/G | 47.6 ± 4.1 | 199/C | 40.4 ± 0.9 | $1.45 \times 10^{-2}$ | -627 | **AT5G24930** | *COL4* | | | 0.3 | $3.05 \times 10^{-2}$ |
| 5 | 15566986 | $5.13 \times 10^{-4}$ | 99/G | 44.6 ± 1.4 | 119/A | 38.1 ± 1 | $1.00 \times 10^{-4}$ | Intron | **AT5G38880** | *AUG5* | -1.1 | $8.00 \times 10^{-3}$ | | |
| 5 | 7392642 | $5.29 \times 10^{-4}$ | 199/A | 41.8 ± 0.9 | 19/T | 32.3 ± 3.4 | $1.23 \times 10^{-3}$ | Intron | **AT5G22330** | *RIN1* | | | | |
| 5 | 7389168 | $6.01 \times 10^{-4}$ | 199/T | 41.8 ± 0.9 | 19/A | 32.5 ± 3.4 | $1.57 \times 10^{-3}$ | -3823 | AT5G22310 | | -2.1 | $1.80 \times 10^{-3}$ | -0.8 | $7.40 \times 10^{-5}$ |
| *Chr5:7386167-Chr5:7392503* | | | | | | | | 1858 | AT5G22330 | *RIN1* | | | | |
| | | | | | | | | Exon | **AT5G22320** | | | | | |
| 5 | 6590926 | $7.47 \times 10^{-4}$ | 25/T | 48.7 ± 3.3 | 193/C | 40 ± 0.9 | $8.51 \times 10^{-4}$ | Intron | **AT5G19530** | *ACL5* | -1.8 | $1.28 \times 10^{-2}$ | 0.3 | $3.05 \times 10^{-2}$ |
| 5 | 3675931 | $7.93 \times 10^{-4}$ | 204/C | 41.5 ± 0.9 | 14/G | 33.7 ± 3.2 | $2.25 \times 10^{-2}$ | -8890 | AT5G11470 | | | | | |
| *Chr5:3666881-Chr5:3683036* | | | | | | | | -4521 | AT5G11510 | *MYB3R-4* | -1.9 | $6.83 \times 10^{-4}$ | | |
| | | | | | | | | Exon | **AT5G11490** | | | | | |
| 5 | 6113789 | $9.49 \times 10^{-4}$ | 70/C | 44.6 ± 1.7 | 148/T | 39.3 ± 1 | $3.40 \times 10^{-3}$ | Intron | **AT5G18440** | *NUFIP* | | | | |

Genes within haploblocks of the most significantly associated single nucleotide polymorphisms (SNPs) in a genome-wide association study of root length variation of worldwide *Arabidopsis thaliana* ecotypes under high pH. Top-ranking SNPs having a *P*-value < $10^{-3}$ are represented in the table. The haploblocks are indicated by bold rectangles. The locations of the SNPs with respect to the nearest genes are shown. Minus and plus signs denote upstream and downstream locations of the SNP with respect to the gene. The relative root length (RRL)s of the *A. thaliana* ecotypes having tolerant and sensitive alleles are shown as average values with standard error, along with the allele frequencies and a *t*-test *P*-value between these groups. The expression levels of the genes in wild type Arabidopsis (Col-0) seedlings under high pH 8.0 (1mM $NaHCO_3$; in-house transcriptome) and under 10 mM $NaHCO_3$, low pH 4.5, and iron deficiency (public transcriptome datasets) are shown as logarithm of the fold-change in expression levels in treatment versus control (pH 5.7-6), along with Benjamini-Hochberg *false discovery rate* (*FDR*).

**Table S4. Functional enrichment of genes detected by a GWAS of high pH tolerance in *Arabidopsis thaliana***

| Enriched GO biological process | Fold Enrichment | *FDR* | Number of GWAS-detected genes | Number of Pathway Genes | AGI codes |
|---|---|---|---|---|---|
| **Growth and development** | | | | | |
| GO:0048507 Meristem development | 13.0 | 0.041 | 2 | 157 | AT1G76870, AT5G22330 |
| GO:0046622 Positive regulation of organ growth | 169.8 | 0.040 | 1 | 6 | AT1G13710 |
| GO:0010338 Leaf formation | 254.7 | 0.035 | 1 | 4 | AT1G13710 |
| GO:0040009 Regulation of growth rate | 254.7 | 0.035 | 1 | 4 | AT1G13710 |
| GO:0080117 Secondary growth | 254.7 | 0.035 | 1 | 4 | AT2G47430 |
| GO:0009826 Unidimensional cell growth | 11.6 | 0.041 | 2 | 175 | AT5G19530, AT5G22310 |
| GO:0030154 Cell differentiation | 8.2 | 0.029 | 5 | 623 | AT1G08000, AT1G76950, AT2G20650, AT3G22750, AT5G22310 |
| GO:0048759 Xylem vessel member cell differentiation | 92.6 | 0.041 | 1 | 11 | AT5G19530 |
| GO:0010087 Phloem or xylem histogenesis | 24.3 | 0.035 | 2 | 84 | AT2G47430, AT5G19530 |
| GO:0048316 Seed development | 8.4 | 0.035 | 4 | 486 | AT1G05640, AT1G75080, AT1G76870, AT5G22320 |
| **Cell division** | | | | | |
| GO:0007062 Sister chromatid cohesion | 72.8 | 0.041 | 1 | 14 | AT2G40550 |
| GO:0007094 Mitotic spindle assembly checkpoint signaling | 101.9 | 0.041 | 1 | 10 | AT2G20635 |
| GO:0051754 Meiotic sister chromatid cohesion-centromeric | 203.8 | 0.035 | 1 | 5 | AT2G20635 |
| GO:0032465 Regulation of cytokinesis | 203.8 | 0.035 | 1 | 5 | AT5G11510 |
| **DNA repair** | | | | | |
| GO:0006301 Postreplication repair | 92.6 | 0.041 | 1 | 11 | AT2G40550 |
| **Epigenetic regulation** | | | | | |
| GO:0031060 Regulation of histone methylation | 339.6 | 0.035 | 1 | 3 | AT5G11470 |
| **Transcriptional regulation** | | | | | |
| GO:0006355 Regulation of transcription-DNA-templated | 4.1 | 0.035 | 6 | 1485 | AT1G08000, AT1G75060, AT1G75080, AT3G60490, AT5G11510, AT5G24930 |
| GO:0032922 Circadian regulation of gene expression | 84.9 | 0.041 | 1 | 12 | AT5G24930 |
| **mRNA decay** | | | | | |
| GO:0071028 Nuclear mRNA surveillance | 127.4 | 0.041 | 1 | 8 | AT3G60500 |
| GO:0034427 Nuclear-transcribed mRNA catabolic process-exonucleolytic-3'-5' | 78.4 | 0.041 | 1 | 13 | AT3G60500 |
| GO:0071042 Nuclear polyadenylation-dependent mRNA catabolic process | 254.7 | 0.035 | 1 | 4 | AT3G60500 |
| GO:0000288 Nuclear-transcribed mRNA catabolic process-deadenylation-dependent decay | 339.6 | 0.035 | 1 | 3 | AT1G02080 |

| | | | | | |
|---|---|---|---|---|---|
| **Splicing** | | | | | |
| GO:0034473 U1 snRNA 3'-end processing | 254.7 | 0.035 | 1 | 4 | AT3G60500 |
| GO:0034476 U5 snRNA 3'-end processing | 254.7 | 0.035 | 1 | 4 | AT3G60500 |
| GO:0034475 U4 snRNA 3'-end processing | 92.6 | 0.041 | 1 | 11 | AT3G60500 |
| GO:0000492 Box C/D snoRNP assembly | 339.6 | 0.002 | 2 | 6 | AT5G18440, AT5G22330 |
| **Protein processing** | | | | | |
| GO:0035304 Regulation of protein dephosphorylation | 113.2 | 0.041 | 1 | 9 | AT1G05640 |
| GO:0000338 Protein deneddylation | 92.6 | 0.041 | 1 | 11 | AT1G02090 |
| GO:0045053 Protein retention in Golgi apparatus | 203.8 | 0.035 | 1 | 5 | AT4G17140 |
| GO:0006417 Regulation of translation | 78.4 | 0.041 | 1 | 13 | AT1G02080 |
| GO:0006893 Golgi to plasma membrane transport | 72.8 | 0.041 | 1 | 14 | AT1G76850 |
| GO:0043666 Regulation of phosphoprotein phosphatase activity | 127.4 | 0.041 | 1 | 8 | AT1G07990 |
| **Metabolism** | | | | | |
| GO:0005975 Carbohydrate metabolic process | 9.2 | 0.035 | 3 | 332 | AT1G05650, AT1G13700, AT4G16260 |
| GO:0009051 Pentose-phosphate shunt-oxidative branch | 72.8 | 0.041 | 1 | 14 | AT1G13700 |
| GO:0006596 Polyamine biosynthetic process | 127.4 | 0.041 | 1 | 8 | AT5G19530 |
| GO:0006574 Valine catabolic process | 72.8 | 0.041 | 1 | 14 | AT3G60510 |
| **Photosynthesis** | | | | | |
| GO:0010206 Photosystem II repair | 78.4 | 0.041 | 1 | 13 | AT5G39830 |
| **Phytohormone signaling** | | | | | |
| GO:0009723 Response to ethylene | 12.8 | 0.041 | 2 | 159 | AT1G02120, AT5G22310 |
| GO:1900458 Negative regulation of brassinosteroid mediated signaling pathway | 145.6 | 0.041 | 1 | 7 | AT1G02100 |
| GO:0009751 Response to salicylic acid | 9.0 | 0.035 | 3 | 338 | AT1G02120, AT3G48320, AT5G11510 |

The results of a functional enrichment analysis of genes detected by a genome-wide association study of high pH tolerance in 218 worldwide *Arabidopsis thaliana* ecotypes are shown. The analysis was conducted in ShinyGO (https://bioinformatics.sdstate.edu/go/). The fold-enrichment of each Gene Ontology (GO) biological process and the Benjamini-Hochberg false discovery rate (*FDR*) are shown along with the frequency of occurrence of each process in the GWAS detected geneset (MLM *P*-value < $10^{-3}$, see Table 1) and in the *A. thaliana* genome.

### Table S5 Local association study of root length variation in 151 worldwide *Arabidopsis thaliana* ecotypes under high pH stress

| Chromosome | Physical position | Frequency of Tolerant allele | Average RRL tolerant allele | Frequency of Sensitive allele | Average RRL sensitive allele | t-test *P*-value | LAS GLM *F*-statistic | LAS GLM *P*-value | LAS MLM *F*-statistic | LAS MLM *P*-value | GWAS-detected SNP | Polymorphism effects | Impact | Gene ID | Gene symbol |
|---|---|---|---|---|---|---|---|---|---|---|---|---|---|---|---|
| 1 | 364798 | 13/G | 42.3 ± 2.1 | 138/GT | 41 ± 1.1 | $7.20 \times 10^{-1}$ | 0.7 | $4.00 \times 10^{-1}$ | 0.8 | $3.59 \times 10^{-1}$ | | Promoter polymorphism | Moderate | AT1G02065 | *SPL8* |
| 1 | 365056 | 132/C | 41.5 ± 1.1 | 19/A | 38.4 ± 1.9 | $3.06 \times 10^{-1}$ | 0.3 | $6.02 \times 10^{-1}$ | 0.2 | $6.88 \times 10^{-1}$ | | Promoter polymorphism | Moderate | | |
| 1 | 365411 | 49/G | 41.1 ± 1.6 | 102/A | 41.1 ± 1.3 | $9.82 \times 10^{-1}$ | 0.4 | $5.54 \times 10^{-1}$ | 1.8 | $1.80 \times 10^{-1}$ | | 5'-UTR polymorphism | Moderate | | |
| 1 | 365682 | 123/A | 41.6 ± 1.2 | 28/C | 39 ± 2.3 | $3.10 \times 10^{-1}$ | 0.7 | $3.92 \times 10^{-1}$ | 0.0 | $9.31 \times 10^{-1}$ | | Amino acid substitution (Asn20His) | Moderate | | |
| 1 | 370752 | 145/CTA | 41.2 ± 1.1 | 4/CTATA | 38.6 ± 2.4 | $6.85 \times 10^{-1}$ | 0.1 | $8.82 \times 10^{-1}$ | 0.1 | $9.39 \times 10^{-1}$ | | **Splice acceptor variant** | **High** | AT1G02070 | |
| 1 | 371066 | 14/C | 42.4 ± 3.1 | 137/A | 41 ± 1.1 | $6.85 \times 10^{-1}$ | 0.1 | $7.78 \times 10^{-1}$ | 0.3 | $5.87 \times 10^{-1}$ | | Promoter polymorphism | Moderate | | |
| 1 | 371098 | 61/G | 44.1 ± 1.6 | 90/A | 39 ± 1.3 | $1.14 \times 10^{-2}$ | **8.0** | **$5.35 \times 10^{-3}$** | 1.8 | $1.79 \times 10^{-1}$ | | Promoter polymorphism | Moderate | | |
| 1 | 371410 | 48/C | 44.7 ± 1.9 | 103/G | 39.4 ± 1.2 | **$1.26 \times 10^{-2}$** | **8.7** | **$3.70 \times 10^{-3}$** | **3.0** | $8.45 \times 10^{-2}$ | ✓ | Promoter polymorphism | Moderate | | |
| 1 | 371484 | 65/T | 43.7 ± 1.5 | 86/A | 39.1 ± 1.3 | **$2.00 \times 10^{-2}$** | **7.6** | **$6.49 \times 10^{-3}$** | 0.5 | $4.64 \times 10^{-1}$ | ✓ | Promoter polymorphism | Moderate | | |
| 1 | 372451 | 135/T | 42.1 ± 1.1 | 16/C | 32.5 ± 3.2 | **$2.50 \times 10^{-3}$** | **10.8** | **$1.28 \times 10^{-3}$** | **5.5** | **$2.01 \times 10^{-2}$** | | Promoter polymorphism | Moderate | AT1G02080 | |
| 1 | 372809 | 72/C | 41.8 ± 1.5 | 79/G | 40.4 ± 1.4 | $4.90 \times 10^{-1}$ | 0.9 | $3.38 \times 10^{-1}$ | 0.1 | $7.14 \times 10^{-1}$ | | Promoter polymorphism | Moderate | | |
| 1 | 372927 | 63/C | 43.2 ± 1.6 | 88/T | 39.6 ± 1.3 | $7.40 \times 10^{-2}$ | **4.6** | **$3.38 \times 10^{-2}$** | 0.5 | $4.67 \times 10^{-1}$ | | Promoter polymorphism | Moderate | | |
| 1 | 372938 | 72/T | 43.3 ± 1.4 | 79/A | 39 ± 1.4 | **$3.07 \times 10^{-2}$** | **6.7** | **$1.08 \times 10^{-2}$** | 0.4 | $5.48 \times 10^{-1}$ | | Promoter polymorphism | Moderate | | |
| 1 | 373035 | 62/A | 43.9 ± 1.7 | 89/T | 39.2 ± 1.3 | **$1.89 \times 10^{-2}$** | **6.5** | **$1.15 \times 10^{-2}$** | 1.6 | $2.05 \times 10^{-1}$ | | Promoter polymorphism | Moderate | | |
| 1 | 373097 | 57/C | 44.3 ± 1.7 | 94/T | 39.2 ± 1.3 | $1.18 \times 10^{-2}$ | **7.4** | **$7.21 \times 10^{-3}$** | 2.2 | $1.40 \times 10^{-1}$ | | Promoter polymorphism | Moderate | | |
| 1 | 373144 | 116/G | 41.7 ± 1.2 | 35/T | 39 ± 1.9 | $2.53 \times 10^{-1}$ | 2.7 | $1.04 \times 10^{-1}$ | 0.1 | $7.30 \times 10^{-1}$ | | Promoter polymorphism | Moderate | | |
| 1 | 373150 | 71/G | 43.2 ± 1.5 | 80/T | 39.2 ± 1.4 | **$4.14 \times 10^{-2}$** | **6.0** | **$1.54 \times 10^{-2}$** | 0.0 | $9.48 \times 10^{-1}$ | | Promoter polymorphism | Moderate | | |
| 1 | 373332 | 15/A | 41.5 ± 3.3 | 136/ATAATG | 41 ± 1.1 | $8.82 \times 10^{-1}$ | 0.2 | $6.66 \times 10^{-1}$ | 0.1 | $8.13 \times 10^{-1}$ | | Promoter polymorphism | Moderate | | |
| 1 | 373685 | 64/C | 43.5 ± 1.6 | 87/A | 39.3 ± 1.3 | **$3.77 \times 10^{-2}$** | **6.0** | **$1.55 \times 10^{-2}$** | 1.0 | $3.23 \times 10^{-1}$ | | Amino acid substitution (Pro51Thr) | Moderate | | |
| 1 | 373691 | 7/T | 44.6 ± 3.3 | 144/C | 40.9 ± 1.1 | $4.41 \times 10^{-1}$ | 0.6 | $4.44 \times 10^{-1}$ | 0.1 | $7.13 \times 10^{-1}$ | | Amino acid substitution (Leu53Phe) | Moderate | | |
| 1 | 374611 | 148/G | 41.1 ± 1.1 | 3/A | 38.4 ± 6.8 | $7.01 \times 10^{-1}$ | 0.0 | $8.51 \times 10^{-1}$ | 0.3 | $5.75 \times 10^{-1}$ | | Amino acid substitution (Asp270Asn) | Moderate | | |
| 1 | 374753 | 148/C | 41.2 ± 1.1 | 3/T | 34.4 ± 2.9 | $3.36 \times 10^{-1}$ | 0.8 | $3.85 \times 10^{-1}$ | 0.1 | $7.10 \times 10^{-1}$ | | Amino acid substitution (Ala317Val) | Moderate | | |
| 1 | 375209 | 6/G | 43.4 ± 3.6 | 145/T | 41 ± 1.1 | $6.34 \times 10^{-1}$ | 0.1 | $7.07 \times 10^{-1}$ | 0.0 | $9.13 \times 10^{-1}$ | | Amino acid substitution (Phe403Val) | Moderate | | |
| 1 | 375372 | 4/C | 42.3 ± 9.2 | 147/G | 41.1 ± 1 | $8.41 \times 10^{-1}$ | 0.0 | $9.88 \times 10^{-1}$ | 0.2 | $6.72 \times 10^{-1}$ | | Amino acid substitution (Met428Ile) | Moderate | | |
| 1 | 375468 | 50/A | 44.6 ± 1.9 | 101/G | 39.3 ± 1.2 | $1.20 \times 10^{-2}$ | **8.7** | **$3.78 \times 10^{-3}$** | 2.4 | $1.25 \times 10^{-1}$ | ✓ | Intron polymorphism | Moderate | | |
| 1 | 375499 | 51/A | 44.5 ± 1.8 | 100/T | 39.3 ± 1.2 | **$1.24 \times 10^{-2}$** | **8.5** | **$4.13 \times 10^{-3}$** | 2.2 | $1.43 \times 10^{-1}$ | ✓ | Intron polymorphism | Moderate | | |
| 1 | 376151 | 6/G | 43.4 ± 3.6 | 145/C | 41 ± 1.1 | $6.34 \times 10^{-1}$ | 0.1 | $7.07 \times 10^{-1}$ | 0.0 | $9.13 \times 10^{-1}$ | | Amino acid substitution (Ser597Cys) | Moderate | | |
| 1 | 376801 | 6/C | 43.4 ± 3.6 | 145/A | 41 ± 1.1 | $6.34 \times 10^{-1}$ | 0.1 | $7.07 \times 10^{-1}$ | 0.0 | $9.13 \times 10^{-1}$ | | Amino acid substitution (Glu751Asp) | Moderate | | |
| 1 | 379604 | 143/T | 41.3 ± 1.1 | 8/C | 37.7 ± 5.4 | $4.22 \times 10^{-1}$ | 0.1 | $7.79 \times 10^{-1}$ | 0.1 | $7.07 \times 10^{-1}$ | | Amino acid substitution (Ile1215Thr) | Moderate | | |
| 1 | 379753 | 146/T | 41.1 ± 1.1 | 5/A | 40.4 ± 6.2 | $8.98 \times 10^{-1}$ | 0.3 | $6.09 \times 10^{-1}$ | 0.4 | $5.22 \times 10^{-1}$ | | Amino acid substitution (Ser1265Thr) | Moderate | | |
| 1 | 380999 | 150/GT | 41.2 ± 1 | 1/G | 31.1 ± 0 | | 1.0 | $3.18 \times 10^{-1}$ | 0.7 | $4.20 \times 10^{-1}$ | | **Splice donor variant** | **High** | | |
| 1 | 381773 | 52/A | 44 ± 1.8 | 99/T | 39.6 ± 1.2 | **$3.47 \times 10^{-2}$** | **5.9** | **$1.62 \times 10^{-2}$** | 0.8 | $3.63 \times 10^{-1}$ | | Amino acid substitution (Thr1616Ser) | Moderate | | |
| 1 | 383526 | 49/G | 44.7 ± 1.9 | 102/A | 39.4 ± 1.2 | $1.10 \times 10^{-2}$ | **9.0** | **$3.24 \times 10^{-3}$** | 2.6 | $1.12 \times 10^{-1}$ | ✓ | Synonymous polymorphism | Moderate | | |
| 1 | 383680 | 145/C | 41.3 ± 1 | 6/G | 35.8 ± 6.8 | $2.80 \times 10^{-1}$ | 0.5 | $4.88 \times 10^{-1}$ | 0.8 | $3.86 \times 10^{-1}$ | | Amino acid substitution (Gln1904Glu) | Moderate | | |
| 1 | 384674 | 60/G | 44 ± 1.6 | 91/T | 39.1 ± 1.3 | **$1.52 \times 10^{-2}$** | **8.2** | **$4.79 \times 10^{-3}$** | 1.0 | $3.10 \times 10^{-1}$ | ✓ | Intron polymorphism | Moderate | | |
| 1 | 384704 | 37/A | 42.9 ± 2.2 | 114/T | 40.5 ± 1.2 | $3.11 \times 10^{-1}$ | 1.5 | $2.27 \times 10^{-1}$ | 1.1 | $2.90 \times 10^{-1}$ | | Amino acid substitution (Asp2019Val) | Moderate | | |

| | | | | | | | | | | | | | | | |
|---|---|---|---|---|---|---|---|---|---|---|---|---|---|---|---|
| 1 | 384788 | 148/G | 41.1 ± 1.1 | 3/T | 38.4 ± 6.8 | $7.01 \times 10^{-1}$ | 0.0 | $8.51 \times 10^{-1}$ | 0.3 | $5.75 \times 10^{-1}$ | | Amino acid substitution (Trp2047Leu) | Moderate | | |
| 1 | 389577 | 59/C | 44.3 ± 1.6 | 92/G | 39 ± 1.3 | **$8.57 \times 10^{-3}$** | **8.3** | **$4.56 \times 10^{-3}$** | 2.0 | $1.58 \times 10^{-1}$ | | 5'-UTR polymorphism | Moderate | AT1G02090 | *FUS5* |
| 1 | 389688 | 77/T | 42.3 ± 1.5 | 74/A | 39.9 ± 1.4 | $2.24 \times 10^{-1}$ | 2.2 | $1.36 \times 10^{-1}$ | 0.5 | $4.82 \times 10^{-1}$ | | 5'-UTR polymorphism | Moderate | | |
| 1 | 389692 | 60/T | 42.5 ± 1.7 | 91/G | 40.1 ± 1.3 | $2.45 \times 10^{-1}$ | 1.8 | $1.76 \times 10^{-1}$ | 0.3 | $5.88 \times 10^{-1}$ | | 5'-UTR polymorphism | Moderate | | |
| 1 | 389750 | 64/G | 43.7 ± 1.7 | 84/A | 39.2 ± 1.3 | **$2.73 \times 10^{-2}$** | 2.8 | $6.18 \times 10^{-2}$ | 0.4 | $6.98 \times 10^{-1}$ | | 5'-UTR polymorphism | Moderate | | |
| 1 | 389853 | 81/T | 42.9 ± 1.4 | 70/C | 38.9 ± 1.4 | **$4.37 \times 10^{-2}$** | 2.4 | $1.24 \times 10^{-1}$ | 0.1 | $7.69 \times 10^{-1}$ | | 5'-UTR polymorphism | Moderate | | |
| 1 | 389949 | 97/C | 42.2 ± 1.3 | 54/T | 39.1 ± 1.7 | $1.31 \times 10^{-1}$ | 1.1 | $2.90 \times 10^{-1}$ | 0.0 | $9.16 \times 10^{-1}$ | | Promoter polymorphism | Moderate | | |
| 1 | 389967 | 84/C | 43.1 ± 1.4 | 67/T | 38.5 ± 1.5 | **$1.96 \times 10^{-2}$** | **3.5** | $6.28 \times 10^{-2}$ | 0.7 | $4.03 \times 10^{-1}$ | | Promoter polymorphism | Moderate | | |
| 1 | 389992 | 145/T | 41.2 ± 1.1 | 6/TTG | 38.4 ± 5.5 | $5.86 \times 10^{-1}$ | 0.3 | $5.82 \times 10^{-1}$ | 0.0 | $8.66 \times 10^{-1}$ | | Promoter polymorphism | Moderate | | |
| 1 | 390211 | 93/T | 43.1 ± 1.3 | 58/A | 37.8 ± 1.5 | **$8.37 \times 10^{-3}$** | **5.7** | **$1.78 \times 10^{-2}$** | 1.0 | $3.19 \times 10^{-1}$ | | Promoter polymorphism | Moderate | | |
| 1 | 390320 | 141/G | 41.1 ± 1.1 | 10/A | 40.5 ± 3.5 | $8.85 \times 10^{-1}$ | 0.0 | $9.91 \times 10^{-1}$ | 0.0 | $8.94 \times 10^{-1}$ | | Promoter polymorphism | Moderate | | |
| 1 | 390415 | 70/C | 43.7 ± 1.6 | 81/T | 38.8 ± 1.2 | **$1.44 \times 10^{-2}$** | **4.1** | **$4.45 \times 10^{-2}$** | 0.6 | $4.49 \times 10^{-1}$ | | Promoter polymorphism | Moderate | | |
| 1 | 390542 | 89/A | 43.5 ± 1.4 | 62/C | 37.6 ± 1.5 | $3.02 \times 10^{-3}$ | **7.0** | **$8.97 \times 10^{-3}$** | 1.0 | $3.18 \times 10^{-1}$ | | Promoter polymorphism | Moderate | | |
| 1 | 1681551 | 18/A | 42.8 ± 3.7 | 133/T | 40.9 ± 1.1 | $5.17 \times 10^{-1}$ | 0.4 | $5.35 \times 10^{-1}$ | 0.7 | $4.10 \times 10^{-1}$ | | Promoter polymorphism | Moderate | AT1G05630 | *5PTASE13* |
| 1 | 1681673 | 8/G | 41.9 ± 6 | 143/A | 41 ± 1 | $8.45 \times 10^{-1}$ | 0.0 | $8.67 \times 10^{-1}$ | 0.1 | $7.46 \times 10^{-1}$ | | Promoter polymorphism | Moderate | | |
| 1 | 1681886 | 19/A | 42.8 ± 3.5 | 132/G | 40.8 ± 1.1 | $5.21 \times 10^{-1}$ | 0.4 | $5.38 \times 10^{-1}$ | 0.6 | $4.26 \times 10^{-1}$ | | Promoter polymorphism | Moderate | | |
| 1 | 1681893 | 130/G | 41.5 ± 1.2 | 21/C | 38.5 ± 2 | $2.96 \times 10^{-1}$ | 0.6 | $4.33 \times 10^{-1}$ | 1.5 | $2.24 \times 10^{-1}$ | | Promoter polymorphism | Moderate | | |
| 1 | 1681895 | 11/T | 41.8 ± 3.4 | 140/A | 41 ± 1.1 | $8.47 \times 10^{-1}$ | 0.1 | $7.95 \times 10^{-1}$ | 0.0 | $9.23 \times 10^{-1}$ | | Promoter polymorphism | Moderate | | |
| 1 | 1682406 | 139/C | 41.5 ± 1.1 | 12/T | 36.1 ± 3.4 | $1.39 \times 10^{-1}$ | 1.0 | $3.16 \times 10^{-1}$ | 1.7 | $1.91 \times 10^{-1}$ | | 5'-UTR polymorphism | Moderate | | |
| 1 | 1682478 | 75/T | 41.6 ± 1.3 | 76/C | 40.6 ± 1.6 | $5.95 \times 10^{-1}$ | 0.5 | $4.77 \times 10^{-1}$ | 0.3 | $6.03 \times 10^{-1}$ | | 5'-UTR polymorphism | Moderate | | |
| 1 | 1682509 | 19/T | 42.8 ± 3.5 | 132/A | 40.8 ± 1.1 | $5.21 \times 10^{-1}$ | 0.4 | $5.38 \times 10^{-1}$ | 0.6 | $4.26 \times 10^{-1}$ | | Amino acid substitution (Glu9Asp) | Moderate | | |
| 1 | 1682537 | 144/C | 41.4 ± 1 | 7/T | 35.5 ± 5.9 | $2.16 \times 10^{-1}$ | **3.1** | $7.85 \times 10^{-2}$ | 2.4 | $1.23 \times 10^{-1}$ | | Amino acid substitution (Pro19Ser) | Moderate | | |
| 1 | 1683078 | 4/T | 52.2 ± 4.5 | 147/C | 40.8 ± 1.1 | $6.29 \times 10^{-2}$ | **3.1** | $7.88 \times 10^{-2}$ | 0.1 | $8.18 \times 10^{-1}$ | | Amino acid substitution (Ala199Val) | Moderate | | |
| 1 | 1683220 | 3/G | 48.4 ± 2.9 | 148/T | 40.9 ± 1.1 | $2.93 \times 10^{-1}$ | 1.3 | $2.61 \times 10^{-1}$ | 1.2 | $2.82 \times 10^{-1}$ | | Amino acid substitution (Asp246Glu) | Moderate | | |
| 1 | 1684104 | 148/G | 41.2 ± 1.1 | 3/A | 37.4 ± 3.8 | $6.01 \times 10^{-1}$ | 0.0 | $9.09 \times 10^{-1}$ | 0.0 | $9.50 \times 10^{-1}$ | | Amino acid substitution (Val481Ile) | Moderate | | |
| 1 | 1686463 | 148/A | 41.2 ± 1.1 | 3/T | 33.2 ± 5.7 | $2.58 \times 10^{-1}$ | 0.6 | $4.43 \times 10^{-1}$ | 0.2 | $6.94 \times 10^{-1}$ | | Amino acid substitution (Ile969Leu) | Moderate | | |
| 1 | 1686485 | 143/T | 41.5 ± 1.1 | 8/C | 33.6 ± 3.7 | $7.19 \times 10^{-2}$ | 2.9 | $8.97 \times 10^{-2}$ | **5.7** | **$1.83 \times 10^{-2}$** | | Amino acid substitution (Leu976Ser) | Moderate | | |
| 1 | 1686725 | 18/T | 49 ± 2.4 | 133/C | 40 ± 1.1 | **$3.06 \times 10^{-3}$** | **7.7** | **$6.39 \times 10^{-3}$** | **4.9** | **$2.78 \times 10^{-2}$** | ✓ | Intron polymorphism | Moderate | | |
| 1 | 1686819 | 146/C | 41.3 ± 1.1 | 5/A | 35.5 ± 3.6 | $2.96 \times 10^{-1}$ | 1.0 | $3.11 \times 10^{-1}$ | 2.6 | $1.06 \times 10^{-1}$ | | Amino acid substitution (Pro1060Gln) | Moderate | | |
| 1 | 1686860 | 5/T | 42.4 ± 3.7 | 146/A | 41 ± 1.1 | $8.11 \times 10^{-1}$ | 0.4 | $5.22 \times 10^{-1}$ | 0.7 | $4.06 \times 10^{-1}$ | | Amino acid substitution (Tyr1074Asn) | Moderate | | |
| 1 | 1687058 | 148/T | 41.2 ± 1.1 | 3/A | 38 ± 8.5 | $6.61 \times 10^{-1}$ | 0.0 | $8.76 \times 10^{-1}$ | 0.1 | $7.24 \times 10^{-1}$ | | Amino acid substitution (Leu1140Ile) | Moderate | | |
| 1 | 1688394 | 10/T | 42.5 ± 3.8 | 141/TA | 41 ± 1.1 | $7.05 \times 10^{-1}$ | 0.4 | $5.31 \times 10^{-1}$ | 0.1 | $7.10 \times 10^{-1}$ | | Frameshift (Arg336fs) | **High** | AT1G05640 | |
| 1 | 1689096 | 140/A | 41.2 ± 1.1 | 11/C | 39.5 ± 2.3 | $6.55 \times 10^{-1}$ | 0.1 | $7.82 \times 10^{-1}$ | 0.0 | $8.83 \times 10^{-1}$ | | Amino acid substitution (Cys136Gly) | Moderate | | |
| 1 | 1689275 | 13/A | 47.4 ± 2.6 | 138/C | 40.5 ± 1.1 | $5.02 \times 10^{-2}$ | 1.9 | $1.74 \times 10^{-1}$ | 0.6 | $4.56 \times 10^{-1}$ | | Amino acid substitution (Arg76Leu) | Moderate | | |
| 1 | 1689335 | 11/G | 44.2 ± 3.6 | 140/C | 40.8 ± 1.1 | $3.85 \times 10^{-1}$ | 1.1 | $2.89 \times 10^{-1}$ | 0.0 | $8.38 \times 10^{-1}$ | | Amino acid substitution (Gly56Ala) | Moderate | | |
| 1 | 1689598 | 135/C | 41.1 ± 1.1 | 16/A | 40.9 ± 3.6 | $9.48 \times 10^{-1}$ | 0.1 | $7.65 \times 10^{-1}$ | 0.2 | $6.18 \times 10^{-1}$ | | 5'-UTR polymorphism | Moderate | | |
| 1 | 1689818 | 137/A | 41.3 ± 1.1 | 14/T | 38.8 ± 3.6 | $4.58 \times 10^{-1}$ | 2.7 | $1.04 \times 10^{-1}$ | 0.9 | $3.38 \times 10^{-1}$ | | 5'-UTR polymorphism | Moderate | | |
| 1 | 1689833 | 131/A | 42 ± 1.1 | 20/G | 35.3 ± 2.1 | **$2.30 \times 10^{-2}$** | **3.7** | $5.64 \times 10^{-2}$ | **4.7** | **$3.23 \times 10^{-2}$** | | 5'-UTR polymorphism | Moderate | | |
| 1 | 1689854 | 105/T | 41.8 ± 1.2 | 46/C | 39.5 ± 2 | $2.80 \times 10^{-1}$ | **3.2** | $7.58 \times 10^{-2}$ | 1.1 | $3.06 \times 10^{-1}$ | | 5'-UTR polymorphism | Moderate | | |
| 1 | 1690016 | 134/C | 41.4 ± 1.1 | 17/CT | 38.3 ± 3 | $3.13 \times 10^{-1}$ | 1.8 | $1.85 \times 10^{-1}$ | 0.9 | $3.45 \times 10^{-1}$ | | Promoter polymorphism | Moderate | | |
| 1 | 1690146 | 13/T | 45.9 ± 2.2 | 138/A | 40.6 ± 1.1 | $1.33 \times 10^{-1}$ | 0.9 | $3.38 \times 10^{-1}$ | 1.5 | $2.16 \times 10^{-1}$ | | Promoter polymorphism | Moderate | | |

| | | | | | | | | | | | | | | | |
|---|---|---|---|---|---|---|---|---|---|---|---|---|---|---|---|
| 1 | 1690150 | 13/T | 45.5 ± 2.1 | 138/A | 40.7 ± 1.1 | $1.75 \times 10^{-1}$ | 0.9 | $3.35 \times 10^{-1}$ | 1.4 | $2.32 \times 10^{-1}$ | | Promoter polymorphism | Moderate | | |
| 1 | 1690155 | 140/A | 41.4 ± 1.1 | 11/T | 36.8 ± 4.8 | $2.22 \times 10^{-1}$ | 2.7 | $1.01 \times 10^{-1}$ | 1.3 | $2.60 \times 10^{-1}$ | | Promoter polymorphism | Moderate | | |
| 1 | 1690357 | 11/C | 41.8 ± 3.4 | 140/A | 41 ± 1.1 | $8.47 \times 10^{-1}$ | 0.1 | $7.95 \times 10^{-1}$ | 0.0 | $9.32 \times 10^{-1}$ | | Promoter polymorphism | Moderate | | |
| 1 | 1690472 | 101/G | 41.8 ± 1.2 | 50/C | 39.7 ± 2 | $3.22 \times 10^{-1}$ | 2.6 | $1.10 \times 10^{-1}$ | 0.6 | $4.59 \times 10^{-1}$ | | Promoter polymorphism | Moderate | | |
| 1 | 1690496 | 108/T | 41.9 ± 1.1 | 43/C | 39 ± 2.2 | $1.82 \times 10^{-1}$ | **3.7** | $5.56 \times 10^{-2}$ | 1.1 | $3.05 \times 10^{-1}$ | | Promoter polymorphism | Moderate | | |
| 1 | 1690501 | 126/AC | 41.2 ± 1.1 | 25/A | 40.4 ± 3 | $7.72 \times 10^{-1}$ | 0.5 | $4.83 \times 10^{-1}$ | 0.0 | $9.38 \times 10^{-1}$ | | Promoter polymorphism | Moderate | | |
| 1 | 1690505 | 20/TA | 42.7 ± 3.1 | 131/T | 40.8 ± 1.1 | $5.27 \times 10^{-1}$ | 0.1 | $7.28 \times 10^{-1}$ | 1.7 | $1.99 \times 10^{-1}$ | | Promoter polymorphism | Moderate | | |
| 1 | 1690524 | 114/G | 41.6 ± 1.1 | 37/A | 39.4 ± 2.4 | $3.31 \times 10^{-1}$ | 2.2 | $1.37 \times 10^{-1}$ | 1.2 | $2.73 \times 10^{-1}$ | | Promoter polymorphism | Moderate | | |
| 1 | 1690571 | 105/G | 41.6 ± 1.2 | 46/T | 40 ± 2 | $4.60 \times 10^{-1}$ | 1.8 | $1.80 \times 10^{-1}$ | 0.2 | $6.58 \times 10^{-1}$ | | Promoter polymorphism | Moderate | | |
| 1 | 1690686 | 100/A | 42 ± 1.2 | 51/G | 39.3 ± 1.9 | $1.86 \times 10^{-1}$ | **3.9** | **$4.88 \times 10^{-2}$** | 1.4 | $2.42 \times 10^{-1}$ | | Promoter polymorphism | Moderate | | |
| 1 | 1690855 | 102/G | 41.8 ± 1.2 | 49/A | 39.5 ± 1.9 | $2.79 \times 10^{-1}$ | **3.3** | $7.13 \times 10^{-2}$ | 1.5 | $2.23 \times 10^{-1}$ | | Promoter polymorphism | Moderate | | |
| 1 | 1690910 | 94/T | 42.2 ± 1.2 | 57/C | 39.2 ± 1.8 | $1.37 \times 10^{-1}$ | 2.7 | $1.00 \times 10^{-1}$ | 1.2 | $2.83 \times 10^{-1}$ | | Promoter polymorphism | Moderate | | |
| 1 | 1690917 | 134/T | 41.7 ± 1.1 | 17/TTCCGA | 36.3 ± 2.8 | $8.69 \times 10^{-2}$ | **6.1** | **$1.45 \times 10^{-2}$** | **6.2** | **$1.41 \times 10^{-2}$** | | Promoter polymorphism | Moderate | | |
| 1 | 1690922 | 108/A | 41.6 ± 1.1 | 43/C | 39.9 ± 2.2 | $4.39 \times 10^{-1}$ | 1.9 | $1.71 \times 10^{-1}$ | 1.0 | $3.13 \times 10^{-1}$ | | Promoter polymorphism | Moderate | | |
| 1 | 1690265 | 1/A | 51.5 ± 0 | 150/C | 41 ± 1 | | 1.7 | $1.93 \times 10^{-1}$ | 0.6 | $4.42 \times 10^{-1}$ | | Stop lost (Ter395Leu ext*) | **High** | AT1G05650 | |
| 1 | 1690472 | 101/G | 41.8 ± 1.2 | 50/C | 39.7 ± 2 | $3.22 \times 10^{-1}$ | 2.6 | $1.10 \times 10^{-1}$ | 1.0 | $3.12 \times 10^{-1}$ | | Amino acid substitution (Thr326Ser) | Moderate | | |
| 1 | 1691013 | 49/T | 41.7 ± 1.9 | 102/A | 40.8 ± 1.2 | $6.71 \times 10^{-1}$ | 0.1 | $8.07 \times 10^{-1}$ | 0.7 | $4.16 \times 10^{-1}$ | | Amino acid substitution (Ser232Thr) | Moderate | | |
| 1 | 1691190 | 19/T | 42.8 ± 3.5 | 132/G | 40.8 ± 1.1 | $5.21 \times 10^{-1}$ | 0.4 | $5.38 \times 10^{-1}$ | 0.6 | $4.26 \times 10^{-1}$ | | Amino acid substitution (Leu173Ile) | Moderate | | |
| 1 | 1692231 | 102/AT | 41.3 ± 1.3 | 49/A | 40.6 ± 1.8 | $7.19 \times 10^{-1}$ | 0.0 | $9.09 \times 10^{-1}$ | 0.1 | $7.75 \times 10^{-1}$ | | Promoter polymorphism | Moderate | | |
| 1 | 1692310 | 19/C | 42.8 ± 3.5 | 132/G | 40.8 ± 1.1 | $5.21 \times 10^{-1}$ | 0.4 | $5.38 \times 10^{-1}$ | 0.6 | $4.26 \times 10^{-1}$ | | Promoter polymorphism | Moderate | | |
| 1 | 1692334 | 119/C | 42.1 ± 1.1 | 32/T | 37.5 ± 2.2 | $6.08 \times 10^{-2}$ | **8.7** | **$3.73 \times 10^{-3}$** | **4.0** | **$4.78 \times 10^{-2}$** | | Promoter polymorphism | Moderate | | |
| 1 | 1692427 | 14/G | 43.7 ± 2.9 | 137/T | 40.8 ± 1.1 | $3.93 \times 10^{-1}$ | 1.0 | $3.14 \times 10^{-1}$ | 0.2 | $6.25 \times 10^{-1}$ | | Promoter polymorphism | Moderate | | |
| 1 | 1692602 | 2/A | 48.9 ± 4.9 | 140/C | 41.1 ± 1.1 | $3.81 \times 10^{-1}$ | 0.5 | $6.16 \times 10^{-1}$ | 0.3 | $7.27 \times 10^{-1}$ | | Promoter polymorphism | Moderate | | |
| 1 | 1692698 | 11/T | 47.3 ± 4.4 | 140/G | 40.6 ± 1.1 | $7.81 \times 10^{-2}$ | 2.9 | $9.19 \times 10^{-2}$ | **5.0** | **$2.63 \times 10^{-2}$** | | Promoter polymorphism | Moderate | | |
| 1 | 1692818 | 20/CT | 41.8 ± 2.8 | 131/C | 41 ± 1.1 | $7.69 \times 10^{-1}$ | 0.3 | $5.69 \times 10^{-1}$ | 1.2 | $2.72 \times 10^{-1}$ | | Promoter polymorphism | Moderate | | |
| 1 | 1692839 | 59/T | 42.6 ± 1.6 | 92/C | 40.1 ± 1.3 | $2.35 \times 10^{-1}$ | 1.2 | $2.74 \times 10^{-1}$ | 1.1 | $2.95 \times 10^{-1}$ | | Promoter polymorphism | Moderate | | |
| 1 | 1692992 | 136/G | 41.6 ± 1.1 | 15/A | 36.7 ± 2.2 | $1.39 \times 10^{-1}$ | 1.6 | $2.08 \times 10^{-1}$ | 2.7 | $1.03 \times 10^{-1}$ | | Promoter polymorphism | Moderate | | |
| 1 | 1692993 | 136/G | 41.6 ± 1.1 | 15/A | 36.7 ± 2.2 | $1.39 \times 10^{-1}$ | 1.6 | $2.08 \times 10^{-1}$ | 2.7 | $1.03 \times 10^{-1}$ | | Promoter polymorphism | Moderate | | |
| 1 | 1692994 | 137/A | 41.6 ± 1.1 | 14/G | 36.5 ± 2.3 | $1.40 \times 10^{-1}$ | 1.6 | $2.06 \times 10^{-1}$ | 2.0 | $1.55 \times 10^{-1}$ | | Promoter polymorphism | Moderate | | |
| 1 | 4695334 | 10/T | 42.1 ± 3.7 | 141/C | 41 ± 1.1 | $7.78 \times 10^{-1}$ | 0.2 | $6.47 \times 10^{-1}$ | 1.0 | $3.18 \times 10^{-1}$ | | Amino acid substitution (Glu62Lys) | Moderate | AT1G13700 | *PGL1* |
| 1 | 4695495 | 148/C | 41.2 ± 1.1 | 3/T | 34.8 ± 5.7 | $3.66 \times 10^{-1}$ | 0.9 | $3.40 \times 10^{-1}$ | 0.4 | $5.23 \times 10^{-1}$ | | Amino acid substitution (Ala36Thr) | Moderate | | |
| 1 | 4695719 | 106/T | 42.2 ± 1.3 | 45/G | 38.4 ± 1.6 | $7.52 \times 10^{-2}$ | 2.0 | $1.58 \times 10^{-1}$ | 0.4 | $5.10 \times 10^{-1}$ | | Promoter polymorphism | Moderate | | |
| 1 | 4695758 | 104/TA | 41.2 ± 1.3 | 47/T | 40.9 ± 1.6 | $8.92 \times 10^{-1}$ | 0.0 | $9.11 \times 10^{-1}$ | 0.1 | $7.77 \times 10^{-1}$ | | Promoter polymorphism | Moderate | | |
| 1 | 4695839 | 61/A | 41.9 ± 1.4 | 90/G | 40.5 ± 1.4 | $5.08 \times 10^{-1}$ | 0.2 | $6.56 \times 10^{-1}$ | 0.4 | $5.31 \times 10^{-1}$ | | Promoter polymorphism | Moderate | | |
| 1 | 4696143 | 128/T | 41.2 ± 1.2 | 23/A | 40.2 ± 2 | $7.19 \times 10^{-1}$ | 0.6 | $4.52 \times 10^{-1}$ | 0.5 | $4.98 \times 10^{-1}$ | | Promoter polymorphism | Moderate | | |
| 1 | 4697671 | 28/A | 46.9 ± 2.8 | 123/G | 39.8 ± 1.1 | **$4.42 \times 10^{-3}$** | **8.4** | **$4.39 \times 10^{-3}$** | **5.7** | **$1.78 \times 10^{-2}$** | ✓ | Synonymous polymorphism | Moderate | | |
| 1 | 4704734 | 11/A | 41.6 ± 3.4 | 140/T | 41.1 ± 1.1 | $8.91 \times 10^{-1}$ | 0.4 | $5.37 \times 10^{-1}$ | 1.3 | $2.56 \times 10^{-1}$ | | Promoter polymorphism | Moderate | AT1G13710 | *CYP78A5* |
| 1 | 4704775 | 140/T | 41.2 ± 1.1 | 11/A | 39.7 ± 3.7 | $7.01 \times 10^{-1}$ | 1.5 | $2.18 \times 10^{-1}$ | 2.3 | $1.33 \times 10^{-1}$ | | Promoter polymorphism | Moderate | | |
| 1 | 4704776 | 139/A | 41.2 ± 1.1 | 12/T | 40.1 ± 3.5 | $7.74 \times 10^{-1}$ | 1.0 | $3.12 \times 10^{-1}$ | 2.0 | $1.63 \times 10^{-1}$ | | Promoter polymorphism | Moderate | | |
| 1 | 4705207 | 138/T | 41.8 ± 1.1 | 13/TG | 33.4 ± 2.9 | **$1.65 \times 10^{-2}$** | **4.1** | **$4.51 \times 10^{-2}$** | 2.1 | $1.52 \times 10^{-1}$ | | Promoter polymorphism | Moderate | | |
| 1 | 4705426 | 138/T | 41.2 ± 1.1 | 13/A | 40.1 ± 3.3 | $7.58 \times 10^{-1}$ | 0.9 | $3.54 \times 10^{-1}$ | 1.5 | $2.23 \times 10^{-1}$ | | Promoter polymorphism | Moderate | | |

| | | | | | | | | | | | | | | | |
|---|---|---|---|---|---|---|---|---|---|---|---|---|---|---|---|
| 1 | 4705512 | 11/A | 41.6 ± 3.4 | 140/C | 41.1 ± 1.1 | $8.91 \times 10^{-1}$ | 0.4 | $5.37 \times 10^{-1}$ | 1.3 | $2.56 \times 10^{-1}$ | | Promoter polymorphism | Moderate | | |
| 1 | 8658652 | 76/G | 43 ± 1.5 | 75/T | 39.2 ± 1.4 | **$5.73 \times 10^{-2}$** | **5.8** | **$1.78 \times 10^{-2}$** | 1.9 | $1.66 \times 10^{-1}$ | | Amino acid substitution (Asp282Glu) | Moderate | AT1G24430 | |
| 1 | 8659236 | 13/C | 47.5 ± 3.7 | 138/T | 40.5 ± 1.1 | **$4.69 \times 10^{-2}$** | **6.9** | **$9.71 \times 10^{-3}$** | 1.3 | $2.61 \times 10^{-1}$ | | Amino acid substitution (Val88Met) | Moderate | | |
| 1 | 8659280 | 16/T | 45 ± 3.3 | 135/A | 40.6 ± 1.1 | $1.74 \times 10^{-1}$ | **3.6** | $6.01 \times 10^{-2}$ | 0.3 | $5.63 \times 10^{-1}$ | | Amino acid substitution (Tyr73Phe) | Moderate | | |
| 1 | 8659392 | 12/C | 48.3 ± 4 | 139/A | 40.5 ± 1.1 | $3.10 \times 10^{-2}$ | **6.4** | **$1.23 \times 10^{-2}$** | **3.1** | $7.91 \times 10^{-2}$ | | Amino acid substitution (Ala36Ser) | Moderate | | |
| 1 | 8659430 | 15/G | 47.4 ± 3.8 | 136/A | 40.4 ± 1.1 | **$3.41 \times 10^{-2}$** | **5.6** | **$1.89 \times 10^{-2}$** | 3.0 | $8.62 \times 10^{-2}$ | | Amino acid substitution (Pro23Leu) | Moderate | | |
| 1 | 8659449 | 16/G | 43.2 ± 3.5 | 135/A | 40.8 ± 1.1 | $4.61 \times 10^{-1}$ | 1.5 | $2.17 \times 10^{-1}$ | 0.4 | $5.09 \times 10^{-1}$ | | Amino acid substitution (Pro17Ser) | Moderate | | |
| 1 | 8659479 | 29/C | 42.8 ± 2.4 | 122/T | 40.7 ± 1.1 | $4.10 \times 10^{-1}$ | 1.6 | $2.08 \times 10^{-1}$ | 0.4 | $5.42 \times 10^{-1}$ | | Amino acid substitution (Val7Ile) | Moderate | | |
| 1 | 8659493 | 27/C | 45.1 ± 2.2 | 124/T | 40.2 ± 1.1 | $6.07 \times 10^{-2}$ | **5.1** | **$2.48 \times 10^{-2}$** | 2.7 | $1.04 \times 10^{-1}$ | | Amino acid substitution (Gly2Glu) | Moderate | | |
| 1 | 8659521 | 22/T | 43 ± 2.6 | 129/C | 40.8 ± 1.1 | $4.32 \times 10^{-1}$ | 1.1 | $2.87 \times 10^{-1}$ | 0.5 | $4.82 \times 10^{-1}$ | | Promoter polymorphism | Moderate | | |
| 1 | 8659542 | 29/C | 43.2 ± 2.2 | 122/A | 40.6 ± 1.2 | $2.89 \times 10^{-1}$ | 1.4 | $2.46 \times 10^{-1}$ | 0.5 | $4.71 \times 10^{-1}$ | | Promoter polymorphism | Moderate | | |
| 1 | 8659552 | 32/C | 41.9 ± 2.3 | 119/A | 40.9 ± 1.2 | $6.57 \times 10^{-1}$ | 0.8 | $3.81 \times 10^{-1}$ | 0.0 | $9.89 \times 10^{-1}$ | | Promoter polymorphism | Moderate | | |
| 1 | 8659563 | 26/A | 42.4 ± 2.7 | 125/G | 40.8 ± 1.1 | $5.33 \times 10^{-1}$ | 0.6 | $4.34 \times 10^{-1}$ | 0.0 | $8.70 \times 10^{-1}$ | | Promoter polymorphism | Moderate | | |
| 1 | 8659565 | 88/AT | 41.2 ± 1.4 | 63/A | 40.9 ± 1.6 | $8.49 \times 10^{-1}$ | 0.0 | $8.27 \times 10^{-1}$ | 0.0 | $8.49 \times 10^{-1}$ | | Promoter polymorphism | Moderate | | |
| 1 | 8659670 | 122/T | 41.3 ± 1.2 | 29/C | 40.3 ± 2.3 | $6.98 \times 10^{-1}$ | 0.0 | $8.55 \times 10^{-1}$ | 0.1 | $7.55 \times 10^{-1}$ | | Promoter polymorphism | Moderate | | |
| 1 | 8659830 | 18/T | 42.1 ± 3.8 | 133/C | 41 ± 1.1 | $7.14 \times 10^{-1}$ | 0.8 | $3.83 \times 10^{-1}$ | 0.1 | $7.79 \times 10^{-1}$ | | Promoter polymorphism | Moderate | | |
| 1 | 8659841 | 117/C | 42.3 ± 1.2 | 34/A | 36.9 ± 1.8 | $2.32 \times 10^{-2}$ | **6.4** | **$1.23 \times 10^{-2}$** | 2.0 | $1.64 \times 10^{-1}$ | ✓ | Promoter polymorphism | Moderate | | |
| 1 | 8659905 | 28/A | 43.2 ± 2.7 | 123/T | 40.6 ± 1.1 | $3.18 \times 10^{-1}$ | 1.5 | $2.23 \times 10^{-1}$ | 0.9 | $3.46 \times 10^{-1}$ | | Promoter polymorphism | Moderate | | |
| 1 | 8659910 | 25/G | 42 ± 2.7 | 126/T | 40.9 ± 1.1 | $6.69 \times 10^{-1}$ | 0.3 | $5.82 \times 10^{-1}$ | 0.5 | $4.95 \times 10^{-1}$ | | Promoter polymorphism | Moderate | | |
| 1 | 8659920 | 132/G | 41.4 ± 1.1 | 19/C | 38.7 ± 3 | $3.64 \times 10^{-1}$ | 0.7 | $4.11 \times 10^{-1}$ | 2.0 | $1.57 \times 10^{-1}$ | | Promoter polymorphism | Moderate | | |
| 1 | 8659923 | 129/T | 41.4 ± 1.1 | 22/A | 39.4 ± 2.8 | $4.77 \times 10^{-1}$ | 0.6 | $4.31 \times 10^{-1}$ | 1.3 | $2.56 \times 10^{-1}$ | | Promoter polymorphism | Moderate | | |
| 1 | 8659930 | 120/A | 41.6 ± 1.2 | 31/T | 39.1 ± 2.2 | $3.08 \times 10^{-1}$ | 0.7 | $4.12 \times 10^{-1}$ | 2.0 | $1.56 \times 10^{-1}$ | | Promoter polymorphism | Moderate | | |
| 1 | 8660127 | 91/TC | 41.5 ± 1.3 | 60/T | 40.4 ± 1.7 | $5.93 \times 10^{-1}$ | 0.4 | $5.18 \times 10^{-1}$ | 0.1 | $7.08 \times 10^{-1}$ | | Promoter polymorphism | Moderate | | |
| 1 | 8660130 | 138/G | 41.5 ± 1.1 | 13/A | 36.9 ± 3.3 | $1.92 \times 10^{-1}$ | 1.4 | $2.39 \times 10^{-1}$ | 1.6 | $2.03 \times 10^{-1}$ | | Promoter polymorphism | Moderate | | |
| 1 | 8660135 | 124/A | 41.7 ± 1.1 | 27/G | 38.1 ± 2.4 | $1.54 \times 10^{-1}$ | 1.4 | $2.41 \times 10^{-1}$ | 2.4 | $1.27 \times 10^{-1}$ | | Promoter polymorphism | Moderate | | |
| 1 | 8660312 | 60/C | 42.2 ± 1.6 | 91/T | 40.3 ± 1.3 | $3.54 \times 10^{-1}$ | 1.1 | $3.01 \times 10^{-1}$ | 0.1 | $7.40 \times 10^{-1}$ | | Promoter polymorphism | Moderate | | |
| 1 | 8660323 | 55/AT | 41.3 ± 1.8 | 96/A | 40.9 ± 1.3 | $8.44 \times 10^{-1}$ | 0.0 | $9.86 \times 10^{-1}$ | 0.0 | $9.53 \times 10^{-1}$ | | Promoter polymorphism | Moderate | | |
| 1 | 8660325 | 69/A | 41.2 ± 1.5 | 82/T | 41 ± 1.4 | $9.43 \times 10^{-1}$ | 0.0 | $8.56 \times 10^{-1}$ | 0.1 | $7.89 \times 10^{-1}$ | | Promoter polymorphism | Moderate | | |
| 1 | 8660343 | 74/T | 41.8 ± 1.5 | 77/A | 40.4 ± 1.4 | $4.73 \times 10^{-1}$ | 0.3 | $6.15 \times 10^{-1}$ | 1.6 | $2.02 \times 10^{-1}$ | | Promoter polymorphism | Moderate | | |
| 1 | 8660367 | 51/C | 42.9 ± 1.8 | 100/G | 40.2 ± 1.2 | $1.98 \times 10^{-1}$ | 2.0 | $1.64 \times 10^{-1}$ | 0.5 | $4.97 \times 10^{-1}$ | | Promoter polymorphism | Moderate | | |
| 1 | 8660407 | 61/C | 43.2 ± 1.6 | 88/G | 39.4 ± 1.4 | $6.58 \times 10^{-2}$ | 2.5 | $8.74 \times 10^{-2}$ | 1.6 | $2.00 \times 10^{-1}$ | | Promoter polymorphism | Moderate | | |
| 1 | 8660437 | 97/A | 41.5 ± 1.2 | 54/C | 40.4 ± 1.8 | $6.17 \times 10^{-1}$ | 1.2 | $2.69 \times 10^{-1}$ | 0.1 | $8.06 \times 10^{-1}$ | | Promoter polymorphism | Moderate | | |
| 1 | 8660438 | 96/C | 41.6 ± 1.2 | 55/T | 40.1 ± 1.8 | $4.71 \times 10^{-1}$ | 2.2 | $1.44 \times 10^{-1}$ | 0.7 | $3.95 \times 10^{-1}$ | | Promoter polymorphism | Moderate | | |
| 1 | 8660455 | 121/T | 41.6 ± 1.1 | 30/C | 39.1 ± 2.6 | $3.30 \times 10^{-1}$ | **4.2** | **$4.25 \times 10^{-2}$** | **3.4** | $6.69 \times 10^{-2}$ | | Promoter polymorphism | Moderate | | |
| 1 | 8660470 | 134/T | 41.2 ± 1.1 | 17/A | 40 ± 3.3 | $6.86 \times 10^{-1}$ | 0.7 | $4.05 \times 10^{-1}$ | 0.0 | $9.13 \times 10^{-1}$ | | Promoter polymorphism | Moderate | | |
| 1 | 8660476 | 20/C | 42.5 ± 3.4 | 131/T | 40.9 ± 1.1 | $5.83 \times 10^{-1}$ | 0.0 | $8.47 \times 10^{-1}$ | 0.4 | $5.20 \times 10^{-1}$ | | Promoter polymorphism | Moderate | | |
| 1 | 8660479 | 21/G | 43.7 ± 3.1 | 130/T | 40.7 ± 1.1 | $2.91 \times 10^{-1}$ | 0.1 | $7.47 \times 10^{-1}$ | 1.7 | $2.00 \times 10^{-1}$ | | Promoter polymorphism | Moderate | | |
| 1 | 8660484 | 20/T | 43.8 ± 3.3 | 131/A | 40.7 ± 1.1 | $2.95 \times 10^{-1}$ | 0.1 | $7.39 \times 10^{-1}$ | 1.1 | $2.98 \times 10^{-1}$ | | Promoter polymorphism | Moderate | | |
| 1 | 23173422 | 39/A | 41.2 ± 2.1 | 112/C | 41.1 ± 1.2 | $9.56 \times 10^{-1}$ | 0.0 | $8.88 \times 10^{-1}$ | 0.2 | $6.42 \times 10^{-1}$ | | Promoter polymorphism | Moderate | AT1G75030 | *TLP-3* |
| 1 | 23173457 | 40/A | 41.5 ± 2.3 | 111/G | 40.9 ± 1.1 | $7.92 \times 10^{-1}$ | 0.0 | $9.18 \times 10^{-1}$ | 0.3 | $5.86 \times 10^{-1}$ | | Promoter polymorphism | Moderate | | |
| 1 | 23173461 | 110/G | 41.8 ± 1.2 | 41/A | 39.3 ± 2.2 | $2.70 \times 10^{-1}$ | 1.5 | $2.18 \times 10^{-1}$ | 2.3 | $1.32 \times 10^{-1}$ | | Promoter polymorphism | Moderate | | |
| 1 | 23173467 | 105/A | 42.3 ± 1.2 | 46/C | 38.3 ± 1.8 | $6.18 \times 10^{-2}$ | 2.7 | $1.02 \times 10^{-1}$ | 1.8 | $1.83 \times 10^{-1}$ | | Promoter polymorphism | Moderate | | |

| | | | | | | | | | | | | | | | |
|---|---|---|---|---|---|---|---|---|---|---|---|---|---|---|---|
| 1 | 23173492 | 15/T | 44.3 ± 2.6 | 136/C | 40.7 ± 1.1 | $2.85 \times 10^{-1}$ | 2.3 | $1.32 \times 10^{-1}$ | 1.0 | $3.14 \times 10^{-1}$ | | Promoter polymorphism | Moderate | | |
| 1 | 23173505 | 11/T | 41.9 ± 3.7 | 140/C | 41 ± 1.1 | $8.16 \times 10^{-1}$ | 0.2 | $6.23 \times 10^{-1}$ | 0.7 | $4.21 \times 10^{-1}$ | | Promoter polymorphism | Moderate | | |
| 1 | 23173513 | 15/T | 42.8 ± 2.6 | 136/C | 40.9 ± 1.1 | $5.63 \times 10^{-1}$ | 1.5 | $2.30 \times 10^{-1}$ | 2.1 | $1.46 \times 10^{-1}$ | | Promoter polymorphism | Moderate | | |
| 1 | 23173568 | 11/A | 41.5 ± 3.7 | 140/G | 41.1 ± 1.1 | $9.17 \times 10^{-1}$ | 0.2 | $6.99 \times 10^{-1}$ | 0.1 | $7.41 \times 10^{-1}$ | | Promoter polymorphism | Moderate | | |
| 1 | 23173583 | 119/G | 41.5 ± 1.2 | 32/A | 39.6 ± 2 | $4.25 \times 10^{-1}$ | 0.4 | $5.07 \times 10^{-1}$ | 0.0 | $9.92 \times 10^{-1}$ | | Promoter polymorphism | Moderate | | |
| 1 | 23173608 | 14/T | 43.6 ± 2.9 | 137/C | 40.8 ± 1.1 | $4.13 \times 10^{-1}$ | 0.8 | $3.58 \times 10^{-1}$ | 1.1 | $2.91 \times 10^{-1}$ | | Promoter polymorphism | Moderate | | |
| 1 | 23173697 | 127/C | 41.6 ± 1.2 | 24/T | 38.5 ± 2.2 | $2.54 \times 10^{-1}$ | 0.9 | $3.58 \times 10^{-1}$ | 0.0 | $8.56 \times 10^{-1}$ | | Promoter polymorphism | Moderate | | |
| 1 | 23173793 | 13/T | 41.5 ± 4.8 | 138/C | 41.1 ± 1 | $9.09 \times 10^{-1}$ | 0.4 | $5.41 \times 10^{-1}$ | 0.4 | $5.35 \times 10^{-1}$ | | Promoter polymorphism | Moderate | | |
| 1 | 23173939 | 17/G | 45.2 ± 2.5 | 134/A | 40.6 ± 1.1 | $1.37 \times 10^{-1}$ | 2.1 | $1.48 \times 10^{-1}$ | 0.9 | $3.38 \times 10^{-1}$ | | Promoter polymorphism | Moderate | | |
| 1 | 23173972 | 103/A | 41.1 ± 1.2 | 48/G | 41.1 ± 1.9 | $9.84 \times 10^{-1}$ | 0.1 | $7.21 \times 10^{-1}$ | 0.0 | $9.26 \times 10^{-1}$ | | Promoter polymorphism | Moderate | | |
| 1 | 23173979 | 103/T | 41.2 ± 1.2 | 48/C | 40.9 ± 1.9 | $8.73 \times 10^{-1}$ | 0.2 | $6.43 \times 10^{-1}$ | 0.1 | $8.18 \times 10^{-1}$ | | Promoter polymorphism | Moderate | | |
| 1 | 28174614 | 137/T | 41.3 ± 1.1 | 14/A | 39 ± 3.5 | $5.04 \times 10^{-1}$ | 0.5 | $4.83 \times 10^{-1}$ | 0.6 | $4.36 \times 10^{-1}$ | | Amino acid substitution (Tyr34Asn) | Moderate | | |
| 1 | 28179461 | 14/A | 41.4 ± 5 | 137/T | 41.1 ± 1 | $9.31 \times 10^{-1}$ | 0.1 | $7.92 \times 10^{-1}$ | 0.0 | $8.76 \times 10^{-1}$ | | Promoter polymorphism | Moderate | AT1G75050 | |
| 1 | 28179649 | 134/G | 41.2 ± 1.1 | 17/C | 40.6 ± 2.6 | $8.53 \times 10^{-1}$ | 0.1 | $8.16 \times 10^{-1}$ | 0.0 | $9.27 \times 10^{-1}$ | | Promoter polymorphism | Moderate | | |
| 1 | 28179813 | 13/G | 48.4 ± 2.3 | 138/T | 40.4 ± 1.1 | **$2.39 \times 10^{-2}$** | 2.7 | $1.03 \times 10^{-1}$ | 1.7 | $1.96 \times 10^{-1}$ | | Promoter polymorphism | Moderate | | |
| 1 | 28180813 | 127/G | 41.9 ± 1.1 | 24/T | 36.8 ± 2.4 | $5.95 \times 10^{-2}$ | **4.2** | **$4.30 \times 10^{-2}$** | 1.1 | $3.07 \times 10^{-1}$ | | Amino acid substitution (Glu164Asp) | Moderate | | |
| 1 | 28181889 | 150/T | 41.1 ± 1 | 1/A | 36.9 ± 0 | | 0.2 | $6.74 \times 10^{-1}$ | 0.1 | $7.51 \times 10^{-1}$ | | Splice acceptor variant | **High** | AT1G75060 | |
| 1 | 28183339 | 25/A | 42.7 ± 3 | 126/C | 40.8 ± 1.1 | $4.82 \times 10^{-1}$ | 0.4 | $5.36 \times 10^{-1}$ | 0.3 | $5.95 \times 10^{-1}$ | | 5'-UTR polymorphism | Moderate | | |
| 1 | 28183613 | 104/T | 41.3 ± 1.3 | 47/G | 40.6 ± 1.8 | $7.50 \times 10^{-1}$ | 0.8 | $3.61 \times 10^{-1}$ | 0.0 | $9.55 \times 10^{-1}$ | | Promoter polymorphism | Moderate | | |
| 1 | 28183757 | 13/G | 49.2 ± 2.3 | 138/T | 40.3 ± 1.1 | $1.18 \times 10^{-2}$ | **3.9** | $5.15 \times 10^{-2}$ | 2.8 | $9.65 \times 10^{-2}$ | | Promoter polymorphism | Moderate | | |
| 1 | 28183796 | 114/T | 42.6 ± 1.2 | 37/C | 36.3 ± 1.8 | **$6.01 \times 10^{-3}$** | **9.8** | **$2.13 \times 10^{-3}$** | **4.4** | **$3.79 \times 10^{-2}$** | | Promoter polymorphism | Moderate | | |
| 1 | 28184035 | 20/AAC | 43.7 ± 3 | 131/A | 40.7 ± 1.1 | $2.96 \times 10^{-1}$ | 0.5 | $4.81 \times 10^{-1}$ | 0.0 | $9.65 \times 10^{-1}$ | | Promoter polymorphism | Moderate | | |
| 1 | 28184066 | 54/T | 42.5 ± 1.9 | 97/C | 40.3 ± 1.2 | $2.78 \times 10^{-1}$ | 1.2 | $2.71 \times 10^{-1}$ | 0.4 | $5.09 \times 10^{-1}$ | | Promoter polymorphism | Moderate | | |
| 1 | 28184206 | 60/G | 42.6 ± 1.8 | 91/T | 40.1 ± 1.2 | $2.29 \times 10^{-1}$ | 1.4 | $2.33 \times 10^{-1}$ | 0.0 | $8.47 \times 10^{-1}$ | | Promoter polymorphism | Moderate | | |
| 1 | 28184486 | 89/A | 43.2 ± 1.4 | 62/G | 38 ± 1.5 | $9.22 \times 10^{-3}$ | **7.3** | **$7.81 \times 10^{-3}$** | **3.2** | $7.50 \times 10^{-2}$ | ✓ | Promoter polymorphism | Moderate | | |
| 1 | 28184722 | 26/C | 41.3 ± 2.8 | 125/T | 41 ± 1.1 | $9.25 \times 10^{-1}$ | 0.0 | $8.61 \times 10^{-1}$ | 0.0 | $8.24 \times 10^{-1}$ | | Promoter polymorphism | Moderate | AT1G75080 | *BZR1* |
| 1 | 28184778 | 27/T | 42.2 ± 2.9 | 124/C | 40.8 ± 1.1 | $5.90 \times 10^{-1}$ | 0.4 | $5.22 \times 10^{-1}$ | 0.1 | $8.23 \times 10^{-1}$ | | Promoter polymorphism | Moderate | | |
| 1 | 28184895 | 53/A | 42.1 ± 1.7 | 98/T | 40.5 ± 1.3 | $4.49 \times 10^{-1}$ | 1.6 | $2.12 \times 10^{-1}$ | 0.7 | $4.06 \times 10^{-1}$ | | Promoter polymorphism | Moderate | | |
| 1 | 28185008 | 39/T | 44.5 ± 2.4 | 112/G | 39.9 ± 1.1 | **$4.37 \times 10^{-2}$** | **3.2** | $7.43 \times 10^{-2}$ | 0.8 | $3.61 \times 10^{-1}$ | | Promoter polymorphism | Moderate | | |
| 1 | 28185029 | 92/A | 41.5 ± 1.4 | 59/G | 40.5 ± 1.6 | $6.13 \times 10^{-1}$ | 0.1 | $7.19 \times 10^{-1}$ | 0.2 | $6.47 \times 10^{-1}$ | | Promoter polymorphism | Moderate | | |
| 1 | 28185167 | 125/T | 41.9 ± 1.1 | 26/C | 37 ± 2.2 | $6.26 \times 10^{-2}$ | **3.3** | $7.06 \times 10^{-2}$ | 0.5 | $4.68 \times 10^{-1}$ | | Promoter polymorphism | Moderate | | |
| 1 | 28185227 | 12/T | 41.5 ± 4.6 | 139/G | 41.1 ± 1.1 | $8.97 \times 10^{-1}$ | 0.2 | $6.19 \times 10^{-1}$ | 0.1 | $7.10 \times 10^{-1}$ | | Promoter polymorphism | Moderate | | |
| 1 | 28185241 | 26/A | 42.1 ± 3 | 125/G | 40.9 ± 1.1 | $6.39 \times 10^{-1}$ | 0.3 | $5.81 \times 10^{-1}$ | 0.0 | $8.84 \times 10^{-1}$ | | Promoter polymorphism | Moderate | | |
| 1 | 28185324 | 21/A | 42 ± 2.3 | 130/C | 40.9 ± 1.1 | $6.99 \times 10^{-1}$ | 0.3 | $6.17 \times 10^{-1}$ | 0.0 | $8.46 \times 10^{-1}$ | | Promoter polymorphism | Moderate | | |
| 2 | 16922707 | 46/C | 42.2 ± 2 | 105/T | 40.6 ± 1.2 | $4.54 \times 10^{-1}$ | 0.2 | $6.75 \times 10^{-1}$ | 0.1 | $7.48 \times 10^{-1}$ | | Promoter polymorphism | Moderate | AT2G40520 | |
| 2 | 16923075 | 9/CT | 44.8 ± 2.9 | 142/C | 40.9 ± 1.1 | $3.50 \times 10^{-1}$ | 1.3 | $2.52 \times 10^{-1}$ | 0.8 | $3.76 \times 10^{-1}$ | | 5'-UTR polymorphism | Moderate | | |
| 2 | 16923451 | 147/T | 41.3 ± 1.1 | 4/C | 34.1 ± 2.5 | $2.46 \times 10^{-1}$ | 1.5 | $2.26 \times 10^{-1}$ | 0.3 | $5.87 \times 10^{-1}$ | | 5'-UTR polymorphism | Moderate | | |
| 2 | 16923512 | 145/TGG | 41.2 ± 1.1 | 6/TGGG | 38.9 ± 5.1 | $6.53 \times 10^{-1}$ | 0.3 | $5.77 \times 10^{-1}$ | 0.9 | $3.51 \times 10^{-1}$ | | 5'-UTR polymorphism | Moderate | | |
| 2 | 16923532 | 6/C | 44.9 ± 5.7 | 145/T | 40.9 ± 1.1 | $4.30 \times 10^{-1}$ | 0.6 | $4.48 \times 10^{-1}$ | 0.1 | $7.72 \times 10^{-1}$ | | 5'-UTR polymorphism | Moderate | | |
| 2 | 16924347 | 144/A | 41.3 ± 1.1 | 7/C | 37.3 ± 4.6 | $4.02 \times 10^{-1}$ | 0.8 | $3.85 \times 10^{-1}$ | 0.4 | $5.28 \times 10^{-1}$ | | Amino acid substitution (Lys169Gln) | Moderate | | |
| 2 | 16925429 | 150/C | 41.2 ± 1 | 1/T | 25.3 ± 0 | | 1.0 | $3.15 \times 10^{-1}$ | 0.0 | $9.24 \times 10^{-1}$ | | Stop gained (Gln452*) | **High** | | |
| 2 | 16934262 | 143/G | 41.3 ± 1.1 | 8/A | 37.7 ± 4.4 | $4.16 \times 10^{-1}$ | 0.6 | $4.27 \times 10^{-1}$ | 0.0 | $8.60 \times 10^{-1}$ | | Promoter polymorphism | Moderate | AT2G40550 | *ETG1* |

| | | | | | | | | | | | | | | |
|---|---|---|---|---|---|---|---|---|---|---|---|---|---|---|
| 2 | 16934522 | 142/G | 41.1 ± 1.1 | 9/GA | 40.9 ± 3.6 | $9.66 \times 10^{-1}$ | 0.2 | $6.45 \times 10^{-1}$ | 1.3 | $2.65 \times 10^{-1}$ | | Promoter polymorphism | Moderate | |
| 2 | 16934686 | 107/T | 41.9 ± 1.3 | 44/TA | 39.2 ± 1.7 | $2.26 \times 10^{-1}$ | 3.0 | $8.61 \times 10^{-2}$ | 1.4 | $2.32 \times 10^{-1}$ | | Promoter polymorphism | Moderate | |
| 2 | 16934776 | 93/G | 42.6 ± 1.4 | 58/C | 38.6 ± 1.5 | **$4.87 \times 10^{-2}$** | **5.2** | **$2.36 \times 10^{-2}$** | 2.9 | $8.92 \times 10^{-2}$ | | Promoter polymorphism | Moderate | |
| 2 | 16934867 | 17/T | 42.3 ± 2.3 | 134/TG | 40.9 ± 1.1 | $6.76 \times 10^{-1}$ | 0.0 | $8.28 \times 10^{-1}$ | 0.0 | $9.50 \times 10^{-1}$ | | 5'-UTR polymorphism | Moderate | |
| 2 | 16934889 | 99/C | 42 ± 1.3 | 52/T | 39.4 ± 1.5 | $2.23 \times 10^{-1}$ | 2.5 | $1.18 \times 10^{-1}$ | 0.6 | $4.41 \times 10^{-1}$ | | 5'-UTR polymorphism | Moderate | |
| 2 | 16936002 | 148/C | 41.2 ± 1 | 3/G | 36.2 ± 8.4 | $4.88 \times 10^{-1}$ | 0.8 | $3.87 \times 10^{-1}$ | 0.4 | $5.14 \times 10^{-1}$ | | Amino acid substitution (Ser183Cys) | Moderate | |
| 2 | 16936277 | 144/A | 41.2 ± 1.1 | 7/T | 39.4 ± 3.6 | $7.08 \times 10^{-1}$ | 0.1 | $8.16 \times 10^{-1}$ | 0.1 | $7.18 \times 10^{-1}$ | | Amino acid substitution (Ser213Cys) | Moderate | |
| 2 | 16936562 | 131/G | 41.3 ± 1.2 | 20/A | 39.5 ± 2 | $5.30 \times 10^{-1}$ | 1.5 | $2.28 \times 10^{-1}$ | 0.7 | $4.17 \times 10^{-1}$ | | Amino acid substitution (Gly274Glu) | Moderate | |
| 2 | 16936596 | 105/A | 42.8 ± 1.3 | 46/G | 37.2 ± 1.6 | **$9.68 \times 10^{-3}$** | **10.0** | **$1.91 \times 10^{-3}$** | **5.3** | **$2.28 \times 10^{-2}$** | ✓ | Synonymous polymorphism | Moderate | |
| 2 | 16937943 | 131/G | 41.1 ± 1.1 | 20/A | 41 ± 2.3 | $9.64 \times 10^{-1}$ | 0.5 | $4.67 \times 10^{-1}$ | 0.1 | $7.87 \times 10^{-1}$ | | Amino acid substitution (Met560Ile) | Moderate | |
| 3 | 6008820 | 24/C | 43.2 ± 3 | 127/G | 40.7 ± 1.1 | $3.64 \times 10^{-1}$ | 0.0 | $8.33 \times 10^{-1}$ | 0.1 | $8.10 \times 10^{-1}$ | | Promoter polymorphism | Moderate | AT3G17570 |
| 3 | 6008940 | 128/G | 41.4 ± 1.1 | 23/T | 39.2 ± 2.5 | $4.11 \times 10^{-1}$ | 0.5 | $4.95 \times 10^{-1}$ | 0.2 | $6.35 \times 10^{-1}$ | | Promoter polymorphism | Moderate | |
| 3 | 6009106 | 18/T | 41.9 ± 3.2 | 133/G | 41 ± 1.1 | $7.55 \times 10^{-1}$ | 1.1 | $2.86 \times 10^{-1}$ | 0.9 | $3.39 \times 10^{-1}$ | | Promoter polymorphism | Moderate | |
| 3 | 6009109 | 131/C | 41.4 ± 1.1 | 20/T | 39.3 ± 2.7 | $4.84 \times 10^{-1}$ | 0.7 | $3.91 \times 10^{-1}$ | 0.4 | $5.11 \times 10^{-1}$ | | Promoter polymorphism | Moderate | |
| 3 | 6009548 | 132/G | 41.3 ± 1.1 | 19/A | 39.4 ± 2.7 | $5.11 \times 10^{-1}$ | 0.6 | $4.23 \times 10^{-1}$ | 0.2 | $6.85 \times 10^{-1}$ | | Amino acid substitution (Val46Met) | Moderate | |
| 3 | 6009606 | 4/GT | 50.7 ± 1.9 | 147/G | 40.8 ± 1.1 | $1.10 \times 10^{-1}$ | 1.80 | $1.82 \times 10^{-1}$ | 2.2 | $1.10 \times 10^{-1}$ | | Frameshift variant (Ser65 Val66fs) | **High** | |
| 3 | 6009710 | 148/G | 41.1 ± 1.1 | 3/T | 38.8 ± 2.6 | $7.43 \times 10^{-1}$ | 0.00 | $9.77 \times 10^{-1}$ | 0.4 | $7.43 \times 10^{-1}$ | | Amino acid substitution (Gly100Cys) | Moderate | |
| 3 | 6009787 | 17/A | 47.6 ± 3.8 | 134/T | 40.3 ± 1 | **$1.89 \times 10^{-2}$** | **6.2** | **$1.38 \times 10^{-2}$** | 2.9 | $9.08 \times 10^{-2}$ | ✓ | Synonymous polymorphism | Moderate | |
| 3 | 6009899 | 6/A | 50.9 ± 5.5 | 145/G | 40.7 ± 1 | **$4.40 \times 10^{-2}$** | 1.9 | $1.68 \times 10^{-1}$ | 2.9 | $9.04 \times 10^{-2}$ | | Amino acid substitution (Asp163Asn) | Moderate | |
| 3 | 6009917 | 150/G | 41.2 ± 1 | 1/GA | 23.2 ± 0 | | 1.8 | $1.81 \times 10^{-1}$ | 2.0 | $1.59 \times 10^{-1}$ | | Frameshift variant (Glu169 Ile170fs) | **High** | |
| 3 | 6009996 | 3/G | 49.9 ± 2.5 | 148/T | 40.9 ± 1.1 | $2.07 \times 10^{-1}$ | 1.3 | $2.64 \times 10^{-1}$ | 1.2 | $2.85 \times 10^{-1}$ | | Amino acid substitution (Val195Gly) | Moderate | |
| 3 | 6010101 | 62/C | 43.5 ± 1.8 | 89/T | 39.4 ± 1.2 | **$4.52 \times 10^{-2}$** | **4.0** | **$4.79 \times 10^{-2}$** | 1.6 | $2.13 \times 10^{-1}$ | | Amino acid substitution (Leu230Pro) | Moderate | |
| 3 | 6010127 | 7/G | 50.3 ± 7.3 | 144/A | 40.6 ± 1 | **$4.05 \times 10^{-2}$** | **5.7** | **$1.80 \times 10^{-2}$** | 2.4 | $1.27 \times 10^{-1}$ | | Amino acid substitution (Asn239Asp) | Moderate | |
| 3 | 6010188 | 131/A | 41.3 ± 1.1 | 20/G | 39.8 ± 2.8 | $6.03 \times 10^{-1}$ | 0.4 | $5.17 \times 10^{-1}$ | 0.1 | $8.02 \times 10^{-1}$ | | Amino acid substitution (Glu259Gly) | Moderate | |
| 3 | 6010444 | 6/G | 49.3 ± 4.9 | 145/T | 40.7 ± 1.1 | $9.29 \times 10^{-2}$ | 0.9 | $3.38 \times 10^{-1}$ | **3.2** | **$4.50 \times 10^{-2}$** | | Stop gained (Tyr344*) | **High** | |
| 3 | 6010464 | 59/T | 43.5 ± 1.8 | 92/G | 39.6 ± 1.2 | $5.26 \times 10^{-2}$ | 3.0 | $8.73 \times 10^{-2}$ | 1.4 | $2.32 \times 10^{-1}$ | | Amino acid substitution (Trp351Leu) | Moderate | |
| 3 | 9670551 | 85/A | 42.9 ± 1.4 | 66/C | 38.7 ± 1.4 | **$3.37 \times 10^{-2}$** | **6.3** | **$1.31 \times 10^{-2}$** | 1.3 | $2.50 \times 10^{-1}$ | | Promoter polymorphism | Moderate | AT3G26420 |
| 3 | 9670560 | 84/G | 42.7 ± 1.4 | 67/A | 39.1 ± 1.4 | $7.69 \times 10^{-2}$ | **4.5** | **$3.49 \times 10^{-2}$** | 0.1 | $6.99 \times 10^{-1}$ | | Promoter polymorphism | Moderate | |
| 3 | 9670788 | 82/T | 43.1 ± 1.5 | 69/C | 38.7 ± 1.3 | **$2.64 \times 10^{-2}$** | **6.8** | **$9.86 \times 10^{-3}$** | 0.9 | $3.49 \times 10^{-1}$ | | Promoter polymorphism | Moderate | |
| 3 | 9670811 | 85/C | 42.9 ± 1.4 | 66/A | 38.8 ± 1.4 | **$4.19 \times 10^{-2}$** | **5.9** | **$1.67 \times 10^{-2}$** | 0.2 | $6.89 \times 10^{-1}$ | | Promoter polymorphism | Moderate | |
| 3 | 9670816 | 77/C | 42.9 ± 1.5 | 74/A | 39.2 ± 1.4 | $6.38 \times 10^{-2}$ | **4.7** | **$3.18 \times 10^{-2}$** | 0.2 | $6.94 \times 10^{-1}$ | | Promoter polymorphism | Moderate | |
| 3 | 9670881 | 83/A | 42.6 ± 1.5 | 68/C | 39.2 ± 1.3 | $8.70 \times 10^{-2}$ | **4.0** | **$4.78 \times 10^{-2}$** | 0.1 | $7.40 \times 10^{-1}$ | | Promoter polymorphism | Moderate | |
| 3 | 9671094 | 93/A | 42.7 ± 1.4 | 58/G | 38.6 ± 1.4 | **$4.49 \times 10^{-2}$** | **6.7** | **$1.07 \times 10^{-2}$** | 0.9 | $3.51 \times 10^{-1}$ | | Promoter polymorphism | Moderate | |
| 3 | 9671106 | 91/A | 42.7 ± 1.4 | 60/T | 38.6 ± 1.4 | **$4.08 \times 10^{-2}$** | **7.0** | **$9.26 \times 10^{-3}$** | 0.5 | $4.70 \times 10^{-1}$ | | Promoter polymorphism | Moderate | |
| 3 | 9671262 | 90/T | 42.5 ± 1.4 | 61/C | 39.1 ± 1.4 | $9.17 \times 10^{-2}$ | **4.7** | **$3.19 \times 10^{-2}$** | 0.1 | $7.04 \times 10^{-1}$ | | Promoter polymorphism | Moderate | |
| 3 | 9671288 | 72/T | 42.7 ± 1.6 | 79/C | 39.7 ± 1.3 | $1.32 \times 10^{-1}$ | **3.9** | **$4.97 \times 10^{-2}$** | 0.0 | $8.58 \times 10^{-1}$ | | Promoter polymorphism | Moderate | |
| 3 | 9671302 | 88/C | 42.4 ± 1.4 | 63/G | 39.3 ± 1.4 | $1.21 \times 10^{-1}$ | **4.2** | **$4.11 \times 10^{-2}$** | 0.0 | $9.33 \times 10^{-1}$ | | Promoter polymorphism | Moderate | |
| 3 | 9671412 | 134/C | 41.4 ± 1.1 | 17/T | 38.7 ± 2.9 | $3.95 \times 10^{-1}$ | 0.9 | $3.51 \times 10^{-1}$ | 0.3 | $5.76 \times 10^{-1}$ | | Promoter polymorphism | Moderate | |
| 3 | 9671428 | 84/A | 42.1 ± 1.4 | 67/T | 39.9 ± 1.4 | $2.79 \times 10^{-1}$ | 2.3 | $1.28 \times 10^{-1}$ | 1.6 | $2.08 \times 10^{-1}$ | | Promoter polymorphism | Moderate | |
| 3 | 9671501 | 37/CT | 45.6 ± 2.1 | 114/C | 39.6 ± 1.2 | **$9.34 \times 10^{-3}$** | **11.1** | **$1.12 \times 10^{-3}$** | **4.5** | **$3.66 \times 10^{-2}$** | | Promoter polymorphism | Moderate | |
| 3 | 9671524 | 6/T | 43 ± 5.5 | 145/C | 41 ± 1.1 | $7.01 \times 10^{-1}$ | 0.2 | $6.56 \times 10^{-1}$ | 0.3 | $5.81 \times 10^{-1}$ | | 5'-UTR polymorphism | Moderate | |
| 3 | 9671576 | 89/G | 42.6 ± 1.4 | 62/T | 38.9 ± 1.4 | $6.10 \times 10^{-2}$ | **5.6** | **$1.91 \times 10^{-2}$** | 0.0 | $9.77 \times 10^{-1}$ | | 5'-UTR polymorphism | Moderate | |

| | | | | | | | | | | | | | | |
|---|---|---|---|---|---|---|---|---|---|---|---|---|---|---|
| 3 | 9671582 | 86/C | 41.9 ± 1.4 | 64/T | 40 ± 1.5 | $3.70 \times 10^{-1}$ | 0.9 | $4.11 \times 10^{-1}$ | 1.7 | $1.94 \times 10^{-1}$ | 5'-UTR polymorphism | Moderate | | |
| 3 | 9671718 | 84/G | 42.4 ± 1.5 | 67/C | 39.5 ± 1.4 | $1.44 \times 10^{-1}$ | **3.3** | $7.29 \times 10^{-2}$ | 0.4 | $5.09 \times 10^{-1}$ | 5'-UTR polymorphism | Moderate | | |
| 3 | 9671733 | 80/T | 42.7 ± 1.4 | 71/G | 39.3 ± 1.4 | $8.78 \times 10^{-2}$ | **4.1** | **$4.56 \times 10^{-2}$** | 0.1 | $8.17 \times 10^{-1}$ | 5'-UTR polymorphism | Moderate | | |
| 3 | 9671744 | 86/A | 42.6 ± 1.4 | 65/T | 39.1 ± 1.5 | $8.56 \times 10^{-2}$ | **4.2** | **$4.25 \times 10^{-2}$** | 0.1 | $7.04 \times 10^{-1}$ | 5'-UTR polymorphism | Moderate | | |
| 3 | 9671747 | 81/C | 42.9 ± 1.5 | 70/G | 39 ± 1.4 | $5.25 \times 10^{-2}$ | **5.1** | **$2.61 \times 10^{-2}$** | 0.8 | $3.75 \times 10^{-1}$ | 5'-UTR polymorphism | Moderate | | |
| 3 | 9671769 | 6/G | 43 ± 5.5 | 145/C | 41 ± 1.1 | $7.01 \times 10^{-1}$ | 0.2 | $6.56 \times 10^{-1}$ | 0.3 | $5.81 \times 10^{-1}$ | 5'-UTR polymorphism | Moderate | | |
| 3 | 9671782 | 90/A | 42.4 ± 1.4 | 61/G | 39.2 ± 1.4 | $1.12 \times 10^{-1}$ | **3.8** | $5.32 \times 10^{-2}$ | 0.0 | $8.42 \times 10^{-1}$ | 5'-UTR polymorphism | Moderate | | |
| 3 | 9671879 | 5/T | 44.2 ± 6.5 | 145/G | 41.1 ± 1.1 | $5.77 \times 10^{-1}$ | 0.7 | $4.78 \times 10^{-1}$ | 1.7 | $1.86 \times 10^{-1}$ | 5'-UTR polymorphism | Moderate | | |
| 3 | 9671888 | 76/A | 42.6 ± 1.5 | 75/G | 39.6 ± 1.4 | $1.29 \times 10^{-1}$ | **3.8** | $5.37 \times 10^{-2}$ | 0.1 | $6.99 \times 10^{-1}$ | 5'-UTR polymorphism | Moderate | | |
| 3 | 9671913 | 91/T | 42.7 ± 1.4 | 60/A | 38.7 ± 1.4 | **$4.62 \times 10^{-2}$** | **5.5** | **$1.99 \times 10^{-2}$** | 0.2 | $6.63 \times 10^{-1}$ | 5'-UTR polymorphism | Moderate | | |
| 3 | 9671916 | 6/T | 43 ± 5.5 | 145/G | 41 ± 1.1 | $7.01 \times 10^{-1}$ | 0.2 | $6.56 \times 10^{-1}$ | 0.3 | $5.81 \times 10^{-1}$ | 5'-UTR polymorphism | Moderate | | |
| 3 | 9671944 | 4/A | 41.3 ± 3.3 | 147/T | 41.1 ± 1.1 | $9.69 \times 10^{-1}$ | 0.0 | $9.42 \times 10^{-1}$ | 0.4 | $5.44 \times 10^{-1}$ | 5'-UTR polymorphism | Moderate | | |
| 3 | 9671964 | 82/G | 42.3 ± 1.5 | 69/T | 39.7 ± 1.4 | $1.98 \times 10^{-1}$ | **3.3** | $7.14 \times 10^{-2}$ | 0.5 | $4.74 \times 10^{-1}$ | Amino acid substitution (Asp4Glu) | Moderate | | |
| 3 | 9672848 | 9/G | 43.6 ± 3.7 | 142/T | 40.9 ± 1.1 | $5.33 \times 10^{-1}$ | 0.7 | $4.11 \times 10^{-1}$ | 0.4 | $5.06 \times 10^{-1}$ | Amino acid substitution (Asp177Glu) | Moderate | | |
| 3 | 9670551 | 85/A | 42.9 ± 1.4 | 66/C | 38.7 ± 1.4 | **$3.37 \times 10^{-2}$** | **6.3** | **$1.31 \times 10^{-2}$** | 1.3 | $2.50 \times 10^{-1}$ | Promoter polymorphism | Moderate | AT3G26430 | *GGL20* |
| 3 | 9670560 | 84/G | 42.7 ± 1.4 | 67/A | 39.1 ± 1.4 | $7.69 \times 10^{-2}$ | **4.5** | **$3.49 \times 10^{-2}$** | 0.1 | $6.99 \times 10^{-1}$ | Promoter polymorphism | Moderate | | |
| 3 | 9670788 | 82/T | 43.1 ± 1.5 | 69/C | 38.7 ± 1.3 | **$2.64 \times 10^{-2}$** | **6.8** | **$9.86 \times 10^{-3}$** | 0.9 | $3.49 \times 10^{-1}$ | Promoter polymorphism | Moderate | | |
| 3 | 9670811 | 85/C | 42.9 ± 1.4 | 66/A | 38.8 ± 1.4 | **$4.19 \times 10^{-2}$** | **5.9** | **$1.67 \times 10^{-2}$** | 0.2 | $6.89 \times 10^{-1}$ | Promoter polymorphism | Moderate | | |
| 3 | 9670816 | 77/C | 42.9 ± 1.5 | 74/A | 39.2 ± 1.4 | $6.38 \times 10^{-2}$ | **4.7** | **$3.18 \times 10^{-2}$** | 0.2 | $6.94 \times 10^{-1}$ | Promoter polymorphism | Moderate | | |
| 3 | 9670881 | 83/A | 42.6 ± 1.5 | 68/C | 39.2 ± 1.3 | $8.70 \times 10^{-2}$ | **4.0** | **$4.78 \times 10^{-2}$** | 0.1 | $7.40 \times 10^{-1}$ | Promoter polymorphism | Moderate | | |
| 3 | 9671094 | 93/A | 42.7 ± 1.4 | 58/G | 38.6 ± 1.4 | **$4.49 \times 10^{-2}$** | **6.7** | **$1.07 \times 10^{-2}$** | 0.9 | $3.51 \times 10^{-1}$ | Promoter polymorphism | Moderate | | |
| 3 | 9671106 | 91/A | 42.7 ± 1.4 | 60/T | 38.6 ± 1.4 | **$4.08 \times 10^{-2}$** | **7.0** | **$9.26 \times 10^{-3}$** | 0.5 | $4.70 \times 10^{-1}$ | Promoter polymorphism | Moderate | | |
| 3 | 9671262 | 90/T | 42.5 ± 1.4 | 61/C | 39.1 ± 1.4 | $9.17 \times 10^{-2}$ | **4.7** | **$3.19 \times 10^{-2}$** | 0.1 | $7.04 \times 10^{-1}$ | Promoter polymorphism | Moderate | | |
| 3 | 9671288 | 72/T | 42.7 ± 1.6 | 79/C | 39.7 ± 1.3 | $1.32 \times 10^{-1}$ | **3.9** | **$4.97 \times 10^{-2}$** | 0.0 | $8.58 \times 10^{-1}$ | Promoter polymorphism | Moderate | | |
| 3 | 9671302 | 88/C | 42.4 ± 1.4 | 63/G | 39.3 ± 1.4 | $1.21 \times 10^{-1}$ | **4.2** | **$4.11 \times 10^{-2}$** | 0.0 | $9.33 \times 10^{-1}$ | Promoter polymorphism | Moderate | | |
| 3 | 9671412 | 134/C | 41.4 ± 1.1 | 17/T | 38.7 ± 2.9 | $3.95 \times 10^{-1}$ | 0.9 | $3.51 \times 10^{-1}$ | 0.3 | $5.76 \times 10^{-1}$ | Promoter polymorphism | Moderate | | |
| 3 | 9671428 | 84/A | 42.1 ± 1.4 | 67/T | 39.9 ± 1.4 | $2.79 \times 10^{-1}$ | 2.3 | $1.28 \times 10^{-1}$ | 1.6 | $2.08 \times 10^{-1}$ | Promoter polymorphism | Moderate | | |
| 3 | 9671501 | 37/CT | 45.6 ± 2.1 | 114/C | 39.6 ± 1.2 | **$9.34 \times 10^{-3}$** | **11.1** | **$1.12 \times 10^{-3}$** | **4.5** | **$3.66 \times 10^{-2}$** | Promoter polymorphism | Moderate | | |
| 3 | 9671524 | 6/T | 43 ± 5.5 | 145/C | 41 ± 1.1 | $7.01 \times 10^{-1}$ | 0.2 | $6.56 \times 10^{-1}$ | 0.3 | $5.81 \times 10^{-1}$ | 5'-UTR polymorphism | Moderate | | |
| 3 | 9671576 | 89/G | 42.6 ± 1.4 | 62/T | 38.9 ± 1.4 | $6.10 \times 10^{-2}$ | **5.6** | **$1.91 \times 10^{-2}$** | 0.0 | $9.77 \times 10^{-1}$ | 5'-UTR polymorphism | Moderate | | |
| 3 | 9671582 | 86/C | 41.9 ± 1.4 | 64/T | 40 ± 1.5 | $3.70 \times 10^{-1}$ | 0.9 | $4.11 \times 10^{-1}$ | 1.7 | $1.94 \times 10^{-1}$ | 5'-UTR polymorphism | Moderate | | |
| 3 | 9671718 | 84/G | 42.4 ± 1.5 | 67/C | 39.5 ± 1.4 | $1.44 \times 10^{-1}$ | **3.3** | $7.29 \times 10^{-2}$ | 0.4 | $5.09 \times 10^{-1}$ | 5'-UTR polymorphism | Moderate | | |
| 3 | 9671733 | 80/T | 42.7 ± 1.4 | 71/G | 39.3 ± 1.4 | $8.78 \times 10^{-2}$ | **4.1** | **$4.56 \times 10^{-2}$** | 0.1 | $8.17 \times 10^{-1}$ | 5'-UTR polymorphism | Moderate | | |
| 3 | 9671744 | 86/A | 42.6 ± 1.4 | 65/T | 39.1 ± 1.5 | $8.56 \times 10^{-2}$ | **4.2** | **$4.25 \times 10^{-2}$** | 0.1 | $7.04 \times 10^{-1}$ | 5'-UTR polymorphism | Moderate | | |
| 3 | 9671747 | 81/C | 42.9 ± 1.5 | 70/G | 39 ± 1.4 | $5.25 \times 10^{-2}$ | **5.1** | **$2.61 \times 10^{-2}$** | 0.8 | $3.75 \times 10^{-1}$ | 5'-UTR polymorphism | Moderate | | |
| 3 | 9671769 | 6/G | 43 ± 5.5 | 145/C | 41 ± 1.1 | $7.01 \times 10^{-1}$ | 0.2 | $6.56 \times 10^{-1}$ | 0.3 | $5.81 \times 10^{-1}$ | 5'-UTR polymorphism | Moderate | | |
| 3 | 9671782 | 90/A | 42.4 ± 1.4 | 61/G | 39.2 ± 1.4 | $1.12 \times 10^{-1}$ | **3.8** | $5.32 \times 10^{-2}$ | 0.0 | $8.42 \times 10^{-1}$ | 5'-UTR polymorphism | Moderate | | |
| 3 | 9671879 | 5/T | 44.2 ± 6.5 | 145/G | 41.1 ± 1.1 | $5.77 \times 10^{-1}$ | 0.7 | $4.78 \times 10^{-1}$ | 1.7 | $1.86 \times 10^{-1}$ | 5'-UTR polymorphism | Moderate | | |
| 3 | 9671888 | 76/A | 42.6 ± 1.5 | 75/G | 39.6 ± 1.4 | $1.29 \times 10^{-1}$ | **3.8** | $5.37 \times 10^{-2}$ | 0.1 | $6.99 \times 10^{-1}$ | 5'-UTR polymorphism | Moderate | | |
| 3 | 9671913 | 91/T | 42.7 ± 1.4 | 60/A | 38.7 ± 1.4 | **$4.62 \times 10^{-2}$** | **5.5** | **$1.99 \times 10^{-2}$** | 0.2 | $6.63 \times 10^{-1}$ | 5'-UTR polymorphism | Moderate | | |
| 3 | 9671916 | 6/T | 43 ± 5.5 | 145/G | 41 ± 1.1 | $7.01 \times 10^{-1}$ | 0.2 | $6.56 \times 10^{-1}$ | 0.3 | $5.81 \times 10^{-1}$ | 5'-UTR polymorphism | Moderate | | |
| 3 | 9671944 | 4/A | 41.3 ± 3.3 | 147/T | 41.1 ± 1.1 | $9.69 \times 10^{-1}$ | 0.0 | $9.42 \times 10^{-1}$ | 0.4 | $5.44 \times 10^{-1}$ | 5'-UTR polymorphism | Moderate | | |

| | | | | | | | | | | | | |
|---|---|---|---|---|---|---|---|---|---|---|---|---|
| 3 | 9671964 | 82/G | 42.3 ± 1.5 | 69/T | 39.7 ± 1.4 | $1.98 \times 10^{-1}$ | **3.3** | $7.14 \times 10^{-2}$ | 0.5 | $4.74 \times 10^{-1}$ | 5'-UTR polymorphism | Moderate |
| 3 | 9671967 | 7/A | 42.5 ± 4.6 | 144/G | 41 ± 1.1 | $7.61 \times 10^{-1}$ | 0.1 | $7.72 \times 10^{-1}$ | 0.5 | $4.89 \times 10^{-1}$ | 5'-UTR polymorphism | Moderate |
| 3 | 9671970 | 8/A | 43.5 ± 6.9 | 143/G | 41 ± 1 | $5.74 \times 10^{-1}$ | 0.5 | $4.75 \times 10^{-1}$ | 0.0 | $8.37 \times 10^{-1}$ | 5'-UTR polymorphism | Moderate |
| 3 | 9672070 | 90/C | 42.4 ± 1.4 | 60/T | 39.1 ± 1.5 | $9.89 \times 10^{-2}$ | 2.1 | $1.27 \times 10^{-1}$ | 0.0 | $9.56 \times 10^{-1}$ | 5'-UTR polymorphism | Moderate |
| 3 | 9672081 | 6/G | 43 ± 5.5 | 145/C | 41 ± 1.1 | $7.01 \times 10^{-1}$ | 0.2 | $6.56 \times 10^{-1}$ | 0.3 | $5.81 \times 10^{-1}$ | 5'-UTR polymorphism | Moderate |
| 3 | 9672140 | 94/T | 42.8 ± 1.4 | 57/C | 38.3 ± 1.4 | **$3.06 \times 10^{-2}$** | **7.7** | **$6.29 \times 10^{-3}$** | 0.6 | $4.47 \times 10^{-1}$ | 5'-UTR polymorphism | Moderate |
| 3 | 9672142 | 6/T | 43 ± 5.5 | 145/C | 41 ± 1.1 | $7.01 \times 10^{-1}$ | 0.2 | $6.56 \times 10^{-1}$ | 0.3 | $5.81 \times 10^{-1}$ | 5'-UTR polymorphism | Moderate |
| 3 | 9672168 | 82/T | 42.8 ± 1.4 | 69/G | 39.1 ± 1.4 | $6.02 \times 10^{-2}$ | **4.5** | **$3.49 \times 10^{-2}$** | 1.2 | $2.74 \times 10^{-1}$ | 5'-UTR polymorphism | Moderate |
| 3 | 9672189 | 60/C | 41.9 ± 1.6 | 91/G | 40.5 ± 1.3 | $5.00 \times 10^{-1}$ | 0.6 | $4.58 \times 10^{-1}$ | 1.0 | $3.23 \times 10^{-1}$ | 5'-UTR polymorphism | Moderate |
| 3 | 9672190 | 64/G | 41.7 ± 1.6 | 87/C | 40.6 ± 1.3 | $5.83 \times 10^{-1}$ | 0.3 | $5.73 \times 10^{-1}$ | 2.2 | $1.41 \times 10^{-1}$ | 5'-UTR polymorphism | Moderate |
| 3 | 9672194 | 74/T | 42.4 ± 1.5 | 77/C | 39.9 ± 1.3 | $2.07 \times 10^{-1}$ | 2.0 | $1.57 \times 10^{-1}$ | 0.4 | $5.48 \times 10^{-1}$ | 5'-UTR polymorphism | Moderate |
| 3 | 9672262 | 39/G | 42.9 ± 2.1 | 112/C | 40.5 ± 1.2 | $2.85 \times 10^{-1}$ | 2.6 | $1.08 \times 10^{-1}$ | 0.7 | $4.17 \times 10^{-1}$ | 5'-UTR polymorphism | Moderate |
| 3 | 9672265 | 78/G | 42.1 ± 1.5 | 73/A | 40 ± 1.3 | $2.81 \times 10^{-1}$ | 2.3 | $1.29 \times 10^{-1}$ | 0.0 | $9.12 \times 10^{-1}$ | 5'-UTR polymorphism | Moderate |
| 3 | 9672276 | 86/G | 42.6 ± 1.5 | 65/A | 39 ± 1.3 | $7.28 \times 10^{-2}$ | **5.1** | $2.50 \times 10^{-2}$ | 0.4 | $5.34 \times 10^{-1}$ | 5'-UTR polymorphism | Moderate |
| 3 | 9672348 | 80/G | 42.7 ± 1.5 | 71/T | 39.2 ± 1.3 | $7.58 \times 10^{-2}$ | **4.4** | **$3.82 \times 10^{-2}$** | 0.0 | $8.82 \times 10^{-1}$ | 5'-UTR polymorphism | Moderate |
| 3 | 9672353 | 80/C | 43.1 ± 1.4 | 71/T | 38.8 ± 1.4 | **$2.73 \times 10^{-2}$** | **6.9** | **$9.36 \times 10^{-3}$** | 1.8 | $1.86 \times 10^{-1}$ | 5'-UTR polymorphism | Moderate |
| 3 | 9672365 | 63/CA | 43.9 ± 1.6 | 88/C | 39.1 ± 1.3 | $1.77 \times 10^{-2}$ | **9.1** | **$3.04 \times 10^{-3}$** | 1.8 | $1.81 \times 10^{-1}$ | 5'-UTR polymorphism | Moderate |
| 3 | 9672400 | 83/G | 42.9 ± 1.4 | 68/A | 38.8 ± 1.5 | **$3.91 \times 10^{-2}$** | **6.3** | **$1.29 \times 10^{-2}$** | 1.5 | $2.21 \times 10^{-1}$ | 5'-UTR polymorphism | Moderate |
| 3 | 9672470 | 6/G | 43 ± 5.5 | 145/A | 41 ± 1.1 | $7.01 \times 10^{-1}$ | 0.2 | $6.56 \times 10^{-1}$ | 0.3 | $5.81 \times 10^{-1}$ | 5'-UTR polymorphism | Moderate |
| 3 | 9672572 | 6/G | 43 ± 5.5 | 145/T | 41 ± 1.1 | $7.01 \times 10^{-1}$ | 0.2 | $6.56 \times 10^{-1}$ | 0.3 | $5.81 \times 10^{-1}$ | 5'-UTR polymorphism | Moderate |
| 3 | 9672611 | 91/C | 42.7 ± 1.4 | 60/T | 38.7 ± 1.5 | $5.10 \times 10^{-2}$ | **5.8** | **$1.75 \times 10^{-2}$** | 0.1 | $8.09 \times 10^{-1}$ | 5'-UTR polymorphism | Moderate |
| 3 | 9672641 | 6/T | 43 ± 5.5 | 145/C | 41 ± 1.1 | $7.01 \times 10^{-1}$ | 0.2 | $6.56 \times 10^{-1}$ | 0.3 | $5.81 \times 10^{-1}$ | 5'-UTR polymorphism | Moderate |
| 3 | 9672659 | 144/G | 41.5 ± 1.1 | 7/T | 33.3 ± 4.4 | $8.19 \times 10^{-2}$ | 2.6 | $1.12 \times 10^{-1}$ | **4.3** | **$4.08 \times 10^{-2}$** | 5'-UTR polymorphism | Moderate |
| 3 | 9672683 | 6/G | 43 ± 5.5 | 145/A | 41 ± 1.1 | $7.01 \times 10^{-1}$ | 0.2 | $6.56 \times 10^{-1}$ | 0.3 | $5.81 \times 10^{-1}$ | 5'-UTR polymorphism | Moderate |
| 3 | 9672848 | 9/G | 43.6 ± 3.7 | 142/T | 40.9 ± 1.1 | $5.33 \times 10^{-1}$ | 0.7 | $4.11 \times 10^{-1}$ | 0.4 | $5.06 \times 10^{-1}$ | 5'-UTR polymorphism | Moderate |
| 3 | 9673112 | 6/A | 43 ± 5.5 | 145/C | 41 ± 1.1 | $7.01 \times 10^{-1}$ | 0.2 | $6.56 \times 10^{-1}$ | 0.3 | $5.81 \times 10^{-1}$ | 5'-UTR polymorphism | Moderate |
| 3 | 9673155 | 83/C | 42.9 ± 1.5 | 68/T | 38.9 ± 1.3 | **$4.28 \times 10^{-2}$** | **5.7** | **$1.86 \times 10^{-2}$** | 0.1 | $7.01 \times 10^{-1}$ | 5'-UTR polymorphism | Moderate |
| 3 | 9673156 | 10/C | 42.6 ± 3.4 | 141/T | 41 ± 1.1 | $6.92 \times 10^{-1}$ | 0.3 | $5.78 \times 10^{-1}$ | 0.5 | $4.81 \times 10^{-1}$ | 5'-UTR polymorphism | Moderate |
| 3 | 9673225 | 6/C | 43 ± 5.5 | 145/T | 41 ± 1.1 | $7.01 \times 10^{-1}$ | 0.2 | $6.56 \times 10^{-1}$ | 0.3 | $5.81 \times 10^{-1}$ | 5'-UTR polymorphism | Moderate |
| 3 | 9673495 | 135/G | 41.3 ± 1.1 | 16/A | 38.9 ± 3.1 | $4.51 \times 10^{-1}$ | 0.7 | $3.93 \times 10^{-1}$ | 0.2 | $6.27 \times 10^{-1}$ | 5'-UTR polymorphism | Moderate |
| 3 | 9673597 | 5/T | 46.1 ± 5.4 | 146/G | 40.9 ± 1.1 | $3.51 \times 10^{-1}$ | 1.0 | $3.20 \times 10^{-1}$ | 1.6 | $2.08 \times 10^{-1}$ | 5'-UTR polymorphism | Moderate |
| 3 | 9673618 | 81/T | 43.3 ± 1.5 | 70/A | 38.5 ± 1.3 | **$1.63 \times 10^{-2}$** | **7.4** | **$7.43 \times 10^{-3}$** | 2.2 | $1.41 \times 10^{-1}$ | 5'-UTR polymorphism | Moderate |
| 3 | 9673743 | 97/AG | 41.8 ± 1.3 | 54/A | 39.8 ± 1.8 | $3.43 \times 10^{-1}$ | 0.3 | $5.95 \times 10^{-1}$ | 0.0 | $8.38 \times 10^{-1}$ | 5'-UTR polymorphism | Moderate |
| 3 | 9673768 | 6/A | 43 ± 5.5 | 145/G | 41 ± 1.1 | $7.01 \times 10^{-1}$ | 0.2 | $6.56 \times 10^{-1}$ | 0.3 | $5.81 \times 10^{-1}$ | 5'-UTR polymorphism | Moderate |
| 3 | 9673807 | 6/C | 43 ± 5.5 | 145/T | 41 ± 1.1 | $7.01 \times 10^{-1}$ | 0.2 | $6.56 \times 10^{-1}$ | 0.3 | $5.81 \times 10^{-1}$ | 5'-UTR polymorphism | Moderate |
| 3 | 9673864 | 145/G | 41.3 ± 1.1 | 6/T | 36.3 ± 6.2 | $3.30 \times 10^{-1}$ | 1.4 | $2.32 \times 10^{-1}$ | **3.1** | $7.86 \times 10^{-2}$ | 5'-UTR polymorphism | Moderate |
| 3 | 9673879 | 76/G | 42.9 ± 1.6 | 75/T | 39.2 ± 1.3 | $6.27 \times 10^{-2}$ | **4.9** | **$2.83 \times 10^{-2}$** | 0.1 | $7.16 \times 10^{-1}$ | 5'-UTR polymorphism | Moderate |
| 3 | 9673903 | 7/G | 46.6 ± 6.1 | 144/GACCTCT | 40.8 ± 1 | $2.21 \times 10^{-1}$ | 1.5 | $2.18 \times 10^{-1}$ | 0.9 | $3.49 \times 10^{-1}$ | 5'-UTR polymorphism | Moderate |
| 3 | 9673937 | 54/G | 43.3 ± 1.7 | 97/A | 39.8 ± 1.3 | $9.14 \times 10^{-2}$ | 2.6 | $1.08 \times 10^{-1}$ | 1.9 | $1.73 \times 10^{-1}$ | 5'-UTR polymorphism | Moderate |
| 3 | 9673939 | 6/G | 43 ± 5.5 | 145/C | 41 ± 1.1 | $7.01 \times 10^{-1}$ | 0.2 | $6.56 \times 10^{-1}$ | 0.3 | $5.81 \times 10^{-1}$ | 5'-UTR polymorphism | Moderate |
| 3 | 9673954 | 114/A | 41.4 ± 1.1 | 37/C | 40.3 ± 2.3 | $6.38 \times 10^{-1}$ | 0.4 | $5.30 \times 10^{-1}$ | **4.7** | **$3.18 \times 10^{-2}$** | 5'-UTR polymorphism | Moderate |
| 3 | 9673960 | 112/G | 41.2 ± 1.1 | 39/A | 40.7 ± 2.3 | $8.08 \times 10^{-1}$ | 0.2 | $6.86 \times 10^{-1}$ | **3.4** | $6.80 \times 10^{-2}$ | 5'-UTR polymorphism | Moderate |

| | | | | | | | | | | | | | |
|---|---|---|---|---|---|---|---|---|---|---|---|---|---|
| 3 | 9673972 | 37/C | 43.4 ± 2.2 | 114/A | 40.3 ± 1.2 | $1.77 \times 10^{-1}$ | 1.0 | $3.29 \times 10^{-1}$ | 0.3 | $5.66 \times 10^{-1}$ | | 5'-UTR polymorphism | Moderate |
| 3 | 9673998 | 20/C | 45 ± 3.5 | 131/T | 40.5 ± 1.1 | $1.24 \times 10^{-1}$ | 2.0 | $1.56 \times 10^{-1}$ | 0.4 | $5.35 \times 10^{-1}$ | | 5'-UTR polymorphism | Moderate |
| 3 | 9674007 | 13/C | 43.7 ± 5 | 138/T | 40.8 ± 1 | $4.25 \times 10^{-1}$ | 0.1 | $7.38 \times 10^{-1}$ | 0.0 | $8.28 \times 10^{-1}$ | | 5'-UTR polymorphism | Moderate |
| 3 | 9674023 | 144/T | 41.2 ± 1 | 7/C | 39.1 ± 6.9 | $6.59 \times 10^{-1}$ | 1.5 | $2.29 \times 10^{-1}$ | **3.4** | $6.62 \times 10^{-2}$ | | 5'-UTR polymorphism | Moderate |
| 3 | 9674036 | 142/T | 41.4 ± 1 | 9/G | 36 ± 5.8 | $2.01 \times 10^{-1}$ | 2.2 | $1.36 \times 10^{-1}$ | **4.4** | **$3.68 \times 10^{-2}$** | | 5'-UTR polymorphism | Moderate |
| 3 | 9674084 | 8/G | 42 ± 4.2 | 142/C | 41.1 ± 1.1 | $8.43 \times 10^{-1}$ | 0.3 | $7.45 \times 10^{-1}$ | 0.2 | $7.92 \times 10^{-1}$ | | 5'-UTR polymorphism | Moderate |
| 3 | 9674100 | 6/A | 43 ± 5.5 | 145/G | 41 ± 1.1 | $7.01 \times 10^{-1}$ | 0.2 | $6.56 \times 10^{-1}$ | 0.3 | $5.81 \times 10^{-1}$ | | 5'-UTR polymorphism | Moderate |
| 3 | 9674111 | 15/G | 44.2 ± 2.5 | 136/A | 40.7 ± 1.1 | $2.94 \times 10^{-1}$ | 1.6 | $2.13 \times 10^{-1}$ | 0.8 | $3.87 \times 10^{-1}$ | | 5'-UTR polymorphism | Moderate |
| 3 | 9674172 | 9/T | 48.1 ± 3.3 | 142/C | 40.6 ± 1.1 | $7.34 \times 10^{-2}$ | **3.8** | $5.47 \times 10^{-2}$ | 2.2 | $1.41 \times 10^{-1}$ | | 5'-UTR polymorphism | Moderate |
| 3 | 9674182 | 10/A | 47.8 ± 3 | 141/G | 40.6 ± 1.1 | $7.08 \times 10^{-2}$ | **3.8** | $5.31 \times 10^{-2}$ | 2.5 | $1.13 \times 10^{-1}$ | | 5'-UTR polymorphism | Moderate |
| 3 | 9674194 | 12/G | 44.7 ± 3.1 | 139/T | 40.8 ± 1.1 | $2.83 \times 10^{-1}$ | 1.3 | $2.61 \times 10^{-1}$ | 0.4 | $5.15 \times 10^{-1}$ | | 5'-UTR polymorphism | Moderate |
| 3 | 9674225 | 18/C | 41.7 ± 3 | 133/T | 41 ± 1.1 | $8.28 \times 10^{-1}$ | 0.2 | $6.28 \times 10^{-1}$ | 0.0 | $9.50 \times 10^{-1}$ | | 5'-UTR polymorphism | Moderate |
| 3 | 9674236 | 135/A | 41.2 ± 1.1 | 16/G | 40.2 ± 2 | $7.65 \times 10^{-1}$ | 0.0 | $9.35 \times 10^{-1}$ | 0.8 | $3.72 \times 10^{-1}$ | | 5'-UTR polymorphism | Moderate |
| 3 | 9674240 | 144/CTT | 41.2 ± 1.1 | 7/C | 38.8 ± 3.7 | $6.18 \times 10^{-1}$ | 0.3 | $6.13 \times 10^{-1}$ | 2.2 | $1.39 \times 10^{-1}$ | | 5'-UTR polymorphism | Moderate |
| 3 | 9674246 | 17/G | 42.5 ± 2.4 | 128/T | 40.8 ± 1.2 | $5.94 \times 10^{-1}$ | 0.4 | $6.90 \times 10^{-1}$ | 0.2 | $8.10 \times 10^{-1}$ | | 5'-UTR polymorphism | Moderate |
| 3 | 9674267 | 36/C | 41.6 ± 1.9 | 115/T | 40.9 ± 1.2 | $7.66 \times 10^{-1}$ | 0.4 | $5.54 \times 10^{-1}$ | 0.0 | $9.63 \times 10^{-1}$ | | 5'-UTR polymorphism | Moderate |
| 3 | 9674311 | 74/A | 41.9 ± 1.4 | 77/T | 40.3 ± 1.5 | $4.40 \times 10^{-1}$ | 1.6 | $2.10 \times 10^{-1}$ | 0.1 | $7.73 \times 10^{-1}$ | | 5'-UTR polymorphism | Moderate |
| 3 | 9674324 | 71/A | 43.1 ± 1.5 | 75/T | 39 ± 1.4 | **$4.20 \times 10^{-2}$** | **3.2** | **$4.48 \times 10^{-2}$** | 0.5 | $6.36 \times 10^{-1}$ | | 5'-UTR polymorphism | Moderate |
| 3 | 9674330 | 5/G | 46.1 ± 5.4 | 146/A | 40.9 ± 1.1 | $3.51 \times 10^{-1}$ | 1.0 | $3.20 \times 10^{-1}$ | 1.6 | $2.08 \times 10^{-1}$ | | 5'-UTR polymorphism | Moderate |
| 3 | 9674332 | 127/T | 41.6 ± 1.1 | 24/C | 38.3 ± 2.7 | $2.21 \times 10^{-1}$ | 1.0 | $3.19 \times 10^{-1}$ | 1.6 | $2.08 \times 10^{-1}$ | | 5'-UTR polymorphism | Moderate |
| 3 | 9674342 | 137/C | 41.3 ± 1.1 | 14/G | 38.6 ± 2.9 | $4.20 \times 10^{-1}$ | 1.0 | $3.09 \times 10^{-1}$ | 0.3 | $6.05 \times 10^{-1}$ | | 5'-UTR polymorphism | Moderate |
| 3 | 9674434 | 113/T | 41.1 ± 1.2 | 38/A | 41 ± 1.9 | $9.69 \times 10^{-1}$ | 0.0 | $8.35 \times 10^{-1}$ | 0.1 | $7.12 \times 10^{-1}$ | | Amino acid substitution (Leu6Met) | Moderate |
| 3 | 9674439 | 6/T | 42.6 ± 2.4 | 145/G | 41 ± 1.1 | $7.59 \times 10^{-1}$ | 0.2 | $6.76 \times 10^{-1}$ | 0.6 | $4.37 \times 10^{-1}$ | | Amino acid substitution (Leu7Phe) | Moderate |
| 3 | 9674440 | 55/A | 41.8 ± 1.6 | 96/G | 40.7 ± 1.3 | $6.03 \times 10^{-1}$ | 1.2 | $2.77 \times 10^{-1}$ | 0.0 | $8.76 \times 10^{-1}$ | | Amino acid substitution (Val8Ile) | Moderate |
| 3 | 9674483 | 5/T | 41.4 ± 5.2 | 146/C | 41.1 ± 1.1 | $9.54 \times 10^{-1}$ | 0.1 | $7.07 \times 10^{-1}$ | 0.0 | $9.56 \times 10^{-1}$ | | Amino acid substitution (Ala22Val) | Moderate |
| 3 | 9674551 | 3/A | 42.8 ± 4.2 | 148/C | 41.1 ± 1.1 | $8.07 \times 10^{-1}$ | 0.1 | $7.57 \times 10^{-1}$ | 0.2 | $6.95 \times 10^{-1}$ | | Amino acid substitution (Leu45Ile) | Moderate |
| 3 | 9674560 | 35/T | 41.9 ± 2.1 | 116/G | 40.8 ± 1.2 | $6.54 \times 10^{-1}$ | 0.4 | $5.37 \times 10^{-1}$ | 0.0 | $9.09 \times 10^{-1}$ | | Amino acid substitution (Ser48Ala) | Moderate |
| 3 | 9674783 | 144/A | 41.2 ± 1.1 | 7/T | 39.4 ± 3.4 | $7.01 \times 10^{-1}$ | 0.2 | $6.90 \times 10^{-1}$ | 0.0 | $9.21 \times 10^{-1}$ | | Amino acid substitution (Asn97Ile) | Moderate |
| 3 | 9674791 | 147/C | 41.2 ± 1.1 | 4/A | 36.2 ± 4.9 | $4.13 \times 10^{-1}$ | 0.8 | $3.87 \times 10^{-1}$ | 0.0 | $9.11 \times 10^{-1}$ | | Amino acid substitution (His100Asn) | Moderate |
| 3 | 9674928 | 1/T | #VALUE! | 150/TC | 41.1 ± 1 | | 0.1 | $7.52 \times 10^{-1}$ | 0.0 | $8.65 \times 10^{-1}$ | | **frameshift_variant** | **High** |
| 3 | 9675121 | 127/T | 41.2 ± 1.1 | 24/A | 40.7 ± 2.4 | $8.52 \times 10^{-1}$ | 0.0 | $9.98 \times 10^{-1}$ | 0.2 | $6.22 \times 10^{-1}$ | | Amino acid substitution (Ser186Thr) | Moderate |
| 3 | 9675127 | 149/CA | 41.2 ± 1.1 | 2/C | 34.7 ± 2.3 | $4.54 \times 10^{-1}$ | 0.7 | $3.98 \times 10^{-1}$ | 0.2 | $6.69 \times 10^{-1}$ | | **frameshift_variant** | **High** |
| 3 | 9675148 | 69/A | 43.6 ± 1.6 | 82/G | 39 ± 1.3 | **$2.00 \times 10^{-2}$** | **6.9** | **$9.64 \times 10^{-3}$** | 1.6 | $2.13 \times 10^{-1}$ | | Amino acid substitution (Asp195Asn) | Moderate |
| 3 | 9675155 | 37/A | 41.3 ± 2.6 | 114/T | 41 ± 1.1 | $9.03 \times 10^{-1}$ | 0.8 | $3.86 \times 10^{-1}$ | 0.0 | $9.97 \times 10^{-1}$ | | Amino acid substitution (His197Leu) | Moderate |
| 3 | 9675163 | 64/T | 42.9 ± 1.7 | 87/C | 39.8 ± 1.3 | $1.26 \times 10^{-1}$ | **3.2** | $7.59 \times 10^{-2}$ | 0.2 | $6.66 \times 10^{-1}$ | | Amino acid substitution (Leu200Phe) | Moderate |
| 3 | 9675173 | 64/C | 43 ± 1.6 | 87/T | 39.7 ± 1.3 | $1.03 \times 10^{-1}$ | **4.3** | **$4.05 \times 10^{-2}$** | 1.3 | $2.58 \times 10^{-1}$ | | Amino acid substitution (Val203Ala) | Moderate |
| 3 | 9675178 | 147/C | 41.1 ± 1.1 | 4/T | 39.6 ± 5.8 | $8.09 \times 10^{-1}$ | 0.7 | $4.17 \times 10^{-1}$ | 0.0 | $8.84 \times 10^{-1}$ | | Amino acid substitution (Arg205Cys) | Moderate |
| 3 | 9675361 | 57/T | 44.1 ± 1.8 | 94/C | 39.3 ± 1.2 | **$1.87 \times 10^{-2}$** | **10.7** | **$1.34 \times 10^{-3}$** | 2.0 | $1.58 \times 10^{-1}$ | ✓ | Amino acid substitution (Pro236Leu) | Moderate |
| 3 | 9675381 | 51/T | 43.6 ± 1.8 | 100/C | 39.8 ± 1.2 | $6.50 \times 10^{-2}$ | **4.5** | **$3.47 \times 10^{-2}$** | 0.1 | $7.43 \times 10^{-1}$ | | Amino acid substitution (His243Tyr) | Moderate |
| 3 | 9675411 | 146/G | 41.1 ± 1 | 5/T | 39.4 ± 7.5 | $7.54 \times 10^{-1}$ | 1.1 | $3.04 \times 10^{-1}$ | 1.1 | $2.95 \times 10^{-1}$ | | Amino acid substitution (Ala253Ser) | Moderate |
| 3 | 9675436 | 81/G | 43 ± 1.5 | 70/A | 38.9 ± 1.3 | **$4.26 \times 10^{-2}$** | **5.8** | **$1.69 \times 10^{-2}$** | 0.2 | $6.40 \times 10^{-1}$ | | Amino acid substitution (Lys261Arg) | Moderate |
| 3 | 9675502 | 146/C | 41.2 ± 1.1 | 5/A | 38.5 ± 3.8 | $6.33 \times 10^{-1}$ | 0.2 | $6.53 \times 10^{-1}$ | 0.0 | $8.90 \times 10^{-1}$ | | Amino acid substitution (Ser283Tyr) | Moderate |

| | | | | | | | | | | | | | | | |
|---|---|---|---|---|---|---|---|---|---|---|---|---|---|---|---|
| 3 | 9675504 | 145/A | 41.2 ± 1.1 | 6/C | 38 ± 3.2 | $5.32 \times 10^{-1}$ | 0.3 | $6.01 \times 10^{-1}$ | 0.0 | $8.30 \times 10^{-1}$ | | Amino acid substitution (Ile284Leu) | Moderate | | |
| 3 | 9675640 | 7/T | 46.7 ± 3.6 | 144/G | 40.8 ± 1.1 | $2.11 \times 10^{-1}$ | 1.5 | $2.23 \times 10^{-1}$ | 1.2 | $2.72 \times 10^{-1}$ | | Amino acid substitution (Arg298Ser) | Moderate | | |
| 3 | 9675748 | 135/G | 41.2 ± 1.1 | 16/C | 40.5 ± 2.5 | $8.33 \times 10^{-1}$ | 0.2 | $6.58 \times 10^{-1}$ | 0.2 | $6.59 \times 10^{-1}$ | | Amino acid substitution (Lys334Asn) | Moderate | | |
| 3 | 9675806 | 19/A | 44.4 ± 3.1 | 132/G | 40.6 ± 1.1 | $1.99 \times 10^{-1}$ | 2.3 | $1.28 \times 10^{-1}$ | 1.3 | $2.64 \times 10^{-1}$ | | Amino acid substitution (Thr354Ala) | Moderate | | |
| 3 | 9675825 | 8/C | 45.5 ± 3.9 | 143/A | 40.8 ± 1.1 | $2.92 \times 10^{-1}$ | 0.4 | $5.22 \times 10^{-1}$ | 0.0 | $8.45 \times 10^{-1}$ | | Amino acid substitution (Gln360Pro) | Moderate | | |
| 3 | 9675827 | 136/C | 41.2 ± 1.1 | 15/A | 40.1 ± 2.9 | $7.49 \times 10^{-1}$ | 0.2 | $6.30 \times 10^{-1}$ | 0.7 | $4.16 \times 10^{-1}$ | | Amino acid substitution (Gln361Lys) | Moderate | | |
| 3 | 9675837 | 58/G | 43 ± 1.8 | 93/A | 39.9 ± 1.2 | $1.20 \times 10^{-1}$ | **3.9** | $5.05 \times 10^{-2}$ | 0.0 | $8.88 \times 10^{-1}$ | | Amino acid substitution (Asp364Gly) | Moderate | | |
| 3 | 9675996 | 47/A | 45.1 ± 2.1 | 104/G | 39.3 ± 1.1 | $6.49 \times 10^{-3}$ | **13.0** | $\mathbf{4.22 \times 10^{-4}}$ | 2.1 | $1.45 \times 10^{-1}$ | ✓ | Synonymous polymorphism | Moderate | | |
| 3 | 22348534 | 125/A | 41.3 ± 1.2 | 26/G | 39.9 ± 2.3 | $5.97 \times 10^{-1}$ | 0.0 | $9.81 \times 10^{-1}$ | 0.0 | $9.96 \times 10^{-1}$ | | Promoter polymorphism | Moderate | AT3G60490 | |
| 3 | 22348535 | 125/C | 41.3 ± 1.2 | 26/T | 40 ± 2 | $6.16 \times 10^{-1}$ | 0.0 | $9.64 \times 10^{-1}$ | 0.0 | $8.81 \times 10^{-1}$ | | Promoter polymorphism | Moderate | | |
| 3 | 22348562 | 131/C | 41.3 ± 1.1 | 20/T | 39.8 ± 2.8 | $6.20 \times 10^{-1}$ | 0.0 | $9.97 \times 10^{-1}$ | 0.0 | $8.52 \times 10^{-1}$ | | Promoter polymorphism | Moderate | | |
| 3 | 22348589 | 124/A | 41.1 ± 1.2 | 27/C | 41 ± 2.1 | $9.60 \times 10^{-1}$ | 0.3 | $5.97 \times 10^{-1}$ | 0.5 | $4.93 \times 10^{-1}$ | | Promoter polymorphism | Moderate | | |
| 3 | 22348652 | 121/T | 41.4 ± 1.2 | 30/C | 39.9 ± 2.1 | $5.66 \times 10^{-1}$ | 0.0 | $9.63 \times 10^{-1}$ | 0.0 | $9.40 \times 10^{-1}$ | | Promoter polymorphism | Moderate | | |
| 3 | 22348678 | 11/C | 41.5 ± 3.4 | 140/A | 41.1 ± 1.1 | $8.97 \times 10^{-1}$ | 0.0 | $8.56 \times 10^{-1}$ | 0.0 | $9.05 \times 10^{-1}$ | | Promoter polymorphism | Moderate | | |
| 3 | 22348707 | 14/G | 44.6 ± 3.9 | 137/GA | 40.7 ± 1.1 | $2.55 \times 10^{-1}$ | 0.6 | $4.31 \times 10^{-1}$ | 0.0 | $9.32 \times 10^{-1}$ | | Promoter polymorphism | Moderate | | |
| 3 | 22348736 | 123/A | 41.9 ± 1.1 | 28/G | 37.5 ± 2.5 | $8.36 \times 10^{-2}$ | 1.3 | $2.49 \times 10^{-1}$ | 0.4 | $5.51 \times 10^{-1}$ | | Promoter polymorphism | Moderate | | |
| 3 | 22348771 | 31/C | 41.3 ± 2.5 | 120/G | 41 ± 1.1 | $9.14 \times 10^{-1}$ | 1.2 | $2.71 \times 10^{-1}$ | 1.8 | $1.85 \times 10^{-1}$ | | Promoter polymorphism | Moderate | | |
| 3 | 22348772 | 48/A | 44.2 ± 1.8 | 103/AT | 39.6 ± 1.2 | $\mathbf{2.94 \times 10^{-2}}$ | 2.9 | $9.07 \times 10^{-2}$ | 1.7 | $1.93 \times 10^{-1}$ | | Promoter polymorphism | Moderate | | |
| 3 | 22348787 | 137/T | 41.2 ± 1.1 | 14/G | 40.2 ± 3.4 | $7.71 \times 10^{-1}$ | 0.1 | $7.78 \times 10^{-1}$ | 0.5 | $4.89 \times 10^{-1}$ | | Promoter polymorphism | Moderate | | |
| 3 | 22348819 | 136/A | 41.1 ± 1.1 | 15/T | 40.6 ± 2.8 | $8.76 \times 10^{-1}$ | 0.2 | $6.97 \times 10^{-1}$ | 0.5 | $4.98 \times 10^{-1}$ | | Promoter polymorphism | Moderate | | |
| 3 | 22348829 | 139/A | 41.2 ± 1.1 | 12/T | 40 ± 2.9 | $7.48 \times 10^{-1}$ | 0.0 | $8.60 \times 10^{-1}$ | 0.2 | $6.81 \times 10^{-1}$ | | Promoter polymorphism | Moderate | | |
| 3 | 22348983 | 11/G | 41.6 ± 3.3 | 140/T | 41 ± 1.1 | $8.78 \times 10^{-1}$ | 0.4 | $5.46 \times 10^{-1}$ | 0.2 | $6.19 \times 10^{-1}$ | | Promoter polymorphism | Moderate | | |
| 3 | 22348989 | 136/G | 41.3 ± 1.1 | 15/C | 38.8 ± 3.1 | $4.48 \times 10^{-1}$ | 0.1 | $7.53 \times 10^{-1}$ | 0.5 | $4.76 \times 10^{-1}$ | | Promoter polymorphism | Moderate | | |
| 3 | 22349041 | 11/T | 41.2 ± 2.7 | 140/A | 41.1 ± 1.1 | $9.80 \times 10^{-1}$ | 0.2 | $6.52 \times 10^{-1}$ | 0.2 | $6.86 \times 10^{-1}$ | | Promoter polymorphism | Moderate | | |
| 3 | 22349074 | 134/A | 41.2 ± 1.1 | 17/T | 39.9 ± 2.9 | $6.63 \times 10^{-1}$ | 0.0 | $8.72 \times 10^{-1}$ | 0.0 | $9.09 \times 10^{-1}$ | | Promoter polymorphism | Moderate | | |
| 3 | 22349168 | 36/A | 45.8 ± 2.1 | 115/T | 39.6 ± 1.1 | $\mathbf{7.68 \times 10^{-3}}$ | **3.9** | $\mathbf{4.99 \times 10^{-2}}$ | 2.6 | $1.09 \times 10^{-1}$ | | Promoter polymorphism | Moderate | | |
| 3 | 22349176 | 33/G | 44.5 ± 2.4 | 118/GATAT | 40.1 ± 1.1 | $6.93 \times 10^{-2}$ | 2.4 | $1.26 \times 10^{-1}$ | 1.6 | $2.02 \times 10^{-1}$ | | Promoter polymorphism | Moderate | | |
| 3 | 22349238 | 129/A | 41.4 ± 1.2 | 22/T | 39.5 ± 2.2 | $5.11 \times 10^{-1}$ | 0.0 | $9.83 \times 10^{-1}$ | 0.0 | $9.82 \times 10^{-1}$ | | Promoter polymorphism | Moderate | | |
| 3 | 22349254 | 11/C | 41.3 ± 3.3 | 140/T | 41.1 ± 1.1 | $9.58 \times 10^{-1}$ | 0.5 | $5.03 \times 10^{-1}$ | 0.5 | $4.91 \times 10^{-1}$ | | Promoter polymorphism | Moderate | | |
| 3 | 22349256 | 140/T | 41.3 ± 1.1 | 11/C | 38.6 ± 4.2 | $4.76 \times 10^{-1}$ | 0.0 | $8.48 \times 10^{-1}$ | 0.2 | $6.54 \times 10^{-1}$ | | Promoter polymorphism | Moderate | | |
| 3 | 22349295 | 138/T | 41.2 ± 1.1 | 13/G | 40.1 ± 3.9 | $7.55 \times 10^{-1}$ | 0.1 | $7.12 \times 10^{-1}$ | 0.2 | $6.72 \times 10^{-1}$ | | Promoter polymorphism | Moderate | | |
| 3 | 22349300 | 136/T | 41.2 ± 1.1 | 15/A | 39.8 ± 3.4 | $6.60 \times 10^{-1}$ | 0.1 | $7.45 \times 10^{-1}$ | 0.5 | $4.76 \times 10^{-1}$ | | Promoter polymorphism | Moderate | | |
| 3 | 22349360 | 32/C | 47.1 ± 2.2 | 119/A | 39.5 ± 1.1 | $\mathbf{1.44 \times 10^{-3}}$ | **6.8** | $\mathbf{1.01 \times 10^{-2}}$ | **3.1** | $7.85 \times 10^{-2}$ | | Promoter polymorphism | Moderate | | |
| 3 | 22349371 | 136/A | 41.1 ± 1.1 | 15/G | 40.9 ± 2.9 | $9.46 \times 10^{-1}$ | 0.4 | $5.05 \times 10^{-1}$ | 1.7 | $1.97 \times 10^{-1}$ | | Promoter polymorphism | Moderate | | |
| 3 | 22349416 | 136/A | 41.4 ± 1.1 | 15/T | 38.4 ± 3.1 | $3.74 \times 10^{-1}$ | 0.1 | $7.17 \times 10^{-1}$ | 0.2 | $6.63 \times 10^{-1}$ | | Promoter polymorphism | Moderate | | |
| 3 | 22349431 | 119/C | 41.3 ± 1.2 | 32/CT | 40.3 ± 2.2 | $6.76 \times 10^{-1}$ | 0.4 | $5.54 \times 10^{-1}$ | 0.8 | $3.60 \times 10^{-1}$ | | Promoter polymorphism | Moderate | | |
| 3 | 22349520 | 132/C | 41.4 ± 1.1 | 19/T | 38.7 ± 2.8 | $3.72 \times 10^{-1}$ | 0.0 | $8.34 \times 10^{-1}$ | 0.0 | $9.78 \times 10^{-1}$ | | Promoter polymorphism | Moderate | | |
| 3 | 22349579 | 130/T | 41.4 ± 1.1 | 21/A | 39.1 ± 2.3 | $4.15 \times 10^{-1}$ | 0.0 | $8.77 \times 10^{-1}$ | 0.2 | $6.55 \times 10^{-1}$ | | Amino acid substitution (Ile18Asn) | Moderate | | |
| 3 | 22349652 | 150/CA | 41.2 ± 1 | 1/C | 22.2 ± 0 | | 2.3 | $1.34 \times 10^{-1}$ | 0.3 | $5.90 \times 10^{-1}$ | | Frameshift variant (Lys43fs) | **High** | | |
| 3 | 22349680 | 137/T | 41.6 ± 1.1 | 14/G | 35.8 ± 2.3 | $8.53 \times 10^{-2}$ | 1.9 | $1.75 \times 10^{-1}$ | 2.1 | $1.48 \times 10^{-1}$ | | Amino acid substitution (Ser52Ala) | Moderate | | |
| 3 | 22350261 | 10/T | 44.6 ± 4 | 132/C | 41.1 ± 1.1 | $3.89 \times 10^{-1}$ | 1.4 | $2.46 \times 10^{-1}$ | 0.3 | $7.68 \times 10^{-1}$ | | Amino acid substitution (Phe245Leu) | Moderate | | |
| 3 | 22352929 | 121/G | 41.1 ± 1.2 | 30/A | 40.9 ± 1.8 | $9.23 \times 10^{-1}$ | 0.0 | $9.14 \times 10^{-1}$ | 0.0 | $9.65 \times 10^{-1}$ | | Promoter polymorphism | Moderate | AT3G60500 | *CER7* |

| | | | | | | | | | | | | | | |
|---|---|---|---|---|---|---|---|---|---|---|---|---|---|---|
| 3 | 22352953 | 130/T | 41.2 ± 1.1 | 21/C | 40.3 ± 2.3 | $7.59 \times 10^{-1}$ | 0.1 | $7.52 \times 10^{-1}$ | 0.2 | $6.53 \times 10^{-1}$ | | Promoter polymorphism | Moderate | |
| 3 | 22352955 | 128/C | 41.3 ± 1.2 | 23/T | 39.7 ± 2.2 | $5.52 \times 10^{-1}$ | 0.4 | $5.29 \times 10^{-1}$ | 0.0 | $9.40 \times 10^{-1}$ | | Promoter polymorphism | Moderate | |
| 3 | 22353010 | 126/C | 41.1 ± 1.2 | 25/T | 41 ± 2.1 | $9.79 \times 10^{-1}$ | 0.0 | $9.67 \times 10^{-1}$ | 0.4 | $5.26 \times 10^{-1}$ | | Promoter polymorphism | Moderate | |
| 3 | 22353027 | 135/C | 41.3 ± 1.1 | **16/T** | 39 ± 2.3 | $4.62 \times 10^{-1}$ | 0.5 | $4.61 \times 10^{-1}$ | 0.0 | $9.42 \times 10^{-1}$ | | Promoter polymorphism | Moderate | |
| 3 | 22353034 | 53/CA | 42.2 ± 1.6 | 98/C | 40.5 ± 1.3 | $4.08 \times 10^{-1}$ | 0.7 | $4.10 \times 10^{-1}$ | 0.0 | $9.53 \times 10^{-1}$ | | Promoter polymorphism | Moderate | |
| 3 | 22353043 | 11/G | 41.6 ± 2.3 | 140/A | 41 ± 1.1 | $8.76 \times 10^{-1}$ | 0.0 | $9.86 \times 10^{-1}$ | 0.8 | $3.77 \times 10^{-1}$ | | Promoter polymorphism | Moderate | |
| 3 | 22353099 | 122/A | 41.4 ± 1.2 | 29/C | 39.6 ± 1.8 | $4.67 \times 10^{-1}$ | 0.7 | $3.90 \times 10^{-1}$ | 0.4 | $5.52 \times 10^{-1}$ | | Promoter polymorphism | Moderate | |
| 3 | 22353107 | 123/G | 41.6 ± 1.2 | 28/C | 39 ± 1.6 | $3.21 \times 10^{-1}$ | 1.1 | $2.90 \times 10^{-1}$ | 0.3 | $6.05 \times 10^{-1}$ | | Promoter polymorphism | Moderate | |
| 3 | 22353179 | 55/A | 44.1 ± 1.7 | 96/G | 39.3 ± 1.3 | **$1.96 \times 10^{-2}$** | **4.1** | **$4.45 \times 10^{-2}$** | 1.4 | $2.36 \times 10^{-1}$ | | Promoter polymorphism | Moderate | |
| 3 | 22353188 | 54/T | 43.8 ± 1.8 | 97/G | 39.6 ± 1.2 | **$3.95 \times 10^{-2}$** | 3.0 | $8.69 \times 10^{-2}$ | 0.7 | $4.13 \times 10^{-1}$ | | Promoter polymorphism | Moderate | |
| 3 | 22353198 | 138/A | 41.7 ± 1.1 | 13/T | 34.3 ± 3.7 | **$3.49 \times 10^{-2}$** | **7.8** | **$5.98 \times 10^{-3}$** | **4.6** | **$3.38 \times 10^{-2}$** | | Promoter polymorphism | Moderate | |
| 3 | 22353203 | 54/G | 44 ± 1.8 | 97/T | 39.5 ± 1.2 | **$2.80 \times 10^{-2}$** | **3.2** | $7.64 \times 10^{-2}$ | 0.5 | $4.99 \times 10^{-1}$ | | Promoter polymorphism | Moderate | |
| 3 | 22353316 | 55/A | 44.8 ± 1.8 | 96/G | 39 ± 1.2 | **$4.42 \times 10^{-3}$** | **6.5** | **$1.16 \times 10^{-2}$** | 2.1 | $1.54 \times 10^{-1}$ | | Promoter polymorphism | Moderate | |
| 3 | 22353390 | 60/A | 44.3 ± 1.7 | 91/C | 38.9 ± 1.2 | **$7.21 \times 10^{-3}$** | **6.4** | **$1.27 \times 10^{-2}$** | 2.0 | $1.56 \times 10^{-1}$ | | Promoter polymorphism | Moderate | |
| 3 | 22353822 | 11/C | 42.3 ± 3.1 | 139/G | 41 ± 1.1 | $7.43 \times 10^{-1}$ | 0.1 | $9.19 \times 10^{-1}$ | 0.1 | $8.62 \times 10^{-1}$ | | Promoter polymorphism | Moderate | |
| 3 | 22353891 | 8/C | 42.3 ± 4 | 143/A | 41 ± 1.1 | $7.67 \times 10^{-1}$ | 0.2 | $6.66 \times 10^{-1}$ | 0.0 | $8.98 \times 10^{-1}$ | | Promoter polymorphism | Moderate | |
| 3 | 22353955 | 6/G | 41.4 ± 5.3 | 145/T | 41.1 ± 1.1 | $9.46 \times 10^{-1}$ | 0.1 | $7.55 \times 10^{-1}$ | 0.2 | $6.32 \times 10^{-1}$ | | 5'-UTR polymorphism | Moderate | |
| 3 | 22353984 | 7/A | 45.8 ± 3.9 | 144/G | 40.9 ± 1.1 | $2.99 \times 10^{-1}$ | 2.3 | $1.29 \times 10^{-1}$ | 1.4 | $2.33 \times 10^{-1}$ | | 5'-UTR polymorphism | Moderate | |
| 3 | 22354029 | 140/G | 41.6 ± 1.1 | 11/A | 34.1 ± 4.4 | **$4.84 \times 10^{-2}$** | **8.3** | **$4.61 \times 10^{-3}$** | **4.9** | **$2.84 \times 10^{-2}$** | | 5'-UTR polymorphism | Moderate | |
| 3 | 22354080 | 9/G | 46.2 ± 4.1 | 142/A | 40.8 ± 1.1 | $1.93 \times 10^{-1}$ | **3.7** | $5.51 \times 10^{-2}$ | 1.5 | $2.19 \times 10^{-1}$ | | 5'-UTR polymorphism | Moderate | |
| 3 | 22354081 | 9/A | 46.2 ± 4.1 | 142/G | 40.8 ± 1.1 | $1.93 \times 10^{-1}$ | **3.7** | $5.51 \times 10^{-2}$ | 1.5 | $2.19 \times 10^{-1}$ | | 5'-UTR polymorphism | Moderate | |
| 3 | 22354096 | 10/T | 44.6 ± 4 | 141/C | 40.8 ± 1.1 | $3.47 \times 10^{-1}$ | 2.4 | $1.26 \times 10^{-1}$ | 0.4 | $5.06 \times 10^{-1}$ | | 5'-UTR polymorphism | Moderate | |
| 3 | 22354173 | 11/T | 44.3 ± 3.7 | 140/C | 40.8 ± 1.1 | $3.67 \times 10^{-1}$ | 2.3 | $1.32 \times 10^{-1}$ | 0.4 | $5.40 \times 10^{-1}$ | | 5'-UTR polymorphism | Moderate | |
| 3 | 22354195 | 9/A | 46.5 ± 4 | 142/G | 40.7 ± 1.1 | $1.71 \times 10^{-1}$ | **4.3** | **$3.99 \times 10^{-2}$** | 1.7 | $1.95 \times 10^{-1}$ | | 5'-UTR polymorphism | Moderate | |
| 3 | 22354196 | 9/C | 46.5 ± 4 | 142/T | 40.7 ± 1.1 | $1.71 \times 10^{-1}$ | **4.3** | **$3.99 \times 10^{-2}$** | 1.7 | $1.95 \times 10^{-1}$ | | 5'-UTR polymorphism | Moderate | |
| 3 | 22354206 | 9/C | 46.5 ± 4 | 142/T | 40.7 ± 1.1 | $1.71 \times 10^{-1}$ | **4.3** | **$3.99 \times 10^{-2}$** | 1.7 | $1.95 \times 10^{-1}$ | | 5'-UTR polymorphism | Moderate | |
| 3 | 22354698 | 140/A | 41.1 ± 1.1 | 11/C | 40.6 ± 4 | $8.95 \times 10^{-1}$ | 1.2 | $2.78 \times 10^{-1}$ | 1.1 | $3.07 \times 10^{-1}$ | | Amino acid substitution (Asn77Thr) | Moderate | |
| 3 | 22354733 | 140/C | 41.1 ± 1.1 | 11/G | 40.6 ± 4 | $8.95 \times 10^{-1}$ | 1.2 | $2.78 \times 10^{-1}$ | 1.1 | $3.07 \times 10^{-1}$ | | Amino acid substitution (Pro89Ala) | Moderate | |
| 3 | 22355783 | 11/C | 44.3 ± 3.7 | 140/T | 40.8 ± 1.1 | $3.67 \times 10^{-1}$ | 2.3 | $1.32 \times 10^{-1}$ | 0.5 | $4.70 \times 10^{-1}$ | | Amino acid substitution (Cys282Arg) | Moderate | |
| 3 | 22355901 | 11/G | 44.3 ± 3.7 | 140/C | 40.8 ± 1.1 | $3.67 \times 10^{-1}$ | 2.3 | $1.32 \times 10^{-1}$ | 0.5 | $4.70 \times 10^{-1}$ | | Amino acid substitution (Ala321Gly) | Moderate | |
| 3 | 22356075 | 9/C | 49.5 ± 4.1 | 142/T | 40.6 ± 1.1 | **$3.20 \times 10^{-2}$** | **6.8** | **$1.02 \times 10^{-2}$** | **6.2** | **$1.40 \times 10^{-2}$** | ✓ | Synonymous polymorphism | Moderate | |
| 3 | 22356350 | 11/T | 44.3 ± 3.7 | 140/A | 40.8 ± 1.1 | $3.67 \times 10^{-1}$ | 2.3 | $1.32 \times 10^{-1}$ | 0.5 | $4.70 \times 10^{-1}$ | | Amino acid substitution (Glu409Asp) | Moderate | |
| 3 | 22357670 | 11/T | 44.3 ± 3.7 | 140/C | 40.8 ± 1.1 | $3.67 \times 10^{-1}$ | 2.3 | $1.32 \times 10^{-1}$ | 0.3 | $5.88 \times 10^{-1}$ | | Amino acid substitution (Leu313Phe) | Moderate | AT3G60510 |
| 3 | 22357865 | 11/T | 44.3 ± 3.7 | 140/G | 40.8 ± 1.1 | $3.67 \times 10^{-1}$ | 2.3 | $1.32 \times 10^{-1}$ | 0.3 | $5.88 \times 10^{-1}$ | | Amino acid substitution (Asp273Glu) | Moderate | |
| 3 | 22358035 | 11/A | 44.3 ± 3.7 | 140/T | 40.8 ± 1.1 | $3.67 \times 10^{-1}$ | 2.3 | $1.32 \times 10^{-1}$ | 0.3 | $5.88 \times 10^{-1}$ | | Amino acid substitution (Lys247Asn) | Moderate | |
| 3 | 22359296 | 11/C | 44.3 ± 3.7 | 140/T | 40.8 ± 1.1 | $3.67 \times 10^{-1}$ | 2.3 | $1.32 \times 10^{-1}$ | 0.3 | $5.88 \times 10^{-1}$ | | Amino acid substitution (Ile100Val) | Moderate | |

| | | | | | | | | | | | | | | | |
|---|---|---|---|---|---|---|---|---|---|---|---|---|---|---|---|
| 3 | 22360258 | 18/G | 42.6 ± 2.4 | 133/C | 40.9 ± 1.1 | $5.85 \times 10^{-1}$ | 1.9 | $1.71 \times 10^{-1}$ | 1.0 | $3.18 \times 10^{-1}$ | | Promoter polymorphism | Moderate | | |
| 3 | 22360700 | 124/G | 41.1 ± 1.1 | 27/A | 40.8 ± 2.5 | $9.08 \times 10^{-1}$ | 1.0 | $3.08 \times 10^{-1}$ | 0.2 | $6.76 \times 10^{-1}$ | | Promoter polymorphism | Moderate | | |
| 3 | 22360715 | 120/C | 42.1 ± 1.1 | 31/T | 37.4 ± 2.3 | $5.56 \times 10^{-2}$ | 2.0 | $1.56 \times 10^{-1}$ | 0.3 | $6.14 \times 10^{-1}$ | | Promoter polymorphism | Moderate | | |
| 3 | 22360820 | 13/A | 44 ± 3.5 | 138/G | 40.8 ± 1.1 | $3.62 \times 10^{-1}$ | 2.4 | $1.20 \times 10^{-1}$ | 0.4 | $5.23 \times 10^{-1}$ | | Promoter polymorphism | Moderate | | |
| 3 | 22360882 | 124/T | 42.3 ± 1.1 | 27/G | 35.4 ± 2.1 | $6.54 \times 10^{-3}$ | **5.9** | **$1.64 \times 10^{-2}$** | 2.0 | $1.63 \times 10^{-1}$ | | Promoter polymorphism | Moderate | | |
| 3 | 22360904 | 130/T | 41.9 ± 1.1 | 21/A | 36.1 ± 2.6 | **$4.08 \times 10^{-2}$** | 2.3 | $1.35 \times 10^{-1}$ | 0.4 | $5.39 \times 10^{-1}$ | | Promoter polymorphism | Moderate | | |
| 3 | 22360964 | 83/G | 42.1 ± 1.4 | 68/A | 39.8 ± 1.5 | $2.50 \times 10^{-1}$ | 0.0 | $8.44 \times 10^{-1}$ | 0.0 | $8.52 \times 10^{-1}$ | | Promoter polymorphism | Moderate | | |
| 4 | 9200675 | 35/G | 46.3 ± 2.5 | 116/A | 39.5 ± 1.1 | **$3.85 \times 10^{-3}$** | **10.3** | **$1.62 \times 10^{-3}$** | **7.8** | **$5.96 \times 10^{-3}$** | ✓ | Intron polymorphism | Moderate | AT4G16260 | |
| 4 | 9200798 | 65/C | 41.9 ± 1.7 | 86/T | 40.5 ± 1.3 | $5.04 \times 10^{-1}$ | 1.0 | $3.17 \times 10^{-1}$ | 0.8 | $3.81 \times 10^{-1}$ | | Amino acid substitution (Val179Ile) | Moderate | | |
| 4 | 9201546 | 18/A | 41.3 ± 2.9 | 133/T | 41.1 ± 1.1 | $9.32 \times 10^{-1}$ | 0.0 | $8.89 \times 10^{-1}$ | 0.2 | $6.21 \times 10^{-1}$ | | 5'-UTR polymorphism | Moderate | | |
| 4 | 9201644 | 71/G | 41.6 ± 1.5 | 80/A | 40.6 ± 1.4 | $6.33 \times 10^{-1}$ | 0.2 | $6.36 \times 10^{-1}$ | 0.3 | $5.88 \times 10^{-1}$ | | Promoter polymorphism | Moderate | | |
| 4 | 9201774 | 131/G | 41.4 ± 1.2 | 20/A | 39 ± 1.6 | $4.06 \times 10^{-1}$ | 0.5 | $4.96 \times 10^{-1}$ | 0.1 | $7.49 \times 10^{-1}$ | | Promoter polymorphism | Moderate | | |
| 4 | 9201791 | 132/T | 41.4 ± 1.1 | 19/G | 39.1 ± 2.9 | $4.44 \times 10^{-1}$ | 0.2 | $6.70 \times 10^{-1}$ | 0.7 | $4.07 \times 10^{-1}$ | | Promoter polymorphism | Moderate | | |
| 4 | 9202034 | 13/G | 50.4 ± 2.9 | 133/A | 40.3 ± 1.1 | **$4.18 \times 10^{-3}$** | **3.8** | **$2.51 \times 10^{-2}$** | 3.0 | $5.47 \times 10^{-2}$ | | Promoter polymorphism | Moderate | | |
| 4 | 9202039 | 140/T | 41.3 ± 1.1 | 11/C | 38.8 ± 3.1 | $5.18 \times 10^{-1}$ | 0.0 | $8.79 \times 10^{-1}$ | 0.1 | $7.26 \times 10^{-1}$ | | Promoter polymorphism | Moderate | | |
| 4 | 9202062 | 133/T | 41.3 ± 1.2 | 18/A | 39.2 ± 1.7 | $4.88 \times 10^{-1}$ | 0.3 | $5.69 \times 10^{-1}$ | 0.0 | $8.27 \times 10^{-1}$ | | Promoter polymorphism | Moderate | | |
| 4 | 9202144 | 133/T | 41.5 ± 1.1 | 18/G | 38.3 ± 3 | $3.11 \times 10^{-1}$ | 0.5 | $4.80 \times 10^{-1}$ | 0.2 | $6.78 \times 10^{-1}$ | | Promoter polymorphism | Moderate | | |
| 4 | 9202188 | 82/T | 41.8 ± 1.4 | 69/C | 40.2 ± 1.5 | $4.27 \times 10^{-1}$ | 0.1 | $7.89 \times 10^{-1}$ | 0.5 | $5.02 \times 10^{-1}$ | | Promoter polymorphism | Moderate | | |
| 4 | 9613640 | 3/T | 48.4 ± 8.4 | 148/C | 40.9 ± 1 | $2.94 \times 10^{-1}$ | 1.5 | $2.16 \times 10^{-1}$ | 0.5 | $4.66 \times 10^{-1}$ | | Amino acid substitution (Gly4210Arg) | Moderate | AT4G17140 | *VPS13B* |
| 4 | 9613758 | 8/T | 49.3 ± 5 | 143/G | 40.6 ± 1 | $5.03 \times 10^{-2}$ | **3.8** | $5.47 \times 10^{-2}$ | 1.9 | $1.75 \times 10^{-1}$ | | Amino acid substitution (Gln4170His) | Moderate | | |
| 4 | 9614901 | 137/T | 41.4 ± 1.1 | 14/A | 38.3 ± 2.5 | $3.62 \times 10^{-1}$ | 1.8 | $1.86 \times 10^{-1}$ | 1.0 | $3.21 \times 10^{-1}$ | | Amino acid substitution (Lys3996Ile) | Moderate | | |
| 4 | 9615303 | 134/C | 41.5 ± 1.1 | 17/T | 37.8 ± 3 | $2.38 \times 10^{-1}$ | 1.9 | $1.71 \times 10^{-1}$ | 1.1 | $2.91 \times 10^{-1}$ | | Amino acid substitution (Val3947Ile) | Moderate | | |
| 4 | 9616210 | 8/C | 44.4 ± 1.9 | 143/G | 40.9 ± 1.1 | $4.35 \times 10^{-1}$ | 0.9 | $3.48 \times 10^{-1}$ | 2.0 | $1.60 \times 10^{-1}$ | | Amino acid substitution (Leu3758Val) | Moderate | | |
| 4 | 9617662 | 121/T | 41.1 ± 1.1 | 30/A | 41 ± 2.7 | $9.52 \times 10^{-1}$ | 0.8 | $3.72 \times 10^{-1}$ | 0.4 | $5.51 \times 10^{-1}$ | | Amino acid substitution (Asn3508Tyr) | Moderate | | |
| 4 | 9617894 | 137/C | 41.4 ± 1.1 | 14/G | 38.3 ± 2.5 | $3.62 \times 10^{-1}$ | 1.8 | $1.86 \times 10^{-1}$ | 1.0 | $3.21 \times 10^{-1}$ | | Amino acid substitution (Glu3464Gln) | Moderate | | |
| 4 | 9618843 | 24/A | 45.1 ± 2.8 | 127/G | 40.3 ± 1.1 | $7.53 \times 10^{-2}$ | **4.3** | **$4.07 \times 10^{-2}$** | 0.1 | $7.55 \times 10^{-1}$ | | Amino acid substitution (Tyr3231His) | Moderate | | |
| 4 | 9618851 | 10/A | 43 ± 1.8 | 141/T | 41 ± 1.1 | $6.08 \times 10^{-1}$ | 0.6 | $4.58 \times 10^{-1}$ | 1.1 | $2.93 \times 10^{-1}$ | | Amino acid substitution (His3228Leu) | Moderate | | |
| 4 | 9618868 | 150/AT | 41.1 ± 1 | 1/A | 33 ± 0 | | 0.3 | $6.16 \times 10^{-1}$ | 0.0 | $9.83 \times 10^{-1}$ | | Frameshift variant (Asn3222fs) | **High** | | |
| 4 | 9620077 | 14/A | 42.3 ± 3.2 | 137/G | 41 ± 1.1 | $6.98 \times 10^{-1}$ | 0.3 | $5.55 \times 10^{-1}$ | 0.2 | $6.93 \times 10^{-1}$ | | Amino acid substitution (Thr2985Ile) | Moderate | | |
| 4 | 9620390 | 147/C | 41.3 ± 1.1 | 4/T | 31.8 ± 4.8 | $1.22 \times 10^{-1}$ | 2.5 | $1.13 \times 10^{-1}$ | 0.2 | $6.19 \times 10^{-1}$ | | Amino acid substitution (Glu2881Lys) | Moderate | | |
| 4 | 9620566 | 3/G | 42.4 ± 11.6 | 147/T | 41 ± 1 | $8.46 \times 10^{-1}$ | 0.3 | $7.59 \times 10^{-1}$ | 0.9 | $4.15 \times 10^{-1}$ | | Amino acid substitution (Lys2822Ile) | Moderate | | |
| 4 | 9620880 | 11/A | 42.3 ± 3.5 | 140/C | 41 ± 1.1 | $7.40 \times 10^{-1}$ | 0.1 | $7.89 \times 10^{-1}$ | 0.5 | $4.72 \times 10^{-1}$ | | Amino acid substitution (Ile2717Met) | Moderate | | |
| 4 | 9621795 | 146/A | 41.3 ± 1.1 | 5/T | 34.7 ± 5.5 | $2.31 \times 10^{-1}$ | 1.9 | $1.74 \times 10^{-1}$ | 0.9 | $3.50 \times 10^{-1}$ | | Amino acid substitution (Val2635Asp) | Moderate | | |
| 4 | 9621831 | 142/T | 41.4 ± 1.1 | 9/A | 36.7 ± 3.5 | $2.61 \times 10^{-1}$ | 0.7 | $3.91 \times 10^{-1}$ | 0.1 | $7.55 \times 10^{-1}$ | | Amino acid substitution (Glu2623Val) | Moderate | | |
| 4 | 9621840 | 144/A | 41.3 ± 1.1 | 7/G | 37.4 ± 4.5 | $4.11 \times 10^{-1}$ | 0.3 | $5.67 \times 10^{-1}$ | 0.1 | $7.38 \times 10^{-1}$ | | Amino acid substitution (Val2620Ala) | Moderate | | |
| 4 | 9622460 | 4/T | 42 ± 9.9 | 147/C | 41.1 ± 1 | $8.82 \times 10^{-1}$ | 0.1 | $7.25 \times 10^{-1}$ | 0.1 | $7.31 \times 10^{-1}$ | | Amino acid substitution (Asp2511Asn) | Moderate | | |
| 4 | 9622463 | 8/T | 44.7 ± 5.2 | 143/C | 40.9 ± 1.1 | $3.92 \times 10^{-1}$ | 0.3 | $5.89 \times 10^{-1}$ | 0.4 | $5.51 \times 10^{-1}$ | | Amino acid substitution (Val2510Ile) | Moderate | | |
| 4 | 9622466 | 122/G | 41.8 ± 1.2 | 28/A | 38.3 ± 2.2 | $1.69 \times 10^{-1}$ | 0.7 | $5.10 \times 10^{-1}$ | 0.5 | $6.32 \times 10^{-1}$ | | Amino acid substitution (Leu2509Val) | Moderate | | |
| 4 | 9623226 | 148/T | 41.3 ± 1 | 3/C | 29.6 ± 7.5 | $9.79 \times 10^{-2}$ | 1.9 | $1.65 \times 10^{-1}$ | **4.4** | **$3.86 \times 10^{-2}$** | | Amino acid substitution (His2378Arg) | Moderate | | |
| 4 | 9623227 | 14/T | 43.4 ± 2.8 | 137/G | 40.8 ± 1.1 | $4.53 \times 10^{-1}$ | 0.9 | $3.55 \times 10^{-1}$ | 1.0 | $3.31 \times 10^{-1}$ | | Amino acid substitution (His2378Asn) | Moderate | | |
| 4 | 9623310 | 3/T | 46.1 ± 2.1 | 148/C | 41 ± 1.1 | $4.76 \times 10^{-1}$ | 0.1 | $7.67 \times 10^{-1}$ | 0.1 | $8.22 \times 10^{-1}$ | | Amino acid substitution (Ser2350Asn) | Moderate | | |
| 4 | 9623762 | 9/C | 45.8 ± 5.8 | 142/G | 40.8 ± 1 | $2.30 \times 10^{-1}$ | 0.6 | $4.41 \times 10^{-1}$ | 2.8 | $9.74 \times 10^{-2}$ | | Amino acid substitution (Ala2229Gly) | Moderate | | |

| | | | | | | | | | | | | | | | |
|---|---|---|---|---|---|---|---|---|---|---|---|---|---|---|---|
| 4 | 9623937 | 22/C | 47.9 ± 2.4 | 129/T | 39.9 ± 1.1 | **$4.43 \times 10^{-3}$** | **9.8** | **$2.11 \times 10^{-3}$** | 2.8 | $9.72 \times 10^{-2}$ | ✓ | Amino acid substitution (Asp2171Asn) | Moderate | | |
| 4 | 9624793 | 145/A | 41.2 ± 1.1 | 6/T | 37.3 ± 5.3 | $4.36 \times 10^{-1}$ | 0.2 | $6.26 \times 10^{-1}$ | 0.1 | $7.60 \times 10^{-1}$ | | Amino acid substitution (Leu2015His) | Moderate | | |
| 4 | 9625252 | 27/A | 42.8 ± 2.4 | 116/G | 40.6 ± 1.2 | $4.06 \times 10^{-1}$ | 0.5 | $6.04 \times 10^{-1}$ | 0.4 | $6.50 \times 10^{-1}$ | | Amino acid substitution (Phe1914Ser) | Moderate | | |
| 4 | 9625325 | 24/T | 45.7 ± 2.5 | 127/C | 40.2 ± 1.1 | **$4.08 \times 10^{-2}$** | **5.9** | **$1.65 \times 10^{-2}$** | 1.3 | $2.63 \times 10^{-1}$ | | Amino acid substitution (Lys1890Glu) | Moderate | | |
| 4 | 9627209 | 21/T | 48.2 ± 2.5 | 130/C | 39.9 ± 1.1 | **$3.84 \times 10^{-3}$** | **9.7** | **$2.26 \times 10^{-3}$** | **3.6** | $6.04 \times 10^{-2}$ | | Amino acid substitution (Thr1536Ala) | Moderate | | |
| 4 | 9627243 | 144/C | 41.2 ± 1.1 | 7/A | 39.4 ± 5.5 | $7.04 \times 10^{-1}$ | 0.7 | $3.94 \times 10^{-1}$ | 1.9 | $1.69 \times 10^{-1}$ | | Amino acid substitution (Glu1524Asp) | Moderate | | |
| 4 | 9629041 | 138/A | 41.3 ± 1.1 | 13/T | 38.9 ± 2.7 | $5.00 \times 10^{-1}$ | 1.3 | $2.65 \times 10^{-1}$ | 0.8 | $3.82 \times 10^{-1}$ | | Amino acid substitution (Ser1143Thr) | Moderate | | |
| 4 | 9629316 | 7/T | 44.1 ± 2.2 | 144/C | 40.9 ± 1.1 | $5.06 \times 10^{-1}$ | 0.8 | $3.75 \times 10^{-1}$ | 1.3 | $2.59 \times 10^{-1}$ | | Amino acid substitution (Ser1086Asn) | Moderate | | |
| 4 | 9629622 | 21/T | 49.3 ± 2.6 | 130/C | 39.8 ± 1.1 | $6.90 \times 10^{-4}$ | **12.5** | **$5.46 \times 10^{-4}$** | **3.9** | **$4.88 \times 10^{-2}$** | ✓ | Intron polymorphism | Moderate | | |
| 4 | 9629911 | 24/C | 46.6 ± 2.4 | 127/A | 40.1 ± 1.1 | **$1.59 \times 10^{-2}$** | **7.8** | **$5.84 \times 10^{-3}$** | 2.2 | $1.43 \times 10^{-1}$ | | Amino acid substitution (Gly940Val) | Moderate | | |
| 4 | 9629912 | 144/C | 41.1 ± 1.1 | 7/A | 40.6 ± 4 | $9.19 \times 10^{-1}$ | 0.0 | $9.65 \times 10^{-1}$ | 0.0 | $8.81 \times 10^{-1}$ | | Amino acid substitution (Gly940Cys) | Moderate | | |
| 4 | 9631341 | 5/T | 47.4 ± 3.5 | 146/C | 40.9 ± 1.1 | $2.39 \times 10^{-1}$ | 1.2 | $2.80 \times 10^{-1}$ | 1.8 | $1.83 \times 10^{-1}$ | | Amino acid substitution (Gly766Ser) | Moderate | | |
| 4 | 9632177 | 22/A | 47.9 ± 2.4 | 129/G | 39.9 ± 1.1 | **$4.43 \times 10^{-3}$** | **9.8** | **$2.11 \times 10^{-3}$** | 2.8 | $9.72 \times 10^{-2}$ | ✓ | Intron polymorphism | Moderate | | |
| 4 | 9633154 | 148/C | 41.3 ± 1.1 | 3/T | 32.9 ± 6.6 | $2.40 \times 10^{-1}$ | 1.4 | $2.37 \times 10^{-1}$ | 0.0 | $9.52 \times 10^{-1}$ | | Amino acid substitution (Gly448Glu) | Moderate | | |
| 4 | 9634010 | 4/A | 45.8 ± 8 | 147/G | 41 ± 1 | $4.31 \times 10^{-1}$ | 0.0 | $8.73 \times 10^{-1}$ | 0.2 | $6.99 \times 10^{-1}$ | | Amino acid substitution (Pro327Ser) | Moderate | | |
| 4 | 9634506 | 122/T | 41.4 ± 1.1 | 29/G | 39.6 ± 2.6 | $4.68 \times 10^{-1}$ | 1.9 | $1.65 \times 10^{-1}$ | 1.5 | $2.28 \times 10^{-1}$ | | Amino acid substitution (Asp221Ala) | Moderate | | |
| 4 | 9636635 | 4/A | 45.8 ± 8 | 147/T | 41 ± 1 | $4.31 \times 10^{-1}$ | 0.0 | $8.73 \times 10^{-1}$ | 0.2 | $6.99 \times 10^{-1}$ | | 5'-UTR polymorphism | Moderate | | |
| 4 | 9636648 | 121/C | 41.1 ± 1.1 | 30/A | 40.9 ± 2.7 | $9.29 \times 10^{-1}$ | 0.7 | $3.92 \times 10^{-1}$ | 0.2 | $6.82 \times 10^{-1}$ | | 5'-UTR polymorphism | Moderate | | |
| 4 | 9636769 | 144/C | 41.3 ± 1.1 | 7/T | 37.4 ± 4.5 | $4.11 \times 10^{-1}$ | 0.3 | $5.67 \times 10^{-1}$ | 0.1 | $7.38 \times 10^{-1}$ | | 5'-UTR polymorphism | Moderate | | |
| 4 | 9636866 | 4/T | 44.6 ± 2.2 | 147/C | 41 ± 1.1 | $5.66 \times 10^{-1}$ | 0.1 | $7.53 \times 10^{-1}$ | 0.2 | $6.51 \times 10^{-1}$ | | 5'-UTR polymorphism | Moderate | | |
| 4 | 9636940 | 65/A | 41.6 ± 1.6 | 86/G | 40.7 ± 1.4 | $6.38 \times 10^{-1}$ | 0.1 | $8.10 \times 10^{-1}$ | 0.0 | $9.18 \times 10^{-1}$ | | Promoter polymorphism | Moderate | | |
| 4 | 9637056 | 14/A | 42.5 ± 3.5 | 137/G | 40.9 ± 1.1 | $6.58 \times 10^{-1}$ | 0.1 | $7.71 \times 10^{-1}$ | 0.0 | $9.81 \times 10^{-1}$ | | Promoter polymorphism | Moderate | | |
| 4 | 9637066 | 136/T | 41.1 ± 1.1 | 15/A | 40.8 ± 3.6 | $9.27 \times 10^{-1}$ | 0.0 | $8.47 \times 10^{-1}$ | 0.4 | $5.52 \times 10^{-1}$ | | Promoter polymorphism | Moderate | | |
| 4 | 9637083 | 81/C | 42.1 ± 1.5 | 70/T | 39.9 ± 1.4 | $2.81 \times 10^{-1}$ | 1.9 | $1.71 \times 10^{-1}$ | 0.5 | $4.79 \times 10^{-1}$ | | Promoter polymorphism | Moderate | | |
| 4 | 9637109 | 135/A | 41.1 ± 1.1 | 16/T | 40.7 ± 2.7 | $8.92 \times 10^{-1}$ | 0.0 | $9.31 \times 10^{-1}$ | 0.3 | $5.92 \times 10^{-1}$ | | Promoter polymorphism | Moderate | | |
| 4 | 9637128 | 11/A | 43.5 ± 5.1 | 140/G | 40.9 ± 1 | $5.01 \times 10^{-1}$ | 0.1 | $7.83 \times 10^{-1}$ | 0.1 | $8.03 \times 10^{-1}$ | | Promoter polymorphism | Moderate | | |
| 4 | 9637162 | 11/T | 43.5 ± 3.4 | 140/C | 40.9 ± 1.1 | $4.90 \times 10^{-1}$ | 0.7 | $4.00 \times 10^{-1}$ | 0.8 | $3.71 \times 10^{-1}$ | | Promoter polymorphism | Moderate | | |
| 4 | 9637331 | 13/A | 43.6 ± 2.3 | 138/G | 40.9 ± 1.1 | $4.47 \times 10^{-1}$ | 1.0 | $3.13 \times 10^{-1}$ | 1.2 | $2.76 \times 10^{-1}$ | | Promoter polymorphism | Moderate | | |
| 4 | 9637393 | 140/G | 41.1 ± 1.1 | 9/T | 40.4 ± 3.6 | $8.65 \times 10^{-1}$ | 0.4 | $7.62 \times 10^{-1}$ | 0.3 | $8.32 \times 10^{-1}$ | | Promoter polymorphism | Moderate | | |
| 4 | 9637398 | 71/G | 41.4 ± 1.5 | 80/A | 40.8 ± 1.4 | $7.72 \times 10^{-1}$ | 0.0 | $9.82 \times 10^{-1}$ | 0.2 | $6.52 \times 10^{-1}$ | | Promoter polymorphism | Moderate | | |
| 4 | 9637411 | 132/G | 41.1 ± 1.1 | 19/A | 41.1 ± 2.3 | $9.92 \times 10^{-1}$ | 0.1 | $7.76 \times 10^{-1}$ | 0.5 | $4.80 \times 10^{-1}$ | | Promoter polymorphism | Moderate | | |
| 4 | 9637418 | 138/GC | 41.6 ± 1.1 | 13/G | 35.4 ± 3.6 | $7.77 \times 10^{-2}$ | **4.2** | **$4.13 \times 10^{-2}$** | 2.2 | $1.44 \times 10^{-1}$ | | Promoter polymorphism | Moderate | | |
| 4 | 9637421 | 129/G | 41.5 ± 1.1 | 22/A | 38.6 ± 3.1 | $3.10 \times 10^{-1}$ | 2.0 | $1.62 \times 10^{-1}$ | 1.0 | $3.24 \times 10^{-1}$ | | Promoter polymorphism | Moderate | | |
| 4 | 9637462 | 54/G | 41.9 ± 1.8 | 96/C | 40.6 ± 1.3 | $5.34 \times 10^{-1}$ | 0.3 | $7.72 \times 10^{-1}$ | 0.0 | $9.93 \times 10^{-1}$ | | Promoter polymorphism | Moderate | | |
| 4 | 9637600 | 138/AT | 41.2 ± 1.1 | 12/A | 39.9 ± 2.2 | $7.29 \times 10^{-1}$ | 0.1 | $8.65 \times 10^{-1}$ | 0.0 | $9.91 \times 10^{-1}$ | | Promoter polymorphism | Moderate | | |
| 4 | 9637612 | 15/A | 42 ± 4.4 | 136/G | 41 ± 1 | $7.67 \times 10^{-1}$ | 0.1 | $7.54 \times 10^{-1}$ | 0.0 | $9.37 \times 10^{-1}$ | | Promoter polymorphism | Moderate | | |
| 5 | 6112374 | 20/A | 44.2 ± 2.5 | 131/G | 40.6 ± 1.1 | $2.21 \times 10^{-1}$ | **3.5** | $6.24 \times 10^{-2}$ | 1.5 | $2.20 \times 10^{-1}$ | | Promoter polymorphism | Moderate | AT5G18440 | *NUFIP* |
| 5 | 6112944 | 30/G | 45.6 ± 2.2 | 121/C | 40 ± 1.1 | $2.34 \times 10^{-2}$ | **5.4** | **$2.17 \times 10^{-2}$** | **3.2** | $7.47 \times 10^{-2}$ | ✓ | 5'-UTR polymorphism | Moderate | | |
| 5 | 6113035 | 145/T | 41.4 ± 1.1 | 6/TC | 33.5 ± 3.7 | $1.17 \times 10^{-1}$ | 2.1 | $1.52 \times 10^{-1}$ | 0.9 | $3.40 \times 10^{-1}$ | | 5'-UTR polymorphism | Moderate | | |
| 5 | 6113036 | 145/A | 41.1 ± 1.1 | 6/G | 40.5 ± 5.6 | $9.02 \times 10^{-1}$ | 0.0 | $9.80 \times 10^{-1}$ | 0.0 | $9.78 \times 10^{-1}$ | | 5'-UTR polymorphism | Moderate | | |
| 5 | 6113038 | 144/A | 41.1 ± 1.1 | 7/C | 40.6 ± 4.8 | $9.10 \times 10^{-1}$ | 0.0 | $9.50 \times 10^{-1}$ | 0.0 | $9.20 \times 10^{-1}$ | | 5'-UTR polymorphism | Moderate | | |
| 5 | 6113044 | 143/A | 41.2 ± 1.1 | 8/T | 38.3 ± 4.7 | $5.05 \times 10^{-1}$ | 0.3 | $6.15 \times 10^{-1}$ | 0.1 | $7.34 \times 10^{-1}$ | | 5'-UTR polymorphism | Moderate | | |

| | | | | | | | | | | | | | | |
|---|---|---|---|---|---|---|---|---|---|---|---|---|---|---|
| 5 | 6113074 | 142/T | 41.4 ± 1.1 | 9/A | 36.6 ± 4.6 | $2.58 \times 10^{-1}$ | 1.1 | $2.93 \times 10^{-1}$ | 1.4 | $2.40 \times 10^{-1}$ | | 5'-UTR polymorphism | Moderate | |
| 5 | 6113279 | 138/T | 41.6 ± 1.1 | 13/A | 36.1 ± 4 | $1.25 \times 10^{-1}$ | 1.7 | $1.94 \times 10^{-1}$ | 1.7 | $1.95 \times 10^{-1}$ | | Amino acid substitution (Ile27Asn) | Moderate | |
| 5 | 6113301 | 138/A | 41.6 ± 1.1 | 13/C | 36.1 ± 4 | $1.25 \times 10^{-1}$ | 1.7 | $1.94 \times 10^{-1}$ | 1.7 | $1.95 \times 10^{-1}$ | | Amino acid substitution (Gln34His) | Moderate | |
| 5 | 6113338 | 138/G | 41.6 ± 1.1 | 13/T | 36.1 ± 4 | $1.25 \times 10^{-1}$ | 1.7 | $1.94 \times 10^{-1}$ | 1.7 | $1.95 \times 10^{-1}$ | | Amino acid substitution (Ala47Ser) | Moderate | |
| 5 | 6113591 | 138/A | 41.6 ± 1.1 | 13/C | 36.1 ± 4 | $1.25 \times 10^{-1}$ | 1.7 | $1.94 \times 10^{-1}$ | 1.7 | $1.95 \times 10^{-1}$ | | Amino acid substitution (His131Pro) | Moderate | |
| 5 | 6113708 | 141/A | 41.5 ± 1.1 | 10/G | 34.8 ± 4.4 | $9.02 \times 10^{-2}$ | 2.6 | $1.07 \times 10^{-1}$ | 2.3 | $1.28 \times 10^{-1}$ | | Amino acid substitution (Gln170Arg) | Moderate | |
| 5 | 6113876 | 139/C | 41.5 ± 1.1 | 12/T | 36.9 ± 4.3 | $2.11 \times 10^{-1}$ | 1.0 | $3.20 \times 10^{-1}$ | 0.8 | $3.76 \times 10^{-1}$ | | Amino acid substitution (Pro192Ser) | Moderate | |
| 5 | 6115420 | 138/A | 41.6 ± 1.1 | 13/C | 36.1 ± 4 | $1.25 \times 10^{-1}$ | 1.7 | $1.94 \times 10^{-1}$ | 1.7 | $1.95 \times 10^{-1}$ | | Amino acid substitution (Lys362Thr) | Moderate | |
| 5 | 6115702 | 139/G | 41.5 ± 1.1 | 12/A | 36.4 ± 4.4 | $1.63 \times 10^{-1}$ | 1.4 | $2.39 \times 10^{-1}$ | 1.4 | $2.34 \times 10^{-1}$ | | Amino acid substitution (Gly456Asp) | Moderate | |
| 5 | 23176756 | 12/C | 48.1 ± 4.4 | 139/T | 40.5 ± 1 | **$3.60 \times 10^{-2}$** | **4.9** | **$2.81 \times 10^{-2}$** | 1.6 | $2.12 \times 10^{-1}$ | | Promoter polymorphism | Moderate | AT5G57200 |
| 5 | 23176761 | 121/G | 42.4 ± 1.2 | 30/T | 36 ± 1.9 | **$9.58 \times 10^{-3}$** | **9.0** | **$3.24 \times 10^{-3}$** | 1.1 | $2.93 \times 10^{-1}$ | | Promoter polymorphism | Moderate | |
| 5 | 23176848 | 87/A | 41.3 ± 1.4 | 64/G | 40.8 ± 1.5 | $7.75 \times 10^{-1}$ | 0.0 | $9.60 \times 10^{-1}$ | 0.1 | $7.21 \times 10^{-1}$ | | Promoter polymorphism | Moderate | |
| 5 | 23176883 | 83/C | 41.4 ± 1.4 | 68/A | 40.7 ± 1.5 | $7.41 \times 10^{-1}$ | 0.0 | $8.64 \times 10^{-1}$ | 0.2 | $6.77 \times 10^{-1}$ | | Promoter polymorphism | Moderate | |
| 5 | 23176900 | 83/A | 41.3 ± 1.4 | 68/T | 40.8 ± 1.5 | $7.84 \times 10^{-1}$ | 0.0 | $9.95 \times 10^{-1}$ | 0.3 | $5.59 \times 10^{-1}$ | | Promoter polymorphism | Moderate | |
| 5 | 23176962 | 67/A | 41.5 ± 1.5 | 84/G | 40.8 ± 1.4 | $7.35 \times 10^{-1}$ | 0.5 | $4.63 \times 10^{-1}$ | 0.3 | $5.88 \times 10^{-1}$ | | Promoter polymorphism | Moderate | |
| 5 | 23176969 | 67/T | 41.5 ± 1.5 | 84/A | 40.8 ± 1.4 | $7.23 \times 10^{-1}$ | 0.5 | $4.81 \times 10^{-1}$ | 0.1 | $7.48 \times 10^{-1}$ | | Promoter polymorphism | Moderate | |
| 5 | 23177060 | 132/G | 41.4 ± 1.1 | 19/C | 38.6 ± 2.5 | $3.43 \times 10^{-1}$ | 0.9 | $3.53 \times 10^{-1}$ | 1.9 | $1.66 \times 10^{-1}$ | | Promoter polymorphism | Moderate | |
| 5 | 23177120 | 132/G | 41.6 ± 1.1 | 19/C | 37.3 ± 2.1 | $1.53 \times 10^{-1}$ | 1.9 | $1.73 \times 10^{-1}$ | **3.5** | $6.27 \times 10^{-2}$ | | Promoter polymorphism | Moderate | |
| 5 | 23177408 | 99/T | 41.9 ± 1.2 | 52/A | 39.6 ± 1.8 | $2.76 \times 10^{-1}$ | 2.8 | $9.78 \times 10^{-2}$ | 1.9 | $1.74 \times 10^{-1}$ | | Promoter polymorphism | Moderate | |
| 5 | 23177435 | 12/A | 48.1 ± 3 | 139/G | 40.5 ± 1.1 | **$3.79 \times 10^{-2}$** | **7.9** | **$5.72 \times 10^{-3}$** | 2.5 | $1.13 \times 10^{-1}$ | | Promoter polymorphism | Moderate | |
| 5 | 23177477 | 13/C | 47.8 ± 3.5 | 138/A | 40.5 ± 1.1 | **$3.78 \times 10^{-2}$** | **7.5** | **$6.82 \times 10^{-3}$** | 2.7 | $1.05 \times 10^{-1}$ | | Promoter polymorphism | Moderate | |
| 5 | 23177556 | 14/T | 43.8 ± 3.3 | 137/A | 40.8 ± 1.1 | $3.83 \times 10^{-1}$ | 2.2 | $1.40 \times 10^{-1}$ | 0.0 | $8.79 \times 10^{-1}$ | | Promoter polymorphism | Moderate | |
| 5 | 23177675 | 74/T | 41.2 ± 1.4 | 77/G | 41 ± 1.5 | $9.29 \times 10^{-1}$ | 0.1 | $7.76 \times 10^{-1}$ | 0.0 | $1.00 \times 10^{-0}$ | | Promoter polymorphism | Moderate | |
| 5 | 23178864 | 3/C | 55.9 ± 3.9 | 148/A | 40.8 ± 1 | **$3.29 \times 10^{-2}$** | 2.1 | $1.53 \times 10^{-1}$ | 2.1 | $1.50 \times 10^{-1}$ | | Amino acid substitution (Tyr211Ser) | Moderate | |
| 5 | 23179225 | 36/G | 44.8 ± 2.3 | 115/A | 39.9 ± 1.1 | $3.67 \times 10^{-2}$ | **7.5** | **$7.02 \times 10^{-3}$** | 1.3 | $2.56 \times 10^{-1}$ | ✓ | Synonymous polymorphism | Moderate | |
| 5 | 23179561 | 10/T | 46.9 ± 5.2 | 141/G | 40.7 ± 1 | $1.20 \times 10^{-1}$ | 2.5 | $1.17 \times 10^{-1}$ | 0.2 | $6.98 \times 10^{-1}$ | | Amino acid substitution (Gln322His) | Moderate | |
| 5 | 23179695 | 148/T | 41.1 ± 1.1 | 3/C | 41.1 ± 5.7 | $9.98 \times 10^{-1}$ | 0.0 | $9.70 \times 10^{-1}$ | 0.4 | $5.27 \times 10^{-1}$ | | Amino acid substitution (Ile367Thr) | Moderate | |
| 5 | 23179883 | 27/G | 43.9 ± 2.8 | 124/T | 40.5 ± 1.1 | $1.79 \times 10^{-1}$ | 2.6 | $1.08 \times 10^{-1}$ | 0.5 | $4.85 \times 10^{-1}$ | | Amino acid substitution (Ser403Ala) | Moderate | |
| 5 | 23180049 | 7/A | 42.1 ± 4 | 144/G | 41 ± 1.1 | $8.19 \times 10^{-1}$ | 0.1 | $8.01 \times 10^{-1}$ | 0.1 | $7.92 \times 10^{-1}$ | | Amino acid substitution (Glu431Lys) | Moderate | |
| 5 | 23180072 | 5/GAAC | 43 ± 10 | 146/G | 41 ± 1 | $7.20 \times 10^{-1}$ | 0.1 | $7.11 \times 10^{-1}$ | 2.2 | $1.42 \times 10^{-1}$ | | **Amino acid insertion (Asn439 Asn440insAsn)** | Moderate | |
| 5 | 23180305 | 39/C | 44.4 ± 2.1 | 112/G | 39.9 ± 1.2 | **$4.79 \times 10^{-2}$** | **6.7** | **$1.04 \times 10^{-2}$** | 0.8 | $3.63 \times 10^{-1}$ | | Amino acid substitution (Ala494Pro) | Moderate | |
| 5 | 23180338 | 39/A | 44.4 ± 2.1 | 112/G | 39.9 ± 1.2 | **$4.79 \times 10^{-2}$** | **6.7** | **$1.04 \times 10^{-2}$** | 0.8 | $3.63 \times 10^{-1}$ | | Amino acid substitution (Val505Met) | Moderate | |
| 5 | 23180426 | 16/G | 46.1 ± 3.6 | 135/T | 40.5 ± 1.1 | $8.24 \times 10^{-2}$ | **3.4** | $6.91 \times 10^{-2}$ | 1.5 | $2.15 \times 10^{-1}$ | | Amino acid substitution (Met534Arg) | Moderate | |
| 5 | 23180438 | 33/T | 45.3 ± 2.4 | 118/G | 39.9 ± 1.1 | **$2.55 \times 10^{-2}$** | **8.1** | **$5.06 \times 10^{-3}$** | 1.2 | $2.75 \times 10^{-1}$ | | Amino acid substitution (Ser538Ile) | Moderate | |
| 5 | 23180439 | 33/G | 45 ± 2.3 | 118/T | 40 ± 1.1 | **$3.54 \times 10^{-2}$** | **7.1** | **$8.39 \times 10^{-3}$** | 0.8 | $3.64 \times 10^{-1}$ | | Amino acid substitution (Ser538Arg) | Moderate | |
| 5 | 23180482 | 146/T | 41.1 ± 1.1 | 5/C | 40.2 ± 5.5 | $8.74 \times 10^{-1}$ | 0.0 | $8.58 \times 10^{-1}$ | 0.3 | $5.73 \times 10^{-1}$ | | Amino acid substitution (Ser553Pro) | Moderate | |
| 5 | 23180575 | 26/C | 45 ± 2.3 | 125/T | 40.3 ± 1.1 | $7.51 \times 10^{-2}$ | **5.2** | **$2.46 \times 10^{-2}$** | 1.2 | $2.83 \times 10^{-1}$ | | Amino acid substitution (Tyr584His) | Moderate | |
| 5 | 23180582 | 17/A | 42.3 ± 2.9 | 134/C | 40.9 ± 1.1 | $6.66 \times 10^{-1}$ | 0.6 | $4.33 \times 10^{-1}$ | 0.4 | $5.35 \times 10^{-1}$ | | Amino acid substitution (Pro586His) | Moderate | |
| 5 | 23181110 | 134/T | 41.1 ± 1.1 | 17/A | 40.8 ± 2.9 | $9.26 \times 10^{-1}$ | 0.0 | $9.50 \times 10^{-1}$ | 0.2 | $6.35 \times 10^{-1}$ | | Amino acid substitution (Asn731Ile) | Moderate | AT5G57210 |
| 5 | 23181140 | 39/C | 44.4 ± 2.1 | 112/T | 39.9 ± 1.2 | **$4.79 \times 10^{-2}$** | **6.7** | **$1.04 \times 10^{-2}$** | 0.8 | $3.63 \times 10^{-1}$ | | Amino acid substitution (Glu721Gly) | Moderate | |
| 5 | 23181290 | 36/G | 45.5 ± 2.1 | 115/C | 39.7 ± 1.1 | **$1.24 \times 10^{-2}$** | **8.8** | **$3.46 \times 10^{-3}$** | 1.4 | $2.43 \times 10^{-1}$ | ✓ | Synonymous polymorphism | Moderate | |
| 5 | 23181357 | 146/G | 41.1 ± 1.1 | 5/A | 40.2 ± 5.5 | $8.74 \times 10^{-1}$ | 0.0 | $8.58 \times 10^{-1}$ | 0.3 | $5.73 \times 10^{-1}$ | | Amino acid substitution (Pro681Leu) | Moderate | |

| | | | | | | | | | | | | |
|---|---|---|---|---|---|---|---|---|---|---|---|---|
| 5 | 23181367 | 16/T | 47.3 ± 3.3 | 135/TTGA | 40.3 ± 1.1 | **$2.97 \times 10^{-2}$** | **7.9** | **$5.68 \times 10^{-3}$** | 1.3 | $2.57 \times 10^{-1}$ | Amino acid substitution (Ser677del) | Moderate |
| 5 | 23181602 | 6/G | 46.7 ± 5.6 | 145/C | 40.9 ± 1.1 | $2.49 \times 10^{-1}$ | 2.2 | $1.42 \times 10^{-1}$ | 0.1 | $8.12 \times 10^{-1}$ | Amino acid substitution (Glu599Asp) | Moderate |
| 5 | 23181738 | 120/G | 41.4 ± 1.2 | 31/T | 39.7 ± 2.2 | $4.79 \times 10^{-1}$ | 0.0 | $8.29 \times 10^{-1}$ | 0.6 | $4.28 \times 10^{-1}$ | Amino acid substitution (His554Pro) | Moderate |
| 5 | 23181758 | 32/A | 44.9 ± 2.4 | 119/C | 40.1 ± 1.1 | **$4.49 \times 10^{-2}$** | **8.0** | **$5.27 \times 10^{-3}$** | 1.5 | $2.23 \times 10^{-1}$ | Amino acid substitution (Lys547Asn) | Moderate |
| 5 | 23181807 | 15/C | 47.1 ± 3.7 | 136/T | 40.4 ± 1.1 | **$4.47 \times 10^{-2}$** | **4.6** | **$3.45 \times 10^{-2}$** | 2.1 | $1.46 \times 10^{-1}$ | Amino acid substitution (Lys531Arg) | Moderate |
| 5 | 23181853 | 37/T | 44.6 ± 2.2 | 114/A | 40 ± 1.1 | **$4.57 \times 10^{-2}$** | **6.6** | **$1.10 \times 10^{-2}$** | 0.8 | $3.82 \times 10^{-1}$ | Amino acid substitution (Leu516Met) | Moderate |
| 5 | 23181927 | 35/A | 45.4 ± 2.2 | 116/T | 39.8 ± 1.1 | **$1.72 \times 10^{-2}$** | **9.1** | **$3.09 \times 10^{-3}$** | 1.6 | $2.01 \times 10^{-1}$ | Amino acid substitution (Glu491Val) | Moderate |
| 5 | 23182020 | 145/C | 41.1 ± 1.1 | 6/T | 41 ± 4.6 | $9.87 \times 10^{-1}$ | 0.0 | $9.95 \times 10^{-1}$ | 0.1 | $7.65 \times 10^{-1}$ | Amino acid substitution (Arg460His) | Moderate |
| 5 | 23182276 | 37/A | 44.5 ± 2.2 | 114/C | 40 ± 1.1 | $5.02 \times 10^{-2}$ | **6.5** | **$1.19 \times 10^{-2}$** | 0.6 | $4.29 \times 10^{-1}$ | Amino acid substitution (Asp375Tyr) | Moderate |
| 5 | 23183686 | 84/A | 41.6 ± 1.4 | 67/G | 40.4 ± 1.5 | $5.57 \times 10^{-1}$ | 0.2 | $6.67 \times 10^{-1}$ | 1.1 | $2.95 \times 10^{-1}$ | Amino acid substitution (Thr7Ile) | Moderate |
| 5 | 23183936 | 27/T | 43.5 ± 1.8 | 124/C | 40.6 ± 1.2 | $2.62 \times 10^{-1}$ | 1.8 | $1.83 \times 10^{-1}$ | 1.1 | $2.90 \times 10^{-1}$ | 5'-UTR polymorphism | Moderate |
| 5 | 23183969 | 136/CT | 41.5 ± 1.1 | 15/C | 37 ± 3.3 | $1.67 \times 10^{-1}$ | **3.9** | **$4.93 \times 10^{-2}$** | 0.2 | $6.56 \times 10^{-1}$ | 5'-UTR polymorphism | Moderate |
| 5 | 23184013 | 125/G | 42.1 ± 1.1 | 26/A | 36 ± 2.2 | **$1.89 \times 10^{-2}$** | **7.5** | **$7.10 \times 10^{-3}$** | 0.6 | $4.50 \times 10^{-1}$ | 5'-UTR polymorphism | Moderate |
| 5 | 23184017 | 13/A | 45.8 ± 3.2 | 138/T | 40.6 ± 1.1 | $1.44 \times 10^{-1}$ | 2.8 | $9.82 \times 10^{-2}$ | 0.4 | $5.42 \times 10^{-1}$ | 5'-UTR polymorphism | Moderate |
| 5 | 23184047 | 110/A | 42.5 ± 1.2 | 40/T | 37.3 ± 1.8 | **$2.04 \times 10^{-2}$** | **6.3** | **$2.32 \times 10^{-3}$** | 2.3 | $1.08 \times 10^{-1}$ | 5'-UTR polymorphism | Moderate |
| 5 | 23184063 | 101/A | 42.1 ± 1.3 | 48/T | 38.9 ± 1.7 | $1.42 \times 10^{-1}$ | 2.5 | $8.72 \times 10^{-2}$ | 1.9 | $1.60 \times 10^{-1}$ | Promoter polymorphism | Moderate |
| 5 | 23184072 | 94/T | 42.3 ± 1.4 | 57/C | 39.1 ± 1.5 | $1.26 \times 10^{-1}$ | **4.3** | **$4.01 \times 10^{-2}$** | 2.8 | $9.85 \times 10^{-2}$ | Promoter polymorphism | Moderate |
| 5 | 23184074 | 89/T | 42 ± 1.4 | 62/A | 39.8 ± 1.5 | $2.85 \times 10^{-1}$ | 3.0 | $8.78 \times 10^{-2}$ | 0.9 | $3.45 \times 10^{-1}$ | Promoter polymorphism | Moderate |
| 5 | 23184164 | 39/A | 43.8 ± 2 | 112/T | 40.1 ± 1.2 | $1.08 \times 10^{-1}$ | **4.3** | **$3.99 \times 10^{-2}$** | 0.5 | $4.89 \times 10^{-1}$ | Promoter polymorphism | Moderate |
| 5 | 23184170 | 128/A | 41.9 ± 1.1 | 23/T | 36.6 ± 2.3 | $5.53 \times 10^{-2}$ | **4.9** | **$2.87 \times 10^{-2}$** | 0.1 | $7.93 \times 10^{-1}$ | Promoter polymorphism | Moderate |
| 5 | 23184208 | 136/CT | 41.4 ± 1.1 | 15/C | 38.5 ± 3.5 | $3.89 \times 10^{-1}$ | 0.6 | $4.59 \times 10^{-1}$ | 0.1 | $7.61 \times 10^{-1}$ | Promoter polymorphism | Moderate |
| 5 | 23184288 | 34/T | 42.8 ± 2.1 | 116/C | 40.7 ± 1.2 | $3.94 \times 10^{-1}$ | 1.7 | $1.81 \times 10^{-1}$ | 1.0 | $3.67 \times 10^{-1}$ | Promoter polymorphism | Moderate |
| 5 | 23184302 | 85/G | 41.4 ± 1.3 | 66/A | 40.7 ± 1.7 | $7.07 \times 10^{-1}$ | 0.7 | $4.02 \times 10^{-1}$ | 0.5 | $5.01 \times 10^{-1}$ | Promoter polymorphism | Moderate |
| 5 | 23184315 | 126/T | 41.8 ± 1.1 | 25/A | 37.7 ± 2.5 | $1.23 \times 10^{-1}$ | **5.7** | **$1.85 \times 10^{-2}$** | 0.2 | $6.82 \times 10^{-1}$ | Promoter polymorphism | Moderate |
| 5 | 23184363 | 71/T | 41.5 ± 1.5 | 80/A | 40.7 ± 1.4 | $7.10 \times 10^{-1}$ | 0.0 | $9.70 \times 10^{-1}$ | 0.6 | $4.22 \times 10^{-1}$ | Promoter polymorphism | Moderate |
| 5 | 23184393 | 17/T | 44.5 ± 3.9 | 134/C | 40.7 ± 1.1 | $2.24 \times 10^{-1}$ | 2.3 | $1.30 \times 10^{-1}$ | 0.0 | $8.90 \times 10^{-1}$ | Promoter polymorphism | Moderate |
| 5 | 23184449 | 123/C | 41.9 ± 1.1 | 28/G | 37.7 ± 2.3 | $1.02 \times 10^{-1}$ | **6.3** | **$1.31 \times 10^{-2}$** | 0.1 | $7.39 \times 10^{-1}$ | Promoter polymorphism | Moderate |
| 5 | 23184461 | 123/G | 41.8 ± 1.1 | 28/C | 37.9 ± 2.3 | $1.19 \times 10^{-1}$ | **5.7** | **$1.84 \times 10^{-2}$** | 0.0 | $9.04 \times 10^{-1}$ | Promoter polymorphism | Moderate |
| 5 | 23184478 | 124/T | 42 ± 1.1 | 27/C | 36.9 ± 2.4 | **$4.83 \times 10^{-2}$** | **8.7** | **$3.74 \times 10^{-3}$** | 0.5 | $4.95 \times 10^{-1}$ | Promoter polymorphism | Moderate |
| 5 | 23184484 | 128/G | 42 ± 1.1 | 23/A | 35.8 ± 2.4 | **$2.42 \times 10^{-2}$** | **6.7** | **$1.08 \times 10^{-2}$** | 0.0 | $8.83 \times 10^{-1}$ | Promoter polymorphism | Moderate |
| 5 | 23184501 | 135/G | 41.1 ± 1.1 | 16/A | 41 ± 3 | $9.89 \times 10^{-1}$ | 0.1 | $8.11 \times 10^{-1}$ | 0.1 | $7.28 \times 10^{-1}$ | Promoter polymorphism | Moderate |
| 5 | 23184503 | 15/G | 41.8 ± 3 | 136/T | 41 ± 1.1 | $8.04 \times 10^{-1}$ | 0.4 | $5.13 \times 10^{-1}$ | 0.1 | $8.07 \times 10^{-1}$ | Promoter polymorphism | Moderate |
| 5 | 23184506 | 111/C | 42.3 ± 1.2 | 40/T | 37.7 ± 1.8 | **$4.21 \times 10^{-2}$** | **4.9** | **$2.81 \times 10^{-2}$** | 1.8 | $1.86 \times 10^{-1}$ | Promoter polymorphism | Moderate |
| 5 | 23184510 | 136/A | 41.1 ± 1.1 | 15/G | 40.6 ± 3.1 | $8.67 \times 10^{-1}$ | 0.0 | $9.92 \times 10^{-1}$ | 1.4 | $2.41 \times 10^{-1}$ | Promoter polymorphism | Moderate |
| 5 | 23184517 | 20/T | 42.7 ± 2.5 | 131/C | 40.8 ± 1.1 | $5.39 \times 10^{-1}$ | 0.7 | $3.98 \times 10^{-1}$ | 0.0 | $9.01 \times 10^{-1}$ | Promoter polymorphism | Moderate |
| 5 | 23184573 | 31/T | 45.8 ± 2.4 | 120/A | 39.9 ± 1.1 | $1.53 \times 10^{-2}$ | **11.7** | **$8.01 \times 10^{-4}$** | **6.6** | **$1.10 \times 10^{-2}$** | Promoter polymorphism | Moderate |
| 5 | 23184635 | 39/G | 44.5 ± 2.1 | 112/C | 39.9 ± 1.2 | **$4.17 \times 10^{-2}$** | **6.7** | **$1.08 \times 10^{-2}$** | 0.9 | $3.57 \times 10^{-1}$ | Promoter polymorphism | Moderate |
| 5 | 23184664 | 140/T | 41.1 ± 1.1 | 11/A | 40.9 ± 3.9 | $9.54 \times 10^{-1}$ | 0.0 | $9.98 \times 10^{-1}$ | 0.2 | $6.46 \times 10^{-1}$ | Promoter polymorphism | Moderate |
| 5 | 23184863 | 53/A | 43.7 ± 1.7 | 98/G | 39.7 ± 1.3 | $5.07 \times 10^{-2}$ | **4.2** | **$4.15 \times 10^{-2}$** | 1.4 | $2.46 \times 10^{-1}$ | Promoter polymorphism | Moderate |

The results of a local association study (LAS) of the variation in relative root length (RRL) in 151 *A. thaliana* ecotypes, in pH 8.0 versus pH 5.8 media, separately for individual genomic regions, are presented. The polymorphisms in coding sequences, 5'-UTRs, and promoters within 1 kb from the start codons of genes within haploblocks of the 20 most significant single-nucleotide polymorphisms (SNPs), detected by a genome-wide association study (GWAS) (see Table 1), were downloaded from the 1001 genomes database (https://tools.1001genomes.org/polymorph/). Data from only 151 out of the 218 ecotypes used for the GWAS (see Figure 1, Table 1) were available in the 1001 genomes database. The frequency of the tolerant and sensitive alleles among the 151 ecotypes is shown with the average RRL for each. The *F*-statistic and *P*-values of a Student's *t*-test, generalized linear model (GLM), as well as the mixed linear model (GLM) methods of LAS calculations are shown.

#### Table S6. Amino acid haplotypes associated with alkalinity tolerance in 151 *Arabidopsis thaliana* ecotypes

| Haplotype | Representative ecotype | Number of ecotypes | RRL | Amino acid haplotypes | | | | | | | | | | | | | |
|---|---|---|---|---|---|---|---|---|---|---|---|---|---|---|---|---|---|
| **AT1G02080.1 (*NOT1*)** | | | | **51** | **53** | **270** | **317** | **403** | **428** | **597** | **751** | **1215** | **1265** | **1616** | **1904** | **2019** | **2047** |
| **H1** | Sij-2 | 1 | 12.9 | Thr | Leu | Asp | Ala | Phe | Met | Ser | Glu | Thr | Ser | Ser | Glu | Val | Trp |
| **H2** | Bsch-0 | 1 | 23.3 | Pro | Leu | Asp | Ala | Phe | Ile | Ser | Glu | Ile | Ser | Thr | Gln | Asp | Trp |
| **H3** | In-0 | 1 | 31.1 | Thr | Leu | Asp | Ala | Phe | Met | Ser | Glu | Ile | Ser | Ser | Gln | Val | Trp |
| **H4** | Timpo-1 | 1 | 31.4 | Thr | Leu | Asp | Val | Phe | Met | Ser | Glu | Ile | Ser | Ser | Gln | Val | Trp |
| **H5** | Ciste-2 | 1 | 31.7 | Thr | Leu | Asp | Val | Phe | Met | Ser | Glu | Ile | Ser | Thr | Gln | Val | Trp |
| **H6** | Sij-1 | 1 | 33.3 | Pro | Leu | Asp | Ala | Phe | Met | Ser | Glu | Thr | Thr | Ser | Glu | Asp | Trp |
| **H7** | Borsk-2 | 1 | 35.6 | Pro | Leu | Asp | Ala | Phe | Met | Ser | Glu | Thr | Ser | Ser | Gln | Asp | Trp |
| **H8** | Yeg-1 | 1 | 40 | Thr | Leu | Asp | Val | Phe | Met | Ser | Glu | Ile | Ser | Ser | Gln | Asp | Trp |
| **H9** | Ms-0 | 1 | 51.2 | Thr | Leu | Asp | Ala | Phe | Met | Ser | Glu | Thr | Ser | Ser | Gln | Val | Trp |
| **H10** | stepn-2 | 1 | 51.4 | Thr | Phe | Asp | Ala | Phe | Met | Ser | Glu | Ile | Ser | Ser | Gln | Val | Trp |
| **H11** | Mnz-0 | 1 | 51.5 | Pro | Leu | Asn | Ala | Phe | Met | Ser | Glu | Ile | Ser | Ser | Gln | Asp | Leu |
| **H12** | Alst-1 | 2 | 31.9 ± 2.6 | Pro | Leu | Asn | Ala | Phe | Met | Ser | Glu | Ile | Ser | Ser | Gln | Val | Leu |
| **H13** | Se-0 | 2 | 32.2 ± 2.4 | Thr | Leu | Asp | Ala | Phe | Met | Ser | Glu | Ile | Ser | Thr | Gln | Val | Trp |
| **H14** | Sorbo | 2 | 36.9 ± 10.4 | Thr | Leu | Asp | Ala | Phe | Met | Ser | Glu | Thr | Thr | Ser | Glu | Val | Trp |
| **H15** | Buckhorn Pass | 55 | 38.6 ± 1.6 | Thr | Leu | Asp | Ala | Phe | Met | Ser | Glu | Ile | Ser | Ser | Gln | Val | Trp |
| **H16** | Lag2-2 | 2 | 40.6 ± 0.6 | Pro | Leu | Asp | Ala | Phe | Met | Ser | Glu | Ile | Ser | Ser | Gln | Asp | Trp |
| **H17** | Lago-1 | 8 | 40.9 ± 3.2 | Pro | Leu | Asp | Ala | Phe | Met | Ser | Glu | Ile | Ser | Ser | Gln | Val | Trp |
| **H18** | Ru3.1-31 | 13 | 42.8 ± 4 | Thr | Leu | Asp | Ala | Phe | Met | Ser | Glu | Ile | Ser | Ser | Gln | Asp | Trp |
| **H19** | Sei-0 | 6 | 43.4 ± 3.6 | Thr | Phe | Asp | Ala | Val | Met | Cys | Asp | Ile | Ser | Ser | Gln | Val | Trp |
| **H20** | Col-0 | 15 | 44.6 ± 3.6 | Pro | Leu | Asp | Ala | Phe | Met | Ser | Glu | Ile | Ser | Thr | Gln | Asp | Trp |
| **H21** | Hh-0 | 30 | 45 ± 2.3 | Pro | Leu | Asp | Ala | Phe | Met | Ser | Glu | Ile | Ser | Thr | Gln | Val | Trp |
| **H22** | Sha | 2 | 47.5 ± 13.5 | Thr | Leu | Asp | Ala | Phe | Met | Ser | Glu | Thr | Thr | Ser | Glu | Asp | Trp |
| **H23** | Petro-1 | 3 | 48.6 ± 9.4 | Pro | Leu | Asp | Ala | Phe | Ile | Ser | Glu | Ile | Ser | Thr | Gln | Val | Trp |
| | | | | **ring→ open-chain** | **Aliphatic→Aromatic** | **Negative→Uncharged** | | **Aromatic→Aliphatic** | | | | **Non-polar→Polar** | | | **Uncharged→Negative** | **Non-polar→Negative** | **Aromatic→Aliphatic** |

| Haplotype | Representative ecotype | Number of ecotypes | RRL | | | | | | | | | | |
|---|---|---|---|---|---|---|---|---|---|---|---|---|---|
| **AT1G05630.1 (*5PTASE13*)** | | | | **9** | **19** | **199** | **246** | **481** | **969** | **976** | **1060** | **1074** | **1140** |
| **H1** | Sij-2 | 5 | 31.2 ± 5.4 | Glu | Pro | Ala | Asp | Val | Ile | Ser | Pro | Asn | Leu |
| **H2** | Si-0 | 3 | 33.2 ± 5.7 | Glu | Pro | Ala | Asp | Val | Leu | Leu | Pro | Asn | Leu |
| **H3** | Do-0 | 5 | 35.5 ± 3.6 | Glu | Pro | Ala | Asp | Val | Ile | Leu | Gln | Asn | Leu |
| **H4** | Aa-0 | 7 | 35.5 ± 5.9 | Glu | Ser | Ala | Asp | Val | Ile | Leu | Pro | Asn | Leu |
| **H5** | Sij-1 | 3 | 37.4 ± 3.8 | Glu | Pro | Ala | Asp | Ile | Ile | Ser | Pro | Asn | Leu |
| **H6** | Ba-1 | 3 | 38 ± 8.5 | Glu | Pro | Ala | Asp | Val | Ile | Leu | Pro | Asn | Ile |
| **H7** | Is-0 | 94 | 41.7 ± 1.3 | Glu | Pro | Ala | Asp | Val | Ile | Leu | Pro | Asn | Leu |
| **H8** | Col-0 | 5 | 42.4 ± 3.7 | Glu | Pro | Ala | Asp | Val | Ile | Leu | Pro | Tyr | Leu |
| **H9** | Buckhorn Pass | 19 | 42.8 ± 3.5 | Asp | Pro | Ala | Asp | Val | Ile | Leu | Pro | Asn | Leu |
| **H10** | Apost-1 | 3 | 48.4 ± 2.9 | Glu | Pro | Ala | Glu | Val | Ile | Leu | Pro | Asn | Leu |
| **H11** | Ha-0 | 4 | 52.2 ± 4.5 | Glu | Pro | Val | Asp | Val | Ile | Leu | Pro | Asn | Leu |
| | | | | | | | | | | **Non-polar→Polar** | | **Non-polar→Polar** | |

| AT1G24430.1 | | | | 2 | 7 | 17 | 23 | 36 | 73 | 88 | 282 |
|---|---|---|---|---|---|---|---|---|---|---|---|
| **H1** | Rovero-1 | 1 | 24.3 | Glu | Val | Pro | Pro | Ser | Phe | Met | Glu |
| **H2** | Slavi-1 | 1 | 29.5 | Glu | Ile | Ser | Leu | Ser | Tyr | Met | Glu |
| **H3** | Gel-1 | 1 | 32.4 | Glu | Val | Pro | Leu | Ser | Phe | Met | Glu |
| **H4** | Ba-1 | 3 | 32.9 ± 7.1 | Glu | Val | Ser | Leu | Ser | Phe | Met | Asp |
| **H5** | Me-0 | 1 | 33.2 | Gly | Val | Ser | Leu | Ser | Tyr | Met | Glu |
| **H6** | Sij-1 | 2 | 33.7 ± 0.4 | Glu | Ile | Pro | Leu | Ser | Phe | Met | Glu |
| **H7** | Sg-1 | 1 | 34 | Gly | Val | Pro | Leu | Ser | Phe | Met | Glu |
| **H8** | Or-0 | 1 | 34.9 | Glu | Ile | Pro | Leu | Ser | Phe | Met | Asp |
| **H9** | Tu-SB30-3 | 1 | 36.9 | Gly | Ile | Ser | Leu | Ala | Phe | Met | Glu |
| **H10** | In-0 | 2 | 36.9 ± 5.9 | Glu | Val | Ser | Leu | Ser | Phe | Met | Glu |
| **H11** | Ru3.1-31 | 3 | 38.4 ± 10.3 | Gly | Ile | Ser | Leu | Ser | Phe | Met | Glu |
| **H12** | Boot-1 | 1 | 39.7 | Gly | Val | Pro | Pro | Ser | Phe | Met | Asp |
| **H13** | Sij-2 | 55 | 39.7 ± 1.7 | Glu | Ile | Ser | Leu | Ser | Phe | Met | Glu |
| **H14** | Lag2-2 | 1 | 40 | Glu | Ile | Ser | Leu | Ser | Tyr | Met | Asp |
| **H15** | Buckhorn Pass | 48 | 41.1 ± 1.9 | Glu | Ile | Ser | Leu | Ser | Phe | Met | Asp |
| **H16** | Ven-1 | 3 | 42.6 ± 2.9 | Glu | Ile | Ser | Leu | Ser | Tyr | Val | Asp |
| **H17** | Rd-0 | 5 | 43.4 ± 3.8 | Gly | Val | Ser | Leu | Ser | Phe | Met | Glu |
| **H18** | Mir-0 | 1 | 44.8 | Gly | Val | Pro | Pro | Ser | Phe | Met | Glu |
| **H19** | Col-0 | 3 | 45.6 ± 0.5 | Gly | Val | Pro | Pro | Ala | Tyr | Val | Asp |
| **H20** | Nok-3 | 2 | 45.8 ± 4.9 | Gly | Val | Ser | Leu | Ser | Phe | Met | Asp |
| **H21** | Ciste-1 | 1 | 45.9 | Glu | Ile | Ser | Pro | Ser | Phe | Met | Glu |
| **H22** | Apost-1 | 4 | 47.5 ± 2.6 | Gly | Ile | Ser | Leu | Ser | Phe | Met | Asp |
| **H23** | Np-0 | 7 | 50.4 ± 0.3 | Gly | Val | Pro | Pro | Ala | Tyr | Val | Asp |
| **H24** | Mer-6 | 1 | 53.7 | Glu | Ile | Ser | Leu | Ala | Phe | Met | Asp |
| **H25** | Gie-0 | 1 | 61.3 | Gly | Val | Pro | Leu | Ser | Phe | Met | Asp |
| **H26** | Castelfed-4 | 1 | 66.8 | Glu | Ile | Ser | Pro | Ser | Phe | Met | Asp |
| | | | | **Non-polar→ Negative** | | | **Polar→ Non-polar** | **Non-polar→ Polar** | | | |

| AT1G75050.1 | | | | 164 |
|---|---|---|---|---|
| H1 | Aa-0 | 24 | 36.8 ± 2.4 | Asp |
| H2 | Col-0 | 127 | 41.9 ± 1.1 | Glu |

| AT2G40550.1 (*ETG1*) | | | | 44-588 MCM Complex | | | |
|---|---|---|---|---|---|---|---|
| | | | | 183 | 213 | 274 | 560 |
| H1 | Sp-0 | 1 | 33.9 | Ser | Ser | Glu | Met |
| H2 | Si-0 | 3 | 36.2 ± 8.4 | Cys | Ser | Gly | Met |
| H3 | Kz-9 | 7 | 39.4 ± 3.6 | Ser | Cys | Gly | Met |
| H4 | Uod-1 | 19 | 39.8 ± 2.1 | Ser | Ser | Glu | Ile |
| H5 | Col-0 | 120 | 41.4 ± 1.2 | Ser | Ser | Gly | Met |
| H6 | Bor-1 | 1 | 63.5 | Ser | Ser | Gly | Ile |

| AT3G17570.1 | | | | | | | | | | | | | | |
|---|---|---|---|---|---|---|---|---|---|---|---|---|---|---|
| | | | | | | **Uncharged→Negative** | | | | | | | | |
| | | | | | | **Kelch-1 motif** | | | | 229-281 Kelch-2 motif | | | 331-379 Kelch-3 motif | |
| | | | | | | 151-199 F-Box domain | | | | | | | | |
| | | | | 46 | 65 | 100 | 163 | 169 | 195 | 230 | 239 | 259 | 344* | 351 |
| H1 | Kz-9 | 1 | 23.2 | Val | Ser | Gly | Asp | Glu | | | | | | |
| H2 | Timpo-1 | 1 | 31.4 | Met | Ser | Gly | Asp | Glu | Val | Leu | Asn | Glu | Tyr | Trp |
| H3 | Do-0 | 4 | 36.5 ± 5.6 | Val | Ser | Gly | Asp | Glu | Val | Pro | Asn | Glu | Tyr | Trp |
| H4 | Col-0 | 61 | 38.8 ± 1.4 | Val | Ser | Gly | Asp | Glu | Val | Leu | Asn | Glu | Tyr | Trp |
| H5 | Wa-1 | 3 | 38.8 ± 2.6 | Val | Ser | Cys | Asp | Glu | Val | Leu | Asn | Glu | Tyr | Trp |
| H6 | Rovero-1 | 2 | 39.4 ± 15.1 | Val | Ser | Gly | Asp | Glu | Val | Leu | Asn | Gly | Tyr | Trp |
| H7 | CIBC-5 | 18 | 39.8 ± 2.8 | Met | Ser | Gly | Asp | Glu | Val | Leu | Asn | Gly | Tyr | Trp |
| H8 | Sij-2 | 39 | 41.3 ± 2 | Val | Ser | Gly | Asp | Glu | Val | Pro | Asn | Glu | Tyr | Leu |
| H9 | Buckhorn Pass | 6 | 47.5 ± 8 | Val | Ser | Gly | Asp | Glu | Val | Pro | Asp | Glu | Tyr | Leu |
| H10 | Kin-0 | 5 | 48.4 ± 5.9 | Val | Ser | Gly | Asn | Glu | Val | Pro | Asn | Glu | * | |
| H11 | Ct-1 | 3 | 49.9 ± 2.5 | Val | Ser | | | | | | | | | |
| H12 | HKT2-4 | 4 | 50 ± 5.6 | Val | Ser | Gly | Asp | Glu | Val | Leu | Asn | Glu | Tyr | Leu |
| H13 | Gy-0 | 1 | 53.1 | Val | Ser | | | | | | | | | |
| H14 | TuWa1-2 | 1 | 53.8 | Val | Ser | Gly | Asp | Glu | Val | Pro | Asn | Glu | * | |
| H15 | Fei-0 | 1 | 63.4 | Val | Ser | Gly | Asn | Glu | Val | Pro | Asn | Glu | Tyr | Trp |
| H16 | Castelfed-4 | 1 | 66.8 | Val | Ser | Gly | Asp | Glu | Val | Pro | Asp | Glu | Tyr | Trp |
| | | | | | **frameshift** | **Non-polar→Polar** | **Negative→Uncharged** | **frameshift** | | **Non-polar→Polar** | **Uncharged→Negative** | **Polar→Non-polar** | | **Aromatic→Aliphatic** |

| | | | | | | | 19-366 SGNH HYDROLASE Domain | | | | | | | | | | | | | | | | | | | | | | | | |
|---|---|---|---|---|---|---|---|---|---|---|---|---|---|---|---|---|---|---|---|---|---|---|---|---|---|---|---|---|---|---|---|
| AT3G26430.1 (*GGL20*) | | | | 6 | 7 | 8 | 22 | 45 | 48 | 97 | 100 | 146 | 186 | 188 | 195 | 197 | 200 | 203 | 205 | 236 | 243 | 253 | 261 | 283 | 284 | 298 | 334 | 354 | 360 | 361 | 364 |
| **H1** | Sij-2 | 1 | 12.9 | Leu | Leu | Ile | Ala | Leu | Ala | Asn | His | Arg | Ser | Gln | Asn | His | Leu | Val | Arg | Pro | His | Ala | Lys | Ser | Ile | Arg | Lys | Ala | Gln | Gln | Asp |
| **H2** | Is-0 | 1 | 14.9 | Met | Leu | Ile | Ala | Leu | Ala | Asn | His | Arg | Ser | Gln | Asp | His | Phe | Val | Arg | Pro | His | Ser | Arg | Ser | Ile | Arg | Lys | Ala | Gln | Gln | Asp |
| **H3** | Buckhorn Pass | 1 | 15.2 | Leu | Leu | Val | Ala | Leu | Ala | Asn | His | Arg | Ser | Gln | Asp | His | Phe | Val | Arg | Leu | Tyr | Ala | Arg | Ser | Ile | Arg | Lys | Ala | Gln | Gln | Gly |
| **H4** | Ru3.1-31 | 1 | 18.3 | Leu | Leu | Val | Ala | Leu | Ala | Asn | His | Arg | Ser | Gln | Asp | His | Leu | Val | Arg | Pro | His | Ala | Lys | Ser | Ile | Arg | Lys | Ala | Gln | Gln | Asp |
| **H5** | Ba-1 | 1 | 21.2 | Leu | Leu | Val | Ala | Leu | Ala | Asn | His | Arg | Ser | Gln | Asp | His | Leu | Val | Arg | Leu | His | Ala | Arg | Ser | Ile | Arg | Lys | Ala | Gln | Lys | Gly |
| **H6** | Kz-9 | 1 | 23.2 | Leu | Leu | Val | Ala | Leu | Ser | Asn | His | Arg | Thr | Gln | Asn | Leu | Phe | Ala | Arg | Pro | His | Ala | Arg | Ser | Ile | Arg | Lys | Ala | Gln | Gln | Gly |
| **H7** | Bsch-0 | 1 | 23.3 | Leu | Leu | Val | Ala | Leu | Ala | Asn | Asn | Arg | Thr | Gln | Asp | Leu | Phe | Ala | Arg | Pro | His | Ala | Arg | Ser | Ile | Arg | Asn | Ala | Gln | Lys | Gly |
| **H8** | Rovero-1 | 1 | 24.3 | Leu | Leu | Val | Ala | Leu | Ser | Asn | His | Arg | Ser | Gln | Asp | Leu | Leu | Val | Cys | Pro | His | Ala | Lys | Ser | Ile | Arg | Lys | Ala | Gln | Gln | Asp |
| **H9** | Sorbo | 1 | 26.6 | Leu | Leu | Val | Ala | Leu | Ser | Asn | His | Arg | Thr | Gln | Asn | Leu | Phe | Ala | Arg | Pro | His | Ala | Arg | Ser | Ile | Arg | Lys | Ala | Gln | Gln | Asp |
| **H10** | Star-8 | 1 | 27.3 | Leu | Leu | Val | Ala | Leu | Ala | Asn | His | Arg | Thr | Gln | Asn | Leu | Leu | Val | Arg | Pro | His | Ala | Arg | Tyr | Leu | Arg | Lys | Ala | Gln | Gln | Asp |
| **H11** | Pn-0 | 1 | 28.5 | Met | Leu | Ile | Ala | Leu | Ala | Asn | His | Arg | Ser | Gln | Asp | His | Leu | Ala | Arg | Leu | Tyr | Ala | Arg | Ser | Ile | Arg | Lys | Ala | Gln | Gln | Gly |
| **H12** | Alst-1 | 1 | 29.3 | Leu | Leu | Val | Ala | Leu | Ala | Asn | His | Arg | Ser | Gln | Asn | Leu | Phe | Ala | Arg | Leu | Tyr | Ala | Arg | Ser | Ile | Arg | Asn | Ala | Pro | Lys | Gly |
| **H13** | Ciste-2 | 1 | 31.7 | Leu | Leu | Val | Ala | Leu | Ser | Ile | His | Arg | Ser | Gln | Asp | Leu | Leu | Val | Arg | Pro | His | Ala | Lys | Ser | Ile | Arg | Lys | Ala | Gln | Gln | Asp |
| **H14** | Sij-1 | 1 | 33.3 | Leu | Leu | Ile | Ala | Leu | Ser | Asn | His | Arg | Thr | Gln | Asn | His | Leu | Val | Arg | Pro | His | Ala | Arg | Ser | Ile | Arg | Lys | Thr | Gln | Gln | Asp |
| **H15** | Sp-0 | 1 | 33.9 | Met | Phe | Ile | Ala | Leu | Ser | Asn | Asn | Arg | Thr | Gln | Asn | Leu | Phe | Ala | Arg | Pro | His | Ala | Arg | Ser | Ile | Arg | Asn | Ala | Gln | Lys | Gly |
| **H16** | Np-0 | 1 | 33.9 | Leu | Leu | Val | Ala | Leu | Ser | Asn | His | Arg | Ser | Gln | Asp | His | Leu | Val | Arg | Pro | His | Ala | Lys | Ser | Ile | Arg | Lys | Ala | Gln | Gln | Asp |
| **H17** | wt-5 | 1 | 33 | Leu | Leu | Val | Ala | Leu | Ala | Asn | His | Arg | Ser | Gln | Asp | His | Phe | Ala | Arg | Leu | Tyr | Ala | Arg | Ser | Ile | Arg | Lys | Ala | Gln | Gln | Gly |
| **H18** | El-0 | 1 | 34.4 | Met | Leu | Ile | Ala | Leu | Ala | Asn | His | Arg | Ser | Gln | Asn | Leu | Phe | Ala | Arg | Leu | Tyr | Ala | Arg | Ser | Ile | Arg | Asn | Ala | Gln | Lys | Gly |
| **H19** | Bolin-1 | 1 | 34.5 | Leu | Leu | Ile | Ala | Leu | Ala | Asn | His | Arg | Ser | Gln | Asn | Leu | Leu | Val | Arg | Leu | Tyr | Ala | Arg | Ser | Ile | Arg | Asn | Thr | Gln | Gln | Asp |
| **H20** | Vezzano-2 | 1 | 34.7 | Met | Leu | Ile | Ala | Leu | Ala | Asn | His | Arg | Ser | Gln | Asp | His | Leu | Val | Arg | Leu | Tyr | Ala | Arg | Ser | Ile | Ser | Asn | Thr | Gln | Gln | Asp |
| **H21** | Gel-1 | 2 | 34.7 ± 2.3 | Leu | Leu | Val | Ala | Leu | Ala | Asn | His | Arg | Ser | Gln | Met | Leu | Ser | Pro | Val | | | | | | | | | | | | |
| **H22** | Or-0 | 1 | 34.9 | Leu | Leu | Val | Ala | Leu | Ala | Asn | His | Arg | Ser | Gln | Asn | His | Phe | Ala | Arg | Leu | Tyr | Ala | Arg | Ser | Ile | Arg | Lys | Ala | Gln | Gln | Gly |
| **H23** | Sha | 1 | 34 | Met | Leu | Ile | Ala | Leu | Ala | Asn | His | Arg | Thr | Gln | Asp | His | Leu | Ala | Arg | Pro | His | Ala | Arg | Ser | Ile | Arg | Lys | Thr | Gln | Gln | Asp |
| **H24** | Borsk-2 | 1 | 35.6 | Leu | Leu | Val | Ala | Leu | Ser | Asn | His | Arg | Thr | Gln | Asn | His | Leu | Val | Arg | Pro | His | Ala | Lys | Ser | Leu | Arg | Lys | Ala | Gln | Gln | Asp |
| **H25** | Ts-1 | 1 | 35.7 | Leu | Leu | Val | Ala | Leu | Ala | Ile | His | Arg | Ser | Gln | Asp | Leu | Leu | Val | Arg | Pro | His | Ala | Lys | Ser | Ile | Arg | Lys | Thr | Gln | Gln | Asp |
| **H26** | Uod-7 | 1 | 35.9 | Met | Leu | Ile | Ala | Leu | Ser | Asn | His | Arg | Ser | Gln | Asn | Leu | Phe | Ala | Arg | Leu | Tyr | Ala | Arg | Ser | Ile | Ser | Asn | Ala | Pro | Lys | Gly |
| **H27** | Baa-1 | 9 | 35.9 ± 3.7 | Met | Leu | Ile | Ala | Leu | Ala | Asn | His | Arg | Ser | Gln | Asn | Leu | Phe | Ala | Arg | Leu | Tyr | Ala | Arg | Ser | Ile | Arg | Lys | Ala | Gln | Gln | Gly |
| **H28** | Rd-0 | 1 | 35 | Leu | Leu | Val | Ala | Ile | Ala | Asn | His | Arg | Ser | Gln | Asp | His | Phe | Ala | Arg | Pro | His | Ala | Arg | Tyr | Leu | Arg | Lys | Ala | Gln | Gln | Asp |
| **H29** | NFA-8 | 1 | 36.6 | Met | Leu | Ile | Ala | Leu | Ala | Asn | His | Arg | Ser | Gln | Asn | Leu | Leu | Val | Arg | Leu | Tyr | Ala | Arg | Ser | Ile | Arg | Asn | Ala | Pro | Lys | Gly |
| **H30** | Pog-0 | 2 | 36.6 ± 1.9 | Leu | Leu | Ile | Ala | Leu | Ala | Asn | His | Arg | Ser | Gln | Asn | Leu | Phe | Ala | Arg | Leu | Tyr | Ala | Arg | Ser | Ile | Arg | Lys | Ala | Gln | Gln | Gly |
| **H31** | Tu-SB30-3 | 1 | 36.9 | Leu | Leu | Val | Ala | Leu | Ala | Asn | His | Arg | Thr | Gln | Asn | Leu | Leu | Ala | Arg | Pro | His | Ala | Arg | Tyr | Leu | Arg | Lys | Ala | Gln | Gln | Asp |
| **H32** | HKT2-4 | 1 | 37.4 | Leu | Leu | Ile | Ala | Leu | Ser | Asn | His | Arg | Ser | Gln | Asn | Leu | Phe | Val | Arg | Pro | His | Ala | Lys | Ser | Ile | Arg | Lys | Thr | Gln | Gln | Asp |
| **H33** | Col-0 | 4 | 37 ± 6 | Leu | Leu | Val | Ala | Leu | Ser | Asn | His | Arg | Ser | Gln | Asp | His | Leu | Val | Arg | Pro | His | Ala | Lys | Ser | Ile | Arg | Lys | Thr | Gln | Gln | Asp |
| **H34** | Da-0 | 2 | 38.7 ± 2.9 | Met | Leu | Ile | Ala | Leu | Ser | Asn | His | Arg | Ser | Gln | Asn | Leu | Phe | Ala | Arg | Leu | Tyr | Ala | Arg | Ser | Ile | Arg | Lys | Ala | Gln | Gln | Gly |
| **H35** | Aa-0 | 34 | 38.8 ± 1.8 | Leu | Leu | Val | Ala | Leu | Ala | Asn | His | Arg | Ser | Gln | Asp | Leu | Leu | Val | Arg | Pro | His | Ala | Lys | Ser | Ile | Arg | Lys | Ala | Gln | Gln | Asp |
| **H36** | Wl-0 | 1 | 38 | Leu | Leu | Val | Ala | Leu | Ala | Asn | His | Arg | Ser | Gln | Asn | Leu | Phe | Ala | Arg | Pro | His | Ala | Lys | Ser | Ile | Arg | Lys | Ala | Gln | Gln | Asp |
| **H37** | Hh-0 | 5 | 40.3 ± 6.4 | Leu | Leu | Val | Ala | Leu | Ser | Asn | His | Arg | Ser | Gln | Asp | Leu | Leu | Val | Arg | Pro | His | Ala | Lys | Ser | Ile | Arg | Lys | Ala | Gln | Gln | Asp |
| **H38** | Wa-1 | 2 | 40.6 ± 6.7 | Met | Leu | Ile | Ala | Leu | Ala | Asn | His | Arg | Thr | Gln | Asn | Leu | Phe | Ala | Arg | Pro | His | Ala | Arg | Ser | Ile | Arg | Lys | Ala | Gln | Gln | Asp |
| **H39** | Ag-0 | 1 | 40.8 | Leu | Leu | Val | Ala | Leu | Ser | Asn | His | Arg | Ser | Gln | Asp | His | Leu | Ala | Arg | Leu | Tyr | Ala | Arg | Ser | Ile | Arg | Lys | Thr | Gln | Gln | Asp |
| **H40** | Si-0 | 3 | 40.8 ± 9.3 | Leu | Leu | Val | Val | Leu | Ala | Asn | His | Arg | Ser | Gln | Asn | Leu | Phe | Ala | Arg | Leu | Tyr | Ala | Arg | Ser | Ile | Arg | Lys | Ala | Gln | Gln | Gly |
| **H41** | Nok-3 | 1 | 40.9 | Met | Leu | Ile | Ala | Leu | Ala | Asn | His | Arg | Ser | Gln | Asp | His | Leu | Val | Arg | Leu | Tyr | Ala | Arg | Ser | Ile | Arg | Lys | Ala | Gln | Lys | Gly |
| **H42** | Lag2-2 | 1 | 40 | Leu | Phe | Ile | Ala | Leu | Ser | Asn | His | Arg | Thr | Gln | Asp | His | Phe | Ala | Arg | Pro | His | Ala | Lys | Ser | Ile | Arg | Lys | Ala | Gln | Gln | Asp |
| **H43** | Aitba 2 | 1 | 41.1 | Leu | Leu | Ile | Ala | Leu | Ala | Asn | His | Arg | Ser | Gln | Asn | Leu | Leu | Val | Arg | Leu | Tyr | Ala | Arg | Ser | Ile | Arg | Asn | Ala | Gln | Gln | Gly |
| **H44** | Lago-1 | 5 | 41.6 ± 4.3 | Leu | Leu | Val | Ala | Leu | Ala | Ile | His | Arg | Ser | Gln | Asp | Leu | Leu | Val | Arg | Pro | His | Ala | Lys | Ser | Ile | Arg | Lys | Ala | Gln | Gln | Asp |
| **H45** | Nie1-2 | 1 | 42.1 | Leu | Leu | Val | Ala | Leu | Ser | Asn | His | Arg | Ser | Gln | Asn | His | Leu | Val | Arg | Pro | His | Ala | Lys | Ser | Ile | Arg | Lys | Thr | Gln | Gln | Asp |
| **H46** | Mh-0 | 1 | 42.5 | Leu | Leu | Val | Val | Leu | Ser | Asn | His | Arg | Ser | Gln | Asn | Leu | Phe | Ala | Arg | Leu | His | Ala | Lys | Ser | Ile | Arg | Lys | Ala | Gln | Gln | Gly |
| **H47** | Ove-0 | 1 | 42.6 | Met | Leu | Ile | Ala | Leu | Ala | Asn | His | Arg | Ser | Gln | Asp | His | Phe | Ala | Arg | Leu | Tyr | Ala | Arg | Ser | Ile | Arg | Lys | Ala | Gln | Gln | Gly |
| **H48** | Stepn-1 | 1 | 42.8 | Leu | Phe | Ile | Ala | Leu | Ala | Asn | His | Ala | | | | | | | | | | | | | | | | | | | |
| **H49** | Kelsterbach-4 | 1 | 42 | Met | Leu | Ile | Val | Leu | Ala | Asn | His | Arg | Ser | Gln | Asn | Leu | Phe | Ala | Arg | Leu | Tyr | Ala | Arg | Ser | Ile | Arg | Lys | Ala | Gln | Gln | Gly |
| **H50** | Gr-1 | 2 | 43.7 ± 1.2 | Met | Phe | Ile | Ala | Leu | Ala | Asn | Asn | Arg | Thr | Gln | Asn | Leu | Phe | Ala | Arg | Pro | His | Ala | Arg | Ser | Ile | Arg | Asn | Ala | Gln | Lys | Gly |
| **H51** | In-0 | 2 | 44.7 ± 13.6 | Met | Leu | Ile | Ala | Leu | Ala | Asn | His | Arg | Thr | Gln | Asn | Leu | Phe | Ala | Arg | Pro | His | Ser | Arg | Ser | Ile | Arg | Lys | Ala | Gln | Gln | Asp |
| **H52** | Ga-0 | 3 | 44.7 ± 3.6 | Leu | Leu | Val | Ala | Leu | Ala | Asn | His | Arg | Ser | Gln | Asp | Leu | Leu | Val | Cys | Pro | His | Ala | Lys | Ser | Ile | Arg | Lys | Ala | Gln | Gln | Asp |
| **H53** | NFA-10 | 1 | 45.1 | Met | Leu | Ile | Ala | Leu | Ala | Asn | His | Arg | Thr | Gln | Asn | Leu | Phe | Val | Arg | Pro | His | Ser | Arg | Ser | Ile | Arg | Lys | Ala | Gln | Gln | Asp |
| **H54** | Ct-1 | 1 | 45.2 | Met | Leu | Ile | Ala | Leu | Ala | Asn | His | Arg | Ser | Gln | Asn | Leu | Phe | Ala | Arg | Leu | Tyr | Ala | Arg | Ser | Ile | Ser | Asn | Ala | Pro | Lys | Gly |
| **H55** | Voeran-1 | 1 | 45.5 | Leu | Leu | Ile | Ala | Leu | Ala | Asn | His | Arg | Ser | Gln | Asn | His | Leu | Val | Arg | Leu | His | Ala | Arg | Ser | Ile | Ser | Asn | Thr | Gln | Gln | Asp |
| **H56** | Bla-1 | 1 | 45.6 | Met | Leu | Ile | Ala | Leu | Ala | Asn | His | Arg | Ser | Gln | Asp | Leu | Leu | Ala | Arg | Leu | Tyr | Ala | Arg | Ser | Ile | Arg | Lys | Ala | Gln | Gln | Gly |
| **H57** | Kondara | 1 | 45 | Leu | Leu | Ile | Ala | Leu | Ala | Asn | His | Arg | Thr | Gln | Asn | Leu | Phe | Ala | Arg | Pro | His | Ala | Arg | Ser | Ile | Arg | Lys | Ala | Gln | Gln | Asp |
| **H58** | Tscha-1 | 2 | 46.7 ± 2.2 | Leu | Leu | Val | Ala | Ile | Ala | Asn | His | Arg | Thr | Gln | Asn | Leu | Phe | Ala | Arg | Pro | His | Ala | Arg | Tyr | Leu | Arg | Lys | Ala | Gln | Gln | Asp |
| **H59** | HR-5 | 1 | 47.6 | Met | Leu | Ile | Ala | Leu | Ser | Asn | His | Arg | Ser | Gln | Asn | Leu | Phe | Ala | Arg | Pro | His | Ser | Arg | Ser | Ile | Arg | Lys | Ala | Gln | Gln | Asp |
| **H60** | Na-1 | 1 | 48.1 | Leu | Leu | Ile | Ala | Leu | Ser | Asn | His | Arg | Ser | Gln | Asn | Leu | Phe | Ala | Arg | Leu | Tyr | Ala | Arg | Ser | Ile | Arg | Lys | Ala | Gln | Lys | Gly |
| **H61** | Yo-0 | 2 | 48.9 ± 2.4 | Met | Leu | Ile | Ala | Leu | Ala | Asn | His | Arg | Ser | Gln | Asp | Leu | Leu | Val | Arg | Pro | His | Ala | Lys | Ser | Ile | Arg | Lys | Ala | Gln | Gln | Asp |
| **H62** | Com-1 | 1 | 50.4 | Leu | Leu | Val | Ala | Leu | Ser | Asn | His | Arg | Ser | Gln | Asn | Leu | Phe | Ala | Arg | Leu | Tyr | Ala | Arg | Ser | Ile | Arg | Lys | Ala | Gln | Gln | Gly |
| **H63** | Lecho-1 | 1 | 50.7 | Leu | Leu | Val | Ala | Leu | Ala | Asn | His | Arg | Ser | Gln | Asp | Leu | Leu | Val | Arg | Pro | His | Ala | Lys | Ser | Ile | Arg | Lys | Thr | Gln | Gln | Asp |
| **H64** | Ms-0 | 1 | 51.2 | Leu | Leu | Val | Ala | Leu | Ala | Asn | His | Arg | Ser | Gln | Asp | His | Leu | Val | Arg | Leu | Tyr | Ala | Arg | Ser | Ile | Ser | Asn | Ala | Pro | Lys | Gly |
| **H65** | stepn-2 | 1 | 51.4 | Leu | Phe | Ile | Ala | Leu | Ser | Asn | His | Arg | Thr | Gln | Asn | His | Leu | Val | Arg | Pro | His | Ala | Lys | Ser | Ile | Arg | Lys | Thr | Pro | Gln | Gly |
| **H66** | Mnz-0 | 1 | 51.5 | Leu | Leu | Val | Ala | Leu | Ser | Asn | His | Arg | Ser | Gln | Asn | Leu | Phe | Ala | Arg | Leu | His | Ala | Arg | Ser | Ile | Arg | Lys | Ala | Gln | Gln | Gly |
| **H67** | En-1 | 1 | 53.2 | Leu | Leu | Val | Ala | Leu | Ala | Asn | His | Arg | Ser | Gln | Asn | Leu | Phe | Ala | Arg | Leu | Tyr | Ala | Arg | Ser | Ile | Ser | Asn | Ala | Pro | Lys | Gly |
| **H68** | Pna-17 | 1 | 53.5 | Leu | Leu | Val | Ala | Leu | Ala | Asn | His | Arg | Ser | Gln | Asp | His | Phe | Ala | Arg | Pro | His | Ala | Lys | Ser | Ile | Arg | Lys | Thr | Gln | Gln | Asp |
| **H69** | TueV-13 | 2 | 53 ± 11.5 | Leu | Leu | Ile | Ala | Leu | Ala | Asn | His | Arg | Ser | Gln | Asn | His | Leu | Val | Arg | Leu | Tyr | Ala | Arg | Ser | Ile | Arg | Lys | Ala | Gln | Gln | Gly |

| | | | | | | | | | | | | | | | | | | | | | | | | | | | | | | | |
|---|---|---|---|---|---|---|---|---|---|---|---|---|---|---|---|---|---|---|---|---|---|---|---|---|---|---|---|---|---|---|---|
| **H70** | Bozen-1 | 1 | 54.4 | Met | Leu | Ile | Ala | Leu | Ala | Asn | His | Arg | Ser | Gln | Asp | Leu | Phe | Val | Arg | Leu | Tyr | Ala | Arg | Ser | Ile | Arg | Lys | Ala | Gln | Gln | Gly |
| **H71** | rou-0 | 1 | 54.5 | Leu | Leu | Val | Ala | Leu | Ser | Asn | His | Arg | Ser | Gln | Asp | His | Phe | Val | Arg | Leu | Tyr | Ala | Arg | Ser | Ile | Arg | Lys | Ala | Gln | Gln | Gly |
| **H72** | Mz-0 | 1 | 55.2 | Met | Leu | Ile | Ala | Leu | Ala | Asn | His | Arg | Ser | Gln | Asn | Leu | Phe | Ala | Arg | Leu | Tyr | Ala | Arg | Ser | Ile | Arg | Lys | Ala | Gln | Gln | Asp |
| **H73** | Pla-0 | 1 | 55 | Met | Leu | Ile | Ala | Leu | Ala | Asn | His | Arg | Ser | Gln | Asp | His | Leu | Val | Arg | Leu | Tyr | Ala | Arg | Ser | Ile | Arg | Lys | Ala | Gln | Gln | Gly |
| **H74** | Chat-1 | 2 | 56.2 ± 14 | Leu | Leu | Val | Ala | Leu | Ala | Asn | His | Arg | Ser | Gln | Asn | Leu | Phe | Ala | Arg | Leu | His | Ala | Arg | Ser | Ile | Arg | Lys | Ala | Gln | Gln | Gly |
| **H75** | Ha-0 | 4 | 57.9 ± 4.3 | Leu | Leu | Val | Ala | Leu | Ala | Asn | His | Arg | Ser | Gln | Asn | Leu | Phe | Ala | Arg | Leu | Tyr | Ala | Arg | Ser | Ile | Arg | Lys | Ala | Gln | Gln | Gly |
| **H76** | Sij-4 | 1 | 60.9 | Leu | Leu | Ile | Ala | Leu | Ala | Asn | His | Arg | Thr | Gln | Asn | His | Phe | Ala | Arg | Pro | His | Ala | Arg | Ser | Ile | Arg | Lys | Ala | Gln | Gln | Asp |
| **H77** | Gie-0 | 1 | 61.3 | Leu | Leu | Val | Ala | Leu | Ser | Asn | His | Arg | Ser | Gln | Asn | His | Leu | Val | Arg | Leu | Tyr | Ala | Arg | Ser | Ile | Ser | Asn | Ala | Pro | Lys | Gly |
| **H78** | Fei-0 | 1 | 63.4 | Leu | Leu | Val | Ala | Leu | Ser | Asn | His | Arg | Ser | Gln | Asp | His | Leu | Val | Arg | Pro | Tyr | Ala | Arg | Ser | Ile | Arg | Lys | Thr | Gln | Gln | Asp |
| **H79** | Bor-1 | 1 | 63.5 | Leu | Leu | Val | Ala | Leu | Ser | Asn | His | Arg | Ser | Gln | Asn | Leu | Phe | Ala | Arg | Leu | Tyr | Ala | Arg | Ser | Ile | Arg | Lys | Thr | Gln | Gln | Gly |
| **H80** | UK-1 | 1 | 65.2 | Leu | Leu | Val | Ala | Leu | Ala | Asn | His | Arg | Thr | Gln | Asn | Leu | Phe | Ala | Arg | Pro | His | Ala | Arg | Ser | Ile | Arg | Lys | Ala | Gln | Gln | Asp |
| **H81** | Castelfed-4 | 1 | 66.8 | Leu | Leu | Ile | Ala | Leu | Ser | Asn | His | Arg | Ser | Gln | Asn | His | Leu | Val | Arg | Leu | Tyr | Ala | Arg | Ser | Ile | Arg | Lys | Ala | Gln | Gln | Asp |
| **H82** | Gu-0 | 1 | 76.1 | Met | Leu | Ile | Ala | Leu | Ala | Asn | His | Arg | Ser | Gln | Asn | His | Phe | Ala | Arg | Leu | His | Ala | Arg | Ser | Ile | Arg | Lys | Thr | Gln | Gln | Gly |
| | | | | | **Non-polar→Polar** | | | | **Polar→Non-polar** | **Polar→Non-polar** | **Positive→Uncharged** | **frameshift** | | **frameshift** | **Negative→Uncharged** | **Positive→Uncharged** | **Aliphatic→Aromatic** | | **Positive→Uncharged** | **Polar→Non-polar** | **Positive→Non-polar** | **Non-polar→Polar** | | **Polar→Non-polar** | | **Positive→Uncharged** | **Positive→Uncharged** | **Polar→Non-polar** | | **Uncharged→Positive** | **Negative→Uncharged** |

| | | | | | | | 800-902 PH (PLECKS | | | | | | | | | | | | | | | | | | | | | | | | | | | | | | | | | | |
|---|---|---|---|---|---|---|---|---|---|---|---|---|---|---|---|---|---|---|---|---|---|---|---|---|---|---|---|---|---|---|---|---|---|---|---|---|---|---|---|---|---|
| **AT4G17140.1 (*VPS13B*)** | | | | **221** | **327** | **448** | **766** | **940** | **940** | **1086** | **1143** | **1524** | **1536** | **1890** | **1914** | **2015** | **2171** | **2229** | **2350** | **2378** | **2378** | **2509** | **2510** | **2511** | **2620** | **2623** | **2635** | **2717** | **2822** | **2881** | **2985** | **3222** | **3228** | **3231** | **3464** | **3508** | **3758** | **3947** | **3996** | **4170** | **4210** |
| **H1** | El-0 | 3 | 5.3 ± 4.1 | Asp | Pro | Gly | Gly | Val | Gly | Ser | Thr | Glu | Ala | Glu | Ser | Leu | Asn | Ala | Ser | His | His | Leu | Val | Asp | Val | Glu | Val | Met | Lys | Glu | Thr | Asn | His | His | Gln | Asn | Leu | Ile | Ile | His | Gly |
| **H2** | Sij-2 | 1 | 12.9 | Asp | Pro | Gly | Gly | Val | Gly | Ser | Ser | Glu | Ala | Glu | Ser | Leu | Asn | Ala | Ser | His | His | Leu | Val | Asp | Val | Glu | Val | Met | Lys | Glu | Thr | Asn | His | Tyr | Glu | Asn | Leu | Ile | Lys | His | Gly |
| **H3** | Ru3.1-31 | 1 | 18.3 | Asp | Pro | Gly | Gly | Val | Gly | Ser | Ser | Glu | Ala | Glu | Phe | Leu | Asn | Ala | Ser | His | His | Leu | Val | Asp | Val | Glu | Val | Met | Lys | Glu | Thr | Asn | His | His | Glu | Asn | Leu | Val | Lys | His | Gly |
| **H4** | Ba-1 | 1 | 21.2 | Asp | Pro | Gly | Gly | Val | Gly | Ser | Ser | Glu | Ala | Glu | Phe | Leu | Asn | Ala | Ser | His | His | Leu | Val | Asp | Val | Glu | Val | Met | Lys | Glu | Ile | Asn | His | His | Glu | Asn | Leu | Val | Lys | His | Gly |
| **H5** | Tha-1 | 1 | 22.2 | Asp | Pro | Gly | Gly | Val | Gly | Ser | Ser | Glu | Ala | Glu | Ser | Leu | Asn | Ala | Ser | His | His | Leu | Val | Asp | Val | Glu | Val | Met | Lys | Glu | Thr | Asn | His | His | Glu | Asn | Leu | Val | Lys | Gln | Gly |
| **H6** | Bsch-0 | 1 | 23.3 | Asp | Pro | Gly | Gly | Val | Gly | Ser | Ser | Glu | Ala | Glu | Ser | His | Asp | Ala | Ser | His | His | Leu | Val | Asp | Ala | Val | Val | Met | Lys | Glu | Thr | Asn | His | His | Glu | Asn | Leu | Val | Lys | His | Gly |
| **H7** | Rovero-1 | 1 | 24.3 | Ala | Pro | Gly | Gly | Val | Gly | Ser | Ser | Glu | Ala | Glu | Phe | Leu | Asn | Ala | Ser | His | His | Leu | Val | Asp | Val | Glu | Val | Met | Lys | Glu | Thr | Asn | His | His | Glu | Tyr | Leu | Val | Lys | His | Gly |
| **H8** | Pt-0 | 1 | 25.3 | Ala | Pro | Gly | Gly | Val | Gly | Ser | Ser | Glu | Ala | Lys | Ser | Leu | Asn | Gly | Ser | His | His | Leu | Val | Asp | Val | Glu | Val | Met | Lys | Glu | Thr | Asn | His | His | Glu | Tyr | Leu | Val | Lys | His | Gly |
| **H9** | Uod-1 | 1 | 25.7 | Asp | Pro | Gly | Gly | Val | Gly | Ser | Thr | Glu | Ala | Glu | Ser | Leu | Asp | Ala | Ser | His | His | Leu | Val | Asp | Val | Glu | Asp | Met | Lys | Glu | Thr | Asn | His | His | Gln | Asn | Leu | Ile | Ile | His | Gly |
| **H10** | Pn-0 | 1 | 28.5 | Ala | Pro | Gly | Gly | Val | Gly | Ser | Ser | Glu | Ala | Glu | Ser | Leu | Asn | Ala | Ser | His | His | Leu | Val | Asp | Val | Glu | Val | Met | Lys | Lys | Thr | Asn | His | His | Glu | Tyr | Leu | Val | Lys | His | Gly |
| **H11** | Slavi-1 | 1 | 29.5 | Asp | Pro | Gly | Gly | Val | Gly | Ser | Ser | Glu | Ala | Glu | Ser | Leu | Asn | Ala | Ser | His | His | Leu | Val | Asp | Val | Glu | Val | Ile | Lys | Glu | Thr | Asn | His | Tyr | Glu | Asn | Leu | Val | Lys | His | Gly |
| **H12** | Is-0 | 3 | 29.6 ± 7.5 | Asp | Pro | Gly | Gly | Val | Gly | Ser | Ser | Glu | Ala | Glu | Ser | Leu | Asn | Ala | Ser | Arg | His | Leu | Val | Asp | Val | Glu | Val | Met | Lys | Glu | Thr | Asn | His | His | Glu | Asn | Leu | Val | Lys | His | Gly |
| **H13** | Se-0 | 1 | 29.9 | Asp | Pro | Gly | Gly | Val | Gly | Ser | Ser | Glu | Ala | Glu | Ser | Leu | Asn | Ala | Ser | His | His | Leu | Val | Asp | Val | Glu | Val | Met | Lys | Glu | Thr | Asn | His | His | Gln | Asn | Leu | Ile | Ile | His | Gly |
| **H14** | Baa-1 | 3 | 30.8 ± 4.6 | Ala | Pro | Gly | Gly | Val | Gly | Ser | Ser | Asp | Ala | Glu | Ser | Leu | Asn | Ala | Ser | His | His | Leu | Val | Asp | Val | Glu | Val | Met | Lys | Glu | Thr | Asn | His | His | Glu | Tyr | Leu | Val | Lys | His | Gly |
| **H15** | Star-8 | 2 | 31 ± 3.8 | Asp | Pro | Gly | Gly | Val | Gly | Ser | Ser | Glu | Ala | Glu | Ser | Leu | Asn | Ala | Ser | His | His | Leu | Val | Asp | Val | Glu | Val | Ile | Thr | Glu | Thr | Asn | His | His | Glu | Asn | Leu | Val | Lys | His | Gly |
| **H16** | Hh-0 | 2 | 31.5 ± 11.1 | Ala | Pro | Glu | Gly | Val | Gly | Ser | Ser | Glu | Ala | Glu | Ser | Leu | Asn | Ala | Ser | His | His | Leu | Val | Asp | Val | Glu | Val | Met | Lys | Lys | Thr | Asn | His | His | Glu | Tyr | Leu | Val | Lys | His | Gly |
| **H17** | Ciste-2 | 1 | 31.7 | Ala | Pro | Gly | Gly | Val | Gly | Ser | Ser | Glu | Ala | Glu | Ser | Leu | Asn | Gly | Ser | His | His | Leu | Val | Asn | Val | Glu | Val | Met | Lys | Glu | Thr | Asn | His | Tyr | Glu | Tyr | Leu | Val | Lys | His | Gly |
| **H18** | Wt-5 | 1 | 33 | Asp | Pro | Gly | Gly | Val | Gly | Ser | Ser | Glu | Ala | Glu | Phe | Leu | Asp | Ala | Ser | His | Asn | Leu | Val | Asp | Val | Glu | Val | Met | Lys | Glu | Ile | Met | Thr | Met | | | | | | | |
| **H19** | Me-0 | 1 | 33.2 | Asp | Pro | Gly | Gly | Val | Gly | Ser | Thr | Glu | Ala | Glu | Phe | Leu | Asn | Ala | Ser | His | His | Leu | Val | Asp | Val | Glu | Val | Met | Lys | Glu | Thr | Asn | His | His | Gln | Asn | Leu | Ile | Ile | His | Gly |
| **H20** | Sij-1 | 1 | 33.3 | Asp | Pro | Gly | Gly | Gly | Gly | Ser | Ser | Glu | Ala | Lys | Ser | Leu | Asn | Ala | Ser | His | His | Leu | Val | Asp | Val | Glu | Val | Met | Lys | Glu | Thr | Asn | His | His | Glu | Asn | Leu | Ile | Lys | His | Gly |
| **H21** | Sp-0 | 1 | 33.9 | Asp | Pro | Gly | Gly | Val | Gly | Ser | Thr | Glu | Ala | Lys | Phe | Leu | Asn | Ala | Ser | His | His | Leu | Val | Asp | Val | Glu | Val | Met | Lys | Glu | Thr | Asn | His | His | Gln | Asn | Leu | Ile | Ile | His | Gly |
| **H22** | Np-0 | 1 | 33.9 | Asp | Pro | Gly | Gly | Val | Gly | Asn | Ser | Glu | Ala | Glu | Tyr | Leu | Asn | Ala | Ser | His | His | Leu | Val | Asp | Val | Glu | Val | Met | Lys | Glu | Thr | Asn | Leu | His | Glu | Asn | Leu | Val | Lys | His | Gly |
| **H23** | Sg-1 | 1 | 34 | Asp | Pro | Gly | Gly | Val | Phe | Ser | Ser | Glu | Ala | Glu | Ser | Leu | Asp | Ala | Ser | His | His | Leu | Val | Asp | Val | Glu | Val | Met | Lys | Glu | Thr | Asn | His | His | Glu | Asn | Leu | Val | Lys | His | Gly |
| **H24** | Ra-0 | 2 | 34.1 ± 1.8 | Asp | Pro | Gly | Gly | Val | Gly | Ser | Ser | Glu | Ala | Glu | Ser | Leu | Asn | Ala | Ser | His | His | Leu | Val | Asp | Val | Val | Val | Met | Lys | Glu | Thr | Asn | His | His | Glu | Asn | Leu | Val | Lys | His | Gly |
| **H25** | Bolin-1 | 1 | 34.5 | Asp | Pro | Gly | Gly | Val | Gly | Ser | Ser | Glu | Ala | Glu | Tyr | Leu | Asn | Ala | Ser | His | His | Val | Val | Asp | Val | Glu | Val | Met | Lys | Glu | Thr | Asn | Leu | His | Glu | Asn | Val | Val | Lys | His | Gly |
| **H26** | Vezzano-2 | 1 | 34.7 | Asp | Pro | Gly | Gly | Val | Gly | Ser | Ser | Glu | Ala | Glu | Ser | Leu | Asn | Ala | Ser | His | His | Leu | Val | Asp | Val | Glu | Val | Ile | Lys | Glu | Thr | Asn | His | His | Glu | Asn | Leu | Val | Lys | His | Gly |
| **H27** | Borsk-2 | 1 | 35.6 | Ala | Pro | Glu | Gly | Gly | Gly | Ser | Ser | Glu | Ala | Glu | Ser | Leu | Asn | Ala | Ser | His | His | Leu | Val | Asp | Val | Glu | Val | Met | Lys | Lys | Thr | Asn | His | His | Glu | Asn | Leu | Val | Lys | His | Gly |
| **H28** | Ts-1 | 1 | 35.7 | Asp | Pro | Gly | Gly | Val | Gly | Ser | Ser | Glu | Ala | Glu | Ser | Leu | Asn | Ala | Ser | His | His | Leu | Val | Asp | Val | Glu | Val | Ile | Lys | Glu | Thr | Asn | His | His | Glu | Asn | Leu | Val | Lys | His | Gly |
| **H29** | Aa-0 | 7 | 35.7 ± 4.2 | Asp | Pro | Gly | Gly | Val | Gly | Ser | Ser | Glu | Ala | Glu | Ser | Leu | Asn | Ala | Ser | His | His | Leu | Val | Asp | Val | Glu | Val | Met | Lys | Glu | Thr | Asn | His | His | Glu | Asn | Leu | Val | Lys | His | Gly |
| **H30** | Petro-1 | 1 | 35.9 | Asp | Pro | Gly | Gly | Val | Gly | Ser | Ser | Glu | Ala | Glu | Phe | Leu | Asp | Ala | Ser | His | Asn | Leu | Val | Asp | Val | Glu | Val | Met | Lys | Glu | Ile | Asn | His | His | Glu | Asn | Leu | Val | Lys | His | Gly |
| **H31** | Sorbo | 3 | 36.7 ± 6.3 | Asp | Pro | Gly | Gly | Val | Gly | Ser | Ser | Glu | Ala | Glu | Ser | Leu | Asn | Ala | Ser | His | His | Leu | Val | Asp | Val | Glu | Val | Met | Lys | Glu | Thr | Asn | His | His | Glu | Asn | Leu | Val | Lys | His | Gly |
| **H32** | Tu-SB30-3 | 1 | 36.9 | Asp | Pro | Gly | Gly | Val | Gly | Ser | Ser | Glu | Ala | Glu | Phe | Leu | Asn | Ala | Ser | His | Asn | Leu | Val | Asp | Val | Glu | Val | Met | Lys | Glu | Thr | Asn | His | His | Glu | Asn | Leu | Val | Lys | His | Gly |

| | | | | | | | | | | | | | | | | | | | | | | | | | | | | | | | | | | | | | | | | | |
|---|---|---|---|---|---|---|---|---|---|---|---|---|---|---|---|---|---|---|---|---|---|---|---|---|---|---|---|---|---|---|---|---|---|---|---|---|---|---|---|---|---|
| **H32** | Tu-SB30-3 | 1 | 36.9 | Asp | Pro | Gly | Gly | Val | Gly | Ser | Ser | Glu | Ala | Glu | Phe | Leu | Asn | Ala | Ser | His | Asn | Leu | Val | Asp | Val | Glu | Val | Met | Lys | Glu | Thr | Asn | His | His | Glu | Asn | Leu | Val | Lys | His | Gly |
| **H33** | Rennes-1 | 4 | 36.9 ± 6.5 | Asp | Pro | Gly | Gly | Val | Gly | Ser | Thr | Glu | Ala | Glu | Ser | Leu | Asn | Ala | Ser | His | His | Leu | Val | Asp | Val | Glu | Asp | Met | Lys | Glu | Thr | Asn | His | His | Gln | Asn | Leu | Ile | Ile | His | Gly |
| **H34** | Wl-0 | 1 | 38 | Asp | Pro | Gly | Gly | Val | Gly | Ser | Ser | Glu | Ala | Glu | Ser | Leu | Asn | Ala | Ser | His | His | Leu | Val | Asp | Ala | Val | Val | Met | Lys | Glu | Thr | Asn | His | His | Glu | Asn | Leu | Val | Lys | His | Gly |
| **H35** | Ga-0 | 1 | 39.4 | Asp | Pro | Gly | Gly | Val | Phe | Ser | Ser | Glu | Ala | Glu | Ser | Leu | Asn | Ala | Ser | His | His | Leu | Val | Asp | Val | Glu | Val | Met | Lys | Glu | Thr | Asn | His | His | Glu | Asn | Leu | Val | Lys | His | Gly |
| **H36** | Lag2-2 | 1 | 40 | Asp | Pro | Gly | Gly | Val | Gly | Ser | Ser | Glu | Ala | Lys | Ser | Leu | Asn | Ala | Ser | His | His | Leu | Val | Asp | Val | Glu | Val | Ile | Lys | Glu | Thr | Asn | His | His | Glu | Asn | Leu | Val | Lys | His | Gly |
| **H37** | Si-0 | 5 | 40.1 ± 5.4 | Asp | Pro | Gly | Gly | Val | Gly | Ser | Ser | Glu | Ala | Glu | Ser | His | Asn | Ala | Ser | His | His | Leu | Val | Asp | Ala | Val | Val | Met | Lys | Glu | Thr | Asn | His | His | Glu | Asn | Leu | Val | Lys | His | Gly |
| **H38** | Aitba 2 | 1 | 41.1 | Asp | Pro | Gly | Gly | Gly | Gly | Ser | Ser | Glu | Ala | Glu | Ser | Leu | Asn | Ala | Ser | His | His | Leu | Val | Asp | Val | Glu | Val | Met | Lys | Glu | Thr | Asn | Leu | His | Glu | Asn | Leu | Val | Lys | His | Gly |
| **H39** | TueV-13 | 1 | 41.5 | Asp | Pro | Gly | Ser | Val | Gly | Ser | Ser | Glu | Ala | Glu | Phe | Leu | Asp | Ala | Ser | His | His | Leu | Val | Asp | Val | Glu | Val | Met | Lys | Glu | Thr | Asn | His | His | Glu | Asn | Leu | Val | Lys | His | Gly |
| **H40** | Do-0 | 8 | 41.9 ± 4.9 | Ala | Pro | Gly | Gly | Val | Gly | Ser | Ser | Glu | Ala | Glu | Ser | Leu | Asn | Ala | Ser | His | His | Leu | Val | Asp | Val | Glu | Val | Met | Lys | Glu | Thr | Asn | His | His | Glu | Tyr | Leu | Val | Lys | His | Gly |
| **H41** | Niel-2 | 1 | 42.1 | Asp | Pro | Gly | Gly | Gly | Gly | Ser | Ser | Glu | Ala | Glu | Phe | Leu | Asp | Ala | Ser | His | His | Leu | Val | Asp | Val | Glu | Val | Met | Lys | Glu | Thr | Asn | His | His | Glu | Asn | Leu | Val | Lys | Gln | Gly |
| **H42** | Sei-0 | 5 | 42.2 ± 5.5 | Asp | Pro | Gly | Gly | Val | Phe | Ser | Ser | Glu | Ala | Glu | Ser | Leu | Asn | Ala | Ser | His | His | Leu | Val | Asp | Val | Glu | Val | Met | Lys | Glu | Thr | Asn | His | His | Glu | Asn | Leu | Val | Lys | His | Gly |
| **H43** | Timpo-1 | 2 | 42.6 ± 11.2 | Asp | Pro | Gly | Gly | Val | Gly | Ser | Ser | Glu | Ala | Glu | Ser | Leu | Asp | Ala | Ser | His | His | Leu | Val | Asp | Val | Glu | Val | Met | Lys | Glu | Thr | Asn | His | His | Glu | Asn | Leu | Val | Lys | His | Gly |
| **H44** | Jl-3 | 1 | 43.2 | Asp | Pro | Gly | Gly | Val | Gly | Ser | Thr | Glu | Ala | Glu | Ser | Leu | Asp | Ala | Asn | His | His | Leu | Val | Asp | Val | Glu | Val | Met | Lys | Glu | Thr | Asn | His | His | Gln | Asn | Leu | Ile | Ile | His | Gly |
| **H45** | Kelsterbach- | 2 | 43.7 ± 1.7 | Asp | Pro | Gly | Gly | Val | Gly | Ser | Ser | Glu | Ala | Glu | Ser | Leu | Asp | Ala | Ser | His | His | Leu | Val | Asp | Val | Glu | Val | Met | Lys | Glu | Thr | Asn | His | His | Glu | Asn | Leu | Val | Lys | His | Gly |
| **H46** | Apost-1 | 1 | 44 | Ala | Pro | Gly | Gly | Val | Gly | Ser | Ser | Glu | Ala | Glu | Ser | Leu | Asn | Ala | Ser | His | His | Leu | Val | Asp | Val | Glu | Val | Met | Lys | Glu | Thr | Asn | His | Tyr | Glu | Tyr | Leu | Val | Lys | His | Gly |
| **H47** | Kz-9 | 8 | 44.6 ± 3.5 | Asp | Pro | Gly | Gly | Val | Gly | Ser | Ser | Glu | Ala | Glu | Phe | Leu | Asn | Ala | Ser | His | Asn | Leu | Val | Asp | Val | Glu | Val | Met | Lys | Glu | Ile | Asn | His | His | Glu | Asn | Leu | Val | Lys | His | Gly |
| **H48** | Buckhorn | 8 | 44.7 ± 5.2 | Asp | Pro | Gly | Gly | Val | Gly | Ser | Ser | Glu | Ala | Glu | Ser | Leu | Asn | Ala | Ser | His | His | Leu | Ile | Asp | Val | Glu | Val | Met | Lys | Glu | Thr | Asn | His | His | Glu | Asn | Leu | Val | Lys | His | Gly |
| **H49** | Je-0 | 13 | 44.7 ± 3.3 | Asp | Pro | Gly | Gly | Gly | Gly | Ser | Ser | Glu | Thr | Lys | Ser | Leu | Asp | Ala | Ser | His | His | Leu | Val | Asp | Val | Glu | Val | Met | Lys | Glu | Thr | Asn | His | Tyr | Glu | Asn | Leu | Val | Lys | His | Gly |
| **H50** | Hn-0 | 5 | 45.2 ± 1.8 | Asp | Pro | Gly | Gly | Val | Gly | Asn | Ser | Glu | Ala | Glu | Tyr | Leu | Asn | Ala | Ser | His | His | Leu | Val | Asp | Val | Glu | Val | Met | Lys | Glu | Thr | Asn | Leu | His | Glu | Asn | Val | Val | Lys | His | Gly |
| **H51** | CIBC-5 | 3 | 45.4 ± 13.1 | Ala | Pro | Gly | Gly | Val | Gly | Ser | Ser | Glu | Ala | Glu | Ser | Leu | Asn | Gly | Ser | His | His | Leu | Val | Asn | Val | Glu | Val | Met | Lys | Glu | Thr | Asn | His | His | Glu | Tyr | Leu | Val | Lys | His | Gly |
| **H52** | Voeran-1 | 1 | 45.5 | Asp | Pro | Gly | Gly | Val | Gly | Ser | Ser | Glu | Ala | Glu | Phe | Leu | Asp | Ala | Ser | His | His | Leu | Val | Asp | Val | Glu | Val | Met | Lys | Glu | Thr | Asn | His | His | Glu | Asn | Leu | Val | Lys | His | Gly |
| **H53** | Mammo-1 | 1 | 45.8 | Asp | Pro | Gly | Gly | Val | Gly | Ser | Ser | Glu | Ala | Glu | Tyr | Leu | Asn | Ala | Ser | His | His | Leu | Val | Asp | Val | Glu | Val | Met | Lys | Glu | Thr | Asn | Leu | His | Glu | Asn | Val | Val | Lys | His | Gly |
| **H54** | Alst-1 | 4 | 45.8 ± 8 | Ala | Ser | Gly | Gly | Val | Gly | Ser | Ser | Asp | Ala | Glu | Ser | Leu | Asn | Ala | Ser | His | His | Leu | Val | Asp | Val | Glu | Val | Met | Lys | Glu | Thr | Asn | His | His | Glu | Tyr | Leu | Val | Lys | His | Gly |
| **H55** | Ciste-1 | 1 | 45.9 | Asp | Pro | Gly | Gly | Val | Gly | Ser | Ser | Glu | Thr | Glu | Ser | Leu | Asn | Ala | Ser | His | His | Leu | Val | Asp | Val | Glu | Val | Met | Lys | Glu | Thr | Asn | His | His | Glu | Tyr | Leu | Val | Lys | His | Gly |
| **H56** | Kas-2 | 1 | 47.2 | Asp | Pro | Gly | Gly | Val | Gly | Ser | Ser | Glu | Ala | Glu | Ser | Leu | Asn | Ala | Ser | His | His | Leu | Val | Asp | Val | Glu | Val | Met | Ile | Glu | Thr | Asn | His | His | Glu | Asn | Leu | Val | Lys | His | Gly |
| **H57** | Bd-0 | 2 | 47.5 ± 2.7 | Asp | Pro | Gly | Gly | Val | Gly | Ser | Thr | Glu | Ala | Glu | Ser | Leu | Asn | Ala | Asn | His | His | Leu | Val | Asp | Val | Glu | Val | Met | Lys | Glu | Thr | Asn | His | His | Gln | Asn | Leu | Ile | Ile | His | Gly |
| **H58** | Rsch-4 | 3 | 48.4 ± 8.4 | Asp | Pro | Gly | Gly | Val | Gly | Ser | Ser | Glu | Ala | Glu | Ser | Leu | Asn | Ala | Ser | His | Asn | Leu | Val | Asp | Val | Glu | Val | Met | Lys | Glu | Ile | Asn | His | His | Glu | Asn | Leu | Val | Lys | His | Arg |
| **H59** | Altenb-2 | 1 | 48.8 | Asp | Pro | Gly | Gly | Val | Gly | Asn | Ser | Glu | Ala | Glu | Phe | Leu | Asn | Ala | Ser | His | His | Leu | Val | Asp | Val | Glu | Val | Met | Lys | Glu | Thr | Asn | Leu | His | Glu | Asn | Val | Val | Lys | His | Gly |
| **H60** | Ws-2 | 4 | 48.9 ± 4 | Asp | Pro | Gly | Ser | Val | Gly | Ser | Ser | Glu | Ala | Glu | Ser | Leu | Asn | Ala | Ser | His | His | Leu | Val | Asp | Val | Glu | Val | Met | Lys | Glu | Thr | Asn | His | His | Glu | Asn | Leu | Val | Lys | His | Gly |
| **H61** | Ove-0 | 3 | 49.6 ± 4.6 | Ala | Pro | Gly | Gly | Val | Gly | Ser | Ser | Glu | Ala | Glu | Ser | Leu | Asn | Gly | Ser | His | His | Leu | Val | Asp | Val | Glu | Val | Met | Lys | Glu | Thr | Asn | His | His | Glu | Tyr | Leu | Val | Lys | His | Gly |
| **H62** | Col-0 | 5 | 52.6 ± 3.5 | Asp | Pro | Gly | Gly | Gly | Gly | Ser | Ser | Glu | Thr | Lys | Phe | Leu | Asp | Ala | Ser | His | His | Leu | Val | Asp | Val | Glu | Val | Ile | Lys | Glu | Thr | Asn | His | Tyr | Glu | Asn | Leu | Val | Lys | Gln | Gly |
| **H63** | Rou-0 | 1 | 54.5 | Asp | Pro | Gly | Gly | Gly | Gly | Ser | Ser | Glu | Thr | Lys | Phe | Leu | Asp | Ala | Ser | His | His | Leu | Val | Asp | Val | Glu | Val | Met | Lys | Glu | Thr | Asn | His | Tyr | Glu | Asn | Leu | Val | Lys | His | Gly |
| **H64** | Sij-4 | 1 | 60.9 | Asp | Pro | Gly | Gly | Val | Gly | Ser | Ser | Glu | Ala | Glu | Ser | Leu | Asn | Ala | Ser | His | His | Leu | Val | Asp | Val | Glu | Val | Met | Lys | Glu | Thr | Asn | His | His | Glu | Asn | Leu | Ile | Lys | His | Gly |
| **H65** | UK-1 | 1 | 65.2 | Asp | Pro | Gly | Gly | Val | Gly | Ser | Ser | Glu | Ala | Glu | Ser | Leu | Asn | Ala | Ser | His | His | Leu | Val | Asp | Val | Glu | Val | Met | Thr | Glu | Thr | Asn | His | His | Glu | Asn | Leu | Val | Lys | His | Gly |
| **H66** | Castelfed-4 | 1 | 66.8 | Asp | Pro | Gly | Gly | Gly | Gly | Ser | Ser | Glu | Thr | Lys | Phe | Leu | Asp | Ala | Ser | His | His | Leu | Val | Asp | Val | Glu | Val | Met | Lys | Glu | Thr | Asn | His | Tyr | Glu | Asn | Leu | Val | Lys | Gln | Gly |
| **H67** | Ema-1 | 1 | 70.1 | Asp | Pro | Gly | Gly | Val | Gly | Ser | Ser | Glu | Ala | Glu | Ser | Leu | Asn | Gly | Ser | His | His | Leu | Val | Asp | Val | Glu | Val | Met | Lys | Glu | Thr | Asn | His | His | Glu | Tyr | Leu | Val | Lys | His | Gly |
| | | | | **Negative→Uncharged** | | **Uncharged→Negative** | **Non-polar→Polar** | | **Aliphatic→Aromat** | | | | **Non-polar→Polar** | **Negative→Positive** | **Non-polar→Polar** | **Uncharged→Positi** | **Negative→Unchar** | | | | **Positive→Uncharge** | | | **Negative→Unchar** | | **Negative→Unchar** | **Uncharged→Negat** | | **Positive→Uncharge** | **Negative→Positive** | **Polar→Non-polar** | **frameshift** | **Positive→Uncharge** | **Non-polar→Polar** | **Negative→Unchar** | **Polar→Non-polar** | | | **Positive→Uncharge** | **Positive→Uncharge** | **Uncharged→Positi** |

| AT5G57200.1 (*PICALM2A*) | | | | 211 | 322 | 367 | 403 | 431 | 439 | 494 | 505 | 534 | 538 | 538 | 553 | 584 | 586 |
|---|---|---|---|---|---|---|---|---|---|---|---|---|---|---|---|---|---|
| **H1** | Si-0 | 1 | 23.1 | Tyr | Gln | Ile | Ala | Lys | Asn | Gly | Gly | Met | Gln | Gln | Phe | Gln | Ser |
| **H2** | Sorbo | 1 | 26.6 | Tyr | Gln | Ile | Ala | Glu | Asn | Gly | Gly | Met | Gln | Gln | Phe | Gln | Ser |
| **H3** | Ciste-2 | 1 | 31.7 | Tyr | Gln | Ile | Ala | Glu | Asn | Pro | Met | Arg | Ser | Ser | Ser | Tyr | Pro |
| **H4** | Pog-0 | 1 | 34.7 | Tyr | Gln | Ile | Ala | Lys | Asn | Gly | Gly | Met | Gln | Gln | Phe | Gln | Ser |
| **H5** | Aa-0 | 5 | 37 ± 6.3 | Tyr | His | Ile | Ala | Glu | Asn | Pro | Met | Arg | Ile | Met | Ser | His | His |
| **H6** | Lago-1 | 2 | 39.3 ± 12.2 | Tyr | Gln | Ile | Ala | Glu | Asn | Pro | Met | Met | Ser | Ser | Ser | Tyr | Pro |
| **H7** | Col-0 | 105 | 39.4 ± 1.2 | Tyr | Gln | Ile | Ser | Glu | Asn | Ala | Val | Met | Ser | Ser | Ser | Tyr | Pro |
| **H8** | Utrecht | 1 | 40.1 | Tyr | Gln | Ile | Ser | Lys | Asn | Pro | Met | Met | Ile | Met | Pro | Tyr | Pro |
| **H9** | Timpo-1 | 2 | 41 ± 6.6 | Tyr | Gln | Ile | Ser | Glu | Asn | Pro | Met | Arg | Ser | Ser | Ser | Tyr | Pro |
| **H10** | Se-0 | 3 | 41.1 ± 5.7 | Tyr | Gln | Thr | Ser | Glu | Asn | Ala | Val | Met | Ser | Ser | Ser | Tyr | Pro |
| **H11** | Old-1 | 4 | 41.3 ± 5.8 | Tyr | Gln | Ile | Ser | Glu | Asn | Pro | Met | Met | Ile | Met | Ser | His | Pro |
| **H12** | Je-0 | 6 | 43.9 ± 4.3 | Tyr | Gln | Ile | Ala | Glu | Asn | Pro | Met | Met | Ile | Met | Ser | His | His |
| **H13** | Apost-1 | 1 | 44 | Tyr | His | Ile | Ala | Glu | Asn | Pro | Met | Arg | Ile | Met | Ser | Tyr | Pro |
| **H14** | Bd-0 | 1 | 44.9 | Tyr | Gln | Ile | Ser | Lys | Asn | Pro | Met | Met | Ile | Met | Ser | Tyr | Pro |
| **H15** | Ciste-1 | 1 | 45.9 | Tyr | Gln | Ile | Ser | Glu | Asn | Pro | Met | Arg | Ser | Arg | Ser | His | Pro |
| **H16** | HR-5 | 1 | 47.6 | Tyr | His | Ile | Ala | Glu | Asn | Pro | Met | Arg | Ile | Met | Ser | His | Pro |
| **H17** | Altenb-2 | 1 | 48.8 | Tyr | Gln | Ile | Ser | Lys | Asn | Ala | Val | Met | Ser | Ser | Ser | Tyr | Pro |
| **H18** | Ll-0 | 1 | 50.6 | Tyr | Gln | Ile | Ser | Glu | Asn | Pro | Met | Met | Ile | Met | Ser | His | His |
| **H19** | Ang-0 | 2 | 51.6 ± 2.9 | Tyr | Gln | Ile | Ala | Lys | Asn | Pro | Met | Met | Ile | Met | Pro | His | His |
| **H20** | Mer-6 | 1 | 53.7 | Tyr | Gln | Ile | Ala | Glu | Asn | Pro | Met | Arg | Ile | Ser | Ser | His | Pro |
| **H21** | Bolin-1 | 3 | 54.7 ± 10.2 | Tyr | Gln | Ile | Ala | Glu | Asn | Pro | Met | Met | Ile | Met | Ser | His | Pro |
| **H22** | Ei-2 | 1 | 55.2 | Tyr | Gln | Ile | Ser | Glu | Asn | Pro | Met | Arg | Ile | Met | Ser | His | His |
| **H23** | Jm-0 | 3 | 55.9 ± 3.9 | Ser | Gln | Ile | Ser | Glu | Asn | Ala | Val | Met | Ser | Ser | Ser | Tyr | Pro |
| **H24** | Gie-0 | 1 | 61.3 | Tyr | His | Ile | Ser | Glu | Asn | Pro | Met | Arg | Ile | Met | Ser | Tyr | Pro |
| **H25** | Rou-0 | 2 | 65.3 ± 10.8 | Tyr | His | Ile | Ala | Glu | Asn | Gly | Gly | Met | Gln | Gln | Phe | Gln | Ser |
| | | | | **Non-polar →Polar** | **Uncharged→ Positive** | **Non-polar →Polar** | **Polar →Non-polar** | **Negative→Positive** | Insertion | **Non-polar →Polar** | | **Uncharged→ Positive** | **Polar →Non-polar** | **Polar →Non-polar** | | **Non-polar →Positive** | **Uncharged→ Positive** |

| AT5G57210.1 | | | | | | | | | | | | | | | 28-375 Rab-Gap TBC | |
|---|---|---|---|---|---|---|---|---|---|---|---|---|---|---|---|---|
| | | | | 731 | 721 | 681 | 677 | 599 | 554 | 547 | 531 | 516 | 491 | 460 | 375 | 7 |
| **H1** | Lago-1 | 1 | 27.1 | Asn | Gly | Pro | Ser | Glu | Pro | Lys | Lys | Met | Glu | Arg | Tyr | Ile |
| **H2** | Timpo-1 | 1 | 31.4 | Asn | Gly | Pro | Ser | Glu | His | Lys | Lys | Met | Glu | Arg | Tyr | Thr |
| **H3** | Ciste-2 | 1 | 31.7 | Asn | Gly | Pro | Ser | Glu | His | Asn | Arg | Leu | Glu | Arg | Tyr | Ile |
| **H4** | Bolin-1 | 1 | 34.5 | Asn | Gly | Pro | Ser | Glu | His | Lys | Lys | Met | Val | Arg | Tyr | Ile |
| **H5** | Aa-0 | 4 | 35 ± 7.6 | Asn | Gly | Pro | Ser | Glu | Pro | Asn | Arg | Met | Val | Arg | Tyr | Ile |
| **H6** | Si-0 | 3 | 37.4 ± 9.2 | Asn | Gly | Leu | Ser | Glu | Pro | Asn | Lys | Met | Val | His | Tyr | Ile |
| **H7** | Col-0 | 23 | 38.2 ± 2.6 | Asn | Glu | Pro | Ser | Glu | His | Lys | Lys | Leu | Glu | Arg | Asp | Thr |
| **H8** | Sorbo | 4 | 38.6 ± 6.3 | Asn | Ala | Gly | Thr | Glu | Pro | Asn | Lys | Met | Val | Arg | Tyr | Ile |
| **H9** | Ru3.1-31 | 46 | 39 ± 1.8 | Asn | Glu | Pro | Ser | Glu | Pro | Lys | Lys | Leu | Glu | Arg | Asp | Ile |
| **H10** | Utrecht | 1 | 40.1 | Asn | Ala | Gly | Thr | Glu | Pro | Asn | Lys | Met | Val | His | Asp | Ile |
| **H11** | Buckhorn Pass | 17 | 40.8 ± 2.9 | Ile | Glu | Pro | Ser | Glu | Pro | Lys | Lys | Leu | Glu | Arg | Asp | Thr |
| **H12** | Sij-2 | 26 | 42.5 ± 2.4 | Asn | Glu | Pro | Ser | Glu | Pro | Lys | Lys | Leu | Glu | Arg | Asp | Thr |
| **H13** | Apost-1 | 1 | 44 | Asn | Gly | Pro | Ser | Glu | His | Asn | Arg | Met | Val | Arg | Tyr | Ile |
| **H14** | Bd-0 | 1 | 44.9 | Asn | Ala | Gly | Thr | Glu | Pro | Asn | Lys | Met | Val | His | Tyr | Ile |
| **H15** | Je-0 | 4 | 44.9 ± 6.1 | Asn | Ala | Gly | Thr | Asp | Pro | Asn | Lys | Met | Val | Arg | Tyr | Ile |
| **H16** | Bla-1 | 1 | 45.6 | Asn | Gly | Pro | Ser | Glu | Pro | Asn | Lys | Met | Val | Arg | Asp | Ile |
| **H17** | Ciste-1 | 2 | 48.3 ± 2.5 | Asn | Gly | Pro | Ser | Glu | His | Lys | Arg | Met | Val | Arg | Tyr | Ile |
| **H18** | Ang-0 | 1 | 48.8 | Asn | Ala | Gly | Thr | Glu | Pro | Asn | Lys | Met | Val | His | Tyr | Ile |
| **H19** | Com-1 | 1 | 50.4 | Asn | Gly | Pro | Ser | Glu | Pro | Lys | Arg | Met | Val | Arg | Tyr | Ile |
| **H20** | Ak-1 | 2 | 50.4 ± 14.9 | Asn | Gly | Pro | Ser | Asp | Pro | Asn | Lys | Met | Val | Arg | Tyr | Ile |
| **H21** | stepn-2 | 1 | 51.4 | Asn | Gly | Pro | Ser | Glu | Pro | Asn | Lys | Leu | Val | Arg | Tyr | Ile |
| **H22** | Ts-1 | 3 | 51.7 ± 8.5 | Asn | Gly | Pro | Ser | Glu | Pro | Asn | Lys | Met | Val | Arg | Tyr | Ile |
| **H23** | Mer-6 | 1 | 53.7 | Asn | Gly | Pro | Ser | Glu | His | Asn | Arg | Met | Glu | Arg | Tyr | Ile |
| **H24** | rou-0 | 1 | 54.5 | Asn | Ala | Gly | Thr | Glu | Pro | Lys | Arg | Met | Val | Arg | Tyr | Ile |
| **H25** | Chat-1 | 3 | 57.9 ± 9.9 | Asn | Ala | Gly | Thr | Glu | Pro | Asn | Arg | Met | Val | Arg | Tyr | Ile |
| **H26** | Gie-0 | 1 | 61.3 | Asn | Ala | Gly | Thr | Glu | His | Asn | Arg | Met | Val | Arg | Tyr | Ile |
| | | | | **Polar→Non-polar** | **Negative→Uncharged** | **Polar→Non-polar** | **Deletion** | | **Positive→Uncharged** | **Positive→Uncharged** | | | **Negative→Uncharged** | | **Polar→Non-polar** | **Polar→Non-polar** |

**Table S6.** Amino acid haplotypes in genes associated with the top 10 most-significant SNPs in the GWAS for high pH tolerance in *Arabidopsis thaliana*. The variants were analyzed with the denser SNP information for 151 out of the 218 ecotypes available in the 1001 genomes database, and those genes having amino acid polymorphisms significantly associated with the phenotype in a local association study (See Table 3) are presented. The relative root length (RRL; %) for ecotypes containing a particular haplotype and the soil pH (at 10 cm depth) of the native geographical location of the ecotype are shown as averages with standard errors. The effects of the polymorphisms, viz., drastic changes in the protein (frameshift, stop gain, and stop loss) and moderate effects like changes in the chemical nature of amino acids, are indicated below. Asterisks indicate a stop codon. Grey cells indicate deviations from the reference haplotype (Col-0). The polymorphism positions with significant associations in the local association study (LAS) are highlighted in pink and functional domains by yellow.

#### Table S7. Genes differentially expressed under high pH stress in *Arabidopsis thaliana*

| GeneID | $log_2FC$ | *FDR* | Gene | Protein names |
|---|---|---|---|---|
| AT4G36700 | 6.40 | 1.51E-37 | *AP22.80* | Vicilin-like seed storage protein At4g36700 (Globulin At4g36700) |
| AT3G12900 | 6.00 | 3.68E-14 | *S8H* | Scopoletin 8-hydroxylase (2-oxoglutarate-dependent dioxygenase S8H) |
| AT2G41810 | 5.90 | 2.90E-14 | *AXX17_At2g39090* | hypothetical protein |
| AT1G07373 | 5.50 | 9.46E-27 | | |
| AT1G07367 | 5.10 | 5.89E-29 | | |
| AT2G41240 | 5.00 | 1.96E-09 | *BHLH100* | Transcription factor bHLH100 (bHLH transcription factor bHLH100) |
| AT1G13608 | 4.70 | 4.40E-18 | *F13B4* | Putative defensin-like protein 288 |
| AT2G43890 | 4.60 | 1.73E-06 | *AXX17_At2g41410* | hypothetical protein |
| AT3G56970 | 4.50 | 1.16E-18 | *ORG2* | Transcription factor ORG2 (bHLH transcription factor bHLH038) |
| AT5G02780 | 4.50 | 7.71E-10 | *GSTL1* | Glutathione S-transferase L1 (GST class-lambda member 1) |
| AT3G56980 | 4.40 | 3.82E-15 | *ORG3* | Transcription factor ORG3 (bHLH transcription factor bHLH039) |
| AT3G58060 | 4.40 | 1.26E-07 | *MTPC3* | Putative metal tolerance protein C3 (AtMTP8) |
| AT1G47400 | 4.30 | 2.79E-37 | *FEP3* | Uncharacterized protein |
| AT3G61930 | 4.30 | 2.29E-09 | *AXX17_At3g56230* | hypothetical protein |
| AT3G47720 | 4.10 | 5.79E-07 | *SRO4* | Probable inactive poly [ADP-ribose] polymerase SRO4 (Protein SIMILAR TO RCD ONE 4) |
| AT2G14247 | 4.10 | 1.25E-10 | *AN1_LOCUS7816* | hypothetical protein |
| AT1G73120 | 4.00 | 1.13E-07 | *AXX17_At1g67320* | Uncharacterized protein |
| AT1G13609 | 3.90 | 4.73E-05 | | Defensin-like family protein |
| AT1G12030 | 3.90 | 1.26E-16 | *AXX17_At1g12410* | Uncharacterized protein |
| AT1G24580 | 3.80 | 1.17E-17 | *AXX17_At1g25830* | RING-type domain-containing protein |
| AT2G30766 | 3.70 | 2.60E-10 | *FEP1* | Uncharacterized protein |
| AT4G37420 | 3.70 | 6.00E-05 | *AT4G37420* | Glycosyltransferase family 92 protein (EC 2.4.1.-) |
| AT5G38820 | 3.60 | 1.32E-04 | *AN1_LOCUS24042* | hypothetical protein |
| AT3G29970 | 3.60 | 1.21E-05 | | B12D protein |
| AT1G01580 | 3.50 | 2.39E-06 | *FRO2* | Ferric reduction oxidase 2 |
| AT5G57540 | 3.40 | 6.00E-05 | *XTH13* | Putative xyloglucan endotransglucosylase/hydrolase protein 13 (EC 2.4.1.207) |
| AT4G33070 | 3.40 | 8.87E-16 | *PDC1* | Pyruvate decarboxylase 1 (EC 4.1.1.1) |
| AT2G19210 | 3.40 | 5.72E-06 | *F27F23.1* | Putative leucine-rich repeat receptor-like protein kinase At2g19210 (EC 2.7.11.1) |
| AT1G47395 | 3.40 | 1.79E-27 | *AXX17_At1g41360* | Uncharacterized protein |
| AT1G12805 | 3.40 | 1.40E-05 | *AN1_LOCUS1389* | Uncharacterized protein |

| | | | | |
|---|---|---|---|---|
| AT5G43360 | 3.30 | 1.78E-02 | *PHT1-3* | Probable inorganic phosphate transporter 1-3 (H(+)/Pi cotransporter) |
| AT1G62420 | 3.30 | 5.58E-06 | *AXX17_At1g55720* | hypothetical protein |
| AT4G19690 | 3.10 | 7.35E-08 | *IRT1* | Fetransport protein 1 transport protein 1) (Iron-regulated transporter 1) |
| AT2G17850 | 3.10 | 5.05E-05 | *T13L16.13* | Rhodanese/Cell cycle control phosphatase superfamily protein |
| AT1G77120 | 3.10 | 1.17E-29 | *ADH1* | Alcohol dehydrogenase class-P (EC 1.1.1.1) |
| AT2G38240 | 3.00 | 4.27E-02 | *JOX4* | Jasmonate-induced oxygenase 4 (Jasmonic acid oxidase 4) |
| AT4G19680 | 3.00 | 3.36E-02 | *IRT2* | Fetransport protein 2 transport protein 2) (Iron-regulated transporter 2) |
| AT5G04150 | 3.00 | 5.25E-05 | *BHLH101* | Basic helix-loop-helix DNA-binding superfamily protein |
| AT5G05250 | 3.00 | 5.37E-33 | *K18I23.5* | AT5g05250/K18I23_5 |
| AT3G13610 | 2.90 | 4.79E-08 | *F6'H1* | Feruloyl CoA ortho-hydroxylase 1 (EC 1.14.11.61) |
| AT3G46900 | 2.90 | 5.25E-07 | *COPT2* | Copper transporter 2 (AtCOPT2) |
| AT5G53450 | 2.80 | 4.67E-35 | *ORG1* | OBP3-responsive protein 1 |
| AT3G12820 | 2.80 | 1.45E-07 | *MYB10* | Myb domain protein 10 |
| AT3G07720 | 2.80 | 4.32E-10 | *F17A17.6* | thiohydroximate-O-sulfate sulfate/sulfur-lyase (EC 4.8.1.5) |
| AT2G20030 | 2.80 | 2.07E-04 | *ATL12* | Putative RING-H2 finger protein ATL12 (RING-type E3 ubiquitin transferase ATL12) |
| AT1G05650 | 2.80 | 3.01E-03 | *F3F20.10* | F3F20.10 protein (Pectin lyase-like superfamily protein) |
| AT3G45060 | 2.60 | 2.75E-05 | *NRT2.6* | High affinity nitrate transporter 2.6 (AtNRT2:6) |
| AT5G01060 | 2.50 | 4.55E-04 | *BSK10* | Serine/threonine-protein kinase BSK (Brassinosteroid-signaling kinase) |
| AT1G01520 | 2.50 | 7.74E-08 | *ASG4* | Homeodomain-like superfamily protein |
| AT3G43190 | 2.30 | 3.82E-13 | *SUS4* | Sucrose synthase 4 (Sucrose-UDP glucosyltransferase 4) |
| AT3G02480 | 2.30 | 3.99E-06 | *AXX17_At3g01700* | Stress-induced protein KIN2-like |
| AT2G44798 | 2.30 | 1.10E-03 | | |
| AT1G43800 | 2.20 | 3.07E-09 | *S-ACP-DES6* | Stearoyl-[acyl-carrier-protein] 9-desaturase 6, chloroplastic (Acyl-[acyl-carrier-protein] desaturase 6) |
| AT3G58810 | 2.20 | 1.62E-05 | *MTPA2* | Metal tolerance protein A2 |
| AT4G01630 | 2.20 | 1.38E-02 | *EXPA17* | Putative expansin-A17 (Ath-ExpAlpha-1.13) |
| AT5G20790 | 2.20 | 1.82E-09 | *T1M15.190* | Transmembrane protein |
| AT1G53635 | 2.20 | 4.85E-03 | | Transmembrane protein |
| AT4G27360 | 2.10 | 1.47E-03 | *F27G19.12* | Dynein light chain |
| AT3G53480 | 2.10 | 5.05E-05 | *ABCG37* | ABC transporter G family member 37 (Protein POLAR AUXIN TRANSPORT INHIBITOR SENSITIVE 1) |
| AT4G33020 | 2.00 | 1.46E-02 | *ZIP9* | ZIP metal ion transporter family |
| AT3G60330 | 2.00 | 4.44E-03 | *HA7* | Plasma membrane ATPase (EC 7.1.2.1) |
| AT3G18290 | 2.00 | 2.44E-13 | *BTS* | Zinc finger protein BRUTUS (Protein EMBRYO DEFECTIVE 2454) |
| AT3G47640 | 2.00 | 8.81E-15 | *BHLH47* | Transcription factor bHLH47 (bHLH transcription factor bHLH047) |
| AT1G23020 | 1.90 | 4.05E-06 | *FRO3* | Ferric reduction oxidase 3 |

| | | | | |
|---|---|---|---|---|
| AT1G14185 | 1.90 | 6.16E-03 | *F7A19.27* | At1g14180/F7A19_27 oxidoreductase family protein) |
| AT5G40590 | 1.90 | 3.51E-05 | *C24_LOCUS24054* | DC1 domain-containing protein |
| AT2G26695 | 1.90 | 2.90E-14 | *AXX17_At2g22560* | hypothetical protein |
| AT5G39890 | 1.80 | 2.14E-04 | *PCO2* | cysteine dioxygenase (EC 1.13.11.20) |
| AT3G48450 | 1.80 | 4.54E-03 | *AN1_LOCUS15189* | hypothetical protein |
| AT2G28710 | 1.80 | 4.51E-04 | *AXX17_At2g24820* | hypothetical protein |
| AT1G01570 | 1.80 | 2.83E-03 | *F22L4.11* | Glycosyltransferase (Transferring glycosyl group transferase (DUF604)) |
| AT1G58684 | 1.70 | 1.00E-03 | *RPS2B* | Small ribosomal subunit protein uS5y/uS5u/uS5v (40S ribosomal protein S2-2) |
| AT4G16370 | 1.70 | 5.25E-16 | *OPT3* | Oligopeptide transporter 3 (AtOPT3) |
| AT5G67370 | 1.70 | 9.03E-08 | *CGLD27* | Protein CONSERVED IN THE GREEN LINEAGE AND DIATOMS 27, chloroplastic |
| AT1G74770 | 1.70 | 2.57E-03 | *BTSL1* | Zinc ion binding protein |
| AT5G66740 | 1.70 | 3.96E-04 | *AXX17_At5g66740* | hypothetical protein |
| AT5G13740 | 1.60 | 6.63E-11 | *ZIF1* | Zinc induced facilitator 1 |
| AT1G03470 | 1.60 | 1.33E-10 | *NET3A* | Protein NETWORKED 3A |
| AT5G36890 | 1.60 | 1.75E-04 | *BGLU42* | Beta-glucosidase 42 (EC 3.2.1.21) |
| AT2G16060 | 1.60 | 2.20E-08 | *AHB1* | Anaerobic nitrite reductase AHB1 (Non-symbiotic hemoglobin 1) |
| AT2G19590 | 1.60 | 1.02E-05 | *ACO1* | ACC oxidase 1 |
| AT3G21240 | 1.60 | 4.21E-04 | *4CL2* | 4-coumarate--CoA ligase (EC 6.2.1.12) |
| AT5G03570 | 1.50 | 1.34E-02 | *IREG2* | Solute carrier family 40 member |
| AT5G03545 | 1.50 | 2.46E-02 | *F12E4_330* | At5g03545 (Expressed in response to phosphate starvation protein) |
| AT3G56360 | 1.50 | 5.25E-07 | *AXX17_At3g50990* | Uncharacterized protein |
| AT5G19890 | 1.40 | 1.08E-02 | *PER59* | Peroxidase 59 (Peroxidase N) |
| AT1G56430 | 1.40 | 7.55E-10 | *NAS4* | Probable nicotianamine synthase 4 (S-adenosyl-L-methionine:S-adenosyl-L-methionine:S-adenosyl-methionine 3-amino-3-carboxypropyltransferase 4) |
| AT3G21890 | 1.40 | 1.52E-03 | *MIP1B* | B-box domain protein 31 (Microprotein 1B) |
| AT1G09560 | 1.40 | 7.72E-03 | *GLP4* | Germin-like protein subfamily 2 member 1 |
| AT4G17680 | 1.40 | 6.38E-04 | *DL4875C* | SBP family protein |
| AT2G42750 | 1.40 | 1.12E-09 | *DJC77* | At2g42750/F7D19.25 (Expressed protein) |
| AT4G06195 | 1.40 | 2.01E-02 | | |
| AT1G65870 | 1.30 | 1.61E-03 | *DIR21* | Dirigent protein 21 (AtDIR21) |
| AT5G05270 | 1.30 | 3.61E-04 | *CHI3* | Probable chalcone--flavanone isomerase 3 (Chalcone isomerase-like 1) |
| AT1G64980 | 1.30 | 3.78E-07 | *CDI* | Nucleotide-diphospho-sugar transferases superfamily protein |
| AT2G29350 | 1.20 | 1.35E-02 | *SAG13* | Senescence-associated protein 13 (EC 1.1.1.-) |
| AT2G37040 | 1.20 | 2.49E-04 | *PAL1* | Phenylalanine ammonia-lyase 1 (EC 4.3.1.24) |
| AT3G03270 | 1.20 | 9.49E-08 | *HRU1* | AT3G03270 protein (Adenine nucleotide alpha hydrolases-like superfamily protein) |

| | | | | |
|---|---|---|---|---|
| AT5G24155 | 1.20 | 1.60E-02 | | Squalene monooxygenase (EC 1.14.14.17) |
| AT4G31320 | 1.20 | 1.48E-02 | *AXX17_At4g35850* | SAUR-like auxin-responsive protein family |
| AT1G66100 | 1.20 | 1.12E-03 | *F15E12.20* | Probable thionin-2.4 |
| AT1G55960 | 1.20 | 3.91E-05 | *F14J16.24* | Polyketide cyclase/dehydrase and lipid transport superfamily protein (Putative membrane related protein CP5) |
| AT3G25190 | 1.10 | 4.81E-02 | *VTL5* | Vacuolar iron transporter homolog 2.1 (AtVTL5) |
| AT4G01080 | 1.10 | 2.46E-02 | *TBL26* | Protein trichome birefringence-like 26 |
| AT5G15950 | 1.10 | 3.14E-05 | *SAMDC2* | S-adenosylmethionine decarboxylase proenzyme 2 [Cleaved into: S-adenosylmethionine decarboxylase 2 alpha chain; S-adenosylmethionine decarboxylase 2 beta chain] |
| AT5G23730 | 1.10 | 4.87E-05 | *RUP2* | WD repeat-containing protein RUP2 (Protein REPRESSOR OF UV-B PHOTOMORPHOGENESIS 2) |
| AT5G67330 | 1.10 | 6.41E-05 | *NRAMP4* | Metal transporter Nramp4 (AtNramp4) |
| AT2G40890 | 1.10 | 7.45E-03 | *CYP98A3* | Cytochrome P450 98A3 (C3'H) |
| AT4G25700 | 1.10 | 5.95E-06 | *BETA-OHASE* | Beta-carotene 3-hydroxylase 1, chloroplastic (EC 1.14.15.24) |
| AT2G27420 | 1.10 | 2.04E-02 | *F10A12.10* | Cysteine proteinase (Putative cysteine proteinase) |
| AT3G55710 | 1.00 | 2.27E-03 | *UGT76F2* | UDP-glycosyltransferase 76F2 (EC 2.4.1.-) |
| AT2G38210 | 1.00 | 2.06E-02 | *PDX1L4* | Pyridoxal 5'-phosphate synthase PDX1-like 4 |
| AT1G32900 | 1.00 | 5.74E-05 | *GBSS1* | Granule-bound starch synthase 1, chloroplastic/amyloplastic (GBSS-I) |
| AT4G32480 | 1.00 | 1.64E-04 | *AN1_LOCUS20066* | Sugar phosphate exchanger, (DUF506) |
| AT4G27450 | 1.00 | 2.27E-02 | *AXX17_At4g31580* | hypothetical protein |
| AT3G47420 | 1.00 | 3.81E-05 | *T21L8.170* | Putative glycerol-3-phosphate transporter 1 (AtPS3) |
| AT2G23030 | -1.00 | 1.37E-02 | *SRK2J* | Serine/threonine-protein kinase SRK2J (SnRK2.9) |
| AT1G53910 | -1.00 | 5.05E-05 | *RAP212* | Related to AP2 12 |
| AT1G70860 | -1.00 | 1.29E-02 | *MLP36* | Major latex-like protein (Polyketide cyclase/dehydrase and lipid transport superfamily protein) |
| AT5G37450 | -1.00 | 1.86E-02 | *T25O11.15* | non-specific serine/threonine protein kinase (EC 2.7.11.1) |
| AT2G16660 | -1.00 | 1.54E-02 | *AN1_LOCUS8015* | hypothetical protein |
| AT1G21670 | -1.00 | 4.70E-03 | *AXX17_At1g22750* | hypothetical protein |
| AT4G11890 | -1.00 | 2.97E-02 | *ARCK1* | Protein kinase superfamily protein |
| AT1G54960 | -1.00 | 1.44E-02 | *ANP2* | Mitogen-activated protein kinase kinase kinase 2 (Arabidopsis NPK1-related protein kinase 2) |
| AT1G68600 | -1.00 | 1.51E-02 | *ALMT5* | Aluminum-activated malate transporter 5 (AtALMT5) |
| AT3G47780 | -1.00 | 2.01E-02 | *ABCA7* | ABC transporter A family member 7 (Probable ABC2 homolog 6) |
| AT5G07100 | -1.10 | 1.46E-02 | *WRKY26* | At5g07100 (WRKY DNA-binding protein 26) |
| AT1G17147 | -1.10 | 4.34E-02 | *VQ1* | VQ motif-containing protein 1 (AtVQ1) |
| AT4G02380 | -1.10 | 4.26E-02 | *SAG21* | Senescence-associated gene 21 |
| AT3G17360 | -1.10 | 2.47E-02 | *POK1* | Phragmoplast orienting kinesin 1 |
| AT2G38380 | -1.10 | 1.51E-04 | *PER22* | Peroxidase 22 (Basic peroxidase E) |
| AT4G02390 | -1.10 | 7.64E-03 | *PARP2* | Poly [ADP-ribose] polymerase (EC 2.4.2.-) |

| | | | | |
|---|---|---|---|---|
| AT4G10380 | -1.10 | 1.91E-02 | *NIP5-1* | Probable aquaporin NIP5-1 (Protein NLM6) |
| AT5G41750 | -1.10 | 1.46E-02 | *MUF8.3* | Disease resistance protein family (Disease resistance protein-like) |
| AT5G06800 | -1.10 | 1.27E-02 | *MPH15.16* | Myb-like HTH transcriptional regulator family protein |
| AT2G01530 | -1.10 | 4.54E-03 | *MLP329* | MLP-like protein 329 |
| AT1G67970 | -1.10 | 5.15E-03 | *HSFA8* | Heat stress transcription factor A-8 (HSTF 5) |
| AT1G58290 | -1.10 | 1.72E-03 | *HEMA1* | Glutamyl-tRNA reductase 1, chloroplastic (EC 1.2.1.70) |
| AT3G09270 | -1.10 | 2.63E-03 | *GSTU8* | Glutathione S-transferase U8 (GST class-tau member 8) |
| AT1G02920 | -1.10 | 8.34E-03 | *GSTF7* | Glutathione S-transferase F7 (Glutathione S-transferase 11) |
| AT5G01600 | -1.10 | 1.68E-03 | *FER1* | Ferritin-1, chloroplastic (EC 1.16.3.1) |
| AT3G26210 | -1.10 | 2.03E-02 | *CYP71B23* | Cytochrome P450 71B23 (EC 1.14.-.-) |
| AT1G20620 | -1.10 | 3.50E-03 | *CAT3* | catalase (EC 1.11.1.6) |
| AT5G41761 | -1.10 | 1.83E-02 | *AXX17_At5g39550* | DUF7722 domain-containing protein |
| AT5G04060 | -1.10 | 1.80E-04 | *F21E1.1* | Probable methyltransferase PMT7 (EC 2.1.1.-) |
| AT4G14746 | -1.10 | 1.02E-04 | | Neurogenic locus notch-like protein |
| AT3G50800 | -1.10 | 2.31E-02 | *AXX17_At3g45050* | Uncharacterized protein |
| AT1G65845 | -1.10 | 1.46E-02 | | At1g65844 (Transmembrane protein) |
| AT1G21520 | -1.10 | 5.29E-04 | *AXX17_At1g22580* | hypothetical protein |
| AT2G46530 | -1.10 | 3.69E-03 | *ARF11* | Auxin response factor 11 |
| AT5G24470 | -1.10 | 3.68E-03 | *APRR5* | Two-component response regulator-like APRR5 (Pseudo-response regulator 5) |
| AT4G24120 | -1.20 | 2.65E-05 | *YSL1* | Metal-nicotianamine transporter YSL1 (AtYSL1) |
| AT4G14365 | -1.20 | 4.37E-06 | *XBAT34* | Putative E3 ubiquitin-protein ligase XBAT34 (RING-type E3 ubiquitin transferase XBAT34) |
| AT5G25810 | -1.20 | 1.51E-02 | *TINY* | Ethylene-responsive transcription factor TINY |
| AT3G55240 | -1.20 | 1.10E-02 | *T26I12.120* | Uncharacterized protein At3g55240 (Uncharacterized protein T26I12.120) |
| AT1G65510 | -1.20 | 1.46E-02 | *STMP7* | Transmembrane protein |
| AT5G24270 | -1.20 | 3.09E-02 | *SOS3* | Calcineurin B-like protein |
| AT4G08390 | -1.20 | 1.73E-06 | *SAPX* | L-ascorbate peroxidase (EC 1.11.1.11) |
| AT2G26560 | -1.20 | 2.95E-02 | *PLP2* | Patatin-like protein 2 (AtPLAIIA) |
| AT2G38390 | -1.20 | 5.86E-06 | *PER23* | Peroxidase 23 (ATP34) |
| AT3G16390 | -1.20 | 5.34E-04 | *NSP3* | Thiohydroximate-O-sulfate sulfur/sulfate-lyase NSP3 (AtNSP3) |
| AT1G01010 | -1.20 | 1.35E-02 | *NAC001* | NAC domain-containing protein 1 (Protein NTM1-like 10) |
| AT2G18690 | -1.20 | 2.24E-02 | *MSF3.7* | Transmembrane protein |
| AT2G48130 | -1.20 | 4.35E-04 | *LTPG15* | Non-specific lipid transfer protein GPI-anchored 15 (AtXYP7) |
| AT1G60960 | -1.20 | 1.82E-03 | *IRT3* | Iron regulated transporter 3 |
| AT1G21100 | -1.20 | 3.50E-02 | *IGMT1* | Indole glucosinolate O-methyltransferase 1 (EC 2.1.1.-) |

| | | | | |
|---|---|---|---|---|
| AT3G50460 | -1.20 | 1.68E-02 | *HR2* | RPW8-like protein 2 (AtHR2) |
| AT1G22770 | -1.20 | 2.63E-03 | *GI* | Protein GIGANTEA |
| AT3G01330 | -1.20 | 2.78E-02 | *E2FF* | E2F transcription factor-like E2FF (E2F-like repressor E2L2) |
| AT3G14680 | -1.20 | 4.98E-03 | *CYP72A14* | Cytochrome P450 72A14 (EC 1.14.-.-) |
| AT4G00970 | -1.20 | 2.10E-04 | *CRK41* | Cysteine-rich RLK 41 |
| AT1G30270 | -1.20 | 1.69E-05 | *CIPK23* | non-specific serine/threonine protein kinase (EC 2.7.11.1) |
| AT4G15610 | -1.20 | 1.51E-02 | *CASPL1D1* | CASP-like protein |
| AT1G66280 | -1.20 | 2.43E-02 | *BGLU22* | Beta-glucosidase 22 (EC 3.2.1.21) |
| AT5G51010 | -1.20 | 1.47E-05 | *AXX17_At5g49800* | Rubredoxin-like domain-containing protein |
| AT5G26280 | -1.20 | 8.58E-04 | *F9D12.7* | AT5g26280/T19G15_130 (TRAF-like family protein) |
| AT2G36885 | -1.20 | 2.85E-02 | | Translation initiation factor |
| AT2G30480 | -1.20 | 4.64E-02 | *T6B20.25* | Uncharacterized protein |
| AT1G68650 | -1.20 | 7.46E-04 | *F24J5.11* | GDT1-like protein 5 |
| AT1G61360 | -1.20 | 2.08E-02 | *T1F9.15* | G-type lectin S-receptor-like serine/threonine-protein kinase At1g61360 (EC 2.7.11.1) |
| AT1G17300 | -1.20 | 3.76E-02 | *T13M22.4* | Uncharacterized protein |
| AT1G11210 | -1.20 | 8.12E-03 | *AN1_LOCUS1215* | DUF4408 domain-containing protein |
| AT5G12420 | -1.30 | 2.18E-03 | *WSD7* | O-acyltransferase family protein |
| AT4G01430 | -1.30 | 1.83E-02 | *UMAMIT29* | WAT1-related protein |
| AT1G78000 | -1.30 | 4.13E-08 | *SULTR1* | Sulfate transporter 12 |
| AT1G21120 | -1.30 | 2.02E-03 | *IGMT2* | O-methyltransferase family protein |
| AT1G21310 | -1.30 | 3.61E-02 | *EXT3* | Extensin-3 (Protein ROOT-SHOOT-HYPOCOTYL-DEFECTIVE) |
| AT2G43150 | -1.30 | 4.30E-02 | *EXT21* | Proline-rich extensin-like family protein (Putative extensin) |
| AT2G18660 | -1.30 | 2.04E-02 | *EGC2* | EG45-like domain containing protein 2 (Ath-ExpGamma-1.2) |
| AT1G56300 | -1.30 | 5.89E-06 | *DJC53* | At1g56300 (DnaJ protein, putative) |
| AT5G23190 | -1.30 | 1.44E-02 | *CYP86B1* | Cytochrome P450 86B1 (EC 1.14.-.-) |
| AT5G52750 | -1.30 | 3.50E-03 | *F6N7.24* | Heavy metal transport/detoxification superfamily protein |
| AT5G07475 | -1.30 | 3.85E-02 | *AN1_LOCUS21568* | Phytocyanin domain-containing protein |
| AT4G01130 | -1.30 | 4.31E-02 | *F2N1.17* | GDSL-like Lipase/Acylhydrolase superfamily protein |
| AT1G63570 | -1.30 | 4.81E-02 | *F2K11.7* | Receptor-like protein kinase-related family protein |
| AT5G57560 | -1.40 | 3.63E-03 | *XTH22* | Xyloglucan endotransglucosylase/hydrolase protein 22 (Touch protein 4) |
| AT5G19600 | -1.40 | 5.70E-06 | *SULTR3;5* | Probable sulfate transporter 3.5 |
| AT2G02120 | -1.40 | 8.52E-06 | *PDF2.1* | Defensin-like protein 4 (Plant defensin 2.1) |
| AT1G34670 | -1.40 | 4.39E-02 | *MYB93* | Transcription factor MYB93 (AtMYB93) |
| AT3G11430 | -1.40 | 6.87E-03 | *GPAT5* | Glycerol-3-phosphate acyltransferase 5 (EC 2.3.1.15) |

| | | | | |
|---|---|---|---|---|
| AT1G64380 | -1.40 | 5.30E-04 | *ERF061* | Ethylene-responsive transcription factor ERF061 |
| AT2G25090 | -1.40 | 3.01E-03 | *CIPK16* | CBL-interacting serine/threonine-protein kinase 16 (SOS2-like protein kinase PKS15) |
| AT1G49910 | -1.40 | 1.18E-04 | *BUB3.2* | Mitotic checkpoint protein BUB3.2 (Protein BUDDING UNINHIBITED BY BENZYMIDAZOL 3.2) |
| AT3G57240 | -1.40 | 1.70E-02 | *BG3* | Probable glucan endo-1,3-beta-glucosidase BG3 (AtBG3) |
| AT4G40070 | -1.40 | 4.14E-06 | *ATL32* | RING-H2 finger protein ATL32 (RING-type E3 ubiquitin transferase ATL32) |
| AT5G06570 | -1.40 | 3.45E-05 | *F15M7.10* | Alpha/beta-Hydrolases superfamily protein |
| AT4G14780 | -1.40 | 5.08E-04 | *AXX17_At4g17090* | Protein kinase domain-containing protein |
| AT3G26470 | -1.40 | 4.26E-02 | *AXX17_At3g28720* | RPW8 domain-containing protein |
| AT3G20380 | -1.40 | 2.97E-02 | | TRAF-like family protein |
| AT1G73810 | -1.40 | 2.46E-02 | *AT1G73810* | Core-2/I-branching beta-1,6-N-acetylglucosaminyltransferase family protein (Glycosyltransferase) |
| AT1G15125 | -1.40 | 1.88E-02 | *AN1_LOCUS1641* | hypothetical protein |
| AT3G23170 | -1.50 | 1.26E-04 | *PRP* | At3g23170 |
| AT1G65390 | -1.50 | 1.27E-02 | *PP2-A5* | Phloem protein 2 A5 |
| AT5G24220 | -1.50 | 2.81E-02 | *MOP9.3* | Gb\|AAD29063.1 (Lipase class 3-related protein) |
| AT2G19800 | -1.50 | 8.17E-03 | *MIOX2* | Inositol oxygenase 2 (MI oxygenase 2) |
| AT1G21110 | -1.50 | 2.92E-02 | *IGMT3* | Indole glucosinolate O-methyltransferase 3 (EC 2.1.1.-) |
| AT5G17170 | -1.50 | 4.41E-08 | *ENH1* | Rubredoxin family protein |
| AT5G57220 | -1.50 | 1.46E-05 | *CYP81F2* | Cytochrome P450 81F2 (Protein INDOLE GLUCOSINOLATE MODIFIER 1) |
| AT4G12330 | -1.50 | 4.71E-03 | *CYP706A7* | Cytochrome P450, family 706, subfamily A, polypeptide 7 (Flavonoid 3, 5-hydroxylase like protein) |
| AT5G26010 | -1.50 | 5.76E-06 | *T1N24.8* | Probable protein phosphatase 2C 72 (EC 3.1.3.16) |
| AT4G21250 | -1.50 | 2.65E-02 | *F7J7.190* | Sulfite exporter TauE/SafE family protein 5 |
| AT3G50400 | -1.50 | 3.70E-02 | *F11C1.240* | GDSL esterase/lipase At3g50400 (Extracellular lipase At3g50400) |
| AT2G15220 | -1.50 | 1.79E-05 | *AXX17_At2g10370* | Plant basic secretory protein family protein |
| AT1G77530 | -1.50 | 1.06E-02 | *AT9943_LOCUS5749* | hypothetical protein |
| AT1G74460 | -1.50 | 4.08E-04 | *F1M20.14* | GDSL esterase/lipase At1g74460 (Extracellular lipase At1g74460) |
| AT1G66465 | -1.50 | 1.60E-02 | | Transmembrane protein |
| AT1G66090 | -1.50 | 1.85E-04 | *F15E12.17* | Disease resistance protein (Putative disease resistance protein) |
| AT1G18940 | -1.50 | 3.61E-02 | *AN1_LOCUS2028* | Uncharacterized protein |
| AT1G07550 | -1.50 | 5.53E-03 | *F22G5.7* | Probable LRR receptor-like serine/threonine-protein kinase At1g07550 (EC 2.7.11.1) |
| AT4G23810 | -1.60 | 2.91E-08 | *WRKY53* | Probable WRKY transcription factor 53 (WRKY DNA-binding protein 53) |
| AT2G38470 | -1.60 | 3.81E-04 | *WRKY33* | Probable WRKY transcription factor 33 (WRKY DNA-binding protein 33) |
| AT5G09520 | -1.60 | 6.09E-04 | *PELPK2* | Protein PELPK2 (Protein Pro-Glu-Leu\|Ile\|Val-Pro-Lys 2) |
| AT3G45710 | -1.60 | 2.36E-04 | *NPF2.5* | Protein NRT1/ PTR FAMILY 2.5 (Protein NAXT1-like 5) |
| AT2G01520 | -1.60 | 1.73E-11 | *MLP328* | MLP-like protein 328 |

| | | | | |
|---|---|---|---|---|
| AT5G60760 | -1.60 | 1.85E-07 | *MAE1.1* | P-loop containing nucleoside triphosphate hydrolases superfamily protein |
| AT2G24720 | -1.60 | 1.62E-05 | *GLR2.2* | Glutamate receptor 2.2 (Ligand-gated ion channel 2.2) |
| AT5G41300 | -1.60 | 4.81E-02 | *CRRSP59* | Cysteine-rich repeat secretory protein 59 |
| AT5G59400 | -1.60 | 1.08E-06 | *F2O15.8* | PGR5-like A protein |
| AT5G46900 | -1.60 | 4.46E-02 | *AXX17_At5g45380* | hypothetical protein |
| AT5G38930 | -1.60 | 3.81E-04 | *K15E6.16* | Germin-like protein subfamily 1 member 10 |
| AT5G12270 | -1.60 | 9.77E-03 | *AXX17_At5g12000* | hypothetical protein |
| AT5G09480 | -1.60 | 2.55E-07 | *AXX17_At5g09020* | Uncharacterized protein |
| AT4G20390 | -1.60 | 2.83E-05 | *F9F13.40* | CASP-like protein 1B2 (AtCASPL1B2) |
| AT4G20000 | -1.60 | 2.46E-02 | *AXX17_At4g23470* | hypothetical protein |
| AT2G35380 | -1.60 | 6.36E-05 | *T32F12.24* | peroxidase (EC 1.11.1.7) |
| AT1G78990 | -1.60 | 6.82E-03 | *AXX17_At1g73710* | HXXXD-type acyl-transferase family protein |
| AT1G55990 | -1.60 | 6.69E-03 | *F14J16.30* | Glycine-rich protein |
| AT1G02850 | -1.60 | 4.97E-02 | *AN1_LOCUS322* | Beta-glucosidase |
| AT5G10760 | -1.60 | 8.22E-04 | *AED1* | Aspartyl protease AED1 (Apoplastic EDS1-dependent protein 1) |
| AT4G04770 | -1.60 | 2.49E-12 | *ABCI8* | Iron-sulfur cluster assembly SufBD family protein ABCI8, chloroplastic (Protein LONG AFTER FAR-RED 6) |
| AT5G13580 | -1.60 | 4.99E-03 | *ABCG6* | ABC transporter G family member 6 (AtWBC6) |
| AT1G73300 | -1.70 | 2.52E-03 | *SCPL2* | Serine carboxypeptidase-like 2 (EC 3.4.16.-) |
| AT3G04070 | -1.70 | 1.88E-02 | *NAC047* | NAC domain containing protein 47 (NAM-like protein (No apical meristem)) |
| AT5G38200 | -1.70 | 1.86E-04 | *MXA21.22* | Class I glutamine amidotransferase-like superfamily protein |
| AT5G40510 | -1.70 | 4.15E-04 | *MNF13.5* | AT5g40510/MNF13_30 (Sucrase/ferredoxin-like family protein) |
| AT2G40300 | -1.70 | 4.05E-13 | *FER4* | Ferritin-4, chloroplastic (EC 1.16.3.1) |
| AT4G23220 | -1.70 | 9.21E-03 | *CRK14* | Cysteine-rich receptor-like protein kinase 14 (EC 2.7.11.-) |
| AT4G37010 | -1.70 | 1.30E-04 | *CEN2* | Centrin 2 |
| AT5G49770 | -1.70 | 1.02E-04 | *K2I5.14* | Probable leucine-rich repeat receptor-like protein kinase At5g49770 (EC 2.7.11.1) |
| AT3G45730 | -1.70 | 1.38E-07 | *AN1_LOCUS14900* | Uncharacterized protein |
| AT2G39920 | -1.70 | 2.39E-03 | *AXX17_At2g36910* | hypothetical protein |
| AT2G25297 | -1.70 | 2.10E-02 | | Transmembrane protein |
| AT2G03260 | -1.70 | 8.44E-03 | *T18E12.7* | EXS family protein |
| AT1G05575 | -1.70 | 8.78E-04 | *AXX17_At1g05020* | hypothetical protein |
| AT5G58660 | -1.80 | 2.08E-03 | *MZN1.11* | 2-oxoglutarate and Fe(II)-dependent oxygenase superfamily protein (Gibberellin oxidase-like protein) |
| AT1G70880 | -1.80 | 7.11E-10 | *MLP15* | Major latex-like protein (Putative Csf-2-related protein) |
| AT1G80240 | -1.80 | 3.51E-05 | *DGR1* | Protein DUF642 L-GALACTONO-1,4-LACTONE-RESPONSIVE GENE 1 (DUF642 L-GalL-RESPONSIVE GENE 1) |
| AT5G58860 | -1.80 | 5.96E-07 | *CYP86A1* | Cytochrome P450 86A1 (Protein HYDROXYLASE OF ROOT SUBERIZED TISSUE) |

| | | | | |
|---|---|---|---|---|
| AT3G43670 | -1.80 | 4.62E-16 | *CuAOgamma2* | Amine oxidase [copper-containing] gamma 2 (EC 1.4.3.21) |
| AT2G32210 | -1.80 | 4.98E-04 | *ATHCYSTM6* | Cysteine-rich/transmembrane domain A-like protein |
| AT4G30670 | -1.80 | 5.05E-05 | *AXX17_At4g35160* | Transmembrane protein |
| AT4G18253 | -1.80 | 2.17E-03 | | Receptor Serine/Threonine kinase-like protein |
| AT4G14450 | -1.80 | 3.31E-02 | *dl3265w* | Uncharacterized protein At4g14450, chloroplastic |
| AT3G46270 | -1.80 | 3.90E-03 | | Receptor protein kinase-like protein |
| AT3G13437 | -1.80 | 4.96E-03 | *AXX17_At3g13750* | Transmembrane protein |
| AT3G06390 | -1.80 | 1.31E-11 | *F24P17.14* | CASP-like protein 1D2 (AtCASPL1D2) |
| AT2G20142 | -1.80 | 3.55E-02 | | Toll-Interleukin-Resistance domain family protein |
| AT1G21230 | -1.90 | 2.33E-02 | *WAK5* | Wall-associated receptor kinase 5 (EC 2.7.11.-) |
| AT5G43580 | -1.90 | 5.56E-06 | *UPI* | Serine protease inhibitor, potato inhibitor I-type family protein |
| AT5G55590 | -1.90 | 7.37E-03 | *QRT1* | pectinesterase (EC 3.1.1.11) |
| AT5G09530 | -1.90 | 4.18E-11 | *PELPK1* | Protein PELPK1 (Protein Pro-Glu-Leu\|Ile\|Val-Pro-Lys 1) |
| AT4G13250 | -1.90 | 1.21E-20 | *NYC1* | NAD(P)-binding Rossmann-fold superfamily protein |
| AT3G02240 | -1.90 | 1.31E-02 | *GLV4* | Protein GOLVEN 4 [Cleaved into: GLV4p] |
| AT2G28850 | -1.90 | 1.18E-03 | *CYP710A3* | Cytochrome P450 710A3 (C-22 sterol desaturase) |
| AT4G23230 | -1.90 | 6.39E-03 | *CRK15* | Cysteine-rich receptor-like protein kinase 15 (EC 2.7.11.-) |
| AT5G52790 | -1.90 | 6.54E-04 | *F6N7.28* | CBS domain protein with a domain protein (DUF21) |
| AT5G37690 | -1.90 | 2.53E-02 | *At5g37700* | SGNH hydrolase-type esterase superfamily protein |
| AT5G11140 | -1.90 | 4.97E-02 | *AN1_LOCUS21908* | Phospholipase-like protein family protein |
| AT4G03540 | -1.90 | 4.73E-02 | *F9H3.17* | CASP-like protein 1C1 (AtCASPL1C1) |
| AT2G28270 | -1.90 | 4.34E-02 | *AXX17_At2g24290* | hypothetical protein |
| AT2G23540 | -1.90 | 2.10E-16 | *F26B6.19* | GDSL esterase/lipase At2g23540 (Extracellular lipase At2g23540) |
| AT1G75040 | -1.90 | 6.23E-09 | *F9E10.11* | Pathogenesis-related protein 5 (PR-5) |
| AT5G10230 | -1.90 | 4.52E-02 | *ANNAT7* | Annexin D7 (AnnAt7) |
| AT4G04990 | -2.00 | 2.30E-02 | *T32N4.7* | AT4g04990 protein (T32N4.7 protein) |
| AT2G22920 | -2.00 | 1.54E-02 | *SCPL12* | Serine carboxypeptidase-like 12 |
| AT3G08860 | -2.00 | 1.06E-02 | *PYD4* | PYRIMIDINE 4 |
| AT3G43850 | -2.00 | 2.62E-02 | *O3L2* | Protein OXIDATIVE STRESS 3 LIKE 2 (AtO3L2) |
| AT1G33910 | -2.00 | 3.53E-02 | *IAN5* | Immune-associated nucleotide-binding protein 5 (AIG1-like protein) |
| AT5G66620 | -2.00 | 1.83E-03 | *DAR6* | DA1-related protein 6 |
| AT1G20630 | -2.00 | 1.85E-17 | *CAT1* | Catalase-1 (EC 1.11.1.6) |
| AT3G54065 | -2.00 | 4.43E-02 | | LOW protein: ankyrin repeat protein |
| AT2G31945 | -2.00 | 1.54E-02 | *AN1_LOCUS9616* | Transmembrane protein |

| | | | | |
|---|---|---|---|---|
| AT2G18450 | -2.10 | 2.08E-03 | *SDH1-2* | Succinate dehydrogenase [ubiquinone] flavoprotein subunit 2, mitochondrial (FP) |
| AT2G24710 | -2.10 | 1.81E-04 | *GLR23* | Glutamate receptor |
| AT5G49780 | -2.10 | 7.80E-04 | *K2I5.15* | Leucine-rich repeat protein kinase family protein |
| AT5G49350 | -2.10 | 2.50E-05 | *K7J8.2* | At5g49350 (Glycine-rich protein family) |
| AT5G26300 | -2.10 | 3.11E-02 | *F9D12.5* | TRAF-like family protein |
| AT4G38080 | -2.10 | 2.31E-08 | *AXX17_At4g43410* | hypothetical protein |
| AT4G33610 | -2.10 | 5.72E-04 | *AXX17_At4g38440* | Glycine-rich protein |
| AT2G29110 | -2.20 | 1.54E-02 | *GLR28* | Glutamate receptor |
| AT2G36690 | -2.20 | 1.55E-13 | *GIM2* | 2-oxoglutarate and Fe(II)-dependent oxygenase superfamily protein |
| AT3G44540 | -2.20 | 1.01E-04 | *FAR4* | Fatty acyl-CoA reductase (EC 1.2.1.84) |
| AT4G23140 | -2.20 | 2.19E-05 | *CRK6* | Cysteine-rich receptor-like protein kinase 6 (Receptor-like protein kinase 5) |
| AT1G09240 | -2.30 | 1.19E-09 | *NAS3* | Nicotianamine synthase 3 (AtNAS3) |
| AT1G26240 | -2.30 | 1.14E-02 | *EXT19* | Proline-rich extensin-like family protein |
| AT3G01420 | -2.30 | 5.22E-11 | *DOX1* | Alpha-dioxygenase 1 (Plant alpha dioxygenase 1) |
| AT4G20450 | -2.30 | 2.34E-03 | *F9F13.100* | Leucine-rich repeat protein kinase family protein |
| AT2G43390 | -2.30 | 1.54E-02 | *AXX17_At2g40880* | Uncharacterized protein |
| AT1G09080 | -2.30 | 1.63E-04 | *AXX17_At1g08900* | hypothetical protein (BIP3) |
| AT4G26200 | -2.30 | 2.34E-02 | *ACS7* | 1-aminocyclopropane-1-carboxylate synthase 7 (S-adenosyl-L-methionine methylthioadenosine-lyase 7) |
| AT1G05675 | -2.40 | 1.57E-03 | *UGT74E1* | UDP-glycosyltransferase 74E1 (EC 2.4.1.-) |
| AT5G65980 | -2.40 | 6.86E-09 | *PILS7* | Auxin efflux carrier family protein |
| AT5G24180 | -2.40 | 8.25E-04 | *K12G2.7* | Gb\|AAD29063.1 (Lipase class 3-related protein) |
| AT3G18400 | -2.40 | 1.62E-02 | *AN1_LOCUS13247* | hypothetical protein |
| AT3G25790 | -2.50 | 6.92E-03 | *HHO1* | Myb-like transcription factor family protein |
| AT2G32200 | -2.60 | 2.27E-05 | *ATHCYSTM5* | Cysteine-rich/transmembrane domain A-like protein |
| AT4G25400 | -2.60 | 1.17E-03 | *T30C3.70* | Basic helix-loop-helix DNA-binding superfamily protein |
| AT1G58225 | -2.60 | 1.20E-07 | | Transmembrane protein |
| AT5G40010 | -2.60 | 9.90E-04 | *AATP1* | AAA-ATPase ASD, mitochondrial (Protein ATPASE-IN-SEED-DEVELOPMENT) |
| AT2G37430 | -2.80 | 9.18E-03 | *ZAT11* | Zinc finger protein ZAT11 |
| AT2G22510 | -2.90 | 6.78E-05 | *AXX17_At2g18110* | Hydroxyproline-rich glycoprotein family protein |
| AT5G08250 | -3.00 | 1.29E-03 | *T22D6_190* | Cytochrome P450 superfamily protein (Cytochrome P450-like protein) |
| AT3G49160 | -3.00 | 5.11E-15 | | pyruvate kinase (EC 2.7.1.40) |
| AT4G08380 | -3.10 | 2.50E-03 | *EXT22* | Extensin-like protein (Proline-rich extensin-like family protein) |
| AT2G34317 | -3.10 | 4.91E-05 | | Avirulence induced family protein |
| AT1G24420 | -3.10 | 9.53E-05 | *F21J9.8* | F21J9.8 (HXXXD-type acyl-transferase family protein) |

| | | | | |
|---|---|---|---|---|
| AT5G51720 | -3.20 | 4.06E-16 | *NEET* | CDGSH iron-sulfur domain-containing protein NEET (At-NEET) |
| AT1G78340 | -3.30 | 8.26E-07 | *GSTU22* | Glutathione S-transferase U22 (GST class-tau member 22) |
| AT3G09220 | -3.80 | 9.53E-09 | *LAC7* | Laccase (Urishiol oxidase) |
| AT1G51830 | -4.10 | 7.78E-20 | *SIF1* | Leucine-rich repeat protein kinase family protein |
| AT5G06900 | -4.50 | 2.88E-06 | *AXX17_At5g06540* | CYP93D1 |
| AT5G22890 | -8.00 | 1.49E-12 | *STOP2* | Protein SENSITIVE TO PROTON RHIZOTOXICITY 2 (Zinc finger protein STOP2) |
